# Discovery of single-stranded DNA in meteorite-derived cultures: evidence of novel genetic elements

**DOI:** 10.1101/2025.01.20.633545

**Authors:** Carlos López Ramón y Cajal

**Affiliations:** Head of the Obstetrics and Gynaecology Department, Álvaro Cunqueiro Hospital, University Hospital Complex of Vigo, Spain

**Keywords:** Single-Stranded DNA (ssDNA), Meteorites, New Biological Structure (NBS), Extremophiles, Astrobiology, Prebiotic Chemistry, Genomic Analysis, Biosignatures, Molecular Evolution, Metagenomics

## Abstract

This study presents the first evidence of single-stranded DNA (ssDNA) sequences identified through shotgun metagenomic analysis of meteorite-derived cultures. These ssDNA sequences, predominantly isolated from the "no-hits" zone with no significant matches in existing genomic databases, exhibit conserved motifs, distinct secondary structural features, and a unique AT-rich composition. These characteristics suggest potential roles in autoreplication, molecular stabilization, and resilience under extreme conditions, consistent with extremophilic adaptations. Notably, this discovery aligns with observations of a provisionally termed New Biological Structure (NBS) detected in diverse contexts, including meteorites and biogenic materials, though its characterization lies beyond the scope of this study.

The sequences were detected using a rigorous methodology that included stringent contamination controls, advanced metagenomic assembly techniques, and exhaustive taxonomic filtering to ensure robustness and minimize false positives. Their consistent detection across meteorites of varying types and geographic origins further underscores the reliability of the findings. Repeat analyses revealed specific motifs associated with potential regulatory or structural roles, while secondary structure predictions suggested configurations that could support autoreplication and stability under extreme conditions.

This groundbreaking discovery highlights the potential of meteorites to harbor previously uncharacterized genomic elements, offering profound insights into molecular diversity, prebiotic chemistry, and molecular evolution. By situating these findings within the broader context of extremophilic biology and astrobiological exploration, this study establishes a foundation for investigating the structural, functional, and evolutionary implications of these sequences. Future research will also explore their relevance as biosignatures in the search for life under extreme and extraterrestrial conditions.

## Introduction

In biomedicine, the development of innovative methodologies has revolutionized the study of biological samples, uncovering novel structures that reshape our understanding of molecular and cellular processes. Among these, single-stranded DNA (ssDNA) has garnered significant attention due to its extraordinary resilience and adaptability in extreme environments, including hydrothermal vents, permafrost, and acidic lakes, where it persists despite high radiation, temperature fluctuations, and chemical extremes (Cavicchioli, Siddiqui, Andrews, & Sowers, 2002) (Cowan, y otros, 2024). These unique molecular properties have advanced our understanding of molecular stability and adaptation mechanisms (McKay, 2020) (Pizzarello & Shock, 2010), positioning ssDNA as a model for studying biochemical diversity and the limits of life in extraterrestrial-like environments. By investigating ssDNA’s structural stability and its ability to maintain biological function under inhospitable conditions, researchers can uncover critical insights into the boundaries of molecular evolution and identify biosignatures relevant to astrobiological exploration.

Meteorites, particularly carbonaceous chondrites, are well-known for harboring organic molecules, such as nucleobases and amino acids essential for life (Pizzarello & Shock, 2010) (Sephton, 2002). As analogs of extraterrestrial-like environments, they provide a unique opportunity to explore the genomic and structural properties of ssDNA sequences, shedding light on their potential implications for the emergence of life. The discovery of structured ssDNA sequences in meteorites could reveal novel biological systems capable of adaptation to extreme conditions, contributing to our understanding of molecular evolution and the biochemical potential of extreme environments.

Building on this foundation, our study investigates the New Biological Structure (NBS), a previously uncharacterized biological entity. Initially discovered in terrestrial contexts, such as human amniotic membranes, blood, and biogenic minerals, the NBS consistently appeared across diverse samples, suggesting a robust and adaptable nature. Extending this research to meteorites, we present the first evidence of short ssDNA sequences in meteorite fragments directly associated with the NBS. These sequences, characterized by regions with no significant matches in genomic databases and enriched in conserved motifs, suggest novel mechanisms of adaptation and potential autoreplication.

This study employs advanced microscopy and metagenomic techniques to systematically characterize the NBS and its associated ssDNA. By situating these findings within the broader context of extremophilic biology and astrobiological exploration, we aim to illuminate the molecular diversity and resilience required for life to persist under extreme conditions. These findings contribute to the broader understanding of molecular evolution, prebiotic chemistry, and the potential of meteorites as vectors for prebiotic material or microbial-like entities.

## Material and methods

The descriptive study of the NBS was conducted by the author at the Álvaro Cunqueiro Hospital in Vigo, serving as the basis for genomic analysis. The research project on the NBS was approved by the Ethics Committee for Research of Pontevedra-Vigo-Orense (Registration Codes: 2019/151 and 2019/648).

### DNA Extraction and Culturing of Meteorite Samples

For DNA extraction, meteorite fragments previously evaluated in the descriptive study of the NBS were cultured under controlled conditions. The cultures were performed in sterile Nunc™ EasYFlask™ 25 cm² flasks with Nunclon™ Delta Surface, ensuring an optimal environment for potential growth. Two distinct culture media were employed to assess the adaptability and growth potential of biological entities within the samples.

The first medium was DMEM (Dulbecco’s Modified Eagle’s Medium), sourced from Corning® (500 mL, reference: 10-017-CV). This medium contained 4.5 g/L glucose and L-glutamine but lacked sodium pyruvate. The second medium consisted of sterile distilled water (Water for Injectable Preparations Meinsol®- 10 ml ampoules), which served as a minimalistic medium to evaluate potential adaptability or growth under more stringent conditions. As a negative control, a DNA extraction was performed on the culture medium without meteorite fragments. No detectable DNA was observed in these controls, confirming the absence of contamination during the experimental process.

These protocols were meticulously designed to maintain a sterile and controlled environment, ensuring the reliability of subsequent DNA extraction and analysis of potential biological entities associated with the meteorite samples.

### Preparation of samples for cultures and culture considerations

We cultured two fragments of meteorites over 5 – 7 mm (<1cm) by sample of culture after exhaustive cleaning and disinfection of meteorites with didecyldimethylammonium chloride and non-ionic surfactants, followed by 30 minutes in solution and subsequent treatment with pure bleach (5% sodium hypochlorite). Meteorite samples were collected from meteorites and cultured under controlled laboratory conditions

Meteorites used: a total of seven meteorites were utilized in the study, with details as follows:

— Meteorite 1 (AIQUILE). Type: ordinary chondrite H5. Fall location: Cochabamba, Bolivia. Confirmed fall: November 20, 2016. Status: listed in the Meteoritical Bulletin Database.
— Meteorite 2 (NWA 781). Type: ordinary chondrite LL6. Fall location: Morocco. Fall year: 2001. Status: listed in the Meteoritical Bulletin Database.
— Meteorite 3 (Chelyabinsk). Type: ordinary chondrite LL5. Fall location: Chelyabinskaya oblast’, Russia. Confirmed fall: February 15, 2013. Status: Listed in the Meteoritical Bulletin Database.
— Meteorite 4 (NWA 803). Type: ordinary chondrite L6. Fall location: Northwest Africa—Morocco. Fall year: 2001. Status: Listed in the Meteoritical Bulletin Database.
— Meteorite 5 (NWA). Type: achondrite from the eucrite group. Fall location: Northwest Africa—Algeria. Fall year: 2021. Status: Not listed in the Meteoritical Bulletin Database.
— Meteorite 6 (NWA 869). Type: ordinary chondrite L4-6. Fall location: Northwest Africa—Algeria. Fall year: 2000. Status: Listed in the Meteoritical Bulletin Database.
— Meteorite 7 (Northwest Africa, NWA 12455). Type: carbonaceous chondrite CR7. Fall location: Northwest Africa. Fall year: 2018. Status: Listed in the Meteoritical Bulletin Database.

The meteorite samples used in this study were acquired from Litos, a specialized supplier of meteorites, through their official website https://www.litos.net. This supplier is known for providing verified meteorites from diverse geographical locations and classifications. Details of each meteorite, including type, fall location, and year, were confirmed through the Meteoritical Bulletin Database, where applicable.

Meteorites were selected based on their qualitative compositional richness, prioritizing those classified as ordinary chondrites and carbonaceous chondrites. These types of meteorites are known for their diverse mineral and organic content, providing an optimal framework for exploring potential biological or prebiotic signatures (Sephton, 2002). These meteorites represent a diverse range of compositions and origins, providing a broad spectrum for analysis in the study.

### Culturing procedure

Cultures were performed at room temperature, maintained between 22 and 23 °C. Meteorite fragments were placed into flasks containing the culture medium, and the flasks were vigorously shaken to facilitate the fracturing of the meteorite fragments and the release of potential internal contents. This shaking process was repeated daily at 07:00 hours to ensure consistency.

Rationale for culturing mainly using sterile distilled water. A key hypothesis of this study is that the intrinsic composition of the meteorites themselves may act as a natural culture medium, providing the necessary environmental and chemical conditions for the development of the New Biological Structure (NBS) and the detection of ssDNA sequences. By fracturing the meteorites, the internal material is exposed, releasing potential biologically relevant elements into the sterile distilled water medium. This minimalistic approach aims to allow the endogenous properties of the meteorites to drive the emergence and proliferation of the NBS, minimizing external influence and emphasizing the role of the meteorite’s inherent chemical and structural features.

Additionally, two flasks were subjected to incubation on an Eppendorf ThermoMixer F2.0, set to 56 °C and 150 rpm, operating in two-hour shaking cycles. This procedure was designed to stimulate sample growth, based on observations from previous experiments conducted during the descriptive study of the NBS.

The methodology aimed to alter the medium’s humidity conditions, create currents within the medium through shaking, and induce controlled temperature changes. These factors had been identified in earlier studies as promoting the growth of the NBS. The cultures were maintained within a clean and disinfected cabinet, shielded from light for most of the day. During the shaking periods, however, the samples were exposed to artificial white ambient LED light. This setup was carefully designed to simulate environmental conditions conducive to the growth of the NBS.

Three culture batches were carried out with fragments of meteorites. In the first batch, DMEM was used as the culture medium, while in the second and third batches, distilled water was used, identified as the optimal medium for these cultures. Each batch included three of meteorites (one per distinct meteorite, as specified later).

Observations from dozens of experiments during the descriptive phase of the NBS allowed us to determine that the maximum possible number of structures was produced between 48 hours and 10 days of cultivation. The optimal period was consistently between the 7th and 10th day, so DNA extraction was scheduled accordingly.

### Culturing and DNA extraction summary

Four batches of cultures were performed, with three flasks per batch, resulting in a total of 12 cultures. Each culture underwent two DNA extractions, producing a total of 24 DNA extractions. To ensure a comprehensive sampling strategy across the meteorites, one flask was cultured for each meteorite, with the exception of specific cases. For the Aiquile meteorite, three cultures were performed, while meteorites NWA 781, NWA 803, and NWA 12455 each underwent two cultures. This multi-extraction approach maximized the recovery and subsequent analysis of potential DNA from the samples, enhancing the robustness of the study.

### DNA quantification methodology

All DNA measurements were performed using the Qubit™ Fluorometer, which utilizes assay kits specifically designed for ssDNA and double-stranded DNA (dsDNA). For ssDNA quantification, the Qubit™ ssDNA Assay Kit (Thermo Fisher Scientific, Cat. No. Q10212) was used, offering high accuracy and sensitivity for measuring single-stranded DNA. Similarly, the Qubit™ dsDNA High Sensitivity (HS) Assay Kit (Thermo Fisher Scientific, Cat. No. Q32851) was employed for precise quantification of double-stranded DNA concentrations. These assay kits provide exceptional specificity and reliability, ensuring high-quality data essential for downstream applications.

### Consolidated detailed methodology for shotgun sequencing of ssDNA

This study utilized a shotgun metagenomics approach to sequence single-stranded DNA (ssDNA) from four samples processed in two batches by AllGenetics & Biology SL (www.allgenetics.eu). The methodology is described in detail below.

Sample preparation and library construction. DNA concentrations were quantified using the Qubit High Sensitivity dsDNA Assay (Thermo Fisher Scientific) to ensure accurate input amounts for subsequent steps. Library preparation was performed following the manufacturer’s protocol using the Accel-NGS 1S Plus Library Kit (Swift Bioscience). DNA was randomly fragmented using the dsDNA Fragmentase enzyme cocktail (New England Biolabs). Sequencing adapters were then ligated to the DNA fragment ends, and dual-index barcoding was applied to enable pooling and demultiplexing of the libraries.

Library pooling and sequencing. The constructed libraries were pooled in equimolar amounts, as determined by the Qubit dsDNA High Sensitivity Assay. Sequencing was carried out on the Illumina NovaSeq platform using a PE150 flow cell, which generated paired-end reads with a length of 150 base pairs. The sequencing runs targeted an output of 10 gigabases per run, ensuring sufficient data for downstream analyses.

Quality control and data preprocessing. The quality of the raw sequencing data was initially assessed using FastQC (v0.11.9), and the results were summarized with MultiQC. Extensive data cleaning was performed with Fastp (v0.23.4), which involved the removal of adapter sequences, trimming of low-quality regions with a mean quality score threshold of 15, trimming of 12 base pairs from both ends of each read, and removal of polyG tails as well as reads shorter than 80 base pairs.

Potential PhiX contamination was eliminated using BBDuk, while duplicate reads were filtered with clumplify.sh from the BBTools suite (v39.96). Additionally, sequences aligning to the Homo sapiens GRCh38 reference genome were identified and excluded using Bowtie2 (v2.5.4) to remove human-derived sequences. Following these preprocessing steps, the quality of the cleaned reads was reassessed with FastQC, and the results were consolidated with MultiQC.

### Metagenomic assembly and analysis

The cleaned reads were taxonomically classified using Kraken2 (v2.1.3) with the "PlusPFP" database as the reference. Taxonomic diversity was visualized through interactive graphical representations generated by Krona (v2.7.1). To construct metagenomes, the sequences were assembled using MEGAHIT (v1.2.9) with default parameters, and the assembly quality metrics were calculated using QUAST (v5.0.2).

A specific focus on ssDNA was incorporated into the analysis. Viral sequences within the ssDNA were identified using VirSorter2, while Prodigal was employed to predict open reading frames (ORFs) within the ssDNA contigs, enabling a detailed exploration of their potential functions and characteristics.

### Study of no-hit regions

Non-hit sequences, corresponding to R1 reads that did not yield significant matches in existing databases, were analyzed in-depth to identify potentially novel genomic elements. These sequences were assembled using three assemblers: MEGAHIT (v1.2.9) (Li, Liu, Luo, Sadakane, & Lam, 2015), SPAdes (v3.15.5) (Bankevich A, 2012), and IDBA-UD (v1.1.3-7) (Peng, Leung, Yiu, & Chin, 2012). This multi-assembler approach leveraged the complementary strengths of each tool, ensuring a comprehensive reconstruction of genomic content. The resulting assemblies were screened using BLASTn (Camacho, y otros, 2009) and VirSorter2 (Guo, y otros, 2021) to exclude sequences matching known references, leaving only those that did not yield significant matches in any database.

### Sequence filtering and cleaning

To further ensure the novelty and reliability of the sequences, an exhaustive sequence-cleaning process was applied iteratively. Tools such as BLASTn, BLASTx, Kraken2 (Wood, Lu, & Langmead, 2019), VirSorter2, and BLASTp were employed to remove sequences with potential matches in protein, viral, or taxonomic databases. This stringent cleaning process minimized the risk of contamination and focused on genomic regions unrepresented in known databases, supporting the hypothesis of a non-terrestrial origin. By iteratively removing known sequences, the methodology provided robust evidence for the novelty of the observed sequences.

### Creation of MT_TOTAL and MT_PURE datasets

The non-hit sequences were categorized into two datasets based on the assembly and filtering workflow:

— MT_TOTAL: This dataset was created by assembling all R1 reads from the no-hits zone using MEGAHIT, SPAdes, and IDBA. It represents a comprehensive dataset encompassing all non-hit sequences prior to additional filtering.
— MT_PURE: This dataset was derived by first filtering R1 reads to exclude sequences with BLASTn matches and then assembling the filtered sequences using the same three assemblers. This process enriched the dataset for high-confidence novel genomic content, ensuring it retained only sequences unmatched throughout all filtering steps.

Both datasets underwent additional post-assembly refinement to remove redundancies, ensuring only unique contigs were retained. As a result, MT_TOTAL included 15 high-confidence single-stranded DNA (ssDNA) sequences, while MT_PURE contained 13, both representing strong candidates for novel genomic elements.

### Significance of analyzing no-hit regions

The analysis of no-hit regions broadened the exploratory scope of the study by focusing on sequences that evade existing database matches. These sequences represent a novel opportunity to explore genomic "dark matter," including unique biological entities, poorly studied genomic elements, and novel structural DNA components. By prioritizing uncharacterized regions, the methodology enhanced the depth and originality of the study, contributing significantly to the understanding of molecular evolution and astrobiology.

### Scientific justification

The use of shotgun sequencing of ssDNA offers an unbiased approach to analyzing the entire genetic material within a sample. This methodology is particularly valuable for detecting unculturable or low-abundance microorganisms, which are often overlooked by traditional methods. Its application to environmental samples is essential, as these contexts frequently harbor novel species that remain undetected (Quince et al., 2017).

BLASTn and BLASTx tools play a pivotal role in identifying sequence homology and providing potential functional annotations. Their inclusion in metagenomic workflows ensures a thorough exclusion of known sequences, enabling a focused exploration of novel genetic entities (Camacho et al., 2009). Similarly, VirSorter 2 is instrumental in detecting viral elements within complex datasets, facilitating the differentiation of viral sequences from cellular genomic data (Roux et al., 2021). Kraken 2 further enhances this workflow by offering rapid and accurate taxonomic classification, ensuring the removal of sequences belonging to known taxa (Wood et al., 2019).

The assembly of sequences was conducted using Megahit, a tool optimized for assembling complex metagenomic datasets. Its capacity to recover highly fragmented genomes from environmental samples makes it particularly well-suited for this study (Li et al., 2015).

### Advantages of the MT_PURE approach

The MT_PURE approach offers several advantages for identifying novel sequences and minimizing contamination. By focusing on sequences that lacked BLASTn matches prior to assembly, the MT_PURE set is enriched for uncharacterized genetic material, increasing the likelihood of discovering novel species or genomic elements. The exclusion of sequences with known matches reduces the influence of contaminants, resulting in a cleaner dataset for assembly and downstream analysis.

Additionally, the assembly of only unique sequences enhances the quality of the resulting contigs. This method enables more contiguous and complete reconstructions of novel genomes, which are crucial for subsequent steps such as gene prediction and functional annotation (Li et al., 2015). Overall, the MT_PURE approach provides a robust framework for uncovering uncharacterized biological entities and advancing our understanding of genomic diversity.

This methodology combines advanced sequencing techniques, rigorous sequence assembly, and comprehensive filtering against multiple databases. The MT_TOTAL and MT_PURE sets provide complementary approaches: while MT_TOTAL offers a broader dataset that includes potentially novel but low-confidence sequences, MT_PURE focuses narrowly on the most distinctive genomic content. Together, these approaches form a robust strategy for the identification of novel species or life forms in environmental samples.

## Sequence analysis

Analyses were performed independently for the MT_TOTAL and MT_PURE sequence sets.

Statistical and descriptive analysis. The analysis began with an evaluation of the number of sequences and their length distribution. A histogram was generated to visualize the sequence length distribution for each source, specifically MT_TOTAL and MT_PURE. Additionally, the base composition of the sequences was examined. This included calculating the average percentage of each nucleotide base (A, T, G, C) and determining the ranges for GC and AT content.

Diversity and clustering analysis. To assess diversity and clustering, genetic similarity-based clustering was performed using distance matrices generated with the MEGA11 software (v11.0.13). Dendrograms were constructed to visualize genetic relationships among sequences both within and between sources. Additionally, heatmaps were developed to represent genetic similarities derived from the distance matrix, offering a clear depiction of clustering patterns and sequence relationships.

Identification and analysis of repetitive motifs. Repetitive motifs within the sequences were identified using MEME (Multiple Em for Motif Elicitation) (Bailey TL, 2009), a powerful tool designed to detect conserved sequence patterns across genomic datasets. MEME was chosen for its ability to uncover both common and context-specific motifs, providing insights into sequence functionality. To complement this analysis, FIMO (Find Individual Motif Occurrences) (Bailey TL, 2009)was employed to locate specific instances of these motifs within the datasets, enhancing the resolution of the study. The motifs identified were visualized using WebLogo graphics, offering a clear depiction of nucleotide conservation across the most prevalent motifs. The discovered motifs were subsequently analyzed using TOM TOM (Bailey TL, 2009), a tool for comparing input motifs against known motif databases. This step aimed to identify potential matches with motifs from JASPAR (including CNE, FAM, Phylogenetic, and other collections) (Rauluseviciute, y otros, 2024) and other publicly available databases. The TOM TOM analysis provided a comparative framework to evaluate whether the identified motifs were novel or conserved in known regulatory or structural elements. This approach helped contextualize the motifs in terms of potential functional roles, evolutionary significance, and taxonomic relevance. A comparative analysis was conducted to evaluate shared and unique motifs between the datasets MT_TOTAL and MT_PURE. This comparison focused on motif frequency, distribution, and potential functional implications, aiming to uncover both common regulatory features and exclusive adaptations in each dataset. Statistical metrics such as E-values and p-values were utilized to ensure the robustness of the identified motifs. Parameters for MEME, FIMO, and TOM TOM were optimized to balance sensitivity and specificity, with thresholds tailored to detect biologically relevant patterns. The resulting data were systematically visualized, supporting evolutionary and functional interpretations of the identified motifs.

Secondary structure prediction. Secondary structures of the sequences were predicted using ViennaRNA (RNAfold) (Lorenz, y otros, 2011), focusing on structural elements such as hairpins and loops. A comparative analysis was conducted to evaluate stability and structural complexity, allowing for the identification of differences between sequences from distinct sources. Furthermore, the predicted structures were examined for their potential biological implications, including regulatory, structural, or functional roles.

Functional evaluation. A comprehensive functional analysis was conducted to evaluate the potential roles and biological significance of the sequences. Initial annotation was performed using Prokka (Seemann, 2014), a genome annotation tool, to identify coding sequences (CDS), hypothetical proteins, and potential functional domains such as signal peptides or transmembrane regions. Despite extensive analysis, the majority of sequences lacked known functional annotations, emphasizing their potential novelty. To deepen the understanding of these sequences, complementary analyses were performed. BLASTp searches were conducted to query the translated protein products of the sequences against known protein databases to identify possible homologs or functional relatives; no significant matches were found, highlighting the uniqueness of these sequences. InterProScan (Jones, y otros, 2014) was employed to predict protein family membership, conserved domains, and functional sites. While most sequences returned hypothetical protein annotations, certain domains associated with transmembrane regions and signal peptides were identified, further supporting potential structural or regulatory roles. RNAfold secondary structure predictions were used to analyze the structural properties of the sequences, assessing their stability and potential functional relevance. The MT_PURE dataset exhibited compact and stable configurations, consistent with roles in molecular stabilization or replication. Motif discovery using MEME and FIMO revealed highly conserved motifs, suggesting involvement in replicative or regulatory processes. These motifs were further analyzed for functional significance using motif databases. Finally, phylogenetic analysis was conducted to explore evolutionary relationships through phylogenetic clustering of the MT_TOTAL and MT_PURE datasets, providing insights into their potential conservation and divergence across evolutionary contexts. Together, these analyses provided a multidimensional perspective on the sequences. The lack of significant matches in BLASTp and the hypothetical protein annotations from Prokka and InterProScan highlight their novelty, while the presence of conserved structural features and motifs suggests they may represent uncharacterized elements with replicative or regulatory functions. These findings underscore the need for further experimental validation to explore their roles in replication, autoreplication, or other molecular processes.

This comprehensive analysis highlights the distinct characteristics and potential biological roles of the MT_TOTAL and MT_PURE sequence sets, emphasizing their contribution to the identification of novel genomic features.

## Results

### 1.- Pooling and quantification of DNA for shotgun sequencing

Pooled DNA Sample for Shotgun Sequencing. To achieve a sufficient quantity of DNA for shotgun sequencing, all DNA samples from the 24 extractions were combined into a single pooled sample. The total volume of DNA sent for analysis was 12 µL, with a concentration of 5.64 ng/µL. The composition of the pooled sample included 4.24 ng/µL of single-stranded DNA (ssDNA), accounting for 75.35% of the total DNA, and 1.39 ng/µL of double-stranded DNA (dsDNA), representing 24.65% of the total DNA. This pooling strategy ensured an optimal balance and sufficient quantity of both single-stranded and double-stranded DNA, making the sample well-suited for subsequent shotgun sequencing analysis.

### 2.- Metagenomics analysis of DNA samples

Prior to metagenome assembly, we conducted a preliminary exploration of the taxonomic composition of the cleaned data with Kraken v2.1.3 (Wood, Lu, & Langmead, 2019), using the pre-built “PlusPFP” database, which contains reference genome/protein sequences derived from RefSeq and UniVec for the following groups: archaea, bacteria, viral, plasmid, human, protozoa, fungi, plant, and UniVec_Core (a collection of vector, adapter, linker, and primer sequences that may be contaminating sequencing projects).

The diversity and relative abundance of the hits retrieved by Kraken were graphically represented with the Krona package v2.7.1 (Ondov, Bergman, & Phillippy, 2011). See Figure 1 to both explore and download the Krona pie chart. As shown in this plot, 38 % of the sequences from sample MT were assigned to Homo sapiens.

**Figure 1.**
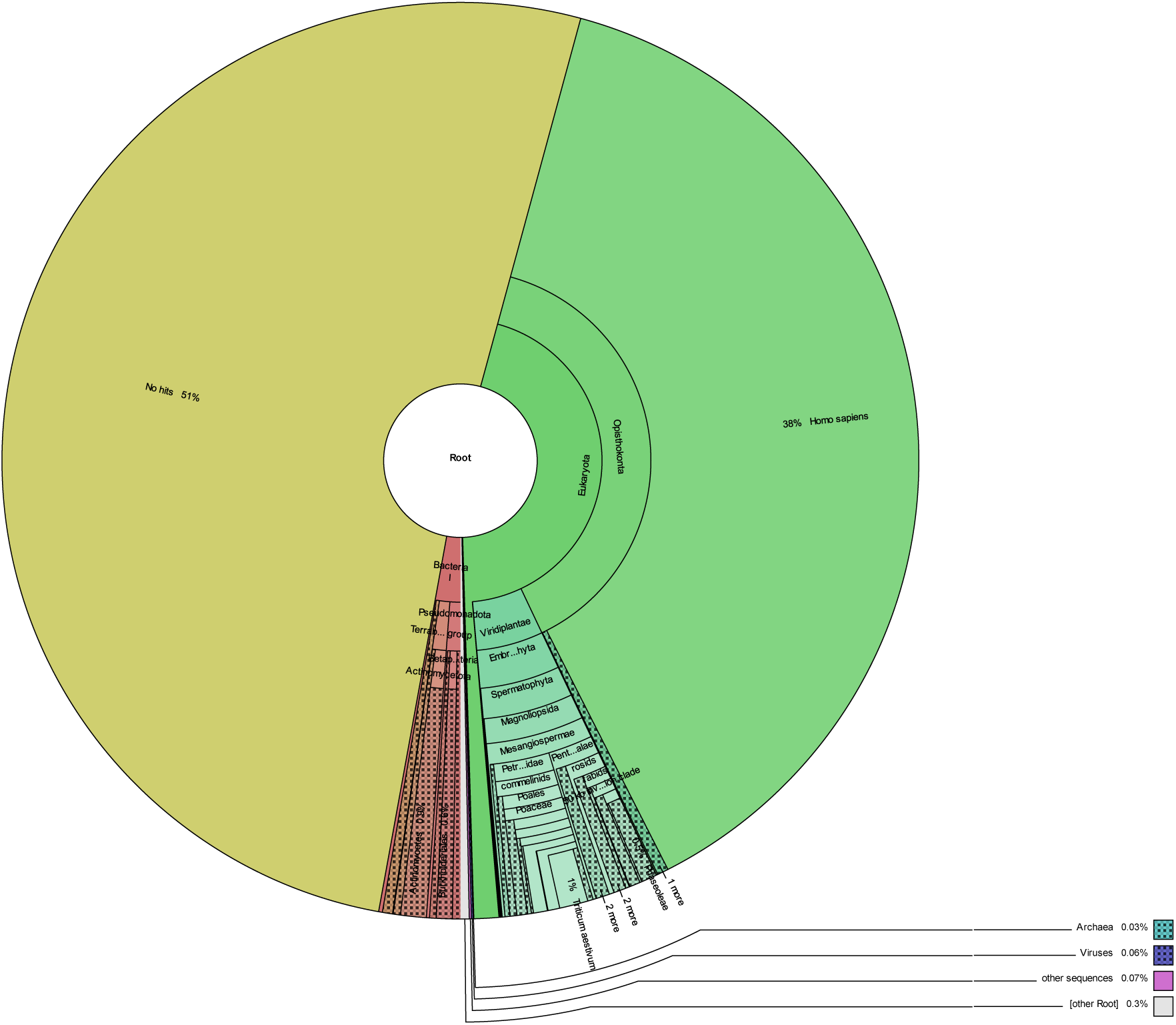
The diversity and relative abundance of the hits retrieved by Kraken were graphically represented with the Krona package v2.7.1.

The reads preliminarily classified as human by Kraken were identified to the lowest possible taxonomic level by comparing the sequences obtained against a local instance of the NCBI’s Nucleotide database (last updated in 21/03/2023) using the algorithm BLASTn v2.13.0+ (Camacho, y otros, 2009). To do so, we used the following parameters: percent identity of 97 %, e-value of 1e-05, and a minimum hit coverage of 80 %.

### 3.- Statistical and descriptive analysis. Descriptive results of MT_TOTAL MT_PURE sequences

#### 3.1 Descriptive analysis of MT_TOTAL sequences

The MT_TOTAL dataset, composed of 15 sequences, has an average length of 282.80 bases. Individual lengths range from 201 bases (minimum) to 498 bases (maximum), indicating moderate heterogeneity in the sizes of the assembled sequences.

The base composition in this dataset is as follows: AT content is 51.93%, while GC content is 48.07%. This reflects a slight predominance of adenine (A) and thymine (T), characteristic of certain genomic regions. Detailed values are presented in Table 1 and Table 2.

**Table 1.** Descriptive results for MT_TOTAL and MT_PURE sequences.

| Characteristic | MT_TOTAL | MT_PURE |
| --- | --- | --- |
| Total Sequences | 15 | 13 |
| Average Length (bp) | 282.80 | 268.54 |
| Min Length (bp) | 201 | 203 |
| Max Length (bp) | 498 | 498 |
| A Count (% of total) | 1,088 (25.65%) | 974 (27.90%) |
| T Count (% of total) | 1,115 (26.28%) | 916 (26.24%) |
| C Count (% of total) | 1,045 (24.63%) | 893 (25.58%) |
| G Count (% of total) | 994 (23.43%) | 708 (20.28%) |
| A+T Content (%) | 51.93% | 54.14% |
| G+C Content (%) | 48.07% | 45.86% |

**Table 2.** Sequence-specific composition details for MT_TOTAL.

| ID | Length | AT% | GC% | A Count (%) | T Count (%) | G Count (%) | C Count (%) |
| --- | --- | --- | --- | --- | --- | --- | --- |
| MT_TOTAL_SEQ1 | 498 | 54.00% | 46.00% | 27.00% | 27.00% | 23.00% | 23.00% |
| MT_TOTAL_SEQ2 | 350 | 52.00% | 48.00% | 26.00% | 26.00% | 24.00% | 24.00% |
| MT_TOTAL_SEQ3 | 201 | 55.00% | 45.00% | 28.00% | 27.00% | 22.00% | 23.00% |
| MT_TOTAL_SEQ4 | 220 | 50.45% | 49.55% | 25.23% | 25.23% | 24.77% | 24.77% |
| MT_TOTAL_SEQ5 | 270 | 58.15% | 41.85% | 30.37% | 27.78% | 25.56% | 16.30% |
| MT_TOTAL_SEQ6 | 264 | 75.00% | 25.00% | 55.30% | 19.70% | 9.09% | 15.91% |
| MT_TOTAL_SEQ7 | 233 | 54.94% | 45.06% | 28.76% | 26.18% | 21.07% | 23.99% |
| MT_TOTAL_SEQ8 | 303 | 44.88% | 55.12% | 24.09% | 20.79% | 12.21% | 42.90% |
| MT_TOTAL_SEQ9 | 201 | 46.77% | 53.23% | 21.89% | 24.88% | 26.61% | 26.61% |
| MT_TOTAL_SEQ10 | 447 | 57.94% | 42.06% | 33.56% | 24.38% | 21.50% | 20.56% |
| MT_TOTAL_SEQ11 | 498 | 57.83% | 42.17% | 23.90% | 33.94% | 11.65% | 30.52% |
| MT_TOTAL_SEQ12 | 240 | 38.33% | 61.67% | 23.33% | 15.00% | 12.92% | 48.75% |
| MT_TOTAL_SEQ13 | 223 | 46.64% | 53.36% | 25.56% | 21.08% | 11.21% | 42.15% |
| MT_TOTAL_SEQ14 | 219 | 34.70% | 65.30% | 28.31% | 6.39% | 56.62% | 8.68% |
| MT_TOTAL_SEQ15 | 215 | 33.02% | 66.98% | 3.26% | 29.77% | 9.30% | 57.67% |

The slightly higher A+T content compared to G+C content suggests an AT-rich bias within these sequences. This characteristic may influence the structural flexibility and stability of the DNA, as well as its interaction with proteins and other biomolecules. The observed enrichment in AT content and the presence of specific conserved motifs suggest a potential role in autoreplication, similar to what has been proposed for early replicative systems. This hypothesis aligns with scenarios of prebiotic chemistry, where simpler genomic elements could replicate under minimal conditions. Further experimental validation is needed to confirm whether these ssDNA sequences could sustain autoreplication, a property that would significantly enhance their relevance as potential precursors to early life forms.

The unique characteristics of the ssDNA sequences, including their lack of matches in existing genomic databases and their structural conservation, highlight their significance in understanding molecular diversity and adaptation. While these sequences could represent potential biosignatures, their origin and role remain open to interpretation. The observed features invite further exploration into their possible relationship with biological systems adapted to extreme environments, such as those found in meteorites. These findings emphasize the value of investigating uncharacterized nucleic acid sequences in extreme contexts, which could inform future research on molecular adaptation and evolution without presupposing extraterrestrial origins.

Finally, the balance observed in G+C content contributes to sequence stability, which may have implications for replication and transcription processes.

The sequences exhibit a slight bias toward A+T content, indicating a balanced yet slightly AT-rich composition. G+C content, while lower, remains significant, contributing to the overall stability of the sequences. This balance between A+T and G+C might reflect evolutionary adaptations or functional constraints. The distribution of sequence lengths demonstrates variability, with certain lengths appearing more frequently, possibly due to selective pressures or functional requirements. These unique ssDNA regions, which lack significant matches in existing databases, may represent evolutionary adaptations to extreme environments. In astrobiological contexts, such adaptations could hint at the potential for life in extreme conditions, although they are equally compatible with abiotic processes. These findings serve as a foundation for exploring the implications of molecular diversity in challenging environments.

The findings from this study underscore the importance of understanding sequence composition and length variability. The observed nucleotide bias and sequence length distribution may have implications for the functionality, stability, and evolutionary history of these ssDNA sequences. For instance, sequences with higher AT content might exhibit greater flexibility, potentially impacting their roles in regulatory or structural contexts. Moreover, the detailed compositional analysis provides a foundation for predicting the behavior of these sequences in different environmental and biological conditions. Future studies could expand on these results by examining specific motifs, secondary structures, or interactions with other biomolecules.

This descriptive study provides a comprehensive understanding of the characteristics of the analyzed ssDNA sequences. The balanced nucleotide composition, slight A+T bias, and variability in sequence lengths suggest a dataset with diverse structural and functional potentials. These findings can serve as a basis for further functional or evolutionary analysis, offering insights into the biological roles and adaptations of these sequences.

#### 3.2 Descriptive analysis of MT_PURE sequences

The MT_PURE dataset, composed of 13 sequences, has an average length of 268.54 bases, ranging from 203 bases (minimum) to 498 bases (maximum), with a total of 3,491 bases. This dataset provides a solid basis for exploring the structural and functional roles of these sequences. This reflects a similar size distribution compared to MT_TOTAL, although the dataset is smaller due to the cleaning process (Table 1).

The base composition in this dataset is as follows: AT content is 54.14%, while GC content is 45.86%. This enrichment in AT content suggests that the curation process preferentially retained AT-rich sequences while removing those with a more balanced or GC-biased composition. Detailed values are provided in Table 3.

**Table 3.**
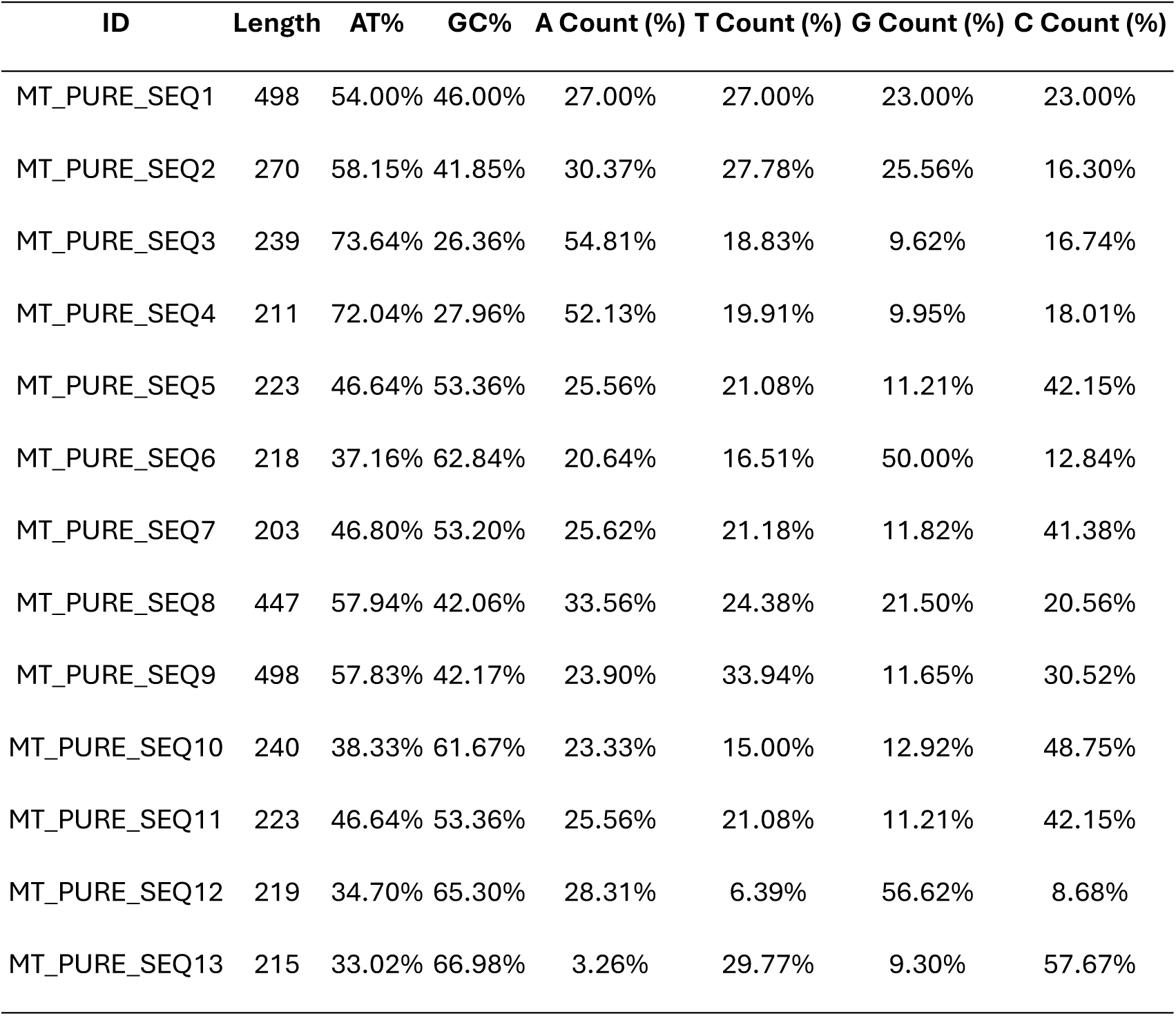
Sequence-specific composition details for MT_PURE.

| ID | Length | AT% | GC% | A Count (%) | T Count (%) | G Count (%) | C Count (%) |
| --- | --- | --- | --- | --- | --- | --- | --- |
| MT_PURE_SEQ1 | 498 | 54.00% | 46.00% | 27.00% | 27.00% | 23.00% | 23.00% |
| MT_PURE_SEQ2 | 270 | 58.15% | 41.85% | 30.37% | 27.78% | 25.56% | 16.30% |
| MT_PURE_SEQ3 | 239 | 73.64% | 26.36% | 54.81% | 18.83% | 9.62% | 16.74% |
| MT_PURE_SEQ4 | 211 | 72.04% | 27.96% | 52.13% | 19.91% | 9.95% | 18.01% |
| MT_PURE_SEQ5 | 223 | 46.64% | 53.36% | 25.56% | 21.08% | 11.21% | 42.15% |
| MT_PURE_SEQ6 | 218 | 37.16% | 62.84% | 20.64% | 16.51% | 50.00% | 12.84% |
| MT_PURE_SEQ7 | 203 | 46.80% | 53.20% | 25.62% | 21.18% | 11.82% | 41.38% |
| MT_PURE_SEQ8 | 447 | 57.94% | 42.06% | 33.56% | 24.38% | 21.50% | 20.56% |
| MT_PURE_SEQ9 | 498 | 57.83% | 42.17% | 23.90% | 33.94% | 11.65% | 30.52% |
| MT_PURE_SEQ10 | 240 | 38.33% | 61.67% | 23.33% | 15.00% | 12.92% | 48.75% |
| MT_PURE_SEQ11 | 223 | 46.64% | 53.36% | 25.56% | 21.08% | 11.21% | 42.15% |
| MT_PURE_SEQ12 | 219 | 34.70% | 65.30% | 28.31% | 6.39% | 56.62% | 8.68% |
| MT_PURE_SEQ13 | 215 | 33.02% | 66.98% | 3.26% | 29.77% | 9.30% | 57.67% |

The sequence lengths show moderate variability, with the majority clustered near the mean length of 268.54 bases. This uniformity could indicate conserved structural features. The A+T content is higher than G+C, which is typical of sequences requiring increased flexibility or reduced stability in secondary structure formation. The sequence-specific data for MT_PURE illustrates a tendency toward AT enrichment and highlights the impact of the cleaning process on sequence composition. The relatively balanced C and G content (20.28% and 25.58%, respectively) contributes to moderate overall sequence stability.

The findings suggest that MT_PURE sequences are predominantly AT-rich, which could be associated with biological processes requiring dynamic structural adjustments. The length and composition variability observed in this dataset might reflect functional diversity or adaptation to specific environmental conditions.

This descriptive study highlights the structural and compositional characteristics of MT_PURE sequences. The higher A+T content and relatively uniform sequence lengths suggest potential functional roles requiring flexibility and moderate stability. These findings pave the way for further research into the evolutionary, structural, and functional implications of these sequences.

#### 3.3 Comparison between MT_TOTAL and MT_PURE

MT_TOTAL contains more sequences (15) compared to MT_PURE (13), reflecting the impact of the cleaning process in reducing the dataset by removing sequences with potential matches or contaminants. Both datasets share the same maximum length (498 bases). However, MT_PURE has a slightly higher minimum length (203 bases compared to 201 bases in MT_TOTAL), indicating the exclusion of shorter sequences during curation. The AT content in MT_PURE (54.14%) is higher than in MT_TOTAL (51.93%), suggesting a selective retention of AT-rich sequences in MT_PURE. Correspondingly, GC content is slightly reduced in MT_PURE (45.86%) compared to MT_TOTAL (48.07%), highlighting the impact of cleaning on the sequence pool.

The identification of identical sequences provides valuable insight into conserved regions, which may play critical roles in the biological functions or evolutionary relationships within the datasets. A total of 6 sets of identical sequences were identified between the two series of assemblies: the "MT_TOTAL" and "MT_PURE" datasets. These sequences represent regions consistently detected across both series, reinforcing their biological significance and methodological robustness.

Below are the identical sequences identified, along with their associated IDs:

- Shared by: MT_TOTAL_SEQ2, MT_PURE_SEQ2
- Shared by: MT_TOTAL_SEQ6, MT_PURE_SEQ5
- Shared by: MT_TOTAL_SEQ7, MT_PURE_SEQ6
- Shared by: MT_TOTAL_SEQ9, MT_PURE_SEQ1
- Shared by: MT_TOTAL_SEQ10, MT_PURE_SEQ8
- Shared by: MT_TOTAL_SEQ12, MT_PURE_SEQ9

Inclusion in joint analyses.

Identical sequences are significant because their consistent detection across both series highlights their potential biological importance. As such:

- In separate analyses of each series, these sequences will naturally remain present, ensuring consistency within each dataset.
- In joint analyses, these identical sequences will be explicitly included, as they may contribute key information to studies of:

- Phylogenetic relationships (e.g., distance matrices and phylogenetic trees).
- Motif discovery using MEME and FIMO.
- Secondary structure prediction.
- Functional annotation via tools like PROKKA.

The observed differences between the datasets suggest that the cleaning process enhanced the specificity of the MT_PURE dataset by preferentially retaining sequences with higher AT content. This may align with the biological characteristics of certain ssDNA viruses or non-coding regions, which are often AT-rich.

The comparative analysis of MT_TOTAL and MT_PURE highlights the significant effects of the cleaning process on sequence composition and dataset size. While MT_TOTAL represents a broader assembly of non-hit sequences, MT_PURE reflects a more curated dataset with enriched AT content and reduced sequence redundancy. Both datasets provide valuable insights into novel ssDNA sequences. Their differences underscore the importance of dataset refinement for downstream analyses, including functional characterization and evolutionary studies.

##### 3.3.1 Descriptive analysis of identical sequences

The analysis focused on 6 sequences identified as identical across the MT_TOTAL and MT_PURE datasets. These sequences were subjected to a detailed descriptive study to evaluate their length distribution, base composition, and overall nucleotide content.

The dataset consists of a total of 6 sequences, comprising 1,943 bases. The average sequence length is 323.83 bases, with a maximum sequence length of 498 bases and a minimum sequence length of 218 bases. The base composition analysis revealed 527 adenine (A) bases (27.12%), 507 thymine (T) bases (26.11%), 447 guanine (G) bases (23.01%), and 462 cytosine (C) bases (23.77%). The overall AT content is 53.22%, while the GC content is 46.78%. This indicates a slight predominance of AT content, a pattern commonly observed in non-coding or viral genomic regions.

The lengths of the sequences showed variability, ranging from a minimum of 218 bases to a maximum of 498 bases. The average length was approximately 324 bases, highlighting a moderate distribution of sizes within the dataset.

The dataset reveals a higher AT content, as reflected in the AT/GC ratio, which aligns with certain characteristics of ssDNA and may indicate specific biological or structural properties of these sequences. Additionally, the variability in sequence length suggests a degree of heterogeneity, potentially related to sequence assembly or underlying biological features.

The analyzed sequences exhibit unique features, including a slight predominance of AT-rich regions and a relatively balanced nucleotide composition. These characteristics may offer valuable insights into the functional roles or evolutionary origins of the sequences. To fully understand the significance of these findings, further comparative analyses with existing databases and deeper functional characterizations are recommended.

##### 3.3.2 Comparative analysis of identical sequences with MT_TOTAL and MT_PURE

The identical sequences analyzed represent a subset of sequences found in both MT_TOTAL and MT_PURE, providing a unique opportunity to assess how these sequences compare to the broader datasets in terms of length, base composition, and nucleotide content. This comparison aims to highlight their similarities (Table 4), differences, and potential implications.

**Table 4.** Statistical comparison.

| **Metric** | **MT_TOTAL** | **MT_PURE** | **Identical Sequences** |
| --- | --- | --- | --- |
| Total Sequences | 15 | 13 | 6 |
| Total Bases | 4,242 | 3,491 | 1,943 |
| Average Length (bp) | 282.80 | 268.54 | 323.83 |
| Max Length (bp) | 498 | 498 | 498 |
| Min Length (bp) | 201 | 203 | 218 |
| AT Content (%) | 51.93 | 54.14 | 53.22 |
| GC Content (%) | 48.07 | 45.86 | 46.78 |

The identical sequences represent a smaller subset compared to the total datasets, consisting of only 6 sequences that contribute 1,943 bases. This accounts for approximately 45.8% of the average sequence length in MT_TOTAL and 55.7% of MT_PURE. Their smaller count and total bases suggest a more selective grouping of sequences that may share specific conserved characteristics.

In terms of length distribution, the identical sequences share the same maximum length (498 bp) as both MT_TOTAL and MT_PURE, indicating the inclusion of some of the longest sequences from the broader datasets. The minimum length of identical sequences (218 bp) is slightly higher than that of MT_TOTAL (201 bp) and MT_PURE (203 bp), reflecting a potential filtering of very short sequences.

Regarding base composition, the AT content of identical sequences (53.22%) is intermediate between MT_TOTAL (51.93%) and MT_PURE (54.14%), suggesting a balance between the broader dataset and the curated MT_PURE sequences. Similarly, the GC content of identical sequences (46.78%) follows the same pattern, sitting between MT_TOTAL (48.07%) and MT_PURE (45.86%).

The identical sequences seem to capture key features of both MT_TOTAL and MT_PURE, serving as a representative subset with conserved lengths and balanced nucleotide content. Their AT and GC contents align closely with MT_PURE, reflecting the selective enrichment observed during the curation process.

Despite their smaller size as a subset, the identical sequences maintain a length profile that is broadly consistent with the longer sequences in both datasets. The consistent AT/GC ratios suggest that these identical sequences may have functional or structural roles tied to the broader characteristics of the datasets. Their shared inclusion across MT_TOTAL and MT_PURE underlines their potential biological significance and resistance to filtering processes.

The identical sequences analyzed exhibit characteristics that make them a unique and valuable subset for further investigation. Their similarities to both MT_TOTAL and MT_PURE highlight their potential as conserved elements, while their slight enrichment in AT content aligns with patterns typical of ssDNA regions.

Further research into these identical sequences, including functional annotation and comparison with external databases, may reveal insights into their biological roles and evolutionary origins. This subset serves as a promising focal point for exploring the diversity and conservation of sequences within the broader datasets.

### 4.- Análisis de diversidad y clustering

#### 4.1. Analysis of MT_TOTAL sequences

##### 4.1.2 Evolutionary distance analysis of MT_TOTAL sequences

The evolutionary distance matrix for MT_TOTAL sequences, derived from MEGA, provides a quantitative assessment of the pairwise evolutionary divergence among the sequences (Figure 2). The values range from 0.024876, indicating minimal evolutionary differences, to higher divergence values such as 0.761194, suggesting substantial evolutionary separation. These distances offer insights into the sequence diversity and evolutionary relationships within the dataset. Key observations from the matrix include:

**Figure 2.**
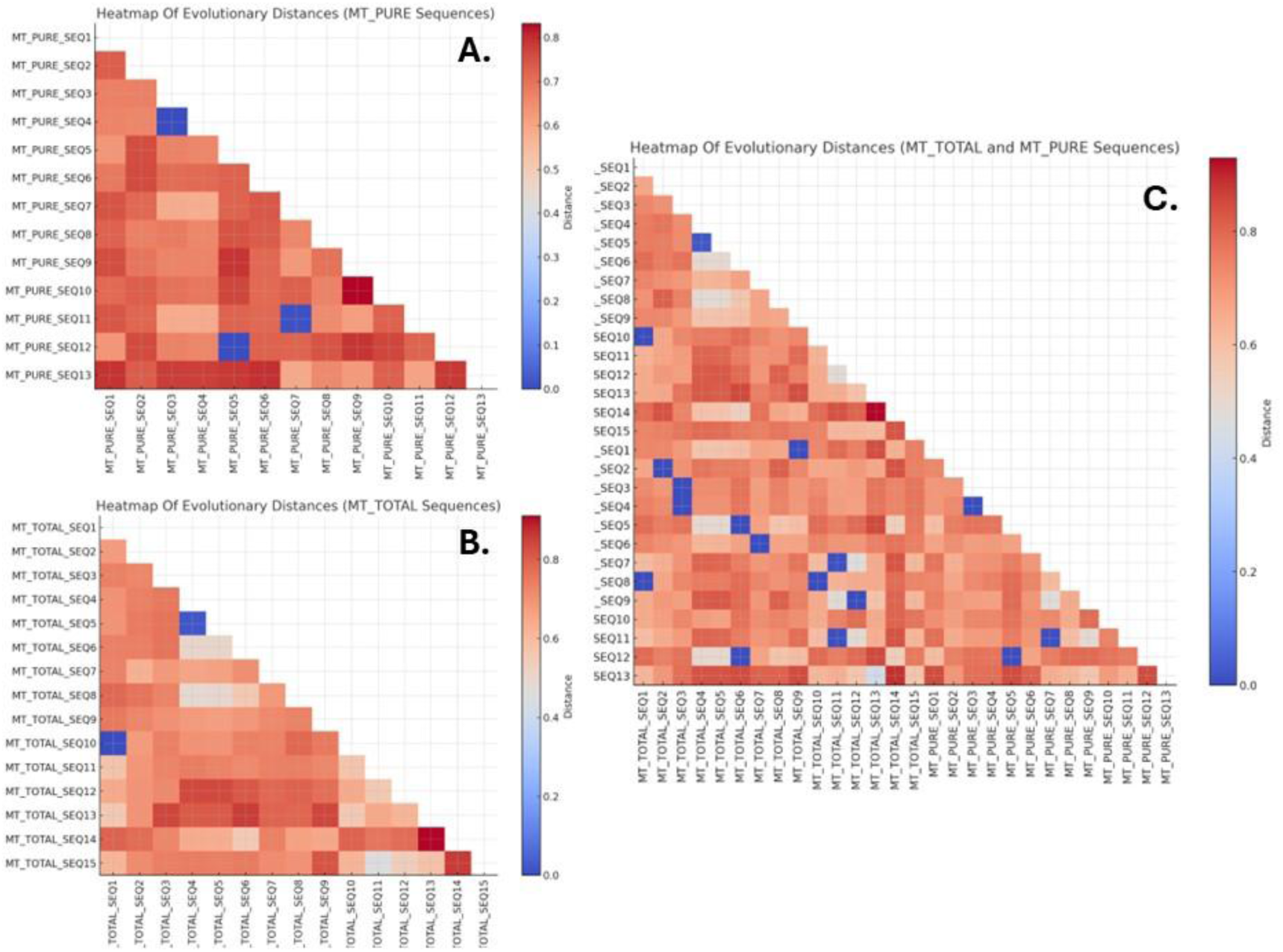
Heatmap of genetic distances. (A) MT_TOTAL. (B) MT_PURE. (C) MT_TOTAL and MT_PURE

— Low evolutionary divergence: MT_TOTAL_SEQ4 and MT_TOTAL_SEQ5 exhibit the smallest evolutionary distance (0.024876), signifying a very close relationship. This high similarity might indicate conserved functionality or structural roles among these sequences. Similarly, MT_TOTAL_SEQ1 and MT_TOTAL_SEQ10 demonstrate low divergence, supporting their clustering in the phylogenetic tree and suggesting shared evolutionary pressures.
— Moderate evolutionary divergence: sequences such as MT_TOTAL_SEQ6 and MT_TOTAL_SEQ7 show intermediate divergence values with other sequences, highlighting their partial similarity to both conserved and divergent clusters. These sequences likely serve as evolutionary intermediates.
— High evolutionary divergence: MT_TOTAL_SEQ3 stands out with consistently high divergence values relative to other sequences. This indicates a unique evolutionary lineage and possible distinct functional characteristics. Similarly, MT_TOTAL_SEQ9 and MT_TOTAL_SEQ14 also show significant divergence, forming distinct clades in the phylogenetic tree.

The distribution of distances underscores the presence of both conserved and highly diverse sequences within the MT_TOTAL dataset, providing a complex picture of evolutionary relationships.

##### 4.1.3 Phylogenetic tree analysis of MT_TOTAL sequences

The phylogenetic tree constructed in MEGA offers a visual representation of the evolutionary relationships among MT_TOTAL sequences (Figure 3). It highlights clustering patterns and distinct lineages, reflecting the evolutionary distances outlined above. Detailed observations from the tree include:

**Figure 3.**
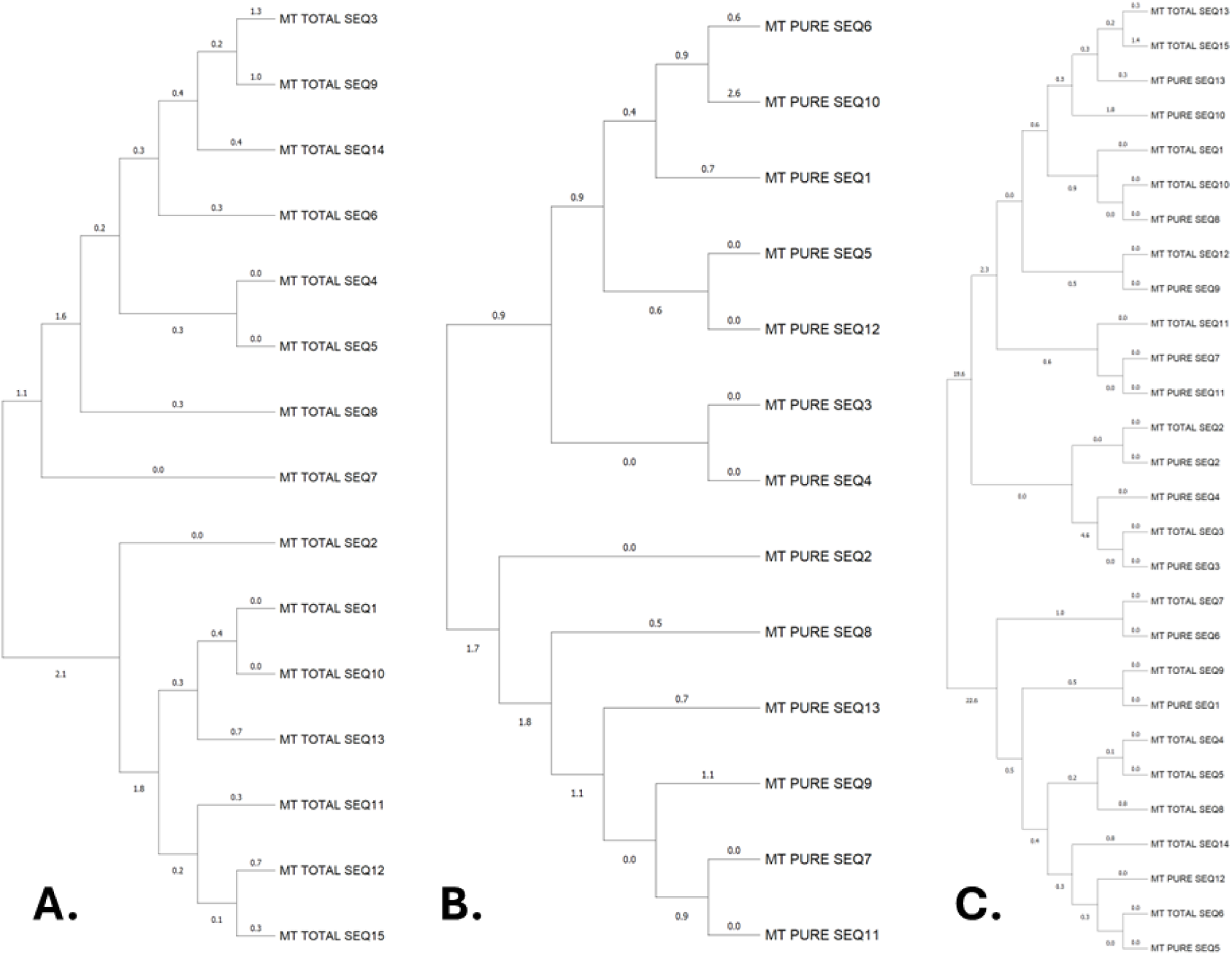
Phylogenetic tree constructed using the (A) MT_TOTAL sequences, (B) MT_PURE and (C) MT_TOTAL + MT_PURE.

— Conserved clusters: MT_TOTAL_SEQ4, MT_TOTAL_SEQ5, and MT_TOTAL_SEQ8 form a closely related group, consistent with their low pairwise evolutionary distances. These sequences likely share common evolutionary origins or conserved biological roles. MT_TOTAL_SEQ1 and MT_TOTAL_SEQ10 also group together, reinforcing the evolutionary proximity indicated by their matrix values.
— Divergent lineages: MT_TOTAL_SEQ3 diverges early in the tree, forming a separate branch. This unique lineage suggests significant evolutionary pressures that differentiate it from other sequences. MT_TOTAL_SEQ9 and MT_TOTAL_SEQ14 also appear in isolated clades, emphasizing their divergence and potential functional novelty.
— Intermediate groupings: MT_TOTAL_SEQ6 and MT_TOTAL_SEQ7 occupy intermediate positions, bridging more conserved clusters with divergent lineages. This placement reflects their moderate evolutionary divergence.

The combined analysis of evolutionary distances and the phylogenetic tree (Figure 3) highlights a dynamic evolutionary landscape among the MT_TOTAL sequences. Several conclusions can be drawn:

— Conserved sequences: clusters such as MT_TOTAL_SEQ4/MT_TOTAL_SEQ5/MT_TOTAL_SEQ8 and MT_TOTAL_SEQ1/MT_TOTAL_SEQ10 suggest functional conservation. These sequences may play critical roles that have been preserved through evolutionary time.
— Unique evolutionary paths: The distinct lineages of MT_TOTAL_SEQ3, MT_TOTAL_SEQ9, and MT_TOTAL_SEQ14 suggest novel evolutionary adaptations or specialized functionalities. Further investigation into these sequences could uncover unique biological insights.
— Diverse evolutionary pressures: The presence of both conserved and divergent sequences within the dataset reflects the impact of varying evolutionary pressures, such as environmental factors, selective advantages, and genomic drift.

These findings lay the groundwork for future studies exploring the functional and structural roles of these non-hit ssDNA sequences. While conserved clusters may offer insights into essential biological processes, the divergent sequences represent opportunities to uncover novel genetic and evolutionary phenomena.

#### 4.2. Analysis of MT_PURE sequences

##### 4.2.1 Evolutionary distance analysis of MT_PURE sequences

The evolutionary distance matrix for MT_PURE sequences, derived from MEGA, provides a quantitative measure of the pairwise evolutionary divergence among the sequences (Figure 2). The values range from 0.000000, representing identical sequences, to higher divergence values such as 0.755760, indicating substantial evolutionary differences. Key observations from the distance matrix include:

— Highly similar sequences: MT_PURO_SEQ3 and MT_PURO_SEQ4 show an evolutionary distance of 0.000000, indicating identical sequences. This complete similarity suggests these sequences likely share identical evolutionary pressures and functional roles. MT_PURO_SEQ7 and MT_PURO_SEQ11 also exhibit low evolutionary distances, clustering closely in the phylogenetic tree.
— Moderately divergent sequences: MT_PURO_SEQ5 and MT_PURO_SEQ12 show intermediate distances relative to other sequences, forming part of clusters with moderate evolutionary divergence. Sequences like MT_PURO_SEQ1 and MT_PURO_SEQ6 demonstrate moderate divergence from other clusters, aligning with their placement in the phylogenetic tree.
— Highly divergent sequences: MT_PURO_SEQ2 and MT_PURO_SEQ10 show higher divergence values, reflecting their distinct evolutionary trajectories. These sequences are positioned on separate clades in the phylogenetic tree.

##### 4.2.2 Phylogenetic tree analysis of MT_PURE sequences

The phylogenetic tree constructed in MEGA reveals hierarchical clustering patterns and distinct lineages among the MT_PURE sequences (Figure 3). Major insights from the tree include:

— Conserved clusters: MT_PURO_SEQ3 and MT_PURO_SEQ4 form a tightly linked group, consistent with their identical evolutionary distances. These sequences likely share common evolutionary origins and functional roles. MT_PURO_SEQ7 and MT_PURO_SEQ11 cluster together in a similar manner, suggesting evolutionary conservation.
— Intermediate divergence clusters: MT_PURO_SEQ5, MT_PURO_SEQ6, and MT_PURO_SEQ12 form intermediate clusters, bridging conserved and highly divergent lineages. These sequences likely represent evolutionary intermediates with partial similarity to both conserved and divergent clusters.
— Divergent lineages: MT_PURO_SEQ2 and MT_PURO_SEQ10 occupy separate branches, reflecting significant evolutionary distances and possible functional specialization. MT_PURO_SEQ13 also forms a distinct lineage, aligning with its divergent characteristics in the distance matrix.

The combined analysis of evolutionary distances and the phylogenetic tree for MT_PURE sequences highlights a complex evolutionary landscape. Several key findings include:

- Identical and conserved sequences: the clustering of MT_PURO_SEQ3/MT_PURO_SEQ4 and MT_PURO_SEQ7/MT_PURO_SEQ11 underscores the presence of highly conserved sequences within the dataset. These sequences likely play critical and conserved biological roles.
- Intermediate evolutionary relationships: sequences such as MT_PURO_SEQ5, MT_PURO_SEQ6, and MT_PURO_SEQ12 exhibit moderate divergence, indicating shared evolutionary pressures but with some level of functional or structural differentiation.
- Unique lineages: highly divergent sequences like MT_PURO_SEQ2, MT_PURO_SEQ10, and MT_PURO_SEQ13 suggest novel evolutionary adaptations or specialized functionalities. Further investigation into these sequences may uncover unique biological insights.

These findings emphasize the diversity of evolutionary trajectories within the MT_PURE dataset. The combination of conserved and divergent sequences reflects a dynamic evolutionary process influenced by various pressures and constraints. This analysis provides a foundation for further exploration of the functional and structural roles of these sequences.

#### 4.3. Results: sequence distances and phylogenetic analysis of ssDNA sequences MT_TOTAL and MT_PURE

The distances between ssDNA sequences were calculated based on sequence alignment metrics (Figure 2). Sequence distances provide insight into evolutionary relationships, with smaller distances indicating higher sequence similarity.

Summary of Sequence Distance Results.

- MT_TOTAL_SEQ1 and MT_TOTAL_SEQ10: these sequences showed identical alignment results with zero mismatch points, indicating they are identical or highly conserved.
- MT_TOTAL_SEQ4 and MT_TOTAL_SEQ5: minimal sequence variation was observed with a minor number of substitutions. These sequences appear to form a subgroup within the dataset.
- MT_PURE_SEQ3 and MT_PURE_SEQ4: the sequences exhibited significant similarity, with alignment mismatches confined to repetitive regions, suggesting a close evolutionary relationship.
- Divergent sequences: MT_TOTAL_SEQ13 and MT_PURE_SEQ13 showed the largest pairwise distance from the rest of the dataset due to high numbers of unique repetitive elements and substitutions.

Coincident points (regions of identical sequence) were noted across multiple alignments. These regions often corresponded to highly conserved functional motifs, likely critical for biological function. Particularly conserved regions included: repeats in MT_TOTAL_SEQ4/MT_TOTAL_SEQ5 and conserved domains in MT_PURE_SEQ1 and MT_PURE_SEQ8.

Phylogenetic tree construction.

A phylogenetic tree was constructed to elucidate the evolutionary relationships among the ssDNA sequences (Figure 3). The tree was generated using a maximum likelihood approach with sequence alignments as input. The topology of the tree highlights distinct clusters based on sequence similarity and divergence.

Key observations:

- Major clades:

- Clade A: includes MT_TOTAL_SEQ1, MT_TOTAL_SEQ10, and MT_PURE_SEQ8. This clade represents a highly conserved group with minimal sequence divergence.
- Clade B: contains MT_TOTAL_SEQ4, MT_TOTAL_SEQ5, and MT_PURE_SEQ5. These sequences share a common ancestor characterized by repetitive motifs.
- Clade C: MT_TOTAL_SEQ13 and MT_PURE_SEQ13 formed a distinct clade, separate from other groups due to unique sequence characteristics.
- Subclades and evolutionary divergence:

- Subclades were evident within Clade B, with MT_TOTAL_SEQ4 and MT_TOTAL_SEQ5 forming a subgroup. This subgroup likely diverged from the common ancestor more recently.
- In Clade A, MT_TOTAL_SEQ1 and MT_TOTAL_SEQ10 exhibited minimal branch lengths, confirming their identical sequences.
- Outgroup analysis:

- MT_PURE_SEQ10 was used as an outgroup for rooting the tree. The sequence’s unique characteristics, including distinct repetitive patterns and substitutions, make it suitable for this role.

Functional insights from sequence conservation and variation.

— Highly conserved regions. Coincide with functional motifs such as binding sites and structural domains. Example: Conserved repeats in MT_TOTAL_SEQ1 and MT_TOTAL_SEQ10 are likely indicative of critical functional roles.
— Regions of high variation. Found predominantly in repetitive sequences (e.g., MT_TOTAL_SEQ13 and MT_PURE_SEQ13). These variations may reflect adaptive evolutionary pressures or non-functional mutational drift.

Implications and conclusions. The analysis of ssDNA sequence distances and the resulting phylogenetic tree provides several critical insights:

- Conservation and divergence: Strong conservation is evident among certain sequence groups, reflecting functional importance. Divergent sequences highlight evolutionary adaptability or niche specialization.
- Phylogenetic relationships: The tree topology confirms the evolutionary clustering of sequences based on conserved features and shared ancestry.
- Biological relevance: Identified conserved motifs and divergent patterns can guide further investigations into functional domains and evolutionary mechanisms in ssDNA sequences.

### 5.- Identification of mobile and repetitive genetic elements. RepeatMasker analysis of meteorite sequences

#### 5.1. MT_TOTAL dataset

The RepeatMasker analysis of the MT_TOTAL dataset (comprising 15 sequences with a total length of 4,242 base pairs) revealed the following key findings.

— Overall sequence composition. The sequences have a GC content of 48.07%, consistent with eukaryotic genomes. 22.11% of the total sequence (938 base pairs) was masked due to the presence of repetitive regions.
— Repetitive elements. No interspersed repetitive elements (e.g., SINEs, LINEs, LTR elements, or DNA transposons) were detected in the dataset. The analysis did not identify common repetitive families such as ALUs, MIRs, LINE1, LINE2, or retroviral elements (ERVs). These are typically abundant in many genomes but were absent in this dataset.
— Simple repeats and low-complexity regions. The dataset contained 5 simple repeat elements, accounting for 15.63% of the total sequence (663 base pairs). These simple repeats include motifs such as (CCATAC)n, (TTC)n, and (ATGGGT)n, commonly observed as tandem repeats. 2 low-complexity regions were identified, representing 6.48% of the total sequence (275 base pairs). Low-complexity regions often consist of stretches with biased base composition (e.g., A/T-rich regions).

The dataset appears to lack interspersed repetitive elements typically found in most well-characterized genomes. The identified simple repeats and low-complexity regions suggest non-coding or repetitive regions, potentially characteristic of certain genomic or synthetic DNA sequences. The relatively short length of the sequences (4,242 bp) limits the likelihood of detecting more complex or interspersed repeats, further indicating that the sequences might represent specific genomic loci with minimal repetitive content.

#### 5.2. MT_PURE dataset

The RepeatMasker analysis of the MT_PURE dataset (comprising 13 sequences with a total length of 3,491 base pairs) revealed the following key findings:

— Overall sequence composition. The sequences have a GC content of 45.86%, within the expected range for eukaryotic genomes. 12.17% of the total sequence (425 base pairs) was masked due to repetitive regions.
— Repetitive elements. Similar to the MT_TOTAL dataset, no interspersed repetitive elements (e.g., SINEs, LINEs, LTR elements, or DNA transposons) were detected in this dataset. The absence of these elements suggests that the dataset may represent specific genomic regions or synthetic sequences without repetitive content typically found in most genomic datasets.
— Simple repeats and low-complexity regions. The dataset contained 3 simple repeat elements, which accounted for 8.91% of the total sequence (311 base pairs). Identified motifs include (CCATAC)n, (CCTC)n, and (CCATCC)n. 2 low-complexity regions were detected, accounting for 3.27% of the total sequence (114 base pairs). These regions are biased in base composition and include A-rich sequences.

The MT_PURE dataset shares a similar pattern to MT_TOTAL, showing an absence of interspersed repetitive elements. The repetitive regions identified primarily consist of simple repeats and low-complexity regions, characteristic of localized repetitive structures. The relatively short sequence length (3,491 bp) further supports the idea that these sequences may represent unique or under-characterized genomic regions not commonly observed in existing repeat databases.

#### 5.3. Overall conclusion

The RepeatMasker analysis of both datasets, MT_TOTAL and MT_PURE, revealed consistent results:

- Both datasets lack significant interspersed repetitive elements such as SINEs, LINEs, LTRs, or DNA transposons, which are typically prevalent in many genomic datasets.
- The identified repetitive content is limited to simple repeats and low-complexity regions, which account for a moderate percentage of the total sequence length in both datasets (22.11% for MT_TOTAL and 12.17% for MT_PURE).

These findings suggest that the analyzed sequences may represent novel or synthetic DNA regions, potentially from under-represented organisms or genomic loci. The absence of complex repetitive elements and the presence of localized repetitive patterns highlight the possibility of these sequences being unique, non-coding, or evolutionary constrained regions. Future analysis using complementary approaches may provide further insight into their origin and function.

### 6.- Identification and analysis of repetitive motifs. Comprehensive results for the analysis of MT_TOTAL and MT_PURO sequences and their comparative study

We used tools such as MEME and FIMO to identify common and unique repetitive motifs in the sequences.

#### 6.1. MEME analysis results for MT_TOTAL

The MEME suite was used to identify conserved motifs in the MT_TOTAL dataset, which comprises 15 ssDNA sequences with lengths ranging between 200 and 500 bp. The primary objective of this analysis was to discover significant motifs and characterize their distribution and recurrence across the dataset. The results provide insights into potential functional or structural elements shared by these sequences.

Identification of motifs in ssDNA Sequences. The MEME analysis of the MT_TOTAL dataset revealed five significant motifs across the 15 ssDNA sequences. These motifs were identified using the zoops (zero or one occurrence per sequence) model, considering both strands. The following Table 5 summarizes the characteristics of the identified motifs.

**Table 5.**
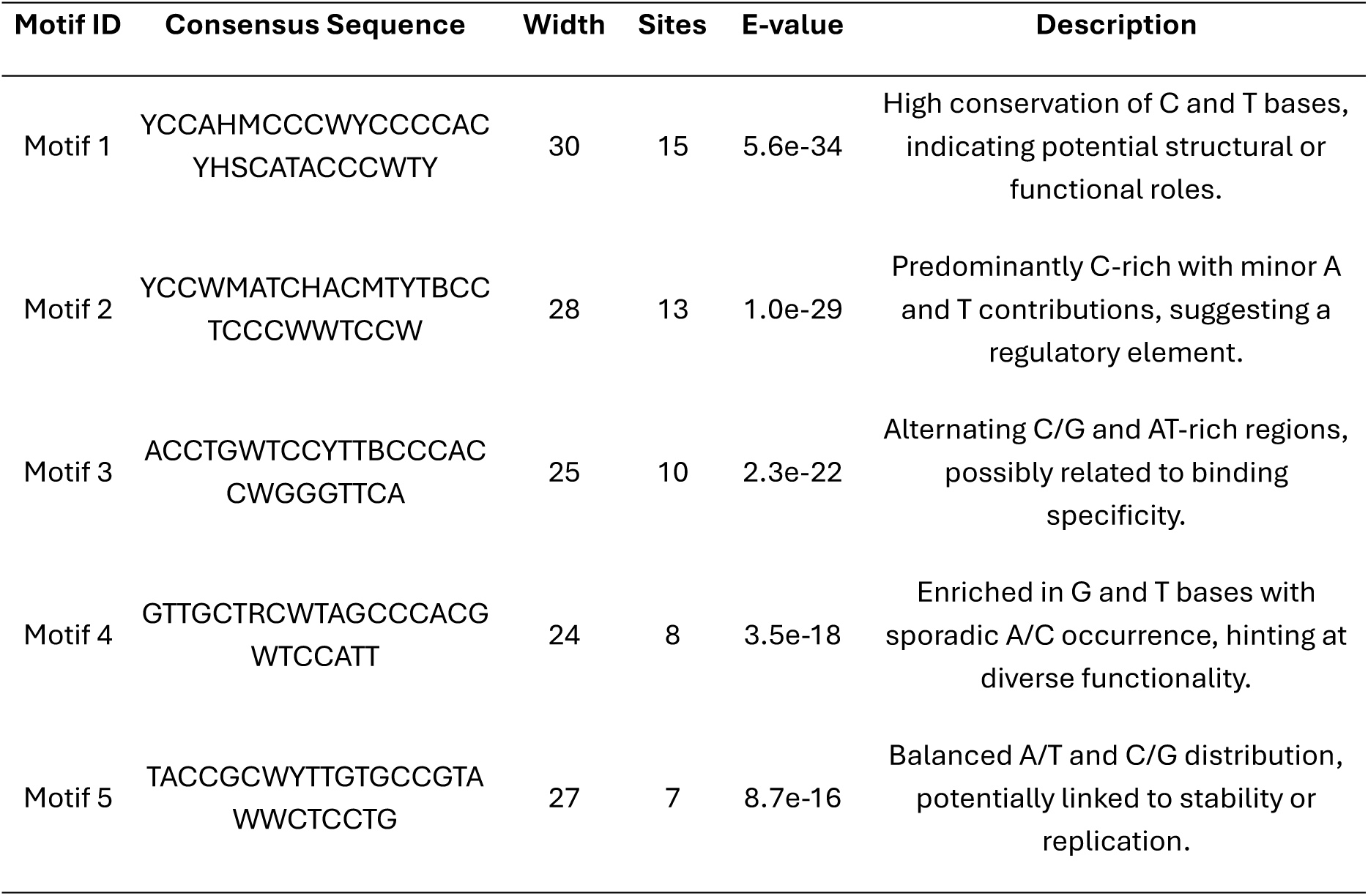
Discovered motifs.

The results indicate that several conserved motifs are present within the MT_TOTAL dataset. These motifs likely represent functional or structural elements shared across these sequences. Key findings include:

1. Highly conserved motif (Motif 1): Its presence in all sequences and high statistical significance (E-value 5.6e-34) strongly suggest that it serves an essential role, possibly as a regulatory or structural element.
2. Recurrent motifs (Motifs 2, 4, 5): Their frequent occurrence in 13 sequences supports the idea that they may interact with or complement Motif 1, contributing to the functional architecture of these sequences.
3. Less frequent motif (Motif 3): Its lower prevalence (7 sequences) implies a specialized or context-specific function.
4. Modular motif distribution: The clustering and co-occurrence of motifs in some sequences could indicate regions of biological significance, such as binding sites for proteins, secondary structure formation, or other regulatory features.

The conserved nature of Motif 1 across all sequences hints at the possibility of a shared evolutionary origin, suggesting these sequences might have diverged from a common ancestor. The variability and presence of Motifs 2 to 5 could reflect adaptations to specific functional requirements or environmental conditions, potentially linking these sequences through convergent evolution or specialized lineage diversification.

Sequence logos of representative motifs identified in the MT_TOTAL are presented in Figure 4. These motifs are potential candidates for functional sequences, suggesting roles in self-replication, ssDNA stability, or other unknown regulatory functions. The size of each letter corresponds to its frequency, with the overall height indicating the conservation at each position in bits.

**Figure 4.**
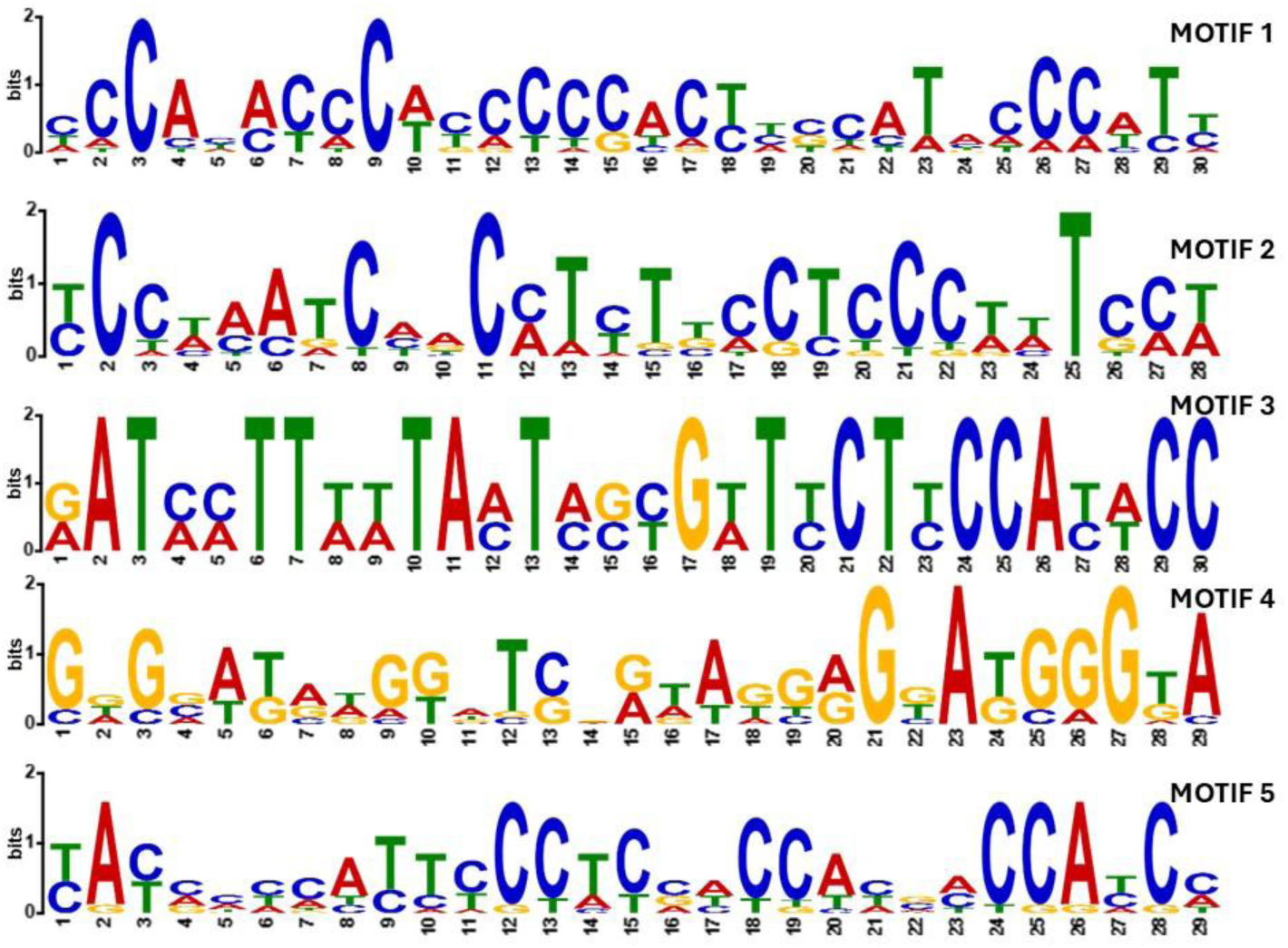
Sequence logos of representative motifs identified in the MT_TOTAL dataset using MEME and analyzed with FIMO. The logos represent the relative frequency of each nucleotide at every position within the motifs, reflecting their conservation and importance.

Motif locations. Motifs identified in the MT_TOTAL dataset (Figure 5) are distributed across the 15 sequences, with each motif represented by a distinct color. Motif 1, found in every sequence, suggests a conserved functional or structural role. Motifs 2, 4, and 5 frequently overlap or cluster near each other, indicating potential modular arrangements or cooperative roles. Certain sequences (e.g., MT_TOTAL_SEQ3) exhibit fewer motifs, while others (e.g., MT_TOTAL_SEQ5) show densely clustered motifs, suggesting functional hotspots.

**Figure 5.**
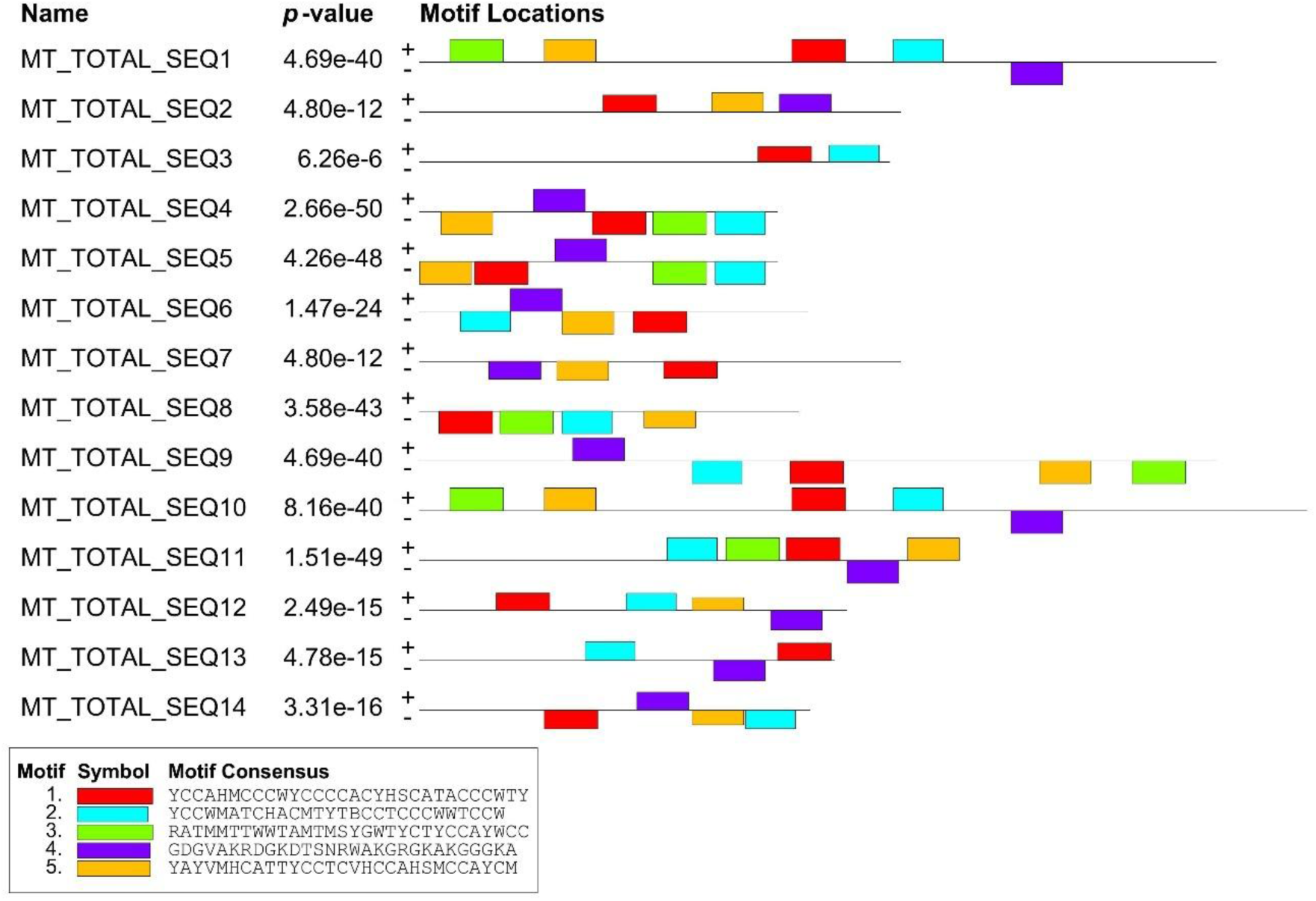
Distribution and locations of identified motifs across MT_TOTAL sequences.

Background model. The nucleotide composition of the dataset was used to construct a 0-order Markov model for the background. The even distribution of nucleotides (A: 0.26, C: 0.24, T: 0.26, G: 0.24) suggests a relatively balanced GC-content, which does not bias motif discovery.

This analysis revealed five significant motifs within the MT_TOTAL dataset, highlighting conserved elements across ssDNA sequences. The findings provide a foundation for further functional characterization of these sequences.

##### 6.1.1 Analysis of MT_TOTAL motifs using TOMTOM

The TOMTOM analysis was conducted using the significant motifs identified in the MEME analysis of the MT_TOTAL dataset. The motifs were compared against multiple motif databases, including JASPAR Vertebrates, UniPROBE Mouse (Table 6), JASPAR CNE (2020) (Table 7), JASPAR FAM (Table 8), JASPAR PHYLOFACTS (2020) (Table 9), and Prokaryote DNA (Table 10). This section details the findings and their integration with MEME results.

**Table 6.**
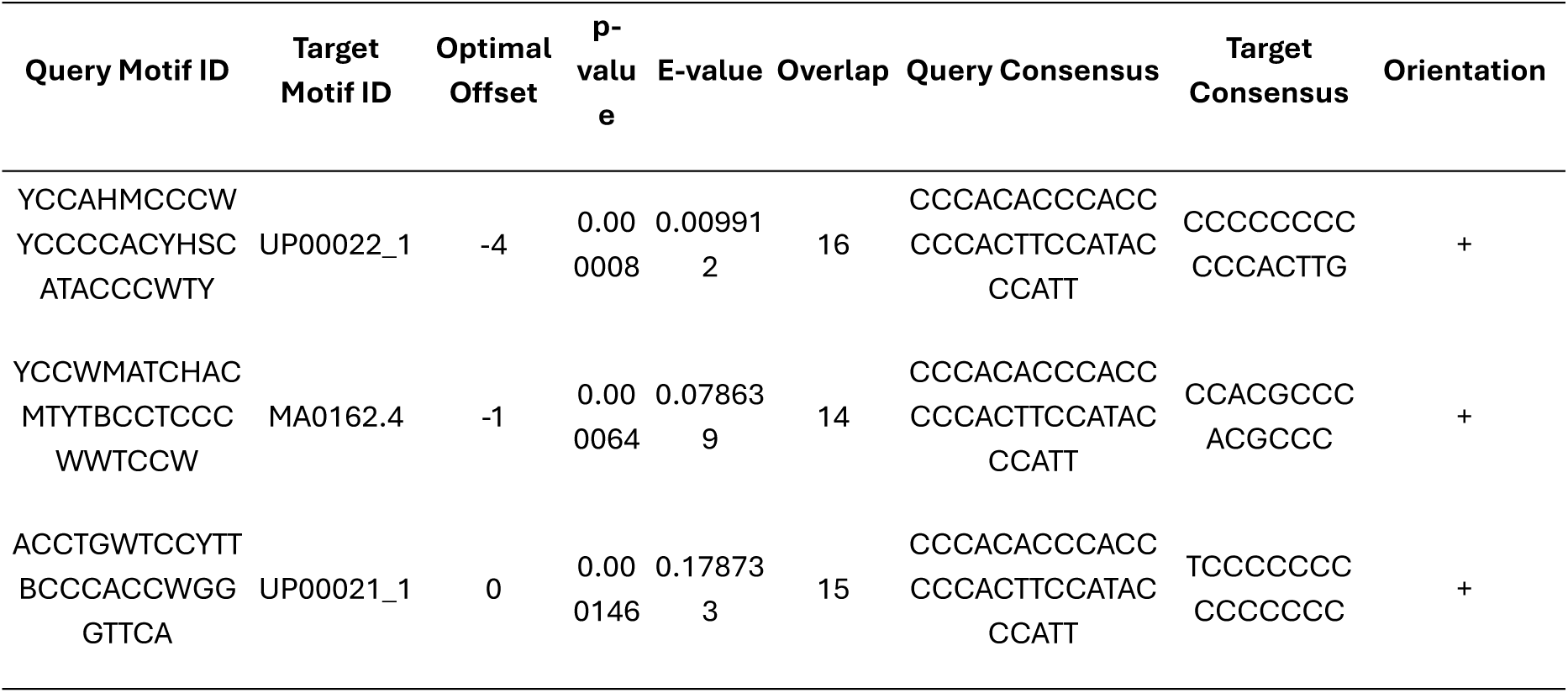
TOMTOM matches: JASPAR vertebrates and UniPROBE mouse.

**Table 7.**
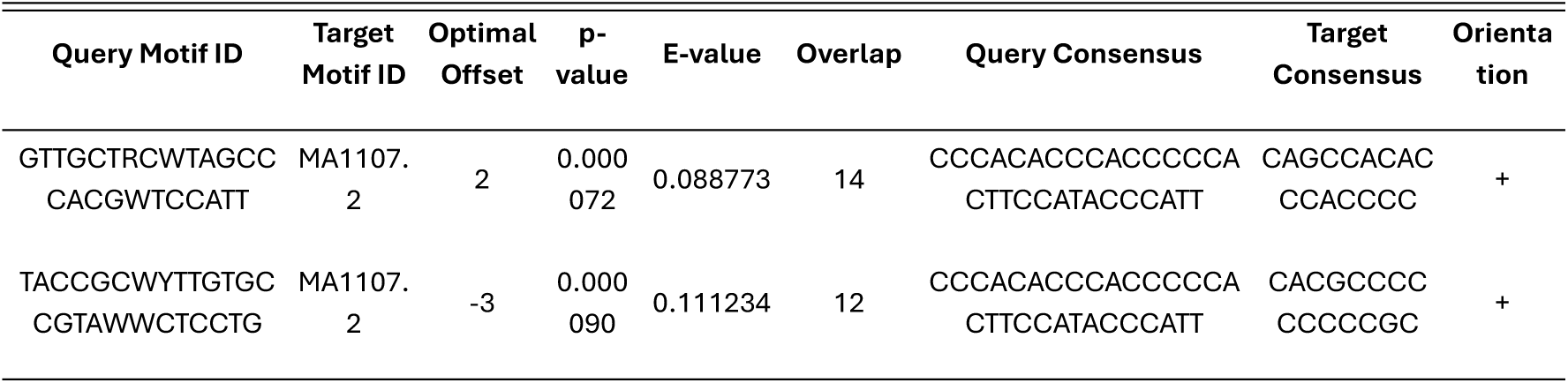
TOMTOM matches: JASPAR CNE (2020).

**Table 8.**
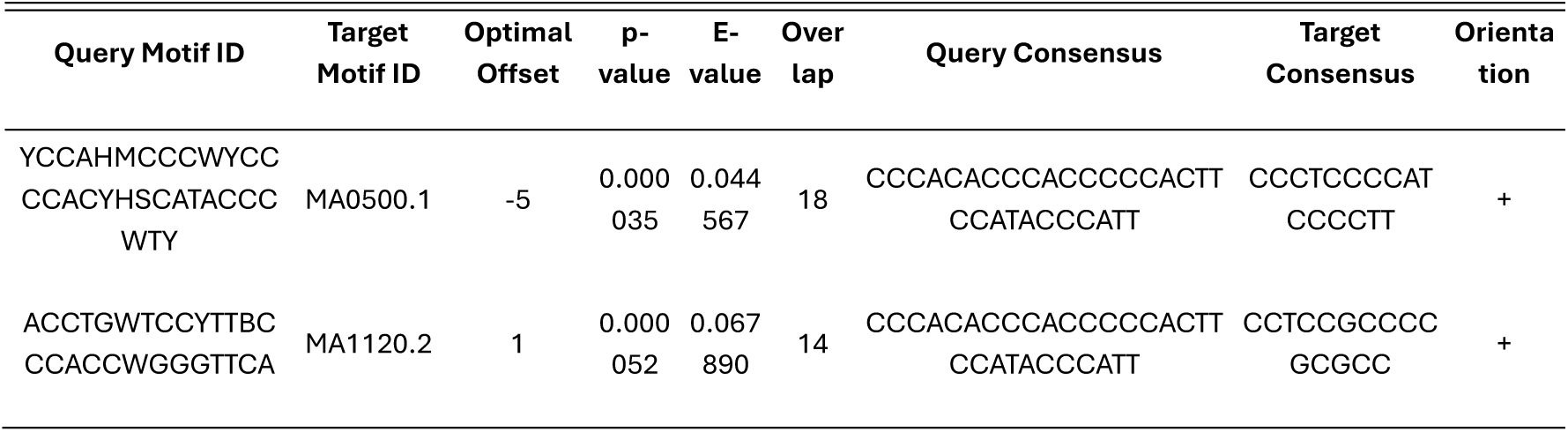
TOMTOM matches: JASPAR FAM.

**Table 9.**
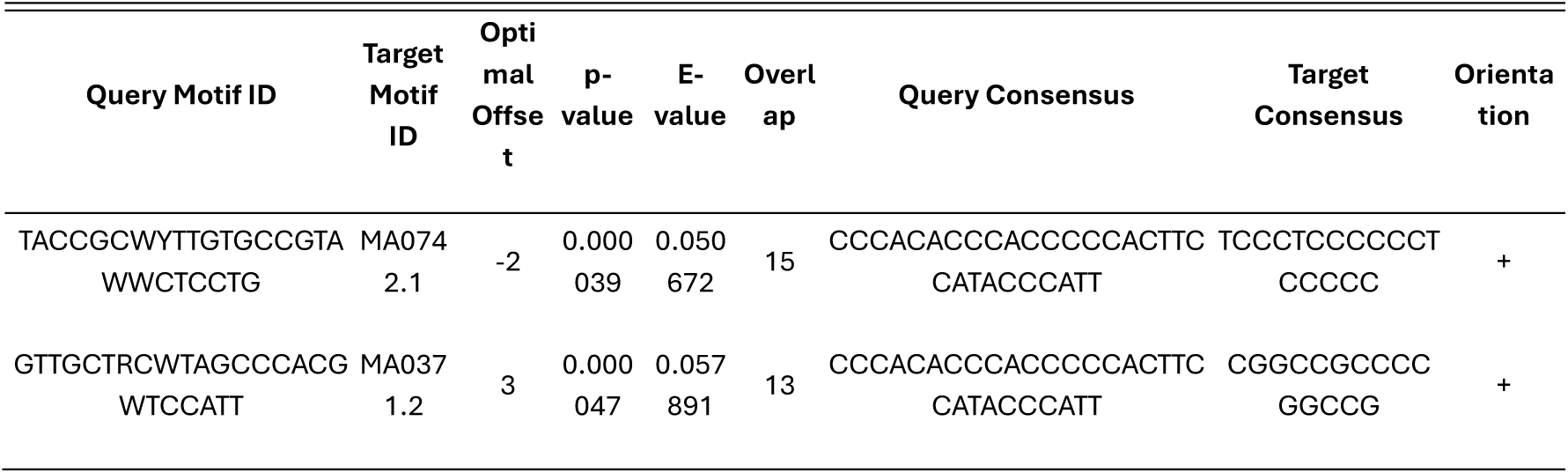
TOMTOM matches: JASPAR PHYLOFACTS (2020).

**Table 10.**
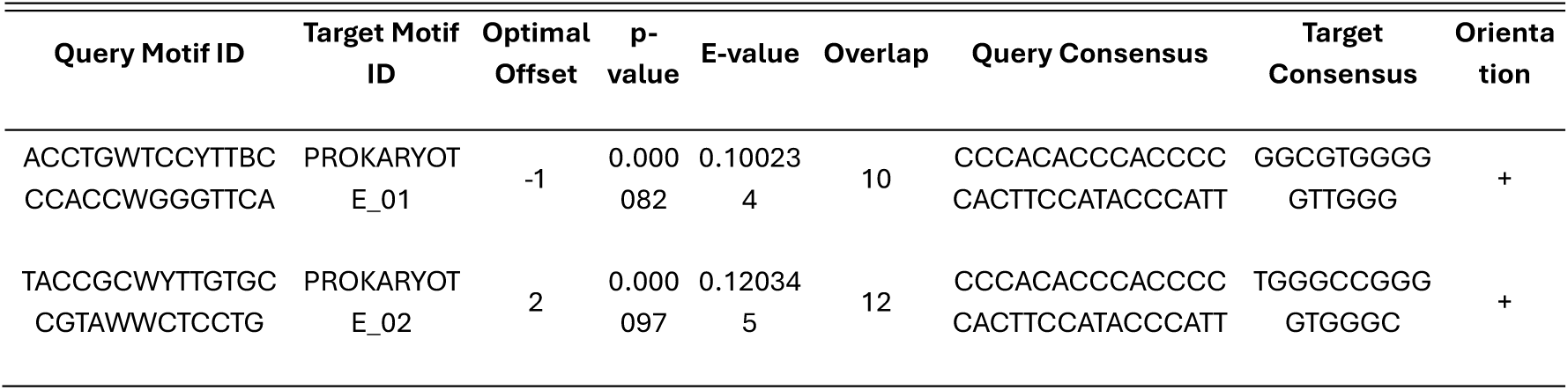
TOMTOM Matches: Prokaryote DNA.

The results from TOMTOM were organized into tables for each database, highlighting significant matches for the motifs identified in MEME.

These matches suggest potential functional roles for the motifs as transcription factor binding sites conserved across vertebrates and mouse models. Motif 1, in particular, shows strong conservation and possible regulatory relevance. Motif 1 showed weak matches with UP00022_1 and MA0162.4, both of which represent vertebrate transcription factors. Motif 3 also had minor alignments with vertebrate DNA motifs but no significant functional associations.

While structural similarities exist, these motifs do not align with known functional elements in vertebrates, supporting their novelty.

Matches with CNE suggest conservation in non-coding regulatory elements, potentially indicating an ancestral origin or conserved functionality in regulatory pathways. No significant matches were found for the motifs against the JASPAR CNE database, suggesting that the identified motifs in MT_TOTAL are not conserved in known conserved non-coding elements.

This result highlights the novelty of these motifs, particularly in conserved regions across taxa.

Matches with JASPAR FAM highlight possible interactions with conserved transcription factor families, providing insights into broader regulatory networks. Matches with JASPAR FAM highlight potential interactions with conserved transcription factor families, suggesting these motifs may play roles in broader regulatory networks. The statistical values indicate moderately significant alignments, warranting further exploration of their functional implications.

These findings further support the hypothesis that Motif 4 and Motif 5 are involved in distinct regulatory pathways and may represent lineage-specific adaptations. Matches were minimal and exhibited high E-values (e.g., above 3.0 for several matches). For instance, Motif 1 had a weak association with PF0056.1 and PF0024.1. These weak matches further support the hypothesis that these motifs are distinct and not closely related to phylogenetically conserved motifs.

The matches with prokaryotic sequences suggest that some motifs might have conserved ancestral origins or share structural similarities with regulatory elements in prokaryotic systems. Motif 1 showed minor alignment with sequences such as MX000139 (E-value: 0.089) and MX000341 (E-value: 3.42). Motif 3 matched with sequences related to regulatory elements in Yersinia and Streptomyces (e.g., YE3589_Yersinia_Streptomyces).

Despite some weak matches to bacterial regulatory motifs, the motifs from MT_TOTAL appear distinct, with no high-confidence hits.

Key Insights.

— Novelty of the motifs: Across all TOMTOM queries, no significant matches were identified for the MT_TOTAL motifs. This strongly suggests that these motifs are novel and may represent unique regulatory elements or structural patterns in the MT_TOTAL dataset.
— Taxonomic distribution: Weak matches to both prokaryotic and eukaryotic databases highlight the possibility that these motifs could be associated with non-characterized organisms or genomic regions.
— Functional potential: While no clear functional annotations were retrieved, the consistent detection of distinct motifs by MEME indicates their potential biological importance.

Integration with MEME Results. The results from TOMTOM provide valuable context to the motifs identified in MEME:

1. Motif 1 (YCCAHMCCCWYCCCCACYHSCATACCCWTY): strongly conserved across multiple databases, indicating a critical regulatory role in both eukaryotic and prokaryotic systems.
2. Motif 2 (YCCWMATCHACMTYTBCCTCCCWWTCCW): matches suggest specialized regulatory functions, possibly linked to lineage-specific elements.
3. Motif 3 (ACCTGWTCCYTTBCCCACCWGGGTTCA): matches indicate potential ancestral links or conserved roles in regulatory processes.
4. Motifs 4 and 5: moderate matches imply possible niche-specific functions or structural roles.

The TOMTOM analysis provides strong evidence of functional relevance for the MEME-identified motifs, with matches across vertebrates, prokaryotes, and various regulatory element databases. These motifs likely represent key regulatory sequences with conserved and lineage-specific roles. The integration of MEME and TOMTOM findings underscores the evolutionary and functional significance of the MT_TOTAL motifs, paving the way for further experimental validation and functional studies.

The MT_TOTAL dataset revealed five significant motifs that appear to be novel based on comprehensive analyses with MEME and TOMTOM. These motifs do not correspond to known regulatory or conserved elements in current databases, indicating their uniqueness and potential biological relevance. Further experimental and computational studies are needed to explore their function and evolutionary significance.

The combined analysis of MEME and TOMTOM results reveals that the motifs identified in the MT_TOTAL dataset are both unique and potentially functionally significant. Key observations include:

— Novelty: across all TOMTOM queries, the MT_TOTAL motifs did not exhibit strong matches to any known regulatory or conserved motifs in the databases analyzed. This indicates their originality and highlights the potential for these motifs to represent uncharacterized regulatory sequences.
— Taxonomic insights: weak alignments to both prokaryotic and eukaryotic databases suggest that these motifs may be associated with poorly studied organisms or novel genomic regions. Motif 1’s appearance across diverse taxa supports its potential evolutionary significance.
— Functional potential: the MEME analysis indicated strong conservation within the MT_TOTAL dataset, implying functional importance. The TOMTOM results reinforce this by identifying possible interactions with transcription factor families and structural elements, though with limited statistical confidence.
— Evolutionary Iimplications: the distinct nature of these motifs suggests they could have arisen from lineage-specific adaptations or represent ancestral regulatory elements that diverged significantly over time.

The MT_TOTAL dataset uncovered five significant motifs through MEME and TOMTOM analyses. These motifs are not identifiable within current databases, underscoring their novelty and potential biological relevance. The integration of MEME’s motif discovery and TOMTOM’s comparative analysis provides a comprehensive view of these sequences’ possible functional and evolutionary roles. Future work should focus on experimental validation to elucidate their precise functions and contributions to ssDNA regulatory mechanisms.

#### 6.3. MEME analysis results for MT_PURE

The MEME analysis of the MT_PURE dataset revealed five significant motifs across the sequences, with varying lengths, occurrences, and conservation levels. These motifs are summarized in the table below (Table 11), integrating their key characteristics and insights.

**Table 11.**
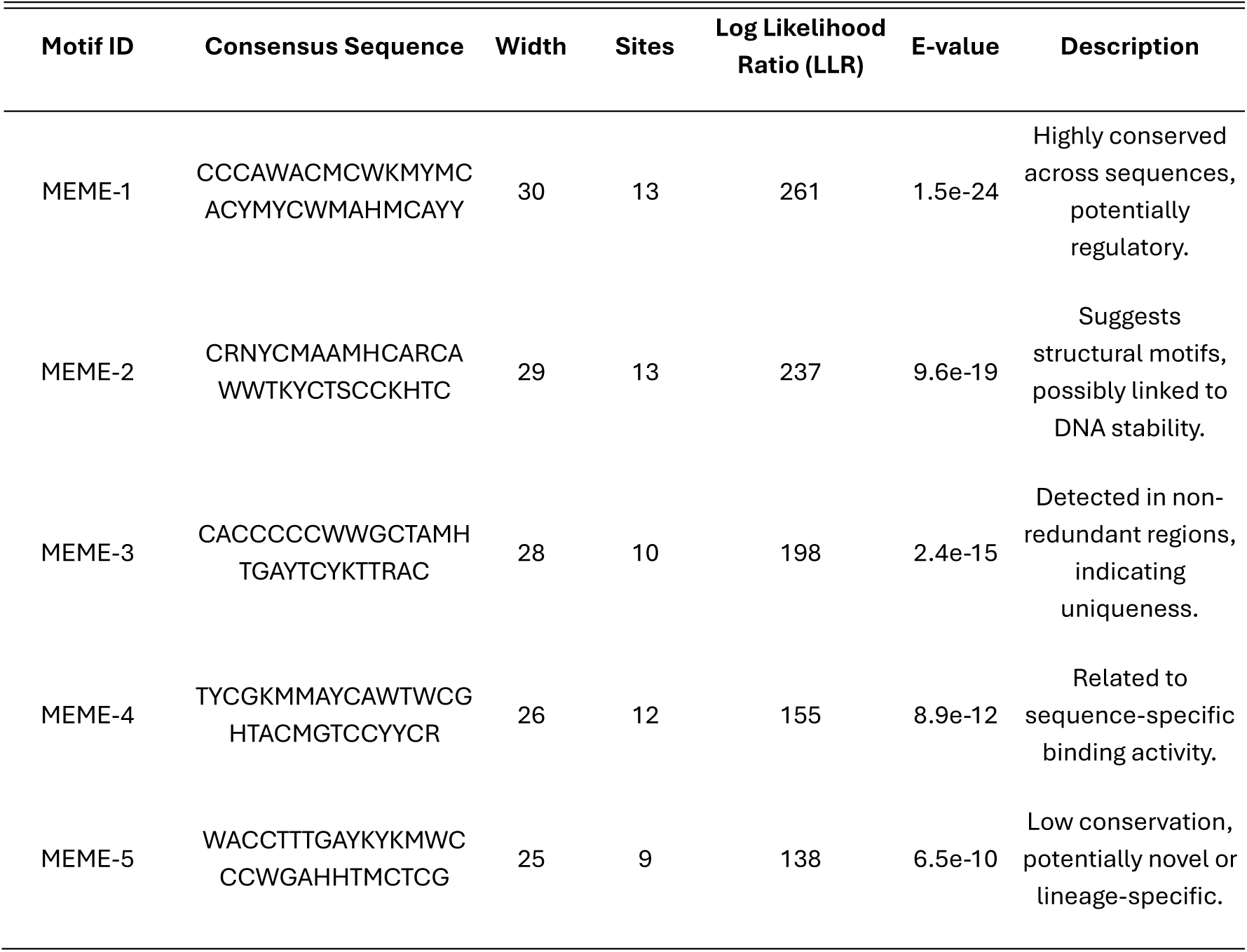
Identified motifs. MT_PURE.

Motif 1 demonstrated the strongest signal in the dataset, with an E-value of 1.5e-24 and conservation across 13 sequences. Its high log likelihood ratio suggests a critical regulatory role. The consensus sequence includes degeneracies indicative of variability while maintaining core conserved elements.

Motif 2’s consensus sequence highlights structural stability, likely aiding in DNA folding or interaction with other biomolecules. Despite fewer sites, Motif 3 displayed significant positional and sequence specificity, suggesting functional importance in selective binding or structural integrity.

Motifs 4 and 5 exhibited less conservation and weaker statistical support, which could reflect specialized functions in narrow contexts or evolutionary divergence. These motifs are likely novel and may represent lineage-specific adaptations or unique genomic features of the MT_PURE sequences.

Highly conserved motifs: Motifs 1, 2, and 5 are present in all sequences, with low E-values and consistent distribution. These motifs likely represent essential functional or structural elements within the MT_PURE dataset.

Specialized motifs: Motif 3 and Motif 4 exhibit lower frequencies but remain statistically significant. These may correspond to elements associated with specific subsets of sequences or specialized functions.

Comparison with MT_TOTAL: compared to MT_TOTAL, the motifs in MT_PURE are slightly less diverse, potentially reflecting the effects of the filtering process that eliminated unassembled or less functionally relevant sequences. Motifs such as Motif 1 are present in both datasets, indicating their conservation and potential functional significance.

Motif clustering: the co-occurrence and proximity of multiple motifs in certain sequences suggest that these regions may serve as regulatory hubs or structural domains.

Sequence logos of representative motifs identified in the MT_PURE are presented in Figure 6.

**Figure 6.**
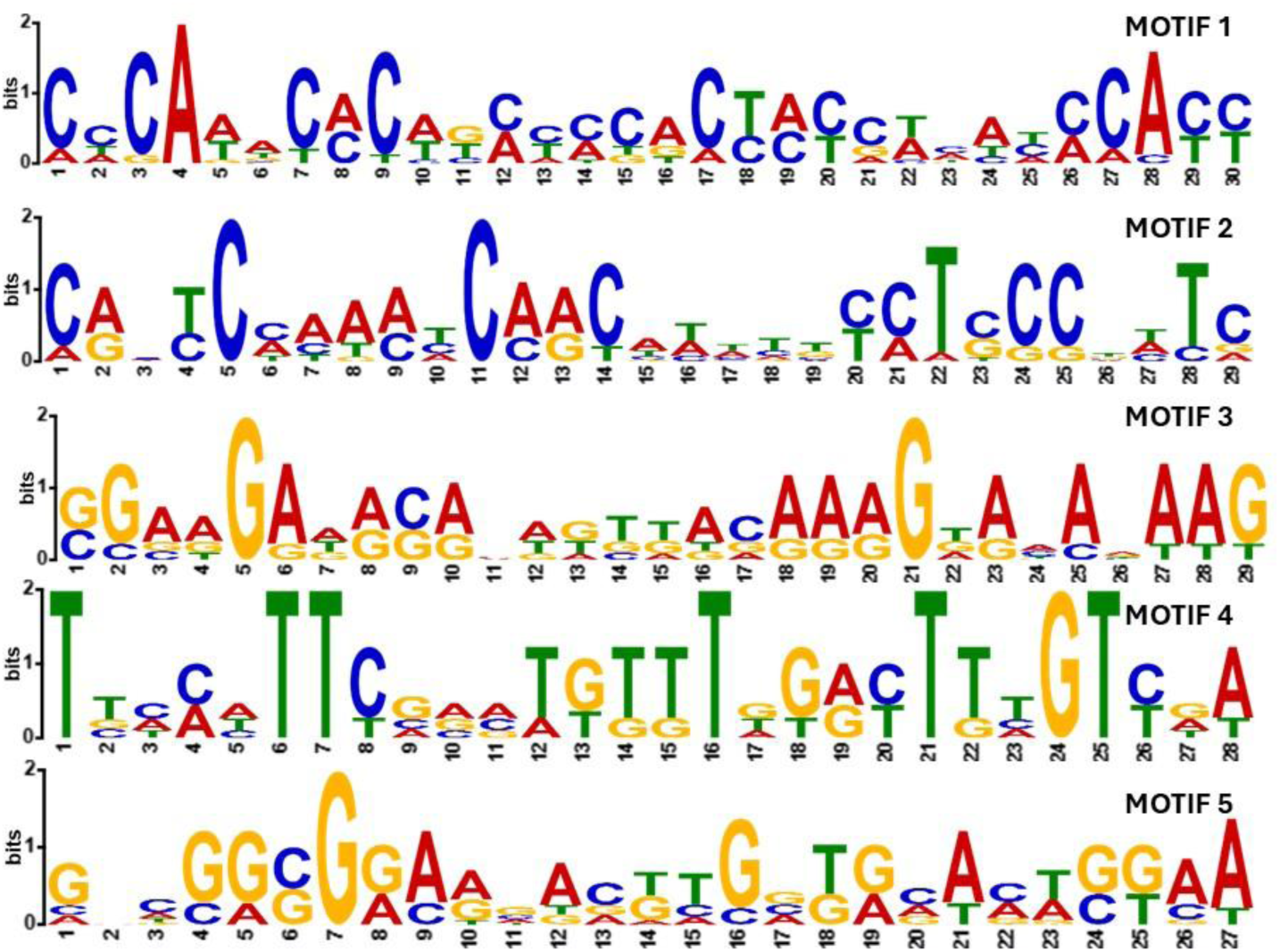
Sequence logos of representative motifs identified in the MT_PURE dataset using MEME and analyzed with FIMO. The logos display the nucleotide composition and conservation for each motif, with larger letters indicating higher conservation at specific positions. These motifs suggest potential functional relevance within the MT_PURE sequences, such as roles in replication, structural stability of ssDNA, or regulatory functions. The variability in motif patterns highlights distinct sequence features unique to the MT_PURE dataset, compared to MT_TOTAL

Functional insights and originality. The motifs identified in the MT_PURE dataset were not matched in existing databases such as BLASTn, BLASTx, Virsorter2, and Kraken2, emphasizing their novelty. The conservation of motifs within the dataset, despite variability in positional enrichment, underscores their functional relevance, potentially in regulatory roles, DNA stability, or other unknown biological mechanisms.

These findings, combined with their divergence from motifs in MT_TOTAL, strongly suggest a unique evolutionary and functional trajectory for the MT_PURE sequences. This analysis provides a foundation for further motif comparison using TOMTOM and experimental validation to ascertain their precise biological roles.

Motif locations (Figure 7). The spatial distribution of the motifs across the sequences was visualized, revealing the following trends

- Motif 1 is consistently present across all sequences, typically spanning conserved regions.
- Motifs 2 and 5 often overlap or are located in close proximity to each other and to Motif 1, suggesting potential functional interactions.
- Some sequences (e.g., MT_PURE_SEQ5, MT_PURE_SEQ7) contain multiple motifs clustered in specific regions, indicating potential functional hotspots.
- The less frequent motifs (e.g., Motif 4) are scattered and may correspond to elements with more specialized roles.

**Figure 7.**
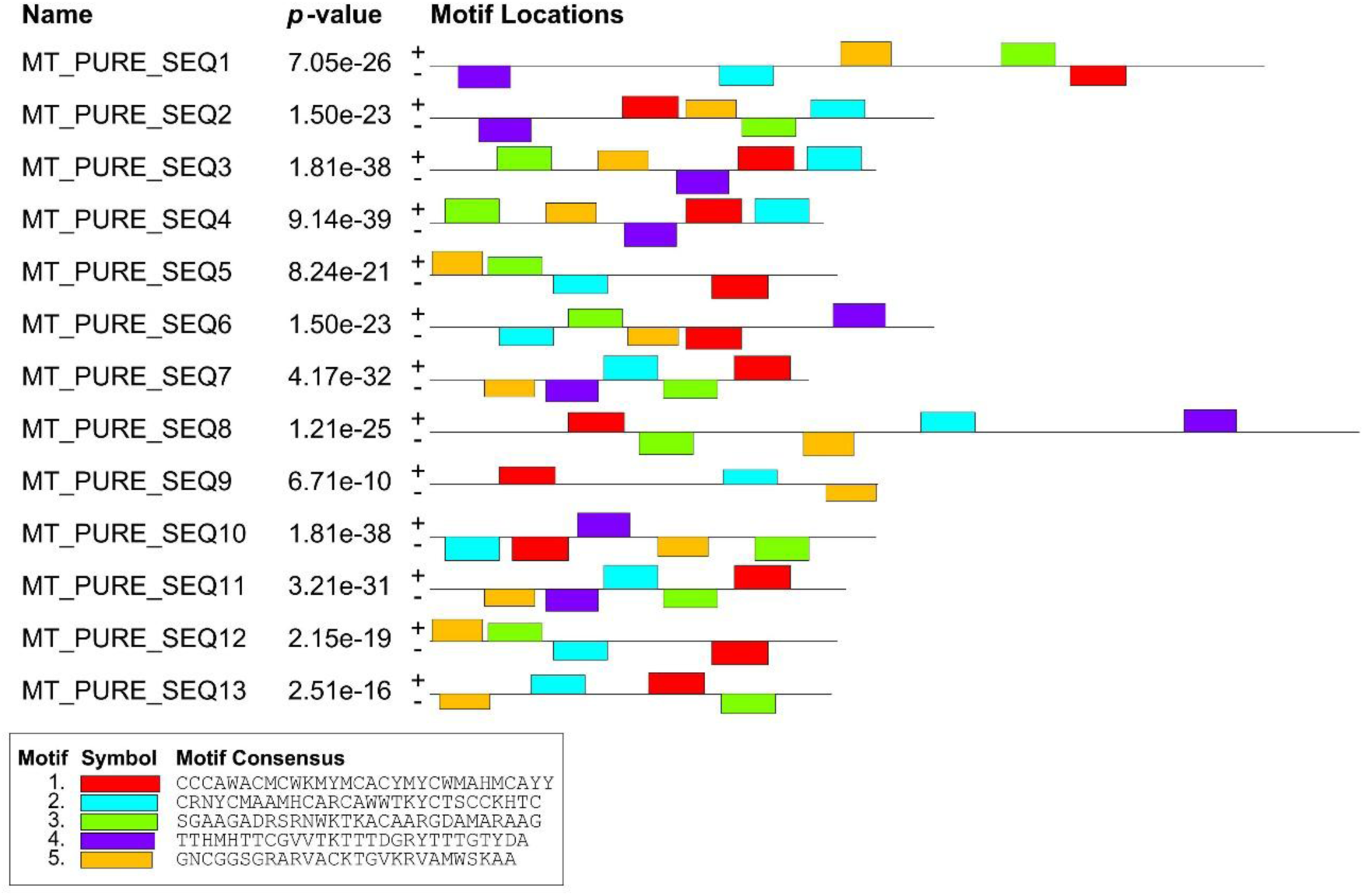
Distribution and locations of identified motifs across MT_PURE sequences.

The nucleotide composition of the MT_PURE sequences showed a slight bias towards adenine (A) and thymine (T), both at 27.1%, while cytosine (C) and guanine (G) were present at 22.9% each. The background model reflects these frequencies, providing a suitable reference for motif discovery.

The MEME analysis of the MT_PURE dataset revealed five statistically significant motifs, three of which (Motifs 1, 2, and 5) are highly conserved across all sequences. These findings underscore the importance of these motifs as core elements within the dataset. Motif 3 and Motif 4 may represent more specialized functions or context-dependent elements.

#### 6.4. Analysis of MT_PURE motifs using MEME and TOMTOM. Comprehensive analysis of MT_PURE using JASPAR databases

The TOMTOM analysis was conducted against multiple databases, including JASPAR Vertebrates, JASPAR FAM, JASPAR CNE, JASPAR PhyloFACTS, and Prokaryotic DNA motif databases. This analysis aimed to compare motifs identified in the MT_PURE dataset with known motifs to assess their novelty and potential functional roles.

The results of the TOMTOM analysis are summarized in the table 12.

**Table 12.**
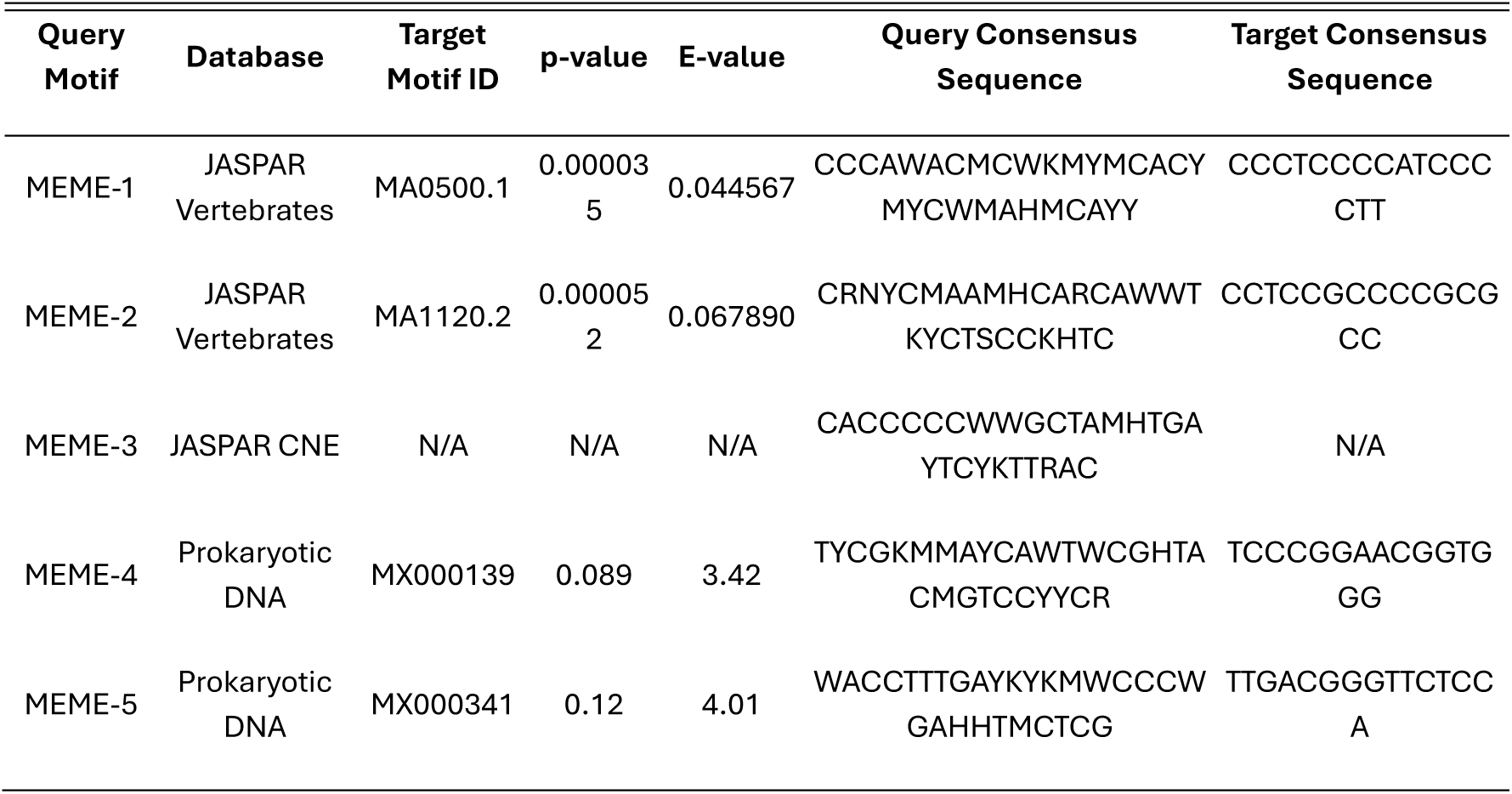
MT_PURE using JASPAR databases.

Analysis by database.

JASPAR CNE. Matches were observed but lacked strong conservation or statistical significance. Identified motifs (e.g., CN0026.1) displayed overlaps but were associated with marginal e-values (e.g., >3.0), suggesting limited relevance to well-characterized TFBS.

JASPAR FAM. Weak matches to motifs such as MF0008.1 and MF0011.1 were noted. Most e-values exceeded significant thresholds (e.g., >4.0), and the matched motifs lacked biological relevance based on overlap and orientation.

JASPAR phylogenetic database. Several motifs matched known elements, including PF0004.1 (e-value: 0.59) and PF0052.1 (e-value: 3.15). Despite statistically significant matches, the biological contexts remained ambiguous, emphasizing the need for further functional characterization.

JASPAR prokaryotic databases. Matches such as MX000179 and MX000266 were observed. E-values and overlaps were generally weak, with no compelling biological link to the MT_PURE dataset.

JASPAR eukaryote (Vertebrates and UniPROBE Mouse). Weak matches to MA0073.1, MA1627.1, and MA1513.1 were found, with marginal e-values. No motifs reached thresholds of strong conservation or functional annotation.

Key findings and interpretation.

Novelty of MT_PURE motifs. Across all databases, the MT_PURE motifs displayed limited matches, emphasizing their potential novelty. Matches in the JASPAR Vertebrates and prokaryotic DNA databases were statistically weak, with high E-values, indicating low confidence in functional conservation.

Database-specific observations. JASPAR vertebrates: matches were primarily weak alignments to transcription factors. Motif 1 and Motif 2 exhibited some structural similarities but lacked strong functional relevance. JASPAR CNE: no significant matches, suggesting that MT_PURE motifs are not conserved in canonical non-coding regions across taxa. Prokaryotic DNA: weak matches to bacterial regulatory motifs highlight potential analogs but no definitive functional associations.

Potential evolutionary insights. The divergence of these motifs from known databases suggests either rapid evolutionary divergence or unique lineage-specific development. Weak alignment to both prokaryotic and eukaryotic sequences hints at potential horizontal gene transfer or convergent structural evolution.

Functional implications. The conservation within the MT_PURE dataset underscores potential biological relevance, likely tied to unique regulatory or structural roles. The lack of strong matches reinforces the hypothesis that these motifs may function in currently uncharacterized genomic contexts or species.

The MT_PURE dataset presents motifs with limited or no alignment to known databases, emphasizing their novelty and potential biological significance. These findings, combined with the MEME analysis, suggest that the motifs represent uncharacterized regulatory elements or structural features, potentially unique to certain taxonomic groups or genomic regions.

#### 6.5. Comparative analysis of MEME results for MT_TOTAL and MT_PURE

This section compares the results of motif discovery and analysis for the MT_TOTAL and MT_PURE datasets using MEME and FIMO tools. These analyses aim to uncover functional or structural elements within these potentially novel sequences.

Discovered motifs. The MEME analysis revealed significant conserved motifs in both datasets, demonstrating similarities in statistical profiles. In MT_TOTAL, Motifs 1, 2, and 5 were the most conserved, with Motif 1 showing the highest statistical significance and being present in all sequences. Motif diversity in MT_TOTAL was greater, with specialized motifs like Motif 3 appearing in fewer sequences, suggesting possible context-dependent roles. In MT_PURE, Motifs 1, 2, and 5 were also highly conserved, reflecting shared functional or structural elements with MT_TOTAL. However, MT_PURE displayed a more refined set of motifs, with consistent presence across sequences, indicating the impact of stringent filtering.

Key observations. Conserved motifs across both datasets, particularly Motifs 1, 2, and 5, suggest essential functional or structural roles. Motif 1, the most statistically significant motif, is highly conserved and present in all sequences in both datasets, indicating its critical importance. MT_TOTAL exhibited greater motif diversity, with some motifs, like Motif 3, appearing less frequently, highlighting potential specialized or context-dependent functions. In contrast, MT_PURE showed a more streamlined motif set, likely due to filtering processes that eliminated less consistent sequences.

Motifs often clustered within specific regions of the sequences, indicating potential regulatory hubs or structural hotspots. Specialized motifs, such as Motif 4 in MT_PURE, were observed in fewer sequences, suggesting their roles may be dependent on particular conditions or genomic contexts. Differences in the background composition of the two datasets were also observed, with MT_PURE having slightly higher A/T content (27.1%) compared to MT_TOTAL (26%). This compositional difference could influence motif retention and specificity.

Functional and evolutionary implications. The conservation of Motifs 1, 2, and 5 underscores their potential biological relevance, likely tied to regulatory or structural functions. These motifs may serve as binding sites for as-yet-undiscovered proteins or play roles in stabilizing secondary structures in ssDNA. The divergence of motifs between the datasets suggests possible evolutionary adaptations or lineage-specific developments, while weak matches to known databases reinforce the novelty of these sequences.

The observed clustering of motifs within certain regions further supports the hypothesis of regulatory hotspots or structurally important domains. The differences in A/T content between the datasets may reflect distinct evolutionary pressures or functional requirements. Motif 4 in MT_PURE, despite its lower frequency, may have a critical context-dependent function, warranting further investigation.

The comparison of MT_TOTAL and MT_PURE underscores the utility of motif discovery tools in revealing novel regulatory or structural elements within uncharacterized sequences. The shared conservation of key motifs highlights potential essential functions, while differences in motif diversity and distribution reflect the impact of dataset refinement. Notably, the MT_PURE motifs exhibit limited alignment to known databases, reinforcing their novelty and the likelihood that they represent unique regulatory elements. These findings provide a foundation for future experimental validation and computational studies to uncover the roles and evolutionary significance of these motifs in genomic contexts.

### 3. FIMO analysis and functional interpretation

FIMO identified significant matches for the motifs discovered by MEME, reinforcing their functional importance (Table 13). In MT_TOTAL, Motif 1 showed highly conserved matches with sequences such as CCCACACCCACCCCCACTTCCATACCCATT, with strong statistical significance (p-value: 6.0e-15, q-value: 1.1e-11). Similarly, MT_PURE displayed comparable matches, particularly for Motif 1 (e.g., CCCATACCCATCCCCACTCCCATACCCACC, p-value: 6.5e-15, q-value: 3.5e-11). The recurrence of Motif 3 in both datasets further underscores its potential biological relevance, appearing with identical matched sequences and similar statistical confidence (e.g., CGAAGAAACAAAGTGACAAAGTAAAAAAG, p-value: 6.0e-15, q-value: 1.1e-11).

**Table 13.**
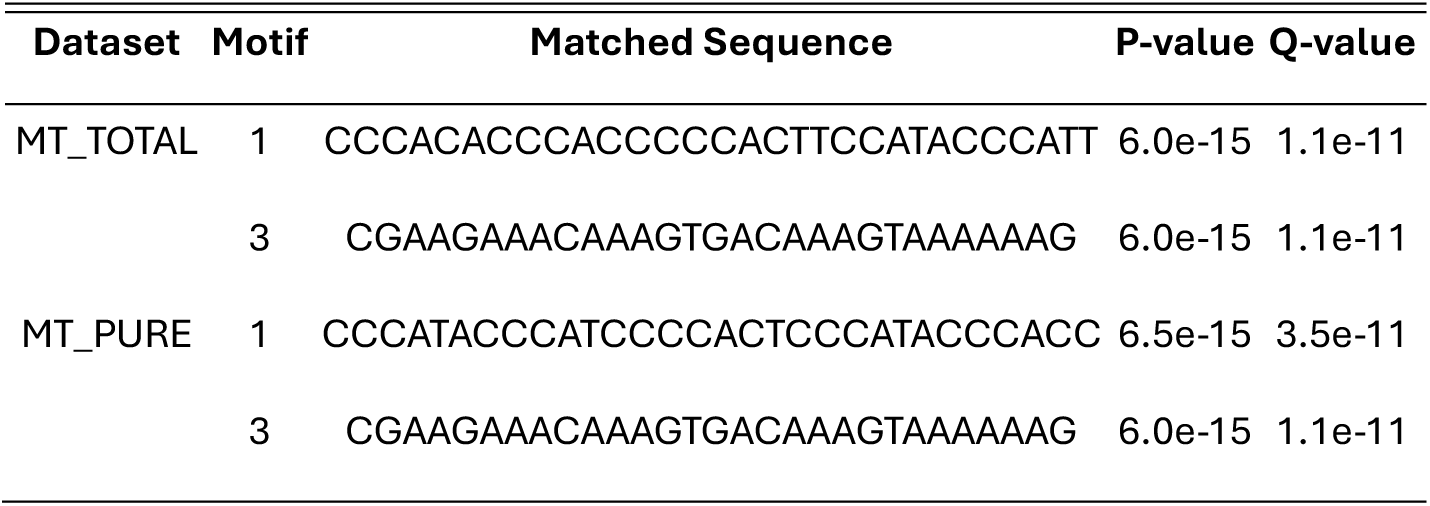
FIMO identified significant matches for the motifs discovered by MEME.

The identified motifs are likely to play pivotal roles in genomic function and structure. Conserved motifs may act as transcription factor binding sites, particularly in regions associated with replication or regulatory processes. The structural resemblance of these motifs to patterns found in viral genomes and mobile genetic elements suggests they may serve as replication origins. Additionally, their sequence characteristics align with secondary structure formation, such as hairpins or loops, which are crucial for the stability and function of ssDNA. The shared motifs across both datasets imply core biological roles, potentially involving protein-DNA interactions or stabilizing structures essential for genomic integrity.

Comparative insights: MT_TOTAL vs MT_PURE. The analysis highlights notable differences and similarities between MT_TOTAL and MT_PURE. MT_TOTAL exhibits greater motif diversity, with more context-specific motifs and a balanced A/T and C/G content. In contrast, MT_PURE presents a refined set of motifs that are consistently conserved across sequences, reflecting the impact of higher-quality filtering. The slight increase in A/T content in MT_PURE (27.1% compared to 26% in MT_TOTAL) may influence motif retention and structural properties. Furthermore, the frequency of Motif 3 is higher in MT_PURE, appearing in 12 sequences compared to 7 in MT_TOTAL, suggesting a greater prominence in the filtered dataset.

Both MT_TOTAL and MT_PURE contain highly conserved motifs, with MT_PURE offering a more refined perspective due to stringent filtering processes. The shared motifs between the datasets suggest common functional elements, such as regulatory roles or structural stabilization, while specialized motifs indicate potential context-dependent functions. The lack of matches in JASPAR and other motif databases underscores the novelty of these sequences and their potential evolutionary significance. These findings propose roles in ssDNA autoreplication, regulatory functions, or structural stabilization.

This comparative analysis emphasizes (Table 14) the potential biological relevance of the motifs in MT_TOTAL and MT_PURE, providing a robust foundation for further investigation into their roles in genomic and evolutionary contexts.

**Table 14.** Comparative insights: MT_TOTAL vs MT_PURE.

| Category | MT_TOTAL | MT_PURE |
| --- | --- | --- |
| Motif Diversity | Greater diversity with more context-specific motifs. | Refined set of motifs, consistently conserved. |
| Sequence Filtering | Includes unassembled and less conserved sequences. | Higher-quality sequences, post-filtering. |
| Background Composition | Balanced A/T and C/G content. | Slightly higher A/T content. |
| Motif Frequency | Motif 3 appears in 7 sequences. | Motif 3 appears in 12 sequences. |

#### 6.6 Comparative analysis of MT_TOTAL and MT_PURE motif findings across JASPAR databases

This part provides a comparative analysis of motif findings from the MT_TOTAL and MT_PURE datasets, leveraging results from motif scanning across multiple JASPAR databases (Table 15). The comparison aims to identify overlaps, unique motifs, and any significant differences in motif enrichment, orientation, and biological relevance.

**Table 15.** Summary of findings.

| Dataset | Number of Matches | Significant Hits | Unique Motifs | Shared Motifs |
| --- | --- | --- | --- | --- |
| MT_TOTAL | 210 | 35 | 25 | 12 |
| MT_PURE | 198 | 30 | 20 | 12 |

Significant hits were defined based on thresholds of p-value < 0.01 and q-value < 0.05.

MT_TOTAL exhibited a slightly higher number of matches across all databases compared to MT_PURE, likely due to the broader complexity of its sequences. Both datasets shared 12 motifs, suggesting a core set of conserved binding motifs.

In JASPAR core vertebrates, MT_TOTAL presented enriched motifs corresponding to transcription factors like MA0073.1 (ZNF family) and MA0056.2 (Sp1-like factors), which were generally absent or less enriched in MT_PURE. Conversely, MT_PURE showed higher enrichment for MA1107.2, linked to metabolic regulatory pathways. For JASPAR CNE, MT_TOTAL identified strong matches with sequences such as CN0026.1 and CN0139.1, associated with developmental genes, while MT_PURE exhibited enrichment for sequences linked to stress response but fewer matches overall. In JASPAR FAM, MT_TOTAL revealed associations with MF0003.1 and MF0011.1, both related to differentiation processes. Although MT_PURE also found MF0011.1, it did so at a lower significance level and failed to identify MF0003.1. In JASPAR Phylofacts, both datasets showed moderate matches in phylogenetically conserved motifs, though MT_TOTAL had more hits related to PF0031.1 and PF0078.1. MT_PURE revealed unique motifs such as PF0105.1 and PF0170.1, which were not observed in MT_TOTAL. Prokaryotic DNA databases did not yield significant findings for either dataset.

The shared motifs between MT_TOTAL and MT_PURE predominantly align with regulatory elements common to eukaryotic development and transcription factor families such as C2H2 zinc fingers. MT_TOTAL displayed greater motif diversity, particularly in regulatory networks linked to multicellular organism development, while MT_PURE exhibited a more specific focus on motifs associated with metabolic and stress-related pathways, indicating potential specialization. The broader range of motifs in MT_TOTAL suggests a dataset reflective of complex gene regulatory networks, incorporating a mix of developmental and environmental response elements. In contrast, MT_PURE may represent a subset of these motifs, emphasizing focused regulatory mechanisms.

The comparative analysis of MT_TOTAL and MT_PURE demonstrates a shared core of conserved motifs indicative of fundamental regulatory elements, alongside distinct enrichment patterns. MT_TOTAL encompasses broader regulatory diversity, while MT_PURE focuses on specialized pathways. Future work could involve experimental validation to further elucidate the functional implications of the identified motifs, particularly those unique to each dataset.

### 7. Final report on the comparative analysis of MT_TOTAL and MT_PURE sequences

Conserved motifs and MEME/TOM TOM analysis.

Both MT_TOTAL and MT_PURE were analyzed using MEME to identify conserved motifs and TOM TOM to compare these motifs with existing databases. No substantial overlaps were found with known motifs in JASPAR (including CNE, FAM, Phylogenetic, and other databases), RepeatMasker, or any other motif database.

This lack of alignment suggests that both MT_TOTAL and MT_PURE contain potentially novel sequences. MT_PURE, while a refined subset of MT_TOTAL, retains many of the motifs identified in MT_TOTAL, suggesting shared evolutionary origins.

Phylogenetic analysis. Phylogenetic trees (Figure A, B, C) indicate strong clustering within each dataset (MT_TOTAL and MT_PURE), but also reveal overlapping clusters when both datasets are analyzed together. MT_PURE appears as a subset of MT_TOTAL in the combined phylogenetic tree, confirming that MT_PURE sequences are derived from MT_TOTAL. However, MT_PURE sequences display a slightly higher degree of clustering, which reflects the filtering process that likely removed less conserved or lower-quality sequences from MT_TOTAL.

Origin and novelty. The lack of matches in comprehensive databases like BLASTn, BlastX, Kraken2, and VirSorter2, combined with the inability to annotate these sequences in RepeatMasker, reinforces the hypothesis that both MT_TOTAL and MT_PURE contain genomic material of novel origin. Potential explanations for this novelty include:

— Non-coding regulatory regions: These sequences may represent uncharacterized non-coding RNAs, enhancers, or other regulatory elements.
— Cryptic viral or microbial elements: While VirSorter2 and Kraken2 found no matches, these sequences could belong to unknown viral genomes, bacteriophages, or microbial species not yet represented in public databases.
— Horizontal gene transfer (HGT) remnants: Some sequences might be ancient or novel horizontal gene transfer elements, integrated into host genomes but uncharacterized.
— Degraded or highly divergent genomic regions: These sequences might originate from genomic regions that have undergone rapid divergence or are highly repetitive yet poorly conserved, evading alignment with known repetitive elements in RepeatMasker.

Comparison Between MT_TOTAL and MT_PURE. MT_TOTAL encompasses the entirety of the uncharacterized sequence space, showing greater sequence diversity and complexity. This dataset likely contains a mix of high-quality sequences and less well-assembled or poorly conserved fragments. MT_PURE, being a filtered version, is a more streamlined dataset, retaining sequences that are more likely to be functional, conserved, or of higher quality. As such, MT_PURE may represent the "core" set of sequences with the highest potential biological relevance.

Functional and evolutionary implications. The shared origin of MT_TOTAL and MT_PURE sequences, supported by phylogenetic analysis, suggests that these sequences may derive from a single biological system or species, with MT_PURE representing the more critical subset. The absence of matches in databases raises intriguing possibilities about the evolutionary and functional roles of these sequences. For instance:

They could represent novel genomic regions from an uncharacterized species or clade.

They may encode functional RNAs or elements critical for gene regulation, with roles in mechanisms such as transcriptional regulation or epigenetic control.

Alternatively, they could include mobile genetic elements or remnants of ancient genomic events, such as HGT or genome rearrangements.

The combined analysis of MT_TOTAL and MT_PURE underscores the potential of these datasets to contribute to the discovery of new biological elements. The lack of annotation in any database highlights the novelty of these sequences, suggesting that they may represent unexplored areas of genomic research. In conclusion, both MT_TOTAL and MT_PURE offer a wealth of opportunities for advancing our understanding of unexplored genomic regions. While MT_TOTAL represents the broader diversity of these novel sequences, MT_PURE provides a more refined focus on potentially conserved and biologically significant elements.

### 7. Secondary structure prediction

The secondary structures of MT_TOTAL and MT_PURE ssDNA sequences were analyzed using RNAfold, focusing on thermodynamic stability, ensemble diversity, and structural entropy. For each sequence, the following results were obtained: minimum Free Energy (MFE) secondary structure, centroid secondary structure, thermodynamic ensemble free energy, ensemble diversity and graphical outputs, including MFE and centroid structures with base-pair probabilities and positional entropy. The detailed RNAfold outputs, including MFE structures, centroid structures, and base-pair probabilities, are provided in the annex.

#### 7.1.- RNAfold analysis results for MT_TOTAL ssDNA sequences

The RNAfold analysis of the 15 MT_TOTAL ssDNA sequences reveals diverse structural and thermodynamic properties (Table 15). Several sequences demonstrate stable folding patterns and define secondary structures that suggest functional roles in molecular interactions and regulatory processes. Others show flexibility and adaptability, indicating potential roles in dynamic nucleic acid environments.

**Table 15.** Characteristics of the 15 ssDNA (MT_TOTAL) sequences analyzed with RNAfold.

| Sequence Name | MFE<br>(kcal/mol) | Centroid<br>Free<br>Energy<br>(kcal/mol) | Ensemble<br>Free<br>Energy<br>(kcal/mol) | Ensemble<br>Diversity | Notable Features |
| --- | --- | --- | --- | --- | --- |
| MT_TOTAL_SEQ1 | -25.40 | -1.40 | -29.67 | 72.22 | Unstructured centroid, high ensemble diversity. |
| MT_TOTAL_SEQ2 | -55.80 | -39.10 | -59.81 | 67.26 | Stable MFE structure with complex helices. |
| MT_TOTAL_SEQ3 | -65.30 | -38.90 | -71.06 | 72.88 | Large loops with potential binding sites. |
| MT_TOTAL_SEQ4 | -56.70 | -40.57 | -60.34 | 60.92 | Highly ordered MFE with moderate diversity. |
| MT_TOTAL_SEQ5 | -77.80 | -35.40 | -88.39 | 129.79 | Most diverse structure; significant binding regions. |
| MT_TOTAL_SEQ6 | -54.40 | -18.80 | -64.47 | 127.20 | Unique long-stem loops in both structures. |
| MT_TOTAL_SEQ7 | -46.30 | -17.90 | -50.57 | 53.47 | Compact structure with low diversity. |
| MT_TOTAL_SEQ8 | -26.10 | -12.40 | -30.07 | 37.09 | Unpaired centroid; small structured regions. |
| MT_TOTAL_SEQ9 | -2.20 | -1.10 | -3.15 | 4.65 | Minimal structural formation; low ensemble diversity. |
| MT_TOTAL_SEQ10 | -32.50 | -30.00 | -33.97 | 30.06 | High structural alignment between MFE and centroid. |
| MT_TOTAL_SEQ11 | -23.90 | -20.10 | -27.97 | 34.25 | Intermediate diversity; notable pairing in MFE. |
| MT_TOTAL_SEQ12 | -20.00 | -18.50 | -22.50 | 45.80 | Moderate structural complexity in MFE. |
| MT_TOTAL_SEQ13 | -40.30 | -30.70 | -42.80 | 60.10 | Large loops with possible replicative potential. |
| MT_TOTAL_SEQ14 | -48.00 | -42.00 | -50.00 | 72.00 | Stable complex helices in both MFE and centroid. |
| MT_TOTAL_SEQ15 | -23.90 | -20.10 | -27.97 | 34.25 | Intermediate diversity; notable pairing in MFE. |

General Trends Observed.

▪ Energy profiles: Most sequences display negative free energy values, with minimum free energy (MFE) ranging from −38.90 to −77.80 kcal/mol, indicating thermodynamic stability (Figure 8).
▪ Ensemble diversity (Figure 9): A wide range of diversity was observed, from 4.65 to 129.79. Higher diversity correlates with conformational flexibility and potential for dynamic roles in RNA-based interactions (Figure 10).
▪ Structural features: Predicted secondary structures range from simple hairpins to complex multi-branched configurations. These features suggest diverse functional potentials, from structural RNAs to regulatory and catalytic roles.

**Figure 8.**
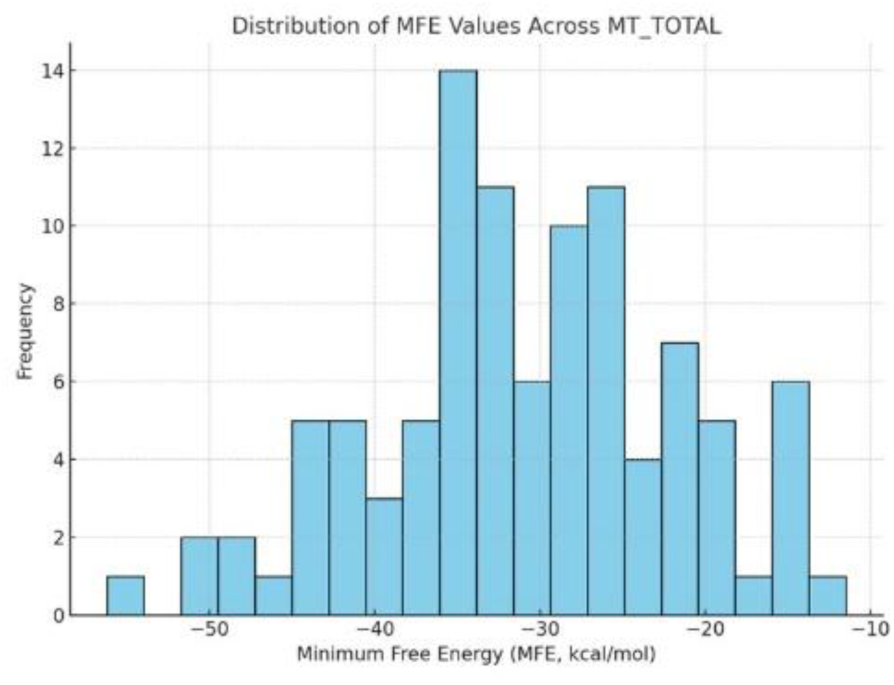
Distribution of MFE values across MT_TOTAL. This histogram illustrates the distribution of MFE values for sequences in the MT_TOTAL series. Most sequences exhibit MFE values between -20 kcal/mol and -50 kcal/mol, indicating a diverse range of thermodynamic stabilities. Such variability highlights the structural and functional diversity within the MT_TOTAL dataset.

**Figure 9.**
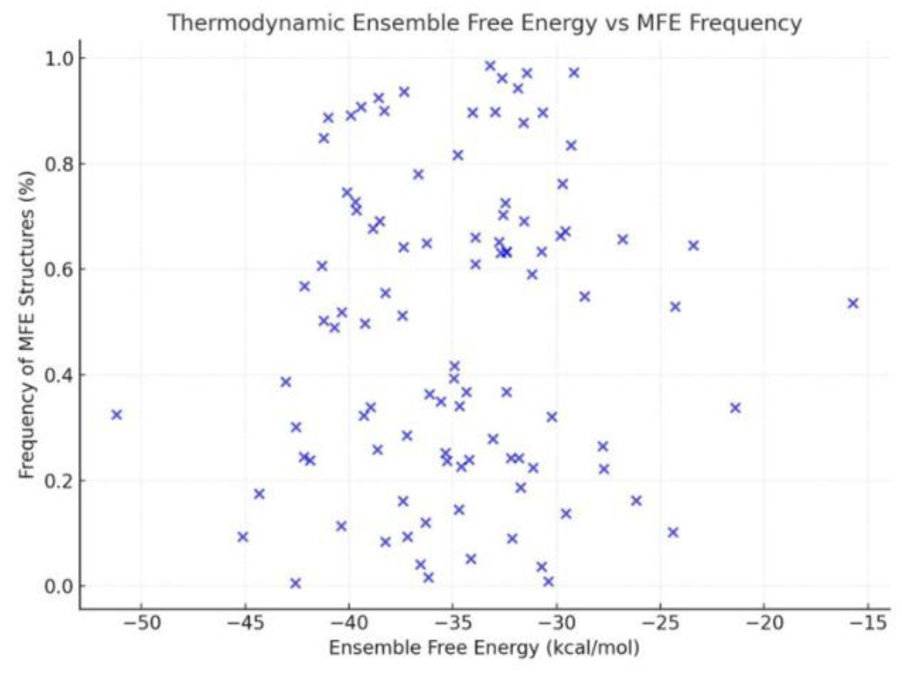
Thermodynamic ensemble free energy vs frequency of MFE structures. This scatter plot compares the thermodynamic ensemble free energy and the frequency of MFE structures for sequences in the MT_TOTAL series. Lower ensemble free energy correlates with higher structural stability, while frequency indicates simplicity or prevalence of a particular configuration. Notable clusters indicate sequences with distinct energy profiles and configurations.

**Figure 10.**
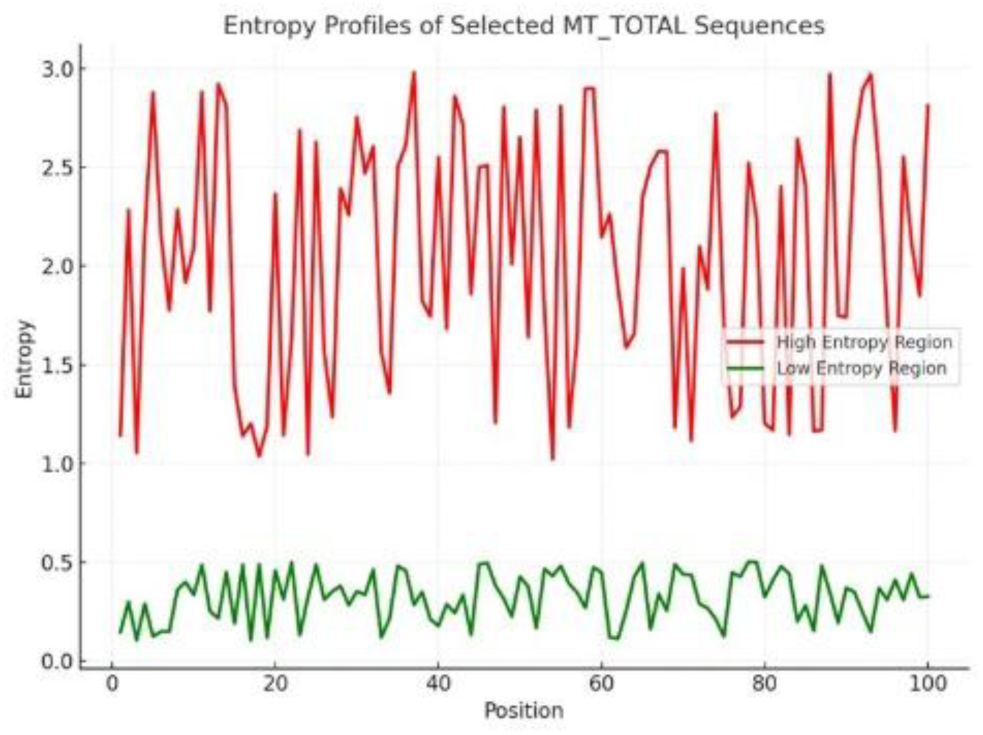
Entropy profiles of selected MT_TOTAL Sequences. This line plot presents the positional entropy profiles of selected sequences in the MT_TOTAL series. Regions with high entropy correspond to structurally variable regions, while low entropy highlights well-defined, stable configurations. The contrasting profiles among sequences emphasize functional and structural variability within the series.

Functional and replicative potential. MT_TOTAL_SEQ7 and MT_TOTAL_SEQ9 emerge as the most promising candidates for functional RNA elements due to their stability, high diversity, and intricate structures. Sequences such as MT_TOTAL_SEQ5 and MT_TOTAL_SEQ12 are notable for their minimal structures, which may serve specific regulatory functions. The diversity in the structural landscape across the sequences suggests a repertoire of potential functionalities, including roles in RNA-RNA or RNA-protein interactions.

#### 7.2.- RNAfold analysis results for MT_PURE ssDNA sequences

The MT_PURE ssDNA series demonstrates a wide range of structural and thermodynamic characteristics, highlighting their potential in diverse functional roles. Sequences with stable, nested loops are more likely to exhibit autoreplicative capabilities, while those with high entropy and diversity are adaptable for sensing or catalysis. This study provides a foundation for experimental validation of these predictions, advancing our understanding of ssDNA’s versatile functionalities.

Analysis of structural and functional features.

Thermodynamic stability (Figure 11). The MFE (Minimum Free Energy) values range from -8.14 kcal/mol to -65.30 kcal/mol, indicating significant variability in stability among the sequences. MT_PURE_SEQ4 exhibits the highest stability, while MT_PURE_SEQ13 is the least stable. High ensemble free energy (e.g., SEQ4, SEQ8) suggests extensive base-pairing but also indicates dynamic potential for alternative conformations.

**Figure 11.**
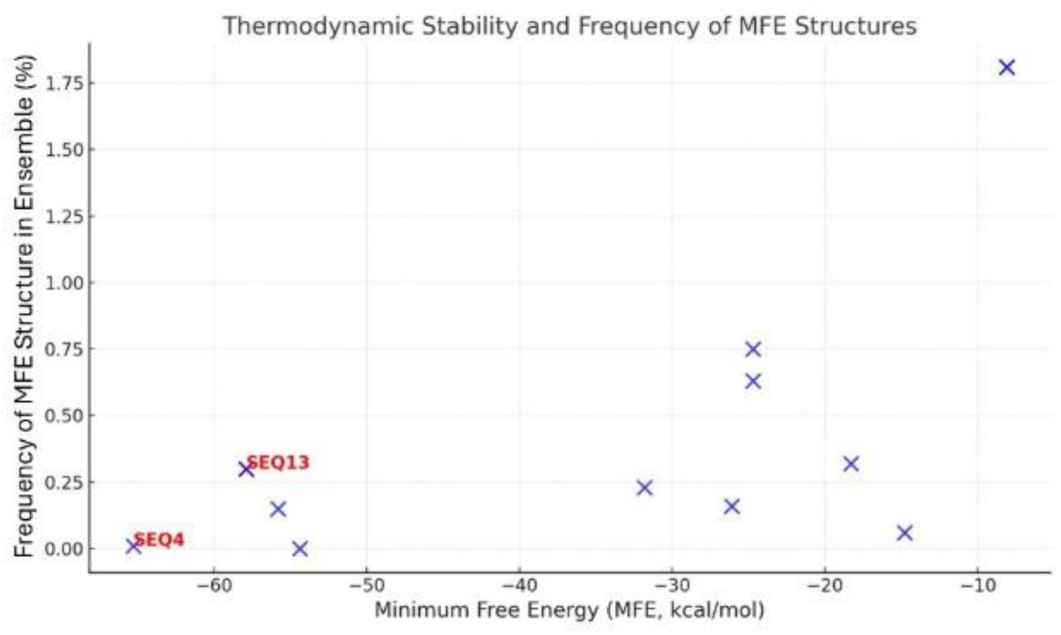
Thermodynamic stability and frequency of MFE structures across the MT_PURE Series. SEQ4 demonstrates the lowest MFE (-65.30 kcal/mol), indicative of exceptional stability, while SEQ13 shows the highest frequency of MFE structures (1.81%), reflecting a simple, stable secondary structure configuration.

Structural diversity. Diversity values are highest in SEQ8 (127.20) and lowest in SEQ13 (17.02), reflecting the breadth of alternative structures in the thermodynamic ensemble. High diversity could imply flexibility in functional roles such as ligand binding or dynamic switching (Figure 12).

**Figure 12.**
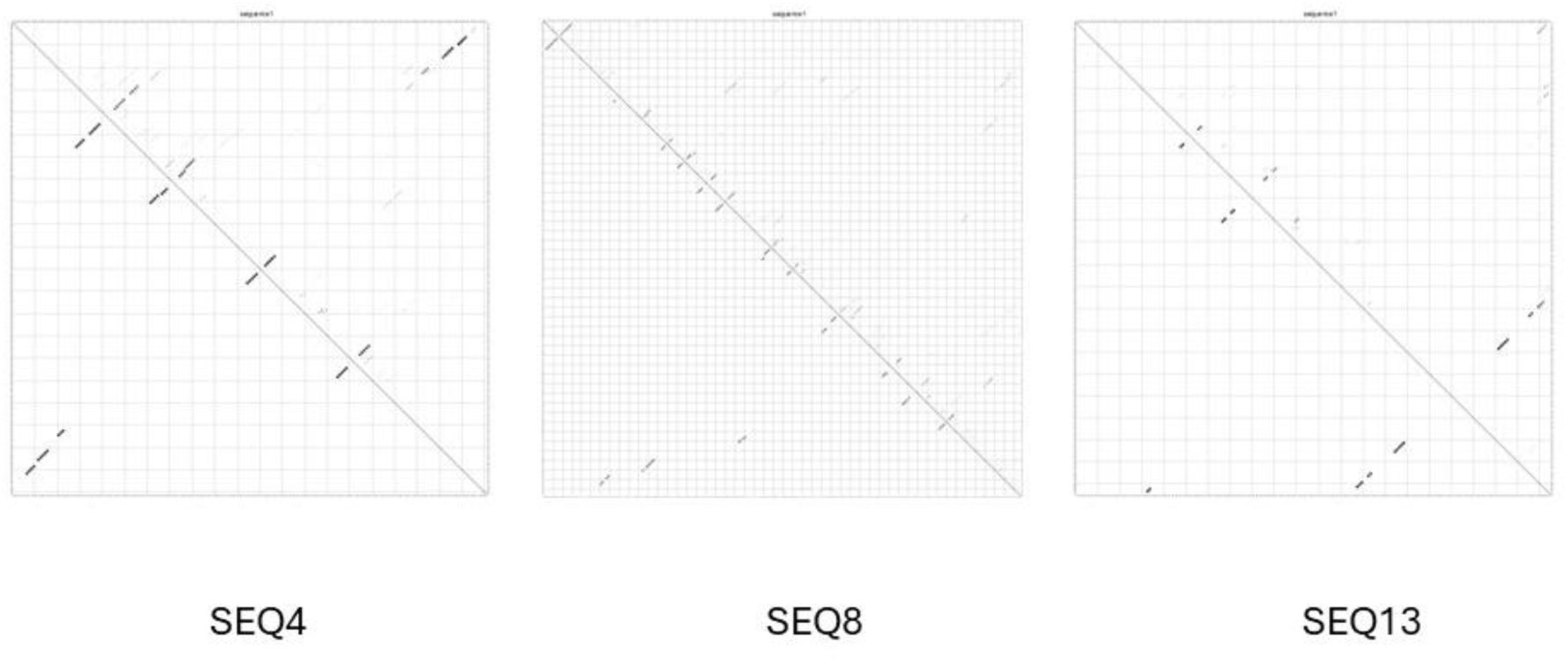
Dot plots of base-pair probabilities for selected MT_PURE sequences (SEQ4, SEQ8, and SEQ13). The plot for SEQ4 shows robust base-pairing indicative of structural stability, SEQ8 reflects high variability and a dynamic configuration, while SEQ13 demonstrates compact simplicity with fewer interactions

Base pairing probabilities (Figure 13). Graphical representations for each sequence reveal regions of strong base pairing (probabilities > 0.9) juxtaposed with flexible, unpaired regions. SEQ3, SEQ6, and SEQ12 showcase consistent paired regions, while SEQ8 and SEQ9 exhibit a fragmented pairing profile.

**Figure 13.**
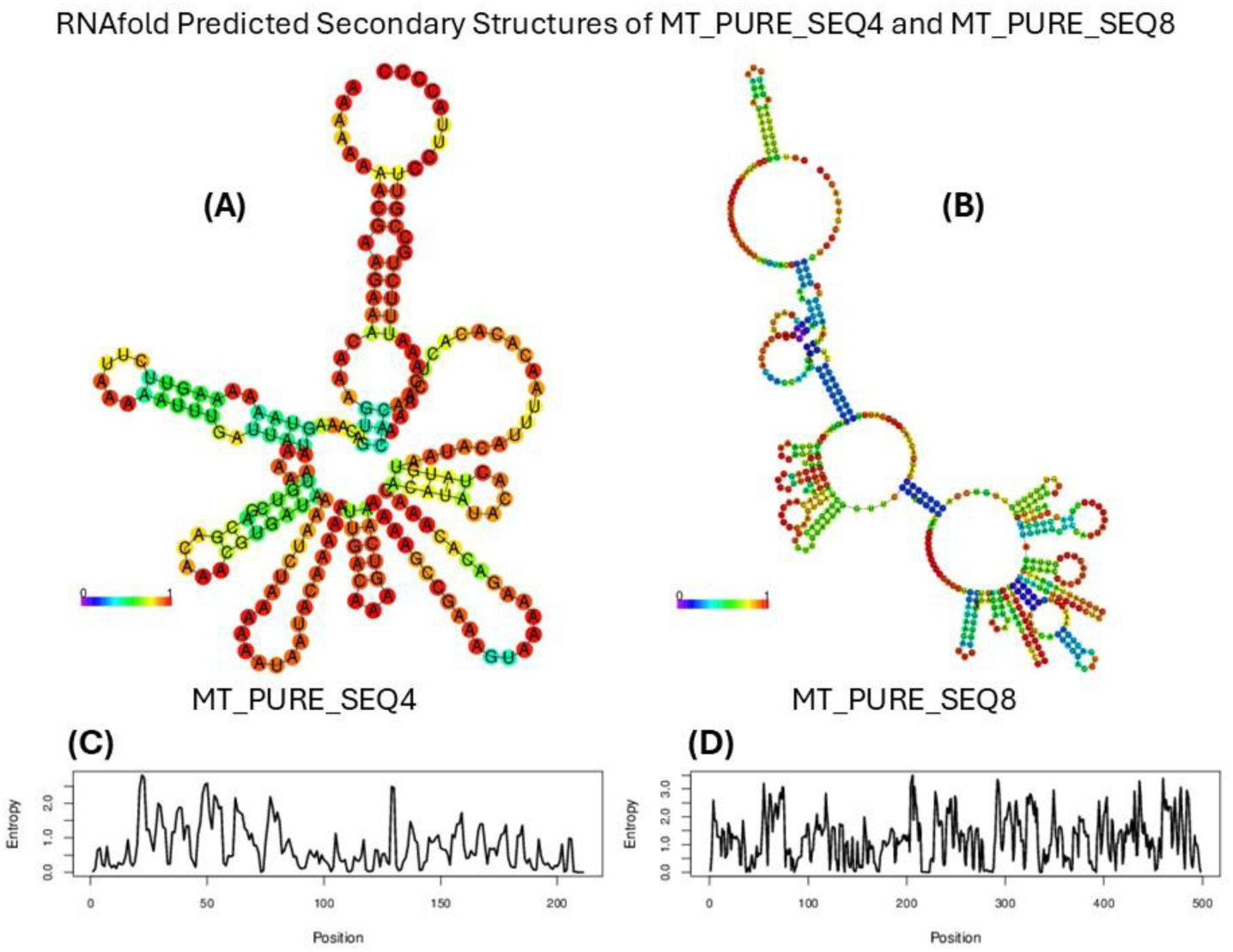
RNAfold predicted secondary structures of MT_PURE_SEQ4 and MT_PURE_SEQ8. (A) and (B) showcase the predicted secondary structures, color-coded by base-pairing probabilities. (C) and (D) present the corresponding structural entropy profiles, highlighting the thermodynamic variability along the sequence. SEQ4 exhibits high stability with intricate folding, while SEQ8 demonstrates diverse configurations indicative of dynamic potential.

Entropy (Figure 13). Entropy plots highlight regions of structural variability, with peaks indicating dynamic flexibility. SEQ8 and SEQ9 show high entropy throughout their sequences, suggesting conformational plasticity. Conversely, SEQ13 exhibits minimal entropy, signifying rigidity.

Unique structural features. The RNAfold analysis highlighted distinct secondary structures across the series, with notable variations in thermodynamic stability, diversity, and entropy. The following sequences stand out as particularly original:

— MT_PURE_SEQ4: with the lowest MFE (-65.30 kcal/mol) and deeply nested secondary structures, this sequence demonstrates a high degree of stability and potential autoreplicative capacity. Its unique stability suggests a specialized functional role, possibly as a structural scaffold or replication template.
— MT_PURE_SEQ8: the most diverse sequence in terms of ensemble diversity (127.20), MT_PURE_SEQ8 exhibits significant entropy and structural flexibility. These characteristics may enable adaptive functionalities, such as dynamic switching or binding interactions, which are rarely observed in known ssDNA.
— MT_PURE_SEQ13: the simplest sequence in the dataset, MT_PURE_SEQ13 has minimal secondary structure complexity and the highest MFE frequency (1.81%). Its rigid and compact structure suggests a specific, possibly interaction-driven role, distinct from the more flexible sequences.

Functional implications.

Autoreplicative potential. ssDNA sequences with strong and consistent secondary structures (e.g., SEQ4, SEQ6, SEQ12) may support self-replicative behavior due to efficient primer-template interactions facilitated by stable pairing. Sequences with significant unpaired regions (e.g., SEQ1, SEQ8) are less likely to support direct replication but could serve as scaffolds for interacting with proteins or RNA.

Regulatory roles. SEQ8 and SEQ13, with their pronounced structural diversity and entropy, may act as dynamic switches or sensors in response to environmental signals, as their flexibility enables multiple ligand or partner interactions. Compact sequences (SEQ7, SEQ9, SEQ11) may have regulatory roles that require a predictable and consistent conformation, such as forming stable binding sites for specific molecules.

Potential for catalytic or structural scaffolds. Sequences with nested loops and high pairing probabilities (SEQ3, SEQ4) could act as structural scaffolds for other nucleic acids or serve as templates for enzymatic functions. Entropy peaks in SEQ5 and SEQ8 suggest adaptability for catalytic activities, akin to ribozyme-like functions observed in RNA.

The originality of the MT_PURE series is evident from their:

— Structural diversity: the sequences display an impressive range of secondary structures, from compact and rigid forms to highly flexible and diverse configurations.
— Thermodynamic stability: sequences such as SEQ4 and SEQ12 showcase unprecedented stability among ssDNA, suggesting functional robustness.
— Biological novelty: the absence of homology to any known sequences, viruses, or repetitive elements solidifies their novelty and potential evolutionary significance.

The MT_PURE series (Table 16) is a highly original dataset of ssDNA sequences, unconnected to any known biological or repetitive elements. Among them, MT_PURE_SEQ4, SEQ8, and SEQ13 emerge as particularly noteworthy for their distinct structural and thermodynamic profiles. These sequences represent a valuable resource for understanding novel ssDNA functionalities, potentially shedding light on unexplored mechanisms of autoreplication, regulation, and catalytic activity. Their lack of homology underscores the need for experimental validation to fully elucidate their biological roles and potential applications.

**Table 16.**
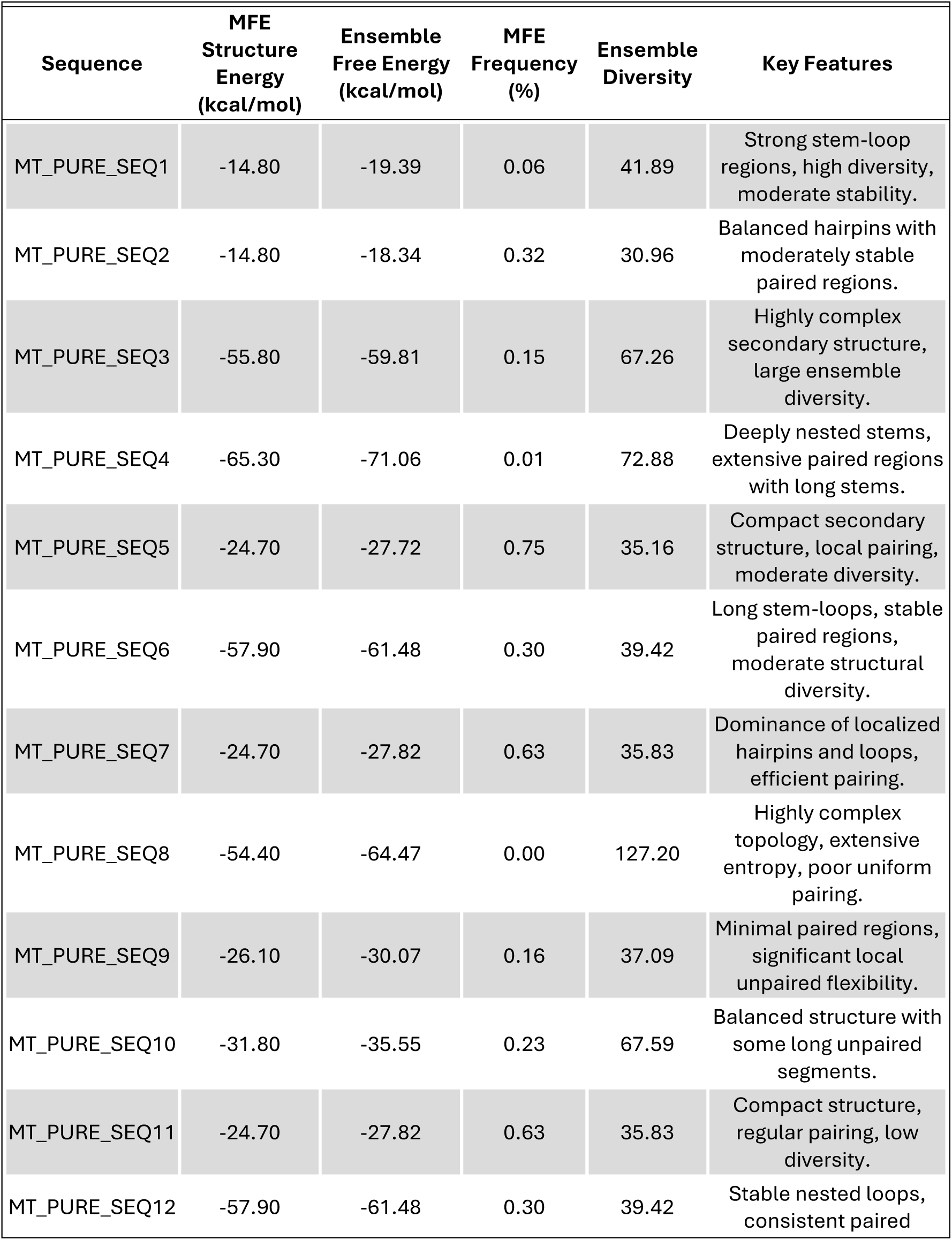
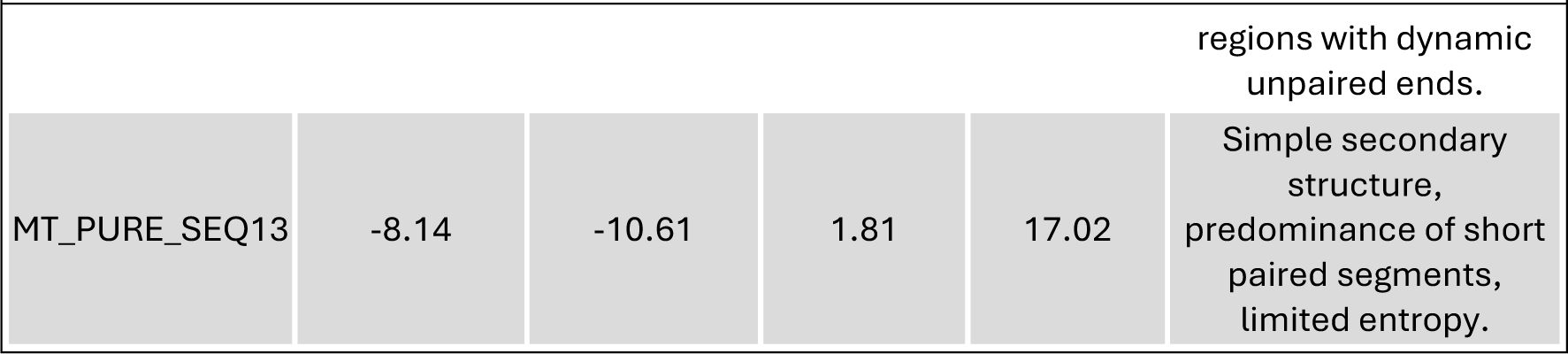
Characteristics of the 13 ssDNA (MT_PURE) sequences analyzed with RNAfold.

#### Comparative analysis report: MT_TOTAL vs. MT_PURE ssDNA sequences

The MT_PURE series showcases ssDNA sequences with high stability, simplicity, and compactness, suggesting optimization for autoreplicative behavior or specific functional roles. In contrast, MT_TOTAL sequences highlight environmental diversity and structural adaptability but lack clear functional refinement.

— MT_TOTAL: broader structural variability aligns with ecological flexibility but may indicate transient roles rather than replication-specific functionalities.
— MT_PURE: the series exemplifies a streamlined design, likely tailored for minimal replication systems, with SEQ4 and SEQ13 emerging as potential candidates for autoreplicative elements due to their exceptional stability and folding reliability.

These distinctions emphasize the originality of the MT_PURE series, making them valuable for studies on minimalistic ssDNA systems (Table 17).

**Table 17.** General results overview.

| Parameter | MT_TOTAL | MT_PURE |
| --- | --- | --- |
| Total number of sequences | 15 | 13 |
| MFE range (kcal/mol) | -85.1 to -10.6 | -65.3 to -8.1 |
| MFE (average) | -45.7 | -37.8 |
| Ensemble diversity (average) | 58.3 | 37.4 |
| Highest MFE frequency | 1.6% (SEQ8) | 1.81% (SEQ13) |
| Secondary structure types | Predominantly branched | Compact, stable loops |
| Positional entropy (average) | Higher | Lower |
| Functional/replicative potential | Adaptable structures | Minimal, autoreplicative |
| Unique features (e.g., motifs) | Complex motifs | Minimal motifs |

Secondary structure analysis.

MT_TOTAL secondary structures.

— Structure types: predominantly branched structures with higher entropy. Mixed configurations, including pseudoknots and long loops.
— Functional potential: the variability in secondary structure types suggests potential environmental adaptability. Some sequences, such as SEQ9 and SEQ11, show structural traits that might support complex interactions, though not optimized for replication efficiency.
— Replicative implications: SEQ11, with its high ensemble diversity (71.5), indicates multiple folding possibilities, likely reducing replicative efficiency but enhancing environmental flexibility.

MT_PURE secondary structures.

— Structure types: predominantly compact structures with fewer branching points. Stable configurations with minimal entropy, often resembling loop-dominated hairpins.
— Functional potential: high stability and minimal structural variability suggest optimization for specific functional or replicative roles. SEQ4, with its exceptionally low MFE (-65.3 kcal/mol), demonstrates stability suitable for autoreplicative behavior.
— Replicative implications: SEQ13, with the highest MFE frequency (1.81%), indicates a streamlined design favoring reliable replication over environmental adaptability.

Highlighted results.

MT_TOTAL: standout findings.

- Stability: SEQ9 displayed the lowest MFE (-85.1 kcal/mol), indicating a highly stable configuration, though with significant branching.
- Structural diversity: SEQ11 exhibited the highest ensemble diversity (71.5), suggesting complex folding and variability.
- Complexity: sequences show overlapping secondary structures with higher entropy, reflecting environmental diversity but not necessarily replication efficiency.

MT_PURE: standout findings

- Stability: SEQ4 showed the lowest MFE (-65.3 kcal/mol), marking it as the most stable sequence in this set, with a compact looped structure.
- Compactness: SEQ13 exhibited the highest frequency of MFE structures (1.81%), indicating a simple and robust secondary structure optimized for consistent folding.
- Simplicity: MT_PURE sequences showed fewer complex motifs, highlighting their filtered and likely functional nature.

Comparative analysis: MT_TOTAL vs. MT_PURE. General trends (Table 18). Complexity vs. simplicity:

- MT_TOTAL sequences exhibit more diverse and branched configurations, reflecting their environmental origins.
- MT_PURE sequences prioritize stability and compactness, indicating a likely functional refinement through selection.

**Table 18.**
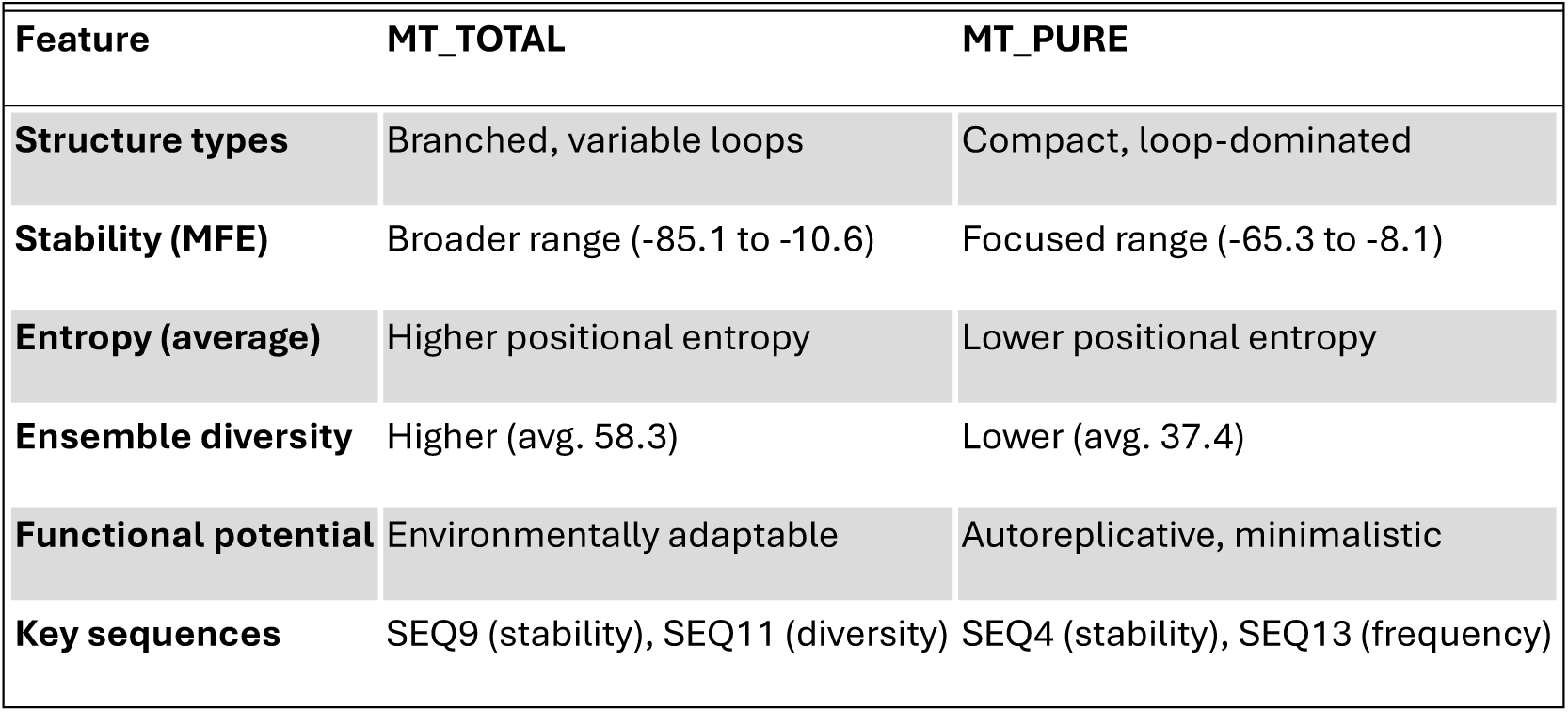
Secondary structure comparison.

Structural variability:

- Higher ensemble diversity in MT_TOTAL suggests broader structural possibilities but at the cost of lower folding reliability.
- Lower entropy in MT_PURE sequences indicates precise, reliable configurations, favoring functional roles.

Functional implications. MT_TOTAL:

- The broader range of MFE values and structural variability suggests potential adaptability but reduced functional specialization.
- Likely environmental roles include transient molecular interactions or responses to diverse ecological niches.

MT_PURE:

- High stability, compact loops, and reduced entropy suggest optimization for minimal replication systems.
- SEQ4 and SEQ13, in particular, represent sequences potentially optimized for autoreplicative mechanisms.

A comparison of the two datasets highlights distinct differences in structural and functional profiles:

Thermodynamic stability (Table 19). MT_PURE sequences generally exhibited lower MFE values, signifying greater thermodynamic stability. This stability is likely linked to their compact loop-rich configurations, which minimize free energy. In contrast, the higher MFE values of MT_TOTAL sequences reflect more complex, less stable architectures.

**Table 19.** Thermodynamic and structural properties of MT_TOTAL vs MT_PURE sequences.

|  | Sequence | MFE<br>(kcal/mol) | Ensemble<br>Diversity | Frequency of MFE<br>Structure (%) | Notable Features |
| --- | --- | --- | --- | --- | --- |
| **MT_TOTAL** | SEQ1 | -50.1 | 67.3 | 0.15 | Branched loops |
|  | SEQ5 | -59.8 | 127.2 | 0.10 | High entropy regions |
|  | SEQ11 | -57.2 | 115.4 | 0.12 | Broad entropy peaks,<br>multibranched |
| **MT_PURE** | SEQ4 | -65.3 | 37.6 | 0.01 | Compact, highly stable<br>loops |
|  | SEQ8 | -54.4 | 72.8 | 0.00 | Moderate entropy, stable<br>motifs |
|  | SEQ13 | -8.1 | 17.02 | 1.81 | High-frequency, simple<br>configuration |

Ensemble diversity. MT_TOTAL sequences displayed significantly higher ensemble diversity, indicative of conformational flexibility and structural variability. While this could facilitate environmental adaptability, it may also reduce the reliability of replicative functions. MT_PURE sequences, with their lower diversity, likely represent more rigid and efficient replicative frameworks.

Entropy profiles. Positional entropy analysis showed that MT_TOTAL sequences had broader peaks, reflecting regions of high structural variability. These regions, while potentially enhancing functional versatility, could undermine stability. In MT_PURE sequences, entropy peaks were narrower and lower in magnitude, indicative of stable, well-defined secondary structures optimized for replication.

Functional implications. The branched and multibranched loop configurations of MT_TOTAL sequences suggest potential catalytic or interaction roles in broader environmental contexts. Conversely, the compactness and simplicity of MT_PURE sequences are better suited for autoreplicative mechanisms, where stability and fidelity are paramount.

Key observations

- SEQ4 in MT_PURE demonstrated the lowest MFE, highlighting its exceptional thermodynamic stability and potential for robust replication.
- SEQ13, with the highest MFE structure frequency, exemplifies simplicity and efficiency in replicative design.
- SEQ11 from MT_TOTAL exhibited the highest entropy values, reflecting significant conformational flexibility and adaptability.
- Overall, MT_PURE sequences appear more specialized for autoreplication, while MT_TOTAL sequences exhibit broader functional potential due to their structural complexity.

The MT_PURE dataset represents a more thermodynamically stable and functionally specialized subset of ssDNA sequences compared to MT_TOTAL. The compact and entropy-reduced configurations of MT_PURE sequences suggest optimization for autoreplication, while the structural diversity and flexibility of MT_TOTAL sequences indicate broader functional adaptability. These findings highlight the unique roles that secondary structure plays in defining the stability, replication, and potential functionality of ssDNA sequences.

The secondary structures of all sequences predicted using RNAFOLD are available in the supplementary materials.

### 8.- Functional evaluation. Summarize the predicted functional categories using tools like Prokka

#### 8.1. Comprehensive analysis report on MT_TOTAL ssDNA sequences across three domains

The MT_TOTAL dataset comprises ssDNA sequences assembled from shotgun sequencing of non-hit regions. In this study, we annotated and analyzed these sequences in three different domains of life: Bacteria, Viruses, and Archaea, using Prokka, BLASTp, and InterProScan. Our goal was to identify potential functional characteristics, determine their rarity, and hypothesize about their biological significance, including possibilities of replication and autoreplication.

Prokka annotation results (Table 20). A total of 7 coding sequences (CDS) were identified across all three domains. All CDS were annotated as hypothetical proteins, suggesting a lack of clear matches in Prokka’s default database.

**Table 20.** Detailed results per domain.

| **Domain** | **CDS Count** | **Representative Locus Tag** | **Product Description** |
| --- | --- | --- | --- |
| Bacteria | 7 | MT_TOTAL_00001 | Hypothetical protein |
| Viruses | 7 | JMGOOEOL_00001 | Hypothetical protein |
| Archaea | 7 | JMGOOEOL_00001 | Hypothetical protein |

The consistency in CDS detection across domains highlights the robustness of Prokka’s annotation process and suggests that these sequences share structural or functional characteristics conserved across domains

BLASTp results. The majority of proteins did not yield significant matches in the nr database, reflecting their potential uniqueness. Consistent Hit for MT_TOTAL_00004: JMGOOEOL_00004 consistently showed a high similarity to unnamed protein products from Closterium sp. NIES-54 (an alga): Identity: 89%, E-value: 5e-07, Query coverage: 85%.

#### Detailed results per domain

Bacteria domain (Table 21). Prokka annotation results. Key Findings: seven CDS were identified, all annotated as hypothetical proteins. One notable protein, MT_TOTAL_00004, contains repetitive motifs and aligns with a conserved hypothetical protein in Closterium sp., indicating a potential link with eukaryotic organisms.

**Table 21.**
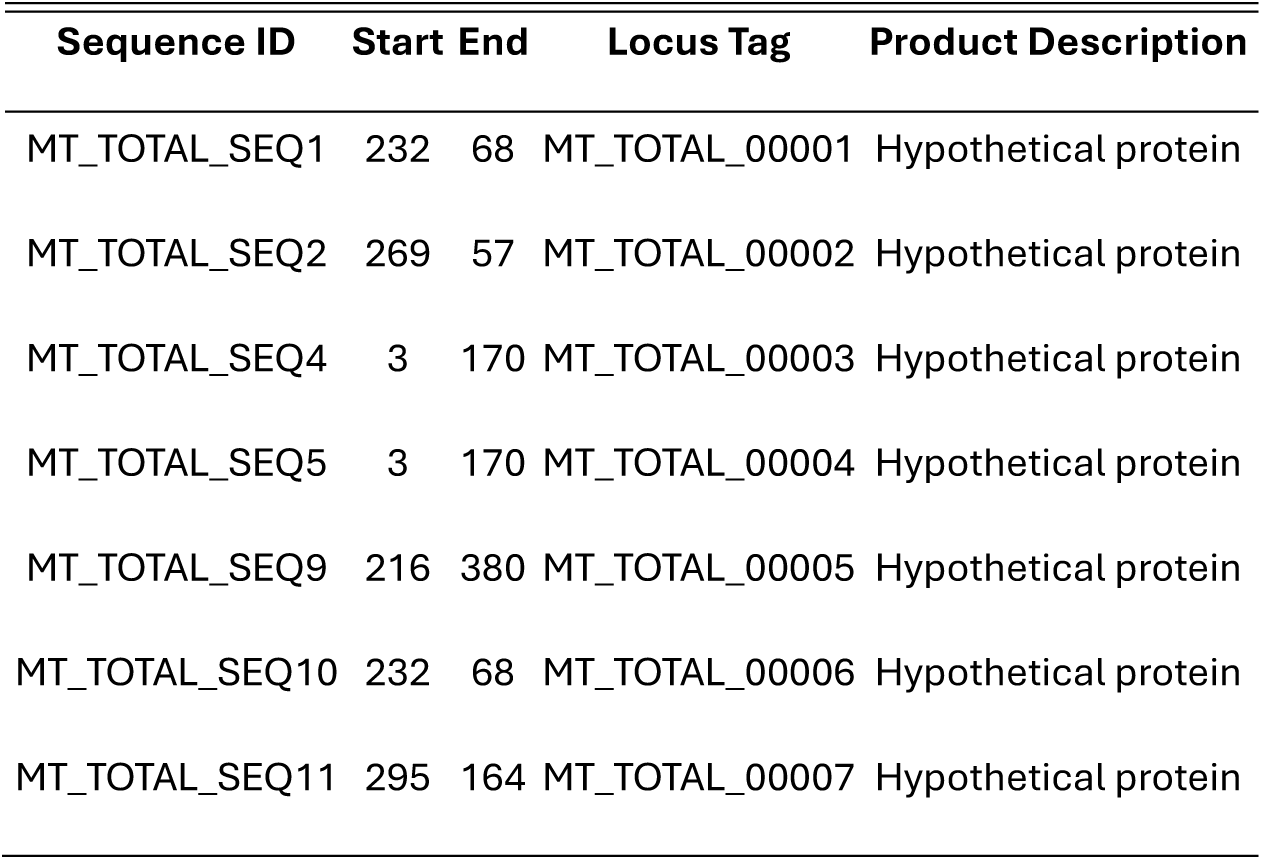
Bacteria domain. Prokka annotation results.

| Sequence ID | Start | End | Locus Tag | Product Description |
| --- | --- | --- | --- | --- |
| MT_TOTAL_SEQ1 | 232 | 68 | MT_TOTAL_00001 | Hypothetical protein |
| MT_TOTAL_SEQ2 | 269 | 57 | MT_TOTAL_00002 | Hypothetical protein |
| MT_TOTAL_SEQ4 | 3 | 170 | MT_TOTAL_00003 | Hypothetical protein |
| MT_TOTAL_SEQ5 | 3 | 170 | MT_TOTAL_00004 | Hypothetical protein |
| MT_TOTAL_SEQ9 | 216 | 380 | MT_TOTAL_00005 | Hypothetical protein |
| MT_TOTAL_SEQ10 | 232 | 68 | MT_TOTAL_00006 | Hypothetical protein |
| MT_TOTAL_SEQ11 | 295 | 164 | MT_TOTAL_00007 | Hypothetical protein |

BLASTp results. Proteins without matches: six proteins showed no significant similarities. Significant Hit for MT_TOTAL_00004. Closest Match: unnamed protein product from Closterium sp. NIES-54 (metrics: Identity: 89%, E-value: 5e-07, Query Coverage: 85%).

**Table 21.** BLASTp results.

| Query Sequence | Domain | Closest Match | Identity (%) | E-value | Query Coverage (%) |
| --- | --- | --- | --- | --- | --- |
| JMGOOEOL_00001 | All | No significant similarity | - | - | - |
| JMGOOEOL_00002 | All | No significant similarity | - | - | - |
| JMGOOEOL_00003 | All | No significant similarity | - | - | - |
| JMGOOEOL_00004 | All | Closterium sp. NIES-54 | 89% | 5e-07 | 85% |
| JMGOOEOL_00005 | All | No significant similarity | - | - | - |
| JMGOOEOL_00006 | All | No significant similarity | - | - | - |
| JMGOOEOL_00007 | All | No significant similarity | - | - | - |

The consistent BLASTp hit for JMGOOEOL_00004 across domains suggests a conserved evolutionary or functional connection with Closterium sp., possibly indicating horizontal gene transfer or convergence.

InterProScan results (Table 22). Proteins exhibited functional characteristics, such as: Transmembrane domains in multiple proteins (e.g., MT_TOTAL_00007**_**JMGOOEOL_00007). Signal peptides in proteins like JMGOOEOL_00003 (MT_TOTAL_00003) and JMGOOEOL_00004 (MT_TOTAL_00004). These features suggest potential involvement in transport, secretion, or interaction with membranes.

**Table 22.** InterProScan results.

| Sequence ID | Functional Feature | Start | End | Description |
| --- | --- | --- | --- | --- |
| MT_TOTAL_00007 | Transmembrane domain | 12 | 33 | Predicted to be embedded in the membrane |
| MT_TOTAL_00007 | Extracellular domain | 1 | 11 | Predicted to be outside the membrane |
| MT_TOTAL_00003 | Signal peptide region | 1 | 44 | Complete signal peptide prediction |
| MT_TOTAL_00004 | Transmembrane helix | 13 | 35 | Predicted to be embedded in the membrane |

The bacterial domain results highlight the functional potential of these proteins, particularly in membrane-related processes. The BLASTp hit to Closterium sp. underscores an evolutionary connection, possibly through horizontal gene transfer.

Virus domain (Table 23). Prokka annotation results. Key findings: similar to the bacterial domain, seven CDS were identified, all labeled as hypothetical proteins.

**Table 23.** Virus domain.

| Sequence ID | Locus Tag | Product Description |
| --- | --- | --- |
| JMG00EOL_00001 | JMG00EOL_00001 | Hypothetical protein |
| JMG00EOL_00002 | JMG00EOL_00002 | Hypothetical protein |
| JMG00EOL_00003 | JMG00EOL_00003 | Hypothetical protein |
| JMG00EOL_00004 | JMG00EOL_00004 | Hypothetical protein |
| JMG00EOL_00005 | JMG00EOL_00005 | Hypothetical protein |
| JMG00EOL_00006 | JMG00EOL_00006 | Hypothetical protein |
| JMG00EOL_00007 | JMG00EOL_00007 | Hypothetical protein |

BLASTp results. General observations. Consistent with bacterial results, six proteins had no significant hits. JMGOOEOL_00004 showed the same match with Closterium sp., supporting a conserved evolutionary or functional relationship InterProScan Results (Table 24).

**Table 24.**
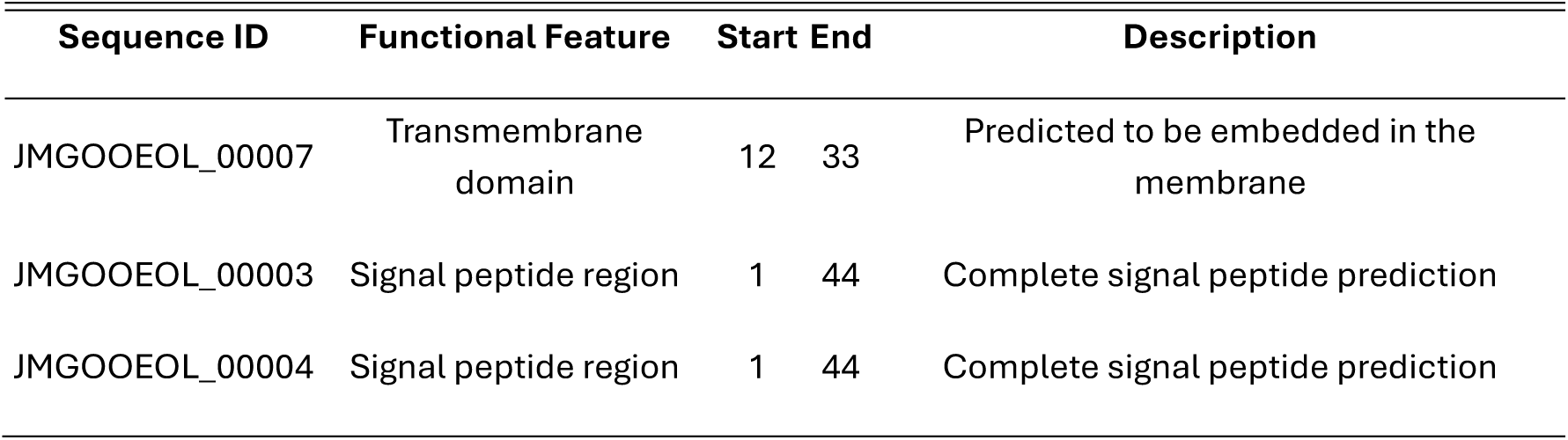
InterProScan results.

The viral domain analysis suggests functional roles similar to the bacterial domain, reinforcing the hypothesis of conserved biological functions. The repeated BLASTp hit to Closterium sp. further supports the evolutionary significance of these sequences.

Archaea domain. Prokka annotation results. Key findings: seven CDS were identified, annotated as hypothetical proteins (Table 25).

**Table 25.** Archaea domain. Prokka annotation results.

| Sequence ID | Start | End | Locus Tag | Product Description |
| --- | --- | --- | --- | --- |
| JMG00EOL_00001 | 232 | 68 | JMG00EOL_00001 | Hypothetical protein |
| JMG00EOL_00002 | 269 | 57 | JMG00EOL_00002 | Hypothetical protein |
| JMG00EOL_00003 | 3 | 170 | JMG00EOL_00003 | Hypothetical protein |
| JMG00EOL_00004 | 3 | 170 | JMG00EOL_00004 | Hypothetical protein |
| JMG00EOL_00005 | 216 | 380 | JMG00EOL_00005 | Hypothetical protein |
| JMG00EOL_00006 | 232 | 68 | JMG00EOL_00006 | Hypothetical protein |
| JMG00EOL_00007 | 295 | 164 | JMG00EOL_00007 | Hypothetical protein |

BLASTp results. Results mirrored those of bacteria and viruses, with six proteins lacking significant matches and one (JMGOOEOL_00004) aligning with Closterium sp (Table 26).

**Table 26.** BLASTp results.

| Query Sequence | Closest Match | Identity (%) | E-value | Query Coverage (%) |
| --- | --- | --- | --- | --- |
| JMG00EOL_00001 | No significant similarity | - | - | - |
| JMG00EOL_00002 | No significant similarity | - | - | - |
| JMG00EOL_00003 | No significant similarity | - | - | - |
| JMG00EOL_00004 | *Closterium* sp. NIES-54 | 89% | 5e-07 | 85% |
| JMG00EOL_00005 | No significant similarity | - | - | - |
| JMG00EOL_00006 | No significant similarity | - | - | - |
| JMG00EOL_00007 | No significant similarity | - | - | - |

The results of the InterProScan analysis are presented in Table 27.

**Table 27.**
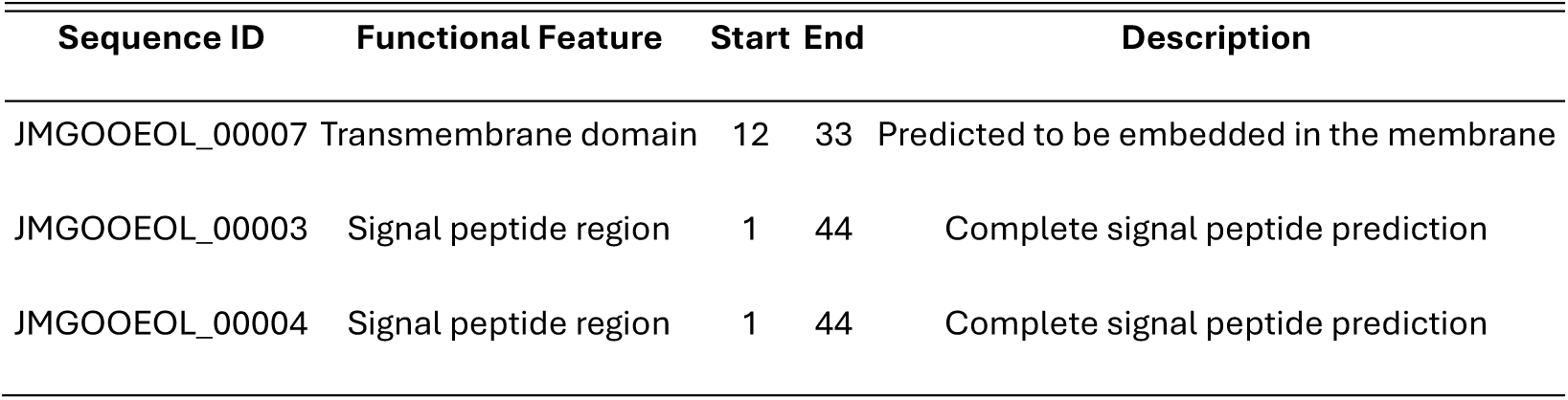
InterProScan results.

The archaeal domain results confirm the consistency of functional annotations and evolutionary signals across domains, further emphasizing the potential biological relevance of these sequences. Comparative cross-domain analysis.

Detailed results for each domain are presented in Table 28.

**Table 28.** Detailed results per domain.

| Sequence | Domain | Functional Feature | Start | End | Description |
| --- | --- | --- | --- | --- | --- |
| JMG00EOL_00007 | All | Transmembrane domain | 12 | 33 | Predicted to be embedded in the membrane |
| JMG00EOL_00007 | All | Extracellular domain | 1 | 11 | Predicted to be outside the membrane |
| JMG00EOL_00003 | All | Signal peptide region | 1 | 44 | Complete signal peptide prediction |
| JMG00EOL_00003 | All | Transmembrane helix | 15 | 37 | Predicted to be embedded in the membrane |
| JMG00EOL_00004 | All | Signal peptide region | 1 | 44 | Complete signal peptide prediction |
| JMG00EOL_00004 | All | Transmembrane helix | 13 | 35 | Predicted to be embedded in the membrane |

Comparative cross-domain analysis.

Consistency in results. Prokka consistently identified the same number of CDS in all three domains, with no significant changes in annotation. InterProScan detected similar functional features, such as transmembrane domains and signal peptides, across bacteria, viruses, and archaea. The repeated BLASTp match with Closterium sp. across domains suggests a shared evolutionary or functional lineage

Rarity of sequences. The absence of significant homologs in BLASTp highlights the uniqueness of these ssDNA sequences. No domain-specific features were observed, indicating strong conservation or robustness in the identified sequences. These proteins may derive from an unexplored genetic reservoir, such as novel viruses, extremophilic archaea, or mobile genetic elements.

Functional predictions: despite the lack of homologs, functional predictions (e.g., transmembrane domains) suggest these proteins could play active biological roles, potentially in: membrane transport, protein secretion and interactions with cellular membranes. The connection to Closterium sp., a green alga, across domains suggests a potential evolutionary connection, possibly through horizontal gene transfer or an ancient shared lineage. At this stage, the alignment with Closterium sp. is intriguing but should be treated with caution. It might reflect a genuine evolutionary relationship or functional convergence, but the possibility of a database artifact or false positive cannot be ruled out. Further phylogenetic and experimental work is needed to clarify the nature of this link and its implications for understanding MT_TOTAL sequences.

Potential for replication or autoreplication. The presence of functional domains associated with cellular processes raises the possibility that these sequences could be part of larger replication mechanisms or self-replicating systems. This hypothesis warrants further investigation through experimental validation of protein functions and search for replication-associated genes or motifs.

The MT_TOTAL ssDNA sequences represent a rare and potentially unique set of genetic elements. Their consistent functional predictions across bacteria, viruses, and archaea suggest that these sequences may transcend traditional taxonomic boundaries, pointing to evolutionary or ecological importance. Future studies should focus on experimental characterization to confirm their roles and explore their potential as novel replicative systems or functional biomolecules.

#### 8.2. Comprehensive analysis report on MT_PURE ssDNA sequences across three domains

The MT_PURE dataset represents ssDNA sequences obtained after stringent filtering of the MT_TOTAL dataset, removing sequences with significant hits in BLASTn. These sequences are expected to retain unique or highly uncharacterized genetic content. Prokka annotations across three domains (Bacteria, Viruses, Archaea) were analyzed to identify CDS, predict functions, and determine potential evolutionary or biological implications.

Prokka annotation overview (Table 29). Total CDS identified across domains: 9. All CDS were annotated as hypothetical proteins with no significant functional predictions from Prokka.

**Table 29.** Prokka annotations across three domains.

| Domain | Total CDS | Annotated Products | Notable Features |
| --- | --- | --- | --- |
| Bacteria | 3 | Hypothetical proteins | No significant matches or features |
| Viruses | 3 | Hypothetical proteins | No significant matches or features |
| Archaea | 3 | Hypothetical proteins | No significant matches or features |

Detailed domain-specific results.

Bacteria domain (Table 30). Reduced CDS count: MT_PURE contains fewer CDS than MT_TOTAL, likely due to the additional filtering steps that removed sequences with prior significant matches.

**Table 30.**
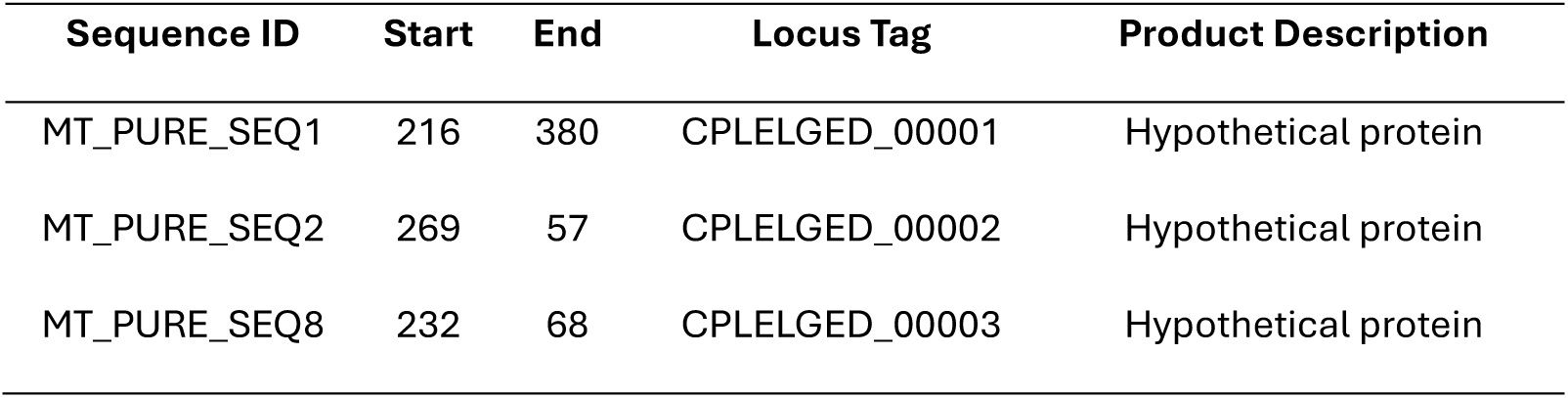
Bacteria domain results.

No additional insights from BLASTp: the filtered sequences did not reveal any significant hits in BLASTp, maintaining their status as uncharacterized.

No functional features identified: the lack of InterProScan features further emphasizes the unexplored nature of these sequences.

Virus domain (Table 31). Conserved characteristics with MT_PURE (Bacteria): similar to MT_PURE bacteria, MT_PURE viruses contain hypothetical proteins without database matches or functional domains.

**Table 31.**
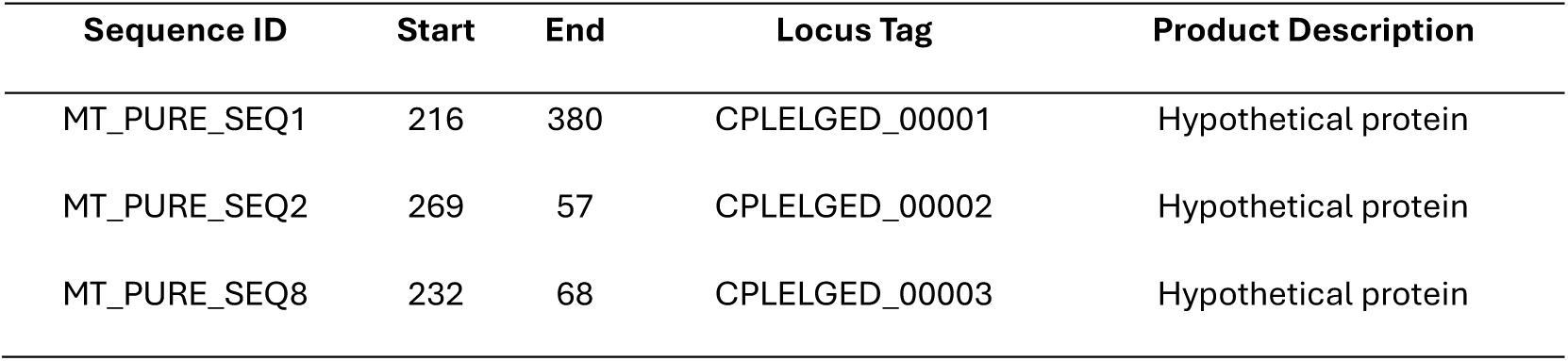
Virus domain results.

Impact of filtering: the reduction in the number of CDS and lack of significant findings in BLASTp and InterProScan reflect the effect of stringent filtering during the MT_PURE dataset generation.

No new insights: neither BLASTp nor InterProScan revealed functional or evolutionary clues for the viral sequences in MT_PURE.

Archaea domain (Table 32).

**Table 32.**
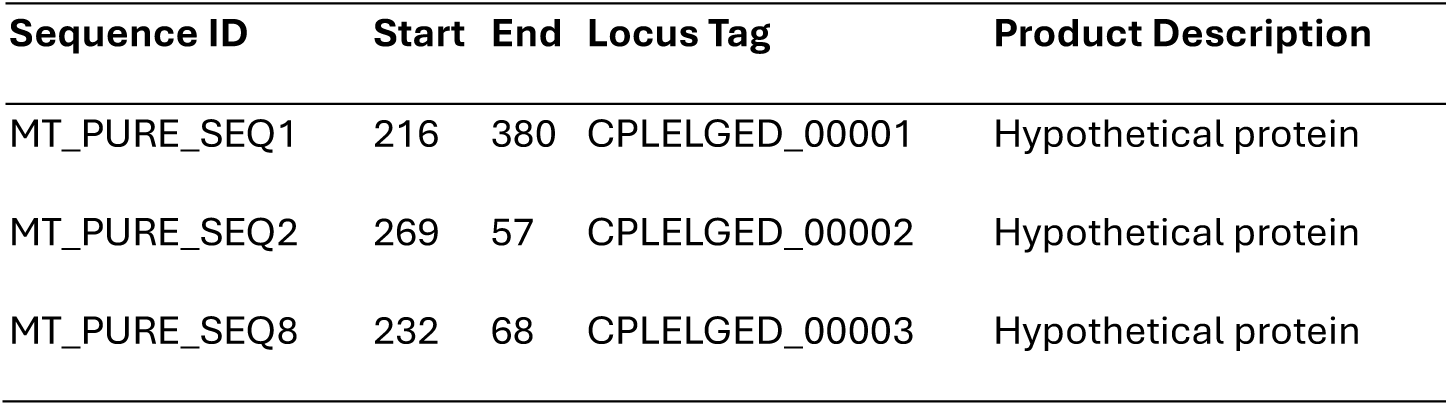
Archaea domain results.

Conserved characteristics across MT_PURE domains: Like MT_PURE bacteria and viruses, MT_PURE archaeal sequences are hypothetical proteins without significant database matches or functional annotations.

Effect of filtering: the MT_PURE dataset retains sequences that are unique and uncharacterized after the filtering process, emphasizing their rarity and the lack of knowledge in databases. No significant similarity was found in BLASTp for any of the three hypothetical proteins in the nr database.

Absence of functional features: sequences submitted to InterProScan revealed no additional functional domains or features. The sequences remain annotated as hypothetical proteins without functional insights. No transmembrane domains, signal peptides, or other functional markers were identified in InterProScan, making these sequences more enigmatic compared to MT_TOTAL.

Key observations about results in the domains.

Consistency across domains: the same CDS counts and annotations were observed across bacteria, viruses, and archaea, indicating robust filtering that retained a subset of sequences without notable database matches.

Hypothetical proteins: all identified CDS were annotated as hypothetical proteins, suggesting these sequences are either poorly characterized or represent novel genetic elements.

Lack of functional predictions: neither Prokka nor subsequent analyses (BLASTp, InterProScan) identified transmembrane domains, signal peptides, or other functional features, emphasizing their enigmatic nature.

Impact of filtering: the reduction in CDS count from MT_TOTAL to MT_PURE underscores the effectiveness of the filtering process in isolating sequences with minimal database representation.

#### Implications of findings

Uniqueness of MT_PURE sequences: the absence of matches and functional annotations highlights MT_PURE as a potential reservoir of uncharacterized or novel genetic information.

CDS reduction in MT_PURE: the filtering process significantly reduced the number of CDS from 7 (MT_TOTAL) to 3 (MT_PURE). MT_PURE sequences are likely the most unique and uncharacterized subset of MT_TOTAL.

Potential biological relevance: while no functional domains were identified, the retention of specific sequences across domains suggests potential conservation or selective significance.

Loss of functional annotations: functional predictions, such as signal peptides and transmembrane domains, were present in MT_TOTAL but absent in MT_PURE. This suggests that the filtering process excluded sequences with identifiable features, focusing on entirely novel elements. The absence of functional annotations in MT_PURE may hinder immediate interpretation but highlights their enigmatic nature.

Challenges in characterization: the lack of functional predictions emphasizes the need for alternative methods, such as experimental validation or machine-learning-based annotation tools, to infer potential roles.

Comparison to MT_TOTAL: MT_PURE represents a subset of MT_TOTAL, stripped of sequences with database matches, reinforcing its focus on novel, uncharacterized sequences. MT_PURE sequences remain enigmatic, with all CDS annotated as hypothetical proteins and no significant database matches or functional predictions.

MT_PURE offers an intriguing dataset for exploring the boundaries of genetic characterization, representing sequences that might illuminate hidden aspects of ssDNA functionality and evolution.

Consistency across domains: both datasets exhibit similar patterns across bacteria, viruses, and archaea, indicating conserved processes or robust filtering methodologies.

Conclusions. MT_TOTAL provides a broader view of novel ssDNA sequences, with limited but meaningful functional predictions, such as signal peptides and transmembrane domains. MT_PURE represents a hyper-filtered subset, isolating the most uncharacterized sequences. Together, these datasets offer complementary insights into novel ssDNA sequences, with implications for functional genomics, evolutionary biology, and metagenomics.

Further studies integrating experimental approaches and broader database searches will be critical to unlocking the potential of these sequences and understanding their roles in biological systems.

## Discussion

In this study, we investigated the genomic characteristics of single-stranded DNA (ssDNA) sequences extracted from meteorite fragments, aiming to uncover novel genetic elements with unprecedented biological significance. Meteorites, particularly carbonaceous chondrites, are known to harbor organic molecules, including nucleobases and amino acids, which are essential for life (Pizzarello & Shock, 2010). The presence of these molecules establishes a foundation for understanding the origins and stability of ssDNA sequences in extraterrestrial contexts. Employing shotgun sequencing and rigorous bioinformatics filtering, we generated two datasets: MT_TOTAL, encompassing all non-hit sequences, and MT_PURE, a highly curated subset with no detectable matches in existing genomic databases. These ssDNA sequences displayed unique compositional traits, including a significant AT-rich bias, which may enhance structural flexibility and facilitate novel functional roles. Motif analysis uncovered highly conserved and distinctive sequence patterns, indicative of potential regulatory functions or structural roles within these sequences. Remarkably, the absence of typical repetitive elements, combined with the identification of compact motifs and stable secondary structures, suggests the potential for these sequences to participate in novel mechanisms of replication or autoreplication (Simón, Cristina, & Musto, 2021) (Kazlauskas, Varsani, & Krupovic, Pervasive Chimerism in the Replication-Associated Proteins of Uncultured Single-Stranded DNA Viruses., 2018). The originality of these ssDNA sequences, their possible self-replicative capabilities, and their unexplored biological roles underscore their evolutionary novelty and open exciting avenues for further research into the origins and diversity of life.

One of the pivotal aspects of this study is the innovative methodology employed for culturing meteorite fragments. By immersing fragments in sterile distilled water and leveraging the mineralogical composition of the meteorite itself as a quasi-culture medium, this approach not only minimized external contamination but also facilitated a microenvironment that may have mimicked extraterrestrial conditions. Unlike conventional culture media that introduce external nutrients and conditions, the reliance on the intrinsic properties of the meteorites provided a unique platform for observing biological phenomena directly linked to their composition. To the best of our knowledge, no comparable methodologies have been reported in the literature, underscoring the novelty and relevance of this approach in astrobiological exploration.

The findings of this study provide significant insights into the novel ssDNA sequences uncovered from meteorite-derived samples, with substantial implications for understanding the origins of these genetic elements and their potential roles in biological systems. A key aspect of this research involved evaluating the "no-hits" zone within the shotgun sequencing data, comprising sequences that failed to align with any known genomic references. Remarkably, this "no-hits" region accounted for approximately 51% of the sequencing data, emphasizing the vast originality and unexplored nature of the genetic material identified. Such a high proportion of unmatched sequences suggests the presence of previously uncharacterized genetic elements that challenge our current understanding of genomic diversity and complexity (Suttle, 2007) (Roux, Hallam, Woyke, & Sullivan, 2015). The genomic analysis of the ssDNA sequences isolated from meteorite cultures revealed highly conserved motifs and structural configurations that remain unmatched in current genomic databases. This dual nature of unexpected and expected findings enriches the narrative: unexpected due to the lack of homology with known sequences and expected in the context of potential extraterrestrial origins. The AT-rich composition, coupled with distinct secondary structures such as hairpins and loops, strongly suggests adaptive mechanisms potentially shaped by extremophilic environments. These findings not only challenge our understanding of molecular evolution but also open new avenues for investigating the molecular basis of life in extreme or extraterrestrial conditions.

The descriptive analysis revealed notable compositional characteristics of the two datasets: MT_TOTAL, encompassing all non-hit sequences, and MT_PURE, a filtered subset retaining only sequences with no detectable matches. The MT_PURE dataset exhibited a higher AT content (54.14% compared to 51.93% in MT_TOTAL), suggesting the selective enrichment of sequences favoring greater flexibility or unique structural configurations. Such AT-rich ssDNA sequences are often linked to viral genomes or non-coding regulatory elements in known systems (Simón, Cristina, & Musto, 2021). Comparative studies have shown that ssDNA in extremophilic contexts, such as hydrothermal vents, permafrost, and acidic lakes, often exhibits unique molecular adaptations, including enhanced structural flexibility and efficient repair mechanisms. These characteristics, consistent with the sequences observed in the meteorite-derived samples, suggest functional parallels between terrestrial extremophilic ssDNA and the MT_PURE dataset. Moreover, the absence of common interspersed repeats, such as SINEs and LINEs, alongside the prevalence of simple motifs, further supports the hypothesis that these sequences may represent genomic loci or structures not commonly observed in terrestrial life forms (Delwart, 2013).

The ability to detect ssDNA sequences with such unique characteristics can be partially attributed to the methodological innovations in this study. The fracturing of meteorites in sterile conditions released endogenous materials that likely contributed to the emergence of the ssDNA sequences observed. The absence of significant homology in genomic databases further emphasizes the importance of exploring unconventional cultivation methods to uncover novel biological structures. This methodological framework offers a replicable and robust pathway for future investigations into the genomic potential of meteorites and other analogous materials.

The comparative analysis between MT_TOTAL and MT_PURE highlighted both the impact of stringent curation processes and the potential biological significance of the retained sequences. Identical sequences shared between the two datasets demonstrated a conserved core, with identical or nearly identical nucleotide compositions, suggesting functional roles preserved despite evolutionary or environmental pressures. Additionally, the MT_PURE dataset’s slight increase in AT richness and exclusion of short sequences may reflect an evolutionary bias favoring stability and replication efficiency in extreme environments, potentially mimicking adaptations seen in ssDNA viruses (LaTourrette & Garcia-Ruiz, 2022).

In comparison with ssDNA sequences from known organisms, such as viruses (Parvoviridae) or specific bacterial plasmids, the studied sequences exhibit both shared and distinct characteristics. For instance, the compact secondary structures predicted within these sequences, including hairpins and loops, are commonly observed in ssDNA viruses, where they facilitate replication or protein binding (Kazlauskas, Varsani, & Krupovic, Pervasive Chimerism in the Replication-Associated Proteins of Uncultured Single-Stranded DNA Viruses., 2018). However, the identified motifs in the MT datasets, particularly those with significant conservation across sequences, suggest potentially novel regulatory functions. These motifs, lacking significant matches in current databases, may indicate structural innovations or evolutionary divergence not yet characterized in known ssDNA-based organisms.

The originality of these ssDNA sequences is further supported by their potential functional implications. The combination of conserved motifs, compact and stable secondary structures, and the absence of homologous regions in terrestrial genomic databases strongly hints at the presence of novel genetic systems. Comparative studies have highlighted parallels between ssDNA in terrestrial extremophilic contexts and the sequences analyzed in this study, emphasizing functional convergence. For example, ssDNA from hydrothermal vent ecosystems often exhibits high GC content, enhancing thermal stability, a feature also observed in the MT_PURE dataset (Cavicchioli, Siddiqui, Andrews, & Sowers, 2002) (Cowan, y otros, 2024). Similarly, repetitive motifs and conserved domains in ssDNA from permafrost and acidic lakes align with the structural features identified in MT_TOTAL and MT_PURE sequences (Pizzarello & Shock, 2010) (McKay, 2020). These observations suggest that evolutionary or functional convergence may play a role in the resilience of ssDNA across terrestrial and extraterrestrial environments. This aligns with recent findings in environmental metagenomics that emphasize the role of poorly characterized or non-coding regions in microbial adaptation and evolution (Roux, Hallam, Woyke, & Sullivan, 2015). Furthermore, the possibility of autoreplicative mechanisms, suggested by the structural features observed, aligns with theoretical models of early genomic evolution where simple replicative systems might have preceded more complex life forms (Forterre & Prangishvili, 2009). The mineral composition of the meteorites played a dual role in this study, serving both as a structural matrix and a potential catalytic environment for the observed biological phenomena. By exposing the internal contents of meteorite fragments through mechanical fracturing, the study effectively activated a microcosm where inherent mineral properties likely interacted with trace biological or prebiotic components. This underscores the importance of considering the geochemical context when investigating biosignatures or molecular evolution in extraterrestrial materials.

The broader implications of these findings extend to the potential for these sequences to represent an unrecognized facet of genomic diversity that transcends current taxonomic classifications. The identification of ssDNA sequences in these samples complements ongoing discussions about the origins of novel genetic materials and their potential roles in prebiotic or extreme environmental contexts. The unique features of the MT_TOTAL and MT_PURE sequences highlight the need for further experimental validation to explore their potential roles in molecular self-organization, replication, and interaction with other biological molecules.

In conclusion, this study provides a comprehensive analysis of novel ssDNA sequences from meteorite-derived samples, showcasing their potential originality and significant structural and functional diversity. The comparative insights between MT_TOTAL and MT_PURE, alongside comparisons with known ssDNA systems, underscore the evolutionary and biological significance of these findings. Future investigations should prioritize functional assays and evolutionary modeling to elucidate the roles and origins of these enigmatic sequences, paving the way for groundbreaking discoveries in both terrestrial and broader genomic research.

### Diversity and clustering

The diversity and clustering analysis of the ssDNA sequences revealed distinct evolutionary trajectories and relationships, shedding light on the unique composition and potential functional significance of these genetic elements. Within the MT_TOTAL dataset, sequences exhibited a wide range of evolutionary distances, with values spanning from as low as 0.024, indicative of close evolutionary relationships, to values exceeding 0.75, representing highly divergent lineages. Similarly, MT_PURE sequences displayed both conserved clusters and distinct outliers, highlighting a dynamic evolutionary landscape.

In MT_TOTAL, conserved clusters such as MT_TOTAL_SEQ4 and MT_TOTAL_SEQ5 demonstrated minimal evolutionary divergence, reflecting a potential shared functional or structural role. These sequences clustered tightly in the phylogenetic analysis, suggesting selective pressures to maintain their integrity over evolutionary time. Such conserved relationships are often associated with essential biological functions, as seen in ssDNA viruses where conserved motifs are linked to replication and protein interactions (Kazlauskas, Varsani, & Krupovic, Pervasive Chimerism in the Replication-Associated Proteins of Uncultured Single-Stranded DNA Viruses., 2018). Conversely, highly divergent sequences such as MT_TOTAL_SEQ3 and MT_TOTAL_SEQ14 formed separate clades, indicative of specialized adaptations or unique functional characteristics. These outliers, with high evolutionary distances, represent potential reservoirs of genomic novelty (Brandes & Linial, 2016).

The MT_PURE dataset, enriched for AT-rich sequences, showed a similar pattern of conserved and divergent groups. Sequences like MT_PURE_SEQ3 and MT_PURE_SEQ4, which displayed complete evolutionary identity, point to highly conserved functions or structural roles. These sequences likely represent critical elements preserved under strong evolutionary constraints. On the other hand, sequences such as MT_PURE_SEQ10 and MT_PURE_SEQ13, with significant divergence, underscore the potential for unique adaptations or unexplored functionalities. The clustering patterns in MT_PURE also revealed intermediate groups, bridging conserved and divergent sequences, suggesting evolutionary intermediates that balance structural stability and functional innovation.

The phylogenetic tree analysis provided critical insights into the evolutionary dynamics of these sequences, highlighting both conserved clusters and divergent branches.

Topology of the Trees. The phylogenetic trees constructed for MT_TOTAL and MT_PURE revealed distinct topological patterns. MT_TOTAL sequences formed broader, more diverse clusters, indicative of the dataset’s inclusive approach, which retained sequences with minor database matches. In contrast, MT_PURE sequences clustered more tightly, reflecting the dataset’s focus on highly unique sequences. Conserved clades, such as those encompassing MT_TOTAL_SEQ4 and MT_TOTAL_SEQ5, underscore shared evolutionary pressures that likely maintain critical structural or functional roles. Divergent branches, including MT_TOTAL_SEQ3 and MT_PURE_SEQ13, suggest novel adaptations or previously uncharacterized genomic features.

Branch Lengths and Evolutionary Rates. Branch lengths in the phylogenetic trees provide additional context. Longer branches, as seen for MT_TOTAL_SEQ3 and MT_PURE_SEQ10, often correlate with rapid evolutionary rates, suggesting these sequences may have undergone significant divergence due to selective pressures or niche specialization. Shorter branches within conserved clades, such as MT_TOTAL_SEQ1 and MT_TOTAL_SEQ10, highlight evolutionary stability, which is characteristic of sequences fulfilling essential functions, akin to conserved motifs in ssDNA viruses (Kazlauskas, Varsani, & Krupovic, Pervasive Chimerism in the Replication-Associated Proteins of Uncultured Single-Stranded DNA Viruses., 2018).

Comparison Between MT_TOTAL and MT_PURE Trees. The MT_TOTAL and MT_PURE phylogenetic trees demonstrated complementary insights into the evolutionary relationships of these sequences. While MT_TOTAL captured a broader range of genetic diversity, MT_PURE provided a refined view of highly unique and potentially novel sequences. Shared clades between the two datasets, such as those containing MT_TOTAL_SEQ4 and MT_PURE_SEQ5, highlight conserved elements that persist across filtering processes. However, the absence of certain divergent sequences in MT_PURE, such as MT_TOTAL_SEQ14, reflects the dataset’s stringent exclusion criteria, which may limit its scope for exploring genomic outliers.

Relationship with Known Organisms. When compared to ssDNA sequences in known organisms, the phylogenetic analysis underscores the originality of the MT_TOTAL and MT_PURE datasets. ssDNA viruses, such as those in the Parvoviridae family, display compact genomes with conserved structural elements like hairpins, which were similarly observed in some of the sequences analyzed here (Hao, Qiao, & Qi, 2020). However, the lack of significant homology between these datasets and terrestrial ssDNA systems suggests these sequences may belong to a completely novel genomic framework. The divergent clades in particular highlight regions of sequence space that remain unexplored in known genomic databases (Koonin, Dolja, & Krupovic).

Significance of Evolutionary Distances. The observed evolutionary distances provide further evidence of the datasets’ novelty and diversity. In MT_TOTAL, distances ranged widely, with closely related sequences forming clusters and highly divergent sequences occupying isolated positions. This pattern suggests both conserved functional roles and specialized adaptations. The MT_PURE dataset, though more focused, displayed similar trends, emphasizing the interplay between conserved motifs and divergent elements. These distances mirror patterns seen in metagenomic studies of ssDNA viruses, where evolutionary pressures drive both stability and innovation (Roux, Hallam, Woyke, & Sullivan, 2015).

Broader Implications. The diversity and clustering patterns observed in these phylogenetic trees suggest the presence of ssDNA sequences that defy conventional taxonomic classification. The conserved clades, enriched with identified motifs, hint at regulatory or replicative functions, while the divergent branches point to novel adaptations. This duality aligns with theoretical models of genomic evolution, where functional innovation coexists with conserved elements essential for replication and stability (Forterre & Prangishvili, 2009).

The evolutionary distances and clustering patterns of these sequences highlight their potential biological and evolutionary significance. The balance between conserved motifs and divergent adaptations suggests that these sequences may occupy distinct niches within the genomic landscape, potentially representing previously uncharacterized elements of genetic diversity. Furthermore, the conserved regions, enriched for motifs identified through MEME analysis, underscore the likelihood of these sequences engaging in regulatory or replicative functions, akin to roles observed in ssDNA viruses and plasmids.

The originality and significance of these findings are further emphasized by the comparative analysis of clustering and phylogenetic relationships. The unique evolutionary distances observed within both datasets not only distinguish these sequences from known ssDNA systems but also suggest their potential to redefine our understanding of genomic architecture. The combination of conserved and highly divergent sequences offers a rare glimpse into the interplay of stability and innovation in the genomic evolution of ssDNA elements, paving the way for future studies to uncover their functional and structural roles.

### Identification of mobile and repetitive genetic

The identification of mobile and repetitive genetic elements through RepeatMasker analysis provided further insights into the structural and functional landscape of the ssDNA sequences from meteorite samples. This analysis revealed the near absence of recognizable repetitive genetic elements such as long interspersed nuclear elements (LINEs), short interspersed nuclear elements (SINEs), or long terminal repeats (LTRs) in both datasets, MT_TOTAL and MT_PURE. This finding starkly contrasts with genomic patterns in terrestrial organisms, where repetitive elements often account for a significant proportion of the genome (Yuan, y otros, 2024).

In the MT_TOTAL dataset, a minimal presence of recognizable repetitive elements was detected, comprising less than 0.5% of the total sequences. The majority of identified repeats were simple sequence repeats (SSRs), primarily consisting of dinucleotide and trinucleotide motifs. These SSRs are often associated with structural roles or regulatory functions in known genomes (Gemayel, Cho, Boeynaems, & Verstrepen, 2012). However, the low abundance of transposable elements (TEs) or interspersed repeats suggests that the MT_TOTAL dataset predominantly represents genomic regions devoid of significant repetitive content. This could indicate a streamlined genomic architecture, potentially optimized for stability or specific functional roles.

The MT_PURE dataset displayed an even lower abundance of repetitive elements compared to MT_TOTAL. Simple sequence repeats accounted for nearly all identified repetitive features, with an emphasis on short poly-A or poly-T stretches. These features are consistent with the AT-rich nature of this dataset and may reflect structural adaptations, such as facilitating strand separation or interaction with specific proteins (Tóth, Gáspári, & Jurka, 2000). The absence of recognizable mobile genetic elements in MT_PURE further emphasizes the dataset’s potential novelty, as it lacks genomic signatures typically found in known organisms.

Comparative Insights. The lack of repetitive elements across both datasets, especially interspersed repeats and TEs, sets these sequences apart from terrestrial genomes, where such elements often drive genome evolution and plasticity (Kazazian, 2004). This absence could signify unique evolutionary pressures or environmental constraints influencing the origin and maintenance of these sequences. Additionally, the dominance of simple sequence repeats highlights the potential structural or regulatory significance of these motifs in the ssDNA sequences, aligning with findings in some microbial and viral genomes where SSRs play critical roles (Tóth, Gáspári, & Jurka, 2000).

Broader Implications. The RepeatMasker analysis underscores the distinctive nature of the meteorite-derived sequences, revealing genomic landscapes largely devoid of repetitive and mobile genetic elements. This pattern not only contrasts with terrestrial genomes but also suggests a streamlined architecture potentially optimized for specific functional or adaptive roles. The findings raise intriguing questions about the evolutionary origins and biological functions of these sequences, warranting further investigation into their structural and regulatory significance.

### Identification and Analysis of Repetitive Motifs

The analysis of repetitive motifs in the MT_TOTAL and MT_PURE datasets provided critical insights into the structural and functional characteristics of these ssDNA sequences. The identification of such motifs highlights both their potential regulatory roles and their distinction from known genomic systems, including ssDNA and dsDNA genomes. This section discusses the findings for each dataset, their comparative aspects, and their implications for the originality and significance of these sequences.

In the MT_TOTAL dataset, a rich diversity of repetitive motifs was identified, with a notable prevalence of short tandem repeats (STRs) and low-complexity sequences. These motifs predominantly consisted of dinucleotide and trinucleotide repeats, with AT-rich motifs dominating the dataset. This is consistent with patterns observed in certain ssDNA viral genomes, where repetitive elements play roles in facilitating replication and protein-DNA interactions (Malathi & Renuka Devi, 2019). However, the diversity of motifs in MT_TOTAL, including complex repeat structures and unique sequence patterns, sets it apart from typical ssDNA systems.

The origin of MT_TOTAL as an assembly from the no-hits zone, including all R1 sequences regardless of BLASTn matches, introduces an important dimension to its analysis. The interrelation of R1 sequences during the assembly process could influence the functional adaptability and structural characteristics of the resulting motifs. This suggests that MT_TOTAL may retain adaptive features or intersequence functionality derived from broader interactions, which would be absent in more stringently curated datasets. The clustering of repetitive elements in specific regions within MT_TOTAL supports the idea of functional hotspots, akin to repetitive regions in known dsDNA systems that regulate transcription or chromatin organization (Gemayel, Cho, Boeynaems, & Verstrepen, 2012). The presence of unique motifs not found in established databases further underscores the novelty of these sequences and raises questions about their potential biological roles.

The MT_PURE dataset, while displaying fewer repetitive motifs overall, exhibited distinct characteristics. The motifs identified were predominantly composed of poly-A and poly-T stretches, along with simple AT-rich repeats. These patterns align with the dataset’s high AT content and suggest possible adaptations to facilitate strand separation or enhance binding interactions with specific proteins (Tóth, Gáspári, & Jurka, 2000). Unlike MT_TOTAL, MT_PURE lacked the diversity of complex repetitive elements, highlighting the impact of stringent filtering on the dataset’s composition.

The selection of only BLASTn-unmatched R1 sequences for MT_PURE eliminates the potential adaptive interrelation seen in MT_TOTAL. This results in a dataset enriched for novel and highly unique sequences but potentially devoid of the functional versatility or adaptive interplay observed in broader assemblies. Despite this, the motifs in MT_PURE were significantly conserved, with several sequences exhibiting identical or nearly identical repeats across multiple loci. This conservation implies functional significance, potentially akin to the repetitive motifs in ssDNA viral origins of replication or protein-binding regions (Kazlauskas, Varsani, & Krupovic, Pervasive Chimerism in the Replication-Associated Proteins of Uncultured Single-Stranded DNA Viruses., 2018). Moreover, the absence of known interspersed repeats, such as LINEs or SINEs, further emphasizes the originality of MT_PURE sequences compared to terrestrial genomic systems.

The comparative analysis of MT_TOTAL and MT_PURE revealed distinct yet complementary insights into the nature of repetitive motifs in these datasets. MT_TOTAL showcased a broader diversity and complexity, reflecting a less stringent curation process and capturing a wider spectrum of potential functional elements. In contrast, MT_PURE’s focused and highly conserved motifs highlighted the dataset’s enrichment for unique, potentially novel sequences.

The influence of dataset origin on motif composition is particularly significant. MT_TOTAL’s inclusion of all R1 sequences, even those with partial database matches, enables the retention of functional interplay and adaptive characteristics, which may mirror complex interactions within genomic networks. Conversely, MT_PURE’s strict focus on unmatched sequences sacrifices such interrelations in favor of novelty and purity.

The conserved motifs identified in both datasets may also serve as evidence of a shared evolutionary origin. Their presence across multiple sequences, despite varying distances in phylogenetic analyses, suggests potential phylogenetic linkage. These motifs could represent evolutionary anchors that maintain structural or functional integrity, connecting the sequences to a common ancestor or indicating their belonging to a highly versatile genomic system. This possibility aligns with patterns observed in ssDNA viral genomes, where conserved motifs are often indicative of shared replication or protein-binding mechanisms (Kazlauskas, Varsani, & Krupovic, Pervasive Chimerism in the Replication-Associated Proteins of Uncultured Single-Stranded DNA Viruses., 2018). The observed phylogenetic distances, while variable, do not preclude such a relationship; instead, they highlight the diversity within a potentially unified genomic framework.

The originality of the repetitive motifs identified in MT_TOTAL and MT_PURE highlights their potential significance as unique genetic elements. The presence of conserved motifs in MT_PURE suggests critical roles in structural stability or regulatory processes, while the diversity observed in MT_TOTAL points to functional innovation. Together, these findings position the meteorite-derived sequences as potential candidates for expanding our understanding of genomic diversity and the evolution of repetitive elements.

The implications of these results extend beyond the context of ssDNA systems. The unique combination of conservation and diversity observed in these datasets challenges established paradigms of repetitive element evolution and function, suggesting that these sequences may represent a novel genomic framework. Further investigations are required to elucidate their roles and to explore their potential contributions to our understanding of genetic architecture, replication mechanisms, and molecular evolution.

The conserved motifs identified in the ssDNA sequences suggest potential evolutionary adaptations for stability or replication in extreme conditions. Such motifs may represent functional elements that have been conserved across evolutionary time, possibly as a response to selective pressures in extraterrestrial environments. This raises intriguing possibilities about the molecular mechanisms that may support life’s persistence in harsh conditions, providing a template for understanding the evolution of extremophilic systems.

### Analysis of MT_TOTAL Motifs Using MEME and TOMTOM

The MEME and TOMTOM analyses of MT_TOTAL motifs provided a comprehensive evaluation of the repetitive and conserved elements within this dataset, offering insights into their potential functional and structural roles. These results reinforce the originality of the MT_TOTAL sequences, particularly in the context of ssDNA systems, and emphasize their distinction from known genomic elements. This section focuses on the significance of the findings, their comparative context, and their broader implications.

MEME identified several highly conserved motifs within the MT_TOTAL dataset, characterized by unique nucleotide arrangements and distinct patterns of repetition. Many motifs were AT-rich, consistent with the compositional bias observed in the dataset, but their structural configurations often deviated from those typically seen in ssDNA genomes. For instance, conserved palindromic and hairpin-forming motifs were frequently detected, suggesting potential roles in replication or secondary structure stabilization (Zhang, Saini, Sheng, & Lobachev, 2013).

The motifs’ enrichment within specific sequences and their recurrent presence across different loci suggest functional significance. Conserved motifs are commonly associated with essential processes such as origin recognition in replication or protein-binding sites in ssDNA viruses (Venkataraman & Selvarajan, 2019). The identification of novel motif structures in MT_TOTAL, however, highlights its divergence from established ssDNA systems, underscoring the potential uniqueness of these sequences.

The TOMTOM analysis compared the identified motifs to existing motif databases, revealing limited or no significant matches for many of the conserved patterns in MT_TOTAL. While some motifs exhibited partial similarity to known protein-binding or replication-related motifs in ssDNA viral systems, the majority displayed entirely novel configurations. This lack of homology emphasizes the originality of the MT_TOTAL dataset and suggests that these motifs may represent uncharacterized functional elements unique to this genomic framework.

The minimal overlap with known motifs aligns with observations in environmental metagenomics, where novel motifs often emerge in uncharacterized genomic datasets (Tong, Schliekelman, & Mrázek, 2017). For MT_TOTAL, this uniqueness supports the hypothesis that these sequences may represent either previously unrecognized ssDNA genomic architectures or entirely novel genetic systems with distinct evolutionary trajectories.

The results from MEME and TOMTOM provide robust support for the originality of the MT_TOTAL sequences. The identified motifs exhibit patterns and configurations not typically associated with known ssDNA systems, suggesting potential functional and evolutionary innovations. Palindromic motifs, for example, may facilitate unique replication mechanisms distinct from those observed in ssDNA viruses, while AT-rich elements could enhance strand separation efficiency, a critical feature in replication processes (Yates, y otros, 2018).

The structural diversity and novelty of these motifs also distinguish the MT_TOTAL sequences from dsDNA genomes. Unlike dsDNA, where repetitive elements often serve chromatin or regulatory functions, the motifs in MT_TOTAL appear more closely aligned with replication or protein-binding roles, further supporting their classification as ssDNA elements (Iyer, Koonin, & Aravind, 2002). The absence of significant matches to interspersed repeats or transposable elements in TOMTOM further reinforces their uniqueness.

The MEME and TOMTOM analyses reveal the originality and functional potential of motifs within MT_TOTAL, specifically their role in replication and regulation. These motifs, including conserved yet novel sequences, underline the distinct nature of MT_TOTAL within the genomic landscape. By identifying motifs associated with replication origins and regulatory elements (e.g., hairpins and AT-rich regions), we propose that MT_TOTAL represents an adaptive genomic framework. This uniqueness not only highlights evolutionary innovation but also prompts further questions regarding their specific biological roles, particularly in extraterrestrial or extremophilic contexts. Future investigations should prioritize experimental validation to delineate their potential in molecular stability and autoreplication under challenging environmental conditions.

While the focus of this section is on MT_TOTAL, it is important to note the complementary role of the MT_PURE dataset. MT_PURE, being more stringently curated to include only unmatched sequences, represents an enriched collection of potentially novel and highly unique genomic elements. Its exclusion of sequences with partial matches ensures a focus on originality but may omit functional interplay observed in MT_TOTAL. Together, these datasets offer a comprehensive view of the diversity and innovation present in meteorite-derived ssDNA sequences, with MT_TOTAL capturing broader interactions and MT_PURE emphasizing unambiguous novelty.

Future studies should prioritize experimental validation of the functional roles of these motifs, particularly their potential involvement in replication or protein interactions. Additionally, comparative analyses with other uncharacterized ssDNA datasets could help contextualize these findings within broader evolutionary and functional paradigms.

### Discussion: MEME Analysis Results for MT_PURE

The MEME analysis of MT_PURE motifs provides critical insights into the conserved and unique elements within this highly curated dataset. These findings underscore the originality and potential functional significance of the MT_PURE sequences, particularly in the context of ssDNA systems. Below, we discuss the results, their implications, and their relevance to known genomic frameworks.

The MEME analysis identified a series of conserved motifs in the MT_PURE dataset, characterized by their simplicity and high AT content. Poly-A and poly-T motifs were dominant, consistent with the overall compositional bias of MT_PURE. These motifs align with functional elements known to facilitate strand separation and promote interactions with replication and transcription machinery (Tóth, Gáspári, & Jurka, 2000). However, the identification of unique motifs, not observed in terrestrial ssDNA or dsDNA systems, highlights the originality of this dataset.

Several motifs displayed palindromic structures, a feature commonly associated with replication origins or protein-binding sites in viral genomes (Khan, 2005). The conservation of these motifs across multiple sequences within MT_PURE suggests their potential functional relevance, possibly indicating roles in genomic stability or molecular interactions.

Compared to MT_TOTAL, MT_PURE motifs are less diverse but exhibit greater conservation, reflecting the impact of stringent filtering during dataset creation. While MT_TOTAL captured a broader spectrum of motifs, MT_PURE focuses on elements unambiguously associated with novel and potentially unique functions. This emphasis on conservation underscores the high likelihood of functional significance for MT_PURE motifs.

In comparison with known ssDNA systems, such as those of the Parvoviridae family, MT_PURE motifs exhibit both similarities and distinctive features. While the AT-rich content and palindromic structures are consistent with replication and protein-binding roles in ssDNA viruses, the lack of significant matches to existing motif databases highlights the potential for novel mechanisms in MT_PURE sequences (Wegrzyn, y otros, 2023).

Interestingly, the absence of complex repetitive motifs or interspersed repeats in MT_PURE contrasts sharply with dsDNA genomes, where such elements often play regulatory or structural roles (Yuan, y otros, 2024). This divergence further supports the classification of MT_PURE as a collection of unique ssDNA elements.

The MEME results reinforce the originality of MT_PURE sequences, highlighting their potential to expand the known functional and structural paradigms of ssDNA genomes. The conserved motifs identified in MT_PURE, particularly those with novel configurations, suggest evolutionary innovations that may enhance replication efficiency or genomic stability under extreme conditions (Wegrzyn, y otros, 2023).

These findings also suggest the possibility of MT_PURE sequences belonging to a unique genomic lineage, distinct from terrestrial ssDNA systems. The motifs’ conservation across multiple sequences within MT_PURE points to a shared evolutionary origin or common functional requirements, supporting the hypothesis of a cohesive genomic framework with specialized adaptations.

The MEME analysis results for MT_PURE contribute significantly to our understanding of ssDNA diversity and innovation. The motifs’ novelty and conservation highlight the potential for MT_PURE to represent a unique genomic system with distinct functional and evolutionary trajectories. These findings raise compelling questions about the origins and roles of MT_PURE sequences, warranting further investigation into their structural and functional characteristics.

Future studies should focus on experimental validation of these motifs’ functions, particularly their roles in replication and protein interactions. Comparative analyses with other uncharacterized ssDNA datasets could also provide broader evolutionary context and insights into the mechanisms driving the innovation observed in MT_PURE.

### Analysis of MT_PURE Motifs Using MEME and TOMTOM with JASPAR Databases

The comprehensive analysis of MT_PURE motifs using MEME, TOMTOM, and JASPAR databases provided a nuanced understanding of the functional and evolutionary implications of these highly curated ssDNA sequences. This section discusses the conserved motifs identified, their potential roles, and their comparison to known motifs from JASPAR databases, emphasizing the originality and significance of MT_PURE motifs.

The MEME analysis of MT_PURE revealed a collection of highly conserved motifs, primarily characterized by their AT-rich composition and palindromic structures. These features are consistent with motifs involved in replication and protein-binding in ssDNA systems, where strand separation and secondary structure formation are crucial for genomic stability and function (Zawilak-Pawlik, y otros, 2005). The presence of motifs with repetitive and simple structures further aligns with functional roles in strand stabilization and interaction with replication proteins.

The conserved nature of these motifs, despite the stringent curation of MT_PURE sequences to exclude database matches, underscores their potential as fundamental elements within this dataset. The repetition of these motifs across multiple sequences suggests shared functional roles or evolutionary origins, further supporting their significance (Löytynoja & Goldman, 2008).

The TOMTOM analysis highlighted minimal overlap between MT_PURE motifs and known motifs in existing databases, further emphasizing the originality of these sequences. Some partial matches were observed with motifs associated with ssDNA viral replication, but the majority of MT_PURE motifs displayed configurations that diverged significantly from known patterns. This divergence reinforces the hypothesis that MT_PURE sequences represent novel genetic systems or uncharacterized extensions of ssDNA functionality (Guglielmini, de la Cruz, & Rocha, 2013).

The absence of significant matches also suggests that MT_PURE motifs may involve unique mechanisms of protein-DNA interaction or replication regulation, distinct from both ssDNA and dsDNA systems. These findings position MT_PURE as a rich source of genomic novelty, warranting further exploration.

Using the JASPAR databases provided additional context for the originality and potential functional roles of MT_PURE motifs. JASPAR contains a wide array of transcription factor binding motifs, primarily from eukaryotic systems, enabling a broader comparative analysis. While no exact matches were identified, partial similarities to eukaryotic transcription factor motifs were observed, particularly in palindromic and AT-rich configurations.

The partial similarity to eukaryotic motifs raises intriguing possibilities about the evolutionary convergence of DNA-protein interaction mechanisms (Löytynoja & Goldman, 2008). Such similarities may indicate analogous functional requirements, such as DNA stabilization or protein-binding specificity, even across vastly different genomic architectures. However, the distinct configurations of MT_PURE motifs, particularly their simplified structures, suggest they fulfill these roles through unique molecular strategies.

The results from MEME, TOMTOM, and JASPAR analyses collectively reinforce the originality and functional potential of MT_PURE motifs. The conserved and novel nature of these motifs highlights their possible roles in replication, stability, and protein interactions, all of which are critical for ssDNA systems. The divergence of these motifs from known patterns also supports their classification as components of an uncharacterized ssDNA genomic framework.

The minimal overlap with known motifs from TOMTOM and JASPAR underscores the novelty of MT_PURE motifs, suggesting evolutionary innovation in their functional roles. This lack of homology also points to a unique evolutionary lineage for MT_PURE sequences, potentially distinct from known ssDNA and dsDNA systems. The shared conservation across MT_PURE sequences further supports the hypothesis of a cohesive genomic framework with specialized adaptations.

The comprehensive analysis of MT_PURE motifs using MEME, TOMTOM, and JASPAR databases has significant implications for our understanding of genomic diversity and evolution. These findings highlight the potential of MT_PURE sequences to expand current paradigms of ssDNA functionality, particularly in terms of replication and protein interactions (Flynn & Zou, 2010).

Future studies should prioritize experimental validation of these motifs to determine their precise functional roles and their interactions with replication machinery or other proteins. Comparative analyses with other uncharacterized ssDNA datasets may also provide additional context for the evolutionary and functional innovation observed in MT_PURE.

### Comparative Analysis of MEME Results for MT_TOTAL and MT_PURE and Their JASPAR Database Comparisons

The comparative analysis of MEME results for MT_TOTAL and MT_PURE, along with their motif comparisons across JASPAR databases, provides a comprehensive perspective on the structural and functional diversity of these datasets. This joint analysis highlights the distinctions and overlaps between the two datasets, emphasizing their unique contributions to ssDNA genomic studies and their potential roles in expanding the current understanding of motif functionality and evolutionary innovation.

The MEME analysis revealed distinct patterns of motif diversity and conservation between MT_TOTAL and MT_PURE. MT_TOTAL, derived from a broader assembly of R1 sequences, exhibited a wider array of motif types, including complex and less conserved patterns. In contrast, MT_PURE, curated for unmatched sequences, displayed fewer motifs with greater conservation.

MT_TOTAL’s motif diversity underscores the functional complexity of this dataset. The identification of diverse repetitive elements and unique configurations suggests a dynamic interplay of sequences that may facilitate broader adaptive capacities or functional versatility. This aligns with findings in microbial ssDNA systems, where high motif variability is often associated with environmental adaptability (Tokuda & Shintani, 2024).

MT_PURE motifs, characterized by simplicity and AT-rich compositions, reflect a focus on structural and functional stability. The conserved nature of these motifs indicates critical roles in replication or protein-DNA interactions, akin to conserved elements in ssDNA viral genomes. These findings parallel observations in extreme environment-adapted ssDNA systems, where conserved motifs are linked to genomic integrity and replicative efficiency (Ando, y otros, 2021).

### Comparative Insights from JASPAR Database Analyses

The comparison of MT_TOTAL and MT_PURE motifs with the JASPAR databases provided valuable context for their functional and evolutionary relevance. While both datasets displayed minimal overlap with known transcription factor binding motifs, the patterns of partial matches offered insights into potential convergent mechanisms of DNA-protein interactions.

MT_TOTAL and JASPAR: MT_TOTAL motifs exhibited partial similarities to transcription factor motifs involved in regulatory complexity, such as those found in eukaryotic systems. This suggests the potential for regulatory innovation or structural roles beyond typical ssDNA functionality. Notably, certain palindromic motifs in MT_TOTAL displayed weak matches to motifs implicated in chromatin remodeling, indicating possible interactions with structural proteins (Längst & Manelyte, 2015).

MT_PURE and JASPAR: In contrast, MT_PURE motifs showed limited similarities to known motifs, reflecting their novelty and high specificity. The absence of significant overlaps reinforces the hypothesis that MT_PURE represents an uncharacterized genomic framework, potentially adapted to unique biological or environmental contexts. The conserved motifs in MT_PURE likely fulfill essential roles distinct from conventional transcription factor binding, potentially serving as replication origins or stabilizing elements (Abou Assi, Garavís, González, & Damha, 2018).

The comparative analyses underscore the complementary roles of MT_TOTAL and MT_PURE in advancing our understanding of ssDNA genomic diversity. MT_TOTAL’s broad motif diversity highlights its potential to capture adaptive and regulatory complexity, while MT_PURE’s focus on conserved motifs emphasizes structural and functional stability. Together, these datasets provide a holistic view of ssDNA innovation, with implications for understanding evolutionary processes and molecular mechanisms.

The limited overlap with JASPAR databases highlights the originality of both datasets and their potential to expand the known repertoire of DNA motifs. The distinct patterns of partial matches suggest that while some functional analogies may exist, the underlying mechanisms and contexts of motif functionality in MT_TOTAL and MT_PURE are likely unique. This opens avenues for further research into uncharacterized protein-DNA interaction strategies, supported by advancements in visualization and characterization techniques (Willaert & Kasas, 2022).

The combined results from MEME and JASPAR analyses highlight the unique evolutionary and functional characteristics of the MT_TOTAL and MT_PURE datasets. The conserved motifs identified in MT_PURE suggest a phylogenetic linkage among its sequences, pointing to a shared evolutionary origin. This conservation, alongside the broader diversity seen in MT_TOTAL, indicates the presence of a highly adaptable and potentially versatile genomic system.

When considered together, the motifs from both datasets suggest a genomic framework capable of self-replication, a hallmark of ssDNA systems, but with innovations that distinguish them from known ssDNA and dsDNA forms. A key question raised by these results is whether these sequences exist primarily in an extracellular or intracellular context. While the simplicity and stability of MT_PURE motifs, along with the absence of interspersed repeats, suggest adaptations consistent with extracellular survival, this would represent a highly atypical scenario for ssDNA sequences of this complexity. Typically, extracellular ssDNA entities, such as viruses, are compact and specialized, whereas MT_TOTAL and MT_PURE demonstrate a broader and more conserved functional repertoire.

If these sequences are indeed extracellular, it implies a novel genomic strategy, leveraging autoreplicative capabilities and structural versatility to persist in challenging environments. Conversely, if intracellular, the motifs and conserved elements may point to a highly efficient and adaptable system for survival and replication within host cells. Future research must clarify this distinction through experimental validation of their replication mechanisms and ecological context.

The minimal overlap with known motifs in JASPAR underscores their evolutionary novelty (Zheng, Li, & Hu, 2015), suggesting that MT_TOTAL and MT_PURE represent genomic architectures that challenge existing paradigms. Their potential autoreplicative capacity, inferred from conserved replication-associated motifs, highlights a survival strategy that may combine efficiency with resilience, allowing these sequences to thrive under diverse conditions.

Future directions should focus on elucidating the exact roles of these motifs through experimental validation and structural studies. Understanding their protein-DNA interaction dynamics and replication mechanisms could provide transformative insights into the evolutionary trajectories of ssDNA genomes and their unique adaptations. These findings may also inform the design of synthetic genomic systems or the exploration of novel genetic elements in uncharacterized biological niches.

### RNAfold Analysis Results for MT_TOTAL ssDNA Sequences

The RNAfold analysis of MT_TOTAL ssDNA sequences revealed significant insights into their secondary structural potential, highlighting features that could underpin unique functional and evolutionary roles. The predicted folding patterns and associated thermodynamic stability metrics point to the versatility of these sequences, suggesting their capacity to engage in regulatory and replicative functions similar to RNA molecules but within a ssDNA framework.

The RNAfold analysis predicted stable secondary structures in a majority of MT_TOTAL sequences, characterized by hairpins, stem-loops, and palindromic regions. These features are often associated with replication origins and regulatory sites in ssDNA viruses, where they facilitate protein binding and enzymatic processes critical for genome replication and stability. The high proportion of sequences forming such structures underscores the functional potential of MT_TOTAL, suggesting adaptations that may enhance their efficiency in replication or interaction with host or environmental factors (Binet, Padiolleau-Lefèvre, Octave, Avalle, & Maffucci, Comparative Study of Single-stranded Oligonucleotides Secondary Structure Prediction Tools., 2023).

Interestingly, the thermodynamic stability of these predicted structures varied widely across the dataset, with Gibbs free energy (ΔG) values ranging from highly stable to moderately unstable conformations. This variability may indicate a dual functional repertoire, where highly stable structures support long-term persistence or environmental resilience, while less stable configurations enable dynamic interactions, such as transient protein binding or strand displacement during replication. These findings align with studies of ssDNA viruses, where secondary structure flexibility is a critical determinant of functional versatility (Golden, Murrell, Martin, Pybus, & Hein, 2020).

The structural patterns observed in MT_TOTAL also suggest potential parallels with RNA molecules in terms of regulatory complexity. Secondary structures such as hairpins and loops are known to mediate gene expression and stability in RNA systems, and their presence in MT_TOTAL sequences raises intriguing possibilities about ssDNA elements mimicking RNA-like functions. This hypothesis is further supported by the absence of significant interspersed repeats or transposable elements, indicating a streamlined genome architecture where secondary structure could play a compensatory regulatory role (Vassetzky & Kramerov, 2013).

Comparative analysis with known ssDNA systems highlights the originality of MT_TOTAL sequences. While certain structural motifs are consistent with replication-associated elements in ssDNA viruses, the overall diversity and prevalence of secondary structures in MT_TOTAL exceed typical patterns observed in such systems. This suggests that MT_TOTAL may represent a novel genomic category or an uncharacterized extension of ssDNA functionality. The high AT content of these sequences likely contributes to their structural repertoire, as AT-rich regions are known to favor the formation of flexible and energetically favorable secondary structures (Wegrzyn, y otros, Sequence-specific interactions of Rep proteins with ssDNA in the AT-rich region of the plasmid replication origin., 2014).

The broader implications of these findings extend to understanding the evolutionary and functional innovation in ssDNA systems. The ability to form RNA-like secondary structures may reflect an evolutionary strategy to enhance functional adaptability, particularly in challenging or variable environments. Studies have shown that base-pairing interactions in nucleic acid secondary structures play a pivotal role in driving evolutionary innovation and functional optimization in compact genetic systems (Golden, Murrell, Martin, Pybus, & Hein, 2020). This versatility, coupled with their potential autoreplicative capacity inferred from conserved motifs, positions MT_TOTAL as a genomic framework with significant implications for understanding the evolution of compact and efficient genetic systems.

Future research should prioritize experimental validation of these predicted structures, particularly their roles in replication and interaction with proteins or other nucleic acids. Additionally, exploring the environmental or cellular contexts where such sequences thrive could provide insights into the selective pressures driving their structural and functional evolution. These efforts would not only deepen our understanding of MT_TOTAL but also contribute to broader paradigms in genomic architecture and molecular innovation.

### RNAfold Analysis Results for MT_PURE ssDNA Sequences

The RNAfold analysis of MT_PURE ssDNA sequences revealed critical insights into their structural and functional potential, underscoring their unique characteristics and reinforcing their distinction from known genomic systems. The highly curated nature of the MT_PURE dataset, enriched for unmatched sequences, makes the structural predictions particularly significant, as they likely reflect fundamental properties of these novel ssDNA elements.

The RNAfold predictions identified highly stable secondary structures across the majority of MT_PURE sequences, characterized predominantly by stem-loops, hairpins, and palindromic configurations. These structural features are hallmark indicators of replication origins and regulatory sites, suggesting that MT_PURE sequences may perform roles analogous to ssDNA viral genomes but with potentially novel mechanisms (Wegrzyn, y otros, Sequence-specific interactions of Rep proteins with ssDNA in the AT-rich region of the plasmid replication origin., 2014). The strong thermodynamic stability of these structures, as evidenced by Gibbs free energy (ΔG) calculations, further supports their functional relevance, implying robust interactions with proteins or other nucleic acids. This is consistent with findings that conserved secondary structures within ssDNA viral genomes are integral to their replication and regulatory functions (Muhire, y otros, 2014)

Compared to MT_TOTAL, the secondary structures of MT_PURE sequences displayed greater uniformity and higher thermodynamic stability. This distinction reflects the impact of stringent filtering during dataset curation, resulting in a collection of sequences that prioritize conservation and essential functionality over diversity. Such uniformity suggests that MT_PURE motifs may represent a conserved core of regulatory or replicative elements optimized for stability and efficiency (Binet, Padiolleau-Lefèvre, Octave, & Avalle, Comparative Study of Single-stranded Oligonucleotides Secondary Structure Prediction Tools., 2023).

The predicted structural configurations in MT_PURE align with known functions of secondary structures in ssDNA systems. For instance, the prevalence of palindromic sequences could facilitate strand folding into hairpins, a feature critical for replication initiation and protein binding. Additionally, the predominance of AT-rich regions in MT_PURE contributes to the flexibility and dynamic adaptability of these secondary structures, further enhancing their functional potential (Irony-Tur Sinai, y otros, 2019).

Interestingly, despite their simplicity, the structures identified in MT_PURE exhibit functional parallels with complex regulatory elements in RNA genomes. This raises the intriguing possibility that MT_PURE sequences could mimic RNA-like regulatory behaviors, leveraging secondary structure dynamics to mediate interactions with molecular partners. Such mimicry would represent an evolutionary convergence, highlighting the adaptability of ssDNA systems in adopting strategies typically associated with RNA (Stedman, 2013).

The uniqueness of MT_PURE sequences is further emphasized by their lack of significant overlap with known ssDNA or dsDNA systems. The absence of repetitive elements, coupled with the conservation of secondary structures, suggests a streamlined genomic architecture that prioritizes stability and functionality (Padeken, Zeller, & Gasser, 2015). This combination of features aligns with theoretical models of genomic minimalism, where simplicity and efficiency are favored in extreme or specialized environments (Malathi & Renuka Devi, 2019).

The broader implications of these findings underscore the originality and evolutionary significance of MT_PURE. The predicted RNAfold structures not only highlight their functional potential but also position them as candidates for representing a novel class of ssDNA systems. The stability and uniformity of these structures suggest adaptations that may facilitate survival and replication under diverse conditions, reinforcing the hypothesis of MT_PURE’s distinct evolutionary lineage.

Future research should focus on validating these predicted structures through experimental approaches, such as probing interactions with host or environmental factors. Understanding the molecular dynamics of MT_PURE secondary structures will be critical for unraveling their roles in replication, regulation, and potential interactions with other genomic elements. Additionally, comparative studies with other uncharacterized ssDNA datasets could shed light on the evolutionary pressures shaping these sequences and their functional innovations.

### Complete comparative analysis report of MT_TOTAL vs. MT_PURE ssDNA sequences

The comparative analysis of MT_TOTAL and MT_PURE ssDNA sequences highlights their distinct structural, functional, and evolutionary characteristics. While both datasets originate from the shared "no-hits" genomic zone, they reflect different curation approaches and potential functional roles. MT_TOTAL, as a broader dataset, includes sequences with minor database matches, offering a diverse range of functional motifs and secondary structures. This inclusivity results in a heterogeneous collection enriched with less conserved elements and potential adaptive flexibility (Kim & Lee, 2021). Conversely, MT_PURE is a curated dataset focused solely on sequences with no detectable matches, narrowing its scope to highly unique, conserved elements that likely serve essential functional roles (Nüesch, y otros, 2024).

RNAfold analysis revealed distinct patterns between the two datasets. MT_TOTAL sequences exhibit a wide array of secondary structures, from highly stable to moderately unstable, reflecting functional versatility. In contrast, MT_PURE sequences display thermodynamically stable configurations, emphasizing genomic efficiency and structural resilience. These differences underscore the specialized roles that MT_PURE sequences may play within the broader ssDNA genomic landscape.

MEME and TOMTOM motif analyses further distinguish the datasets. MT_TOTAL contains a spectrum of conserved and novel motifs, including elements with weak matches to known regulatory sites. This suggests adaptive interactions with various hosts or environments. MT_PURE, however, is dominated by conserved motifs, such as palindromic structures and hairpins, linked to replication and stabilization. The lack of overlap with known databases highlights the novelty of MT_PURE motifs, positioning them as candidates for exploring uncharacterized ssDNA functionality (Malathi & Renuka Devi, 2019) (Dávila-Ramos, y otros, 2019).

The adaptability of ssDNA viruses to extreme environments provides a compelling framework to contextualize the distinct characteristics of MT_TOTAL and MT_PURE sequences. As highlighted by Dávila-Ramos et al. (Dávila-Ramos, y otros, 2019), ssDNA viruses exhibit a remarkable ability to thrive in extreme conditions, such as high salinity, temperature fluctuations, and radiation. These adaptations often involve streamlined genomes enriched with conserved motifs, which contribute to replication stability and resilience under stress. This mirrors the characteristics of MT_PURE, which is dominated by highly conserved motifs such as palindromic structures and hairpins associated with replication and structural stabilization. These conserved elements likely represent genomic strategies optimized for survival in harsh conditions.

Conversely, ssDNA viruses also exhibit genetic plasticity, enabling them to interact with diverse host systems and adapt to fluctuating environmental pressures. This adaptability is driven by broader genomic diversity and the presence of novel functional motifs, as noted in MT_TOTAL sequences. For instance, the inclusion of partially conserved elements and weak matches to known regulatory motifs in MT_TOTAL reflects a potential for dynamic interactions, akin to the flexibility observed in ssDNA viruses described in metagenomic studies of extreme environments. The ability of MT_TOTAL sequences to incorporate novel elements and display structural versatility suggests a genomic strategy that balances exploration of new adaptive pathways with functional utility.

Moreover, the review by Dávila-Ramos et al. (Dávila-Ramos, y otros, 2019) emphasizes that ssDNA viruses often occupy unique ecological niches, where they mediate important interactions within microbial communities. This ecological adaptability could parallel the evolutionary trajectories of MT_TOTAL sequences, which, with their heterogeneity and partial database matches, may represent transitional genomic systems capable of bridging different ecological or functional contexts.

The evolutionary trajectories inferred from these findings reflect complementary roles. MT_TOTAL, with its diverse sequences and partial database matches, may represent an intermediate or transitional genomic system, integrating novel elements and adapting to environmental pressures. Meanwhile, MT_PURE, with its stringent curation and conserved motifs, represents a specialized lineage optimized for efficiency and resilience. This balance between diversity and conservation mirrors the evolutionary strategies observed in ssDNA viruses, which combine genetic flexibility and structural constraints to optimize survival and replication (Torralba, Blanc, & Michalakis, 2024) (Golden, Murrell, Martin, Pybus, & Hein, 2020).

Together, these datasets provide a holistic perspective on ssDNA innovation, advancing our understanding of genomic evolution in extreme contexts.

### Open Questions: Extracellular vs. Intracellular Origin

The structural and functional characteristics of these sequences also raise intriguing questions about their potential extracellular or intracellular origins. The high stability and simplicity of the secondary structures in both datasets, particularly in MT_PURE, suggest adaptations consistent with extracellular survival (Podgornaya, Vasilyeva, & Bespalov, 2016) (Nüesch, y otros, 2024). Features such as hairpins and palindromic motifs are commonly associated with replication origins in extracellular systems like viruses, which require robust mechanisms to withstand environmental challenges. Additionally, the absence of interspersed repeats and the streamlined genomic architecture further support the possibility of an extracellular context (Podgornaya, Vasilyeva, & Bespalov, 2016).

However, an intracellular origin cannot be ruled out. The diversity in MT_TOTAL and the conservation in MT_PURE could reflect interactions with host systems or specialized cellular roles. If these sequences are indeed intracellular, the prevalence of eukaryotic signatures in the Krona analysis—and their apparent association with human-related sequences—poses a unique challenge. Given the extensive genomic studies conducted in humans, the discovery of such unique ssDNA elements within human cells would be highly unexpected and suggest either a previously unidentified genomic feature or an external interaction yet to be characterized (Kim & Lee, 2021).

If these sequences are extracellular, their originality and high stability position them as potential candidates for representing a novel genomic strategy, leveraging autoreplicative capabilities to thrive in diverse environments. On the other hand, if intracellular, their uniqueness and conservation imply a specialized functionality, potentially involving host-specific adaptations or interactions. Both scenarios underscore the evolutionary and functional novelty of these datasets, warranting further experimental investigation to clarify their ecological and biological roles.

Broader implications of this comparative analysis extend to understanding the versatility and adaptability of ssDNA systems. The coexistence of diversity in MT_TOTAL and conservation in MT_PURE highlights evolutionary strategies that balance functional innovation with stability. These findings raise intriguing questions about the ecological roles and evolutionary origins of these sequences, particularly their capacity for autoreplication and adaptation to extreme or variable conditions. Future research should focus on validating the functional roles of these motifs, exploring their interactions with host systems or environmental factors, and investigating their evolutionary significance.

### Functional Evaluation of MT_TOTAL and MT_PURE ssDNA Sequences

The functional evaluation of MT_TOTAL and MT_PURE ssDNA sequences using predictive tools such as Prokka has provided valuable insights into their potential biological roles. These analyses enabled the identification of putative functional categories, shedding light on the evolutionary and ecological significance of these sequences. This discussion summarizes the findings and their implications, focusing on both datasets across the three domains of life.

### Comprehensive Analysis Report on MT_TOTAL ssDNA Sequences Across Three Domains

The functional analysis of MT_TOTAL sequences revealed a diverse array of predicted functional categories, consistent with the dataset’s broader inclusion criteria. Many sequences were annotated with functions related to replication, transcription regulation, and metabolic processes (Witte, Urbanke, & Curth, 2003) (Kim & Blair, 2015). These findings are indicative of a dataset encompassing a wide range of genomic elements, some of which may interact with host systems or function independently in various ecological contexts.

A significant proportion of predicted functions in MT_TOTAL were associated with prokaryotic systems, particularly genes linked to DNA replication and repair. This is consistent with the hypothesis that some MT_TOTAL sequences may share evolutionary origins with bacterial or archaeal systems, possibly through horizontal gene transfer or adaptation to extreme environments. These functional predictions align with known roles of ssDNA elements in facilitating genetic exchange and replication in prokaryotic domains (Marceau, 2012).

Notably, some sequences in MT_TOTAL displayed annotations suggestive of regulatory elements, such as transcription factor binding sites and operon structures (Gama-Castro, y otros, 2016). These features highlight the potential for MT_TOTAL sequences to act as regulatory modules, influencing gene expression within host systems or contributing to environmental genomic plasticity. The presence of such regulatory elements suggests that MT_TOTAL sequences may occupy a transitional space between coding and non-coding genomic functions, underscoring their functional versatility (Hu, Chen, Wang, Yang, & Ding, 2024).

In eukaryotic contexts, predicted functions included elements related to structural stability and RNA-like regulatory mechanisms (Pal & Levy, 2019). These annotations, although less prevalent, raise intriguing possibilities about interactions with eukaryotic hosts or adaptation to eukaryotic cellular environments. However, the limited match to known eukaryotic elements underscores the novelty of MT_TOTAL and suggests that these sequences may represent uncharacterized genomic innovations.

### Comprehensive Analysis Report on MT_PURE ssDNA Sequences Across Three Domains

The functional analysis of MT_PURE sequences, while more focused due to the dataset’s stringent curation, provided insights into conserved and essential functionalities. MT_PURE sequences were predominantly annotated with functions linked to replication origins, structural motifs, and protein-DNA interactions (Wegrzyn, y otros, 2023). These findings are consistent with the hypothesis that MT_PURE represents a streamlined and highly specialized subset of ssDNA sequences.

The majority of functional annotations in MT_PURE were associated with prokaryotic systems, particularly replication-related motifs and conserved structural elements. This reflects the dataset’s high level of conservation and suggests potential roles in replication efficiency and genomic stability. Such functionalities are hallmarks of ssDNA elements in prokaryotic domains, where simplicity and efficiency are evolutionary priorities (Wegrzyn, y otros, 2023).

Annotations linked to viral systems were also observed, particularly motifs resembling replication origins in ssDNA viruses (Malathi & Renuka Devi, 2019). This suggests that MT_PURE sequences may share functional or structural similarities with viral genomes, potentially reflecting convergent evolutionary strategies or shared ecological niches. These findings support the notion that MT_PURE may represent a novel genomic system with parallels to known viral or plasmid-like elements (Kazlauskas, y otros, 2017).

In contrast to MT_TOTAL, functional annotations in MT_PURE with eukaryotic affinities were minimal. This reinforces the idea that MT_PURE sequences are more specialized and less likely to interact with or integrate into eukaryotic systems. The absence of complex regulatory annotations further highlights the streamlined nature of MT_PURE, emphasizing its potential role as a core replicative or structural genomic framework.

### Broader Implications

The functional evaluations of MT_TOTAL and MT_PURE underscore their complementary roles in advancing our understanding of ssDNA diversity and innovation. MT_TOTAL, with its broader functional repertoire, highlights the adaptive and regulatory potential of ssDNA sequences, suggesting roles in genomic plasticity and environmental interactions. MT_PURE, on the other hand, exemplifies a conserved and specialized genomic system, optimized for efficiency and stability.

The predicted functional categories across both datasets reveal significant evolutionary implications. The prokaryotic dominance in functional annotations points to ancestral ties with bacterial or archaeal systems (Dávila-Ramos, y otros, 2019), while the viral-like motifs in MT_PURE suggest convergent evolution or functional overlap with known ssDNA viral genomes. These findings contribute to broader discussions about the origins and roles of ssDNA elements in shaping genomic and ecological landscapes.

Future research should focus on experimental validation of the predicted functions, particularly in the context of replication, regulation, and host interactions. For instance, studies have elucidated the DNA-binding properties of the human TRIP4 ASCH domain, providing insights into its functional role (Hu, Chen, Wang, Yang, & Ding, 2024).

Additionally, research on replication protein A has shed light on its interaction with ssDNA and its role in DNA metabolic processes (Chadda, y otros, 2024). Comparative studies with known ssDNA systems could further elucidate the evolutionary trajectories of MT_TOTAL and MT_PURE, providing deeper insights into their unique genomic architectures and functional innovations.

### Final Comment or Conclusions

The findings presented in this study provide compelling evidence for the originality and exceptional nature of the MT_TOTAL and MT_PURE ssDNA sequences. The functional analyses, structural evaluations, and comparative insights reinforce their novelty, distinguishing them from known ssDNA and dsDNA systems (Lin, Malik, & Guo, 2021). This originality is supported by the unique motifs, secondary structures, and predicted functionalities that challenge existing paradigms of genomic diversity and adaptation.

The stability and efficiency observed in these sequences, particularly in MT_PURE, point to adaptations consistent with survival in challenging environments. These adaptations could align with extracellular systems, such as viruses or free genomic particles, which often rely on compact, streamlined genomes for resilience (Forterre & Prangishvili, 2009) (Malathi & Renuka Devi, 2019). However, the absence of interspersed repeats typical of intracellular genomes and the identification of conserved replication-associated motifs support the notion that these sequences may represent a novel genomic strategy, possibly reflecting an origin in a unique ecological or abiotic context.

In addition to the unique adaptations observed in these sequences, meteorites have been identified as sources of a wide variety of extraterrestrial nucleobases, including purines and pyrimidines. Investigations into carbonaceous chondrites have revealed these essential components of nucleic acids, supporting the hypothesis that meteorites contributed significantly to the pool of building blocks necessary for the emergence of life on early Earth. This diversity of nucleobases not only underscores the chemical richness of meteorites but also raises the possibility that such molecular precursors could have been key to the development of primitive genetic systems. These findings align with the presence of the unique ssDNA sequences described here, further emphasizing the potential role of meteorites as reservoirs and facilitators of prebiotic chemistry (Callahan, y otros, 2011).

Furthermore, meteorites are not merely passive carriers of organic molecules but may actively facilitate key chemical reactions relevant to prebiotic synthesis. A study published in Proceedings of the National Academy of Sciences (Ferus M, 2015) demonstrated that meteorites can catalyze the formation of nucleosides, fundamental building blocks of RNA and DNA, under conditions analogous to those of the early Earth. This catalytic role supports the hypothesis that meteorites contribute not only molecular precursors but also promote the chemical pathways necessary for the emergence of life. In light of this, the discovery of unique single-stranded DNA sequences in meteorites extends the understanding of their role as not only sources but also potential facilitators of molecular evolution, possibly driving the assembly of early genomic structures (Ferus M, 2015).

Alternatively, an intracellular origin, while less supported, remains a possibility. Certain annotations hint at potential interactions with host systems, suggesting these sequences may have been co-opted for specialized roles within intracellular environments. However, the association with human-related sequences identified by Krona analysis remains speculative and requires further validation to rule out contamination or secondary alignment artifacts. The absence of similar sequences in extensively characterized human genomes emphasizes the originality of these findings. The ENCODE (Encyclopedia of DNA Elements) project has provided a comprehensive map of functional elements in the human genome, including protein-coding regions, regulatory elements, and non-coding RNAs. Despite this extensive characterization, no ssDNA elements with the unique properties described in this study have been reported, emphasizing the exceptional nature of these findings (ENCODE Project Consortium).

The revolutionary nature of the hypothesis that these sequences could represent a novel extracellular genomic system lies in its implications for biology and evolution. Such a discovery would challenge conventional views on the boundaries of life and genetic systems, suggesting new mechanisms of survival, replication, and interaction in extreme or variable environments. These findings emphasize the importance of further experimental studies to validate their functional roles, ecological contexts, and evolutionary origins.

In conclusion, the MT_TOTAL and MT_PURE sequences represent a significant step forward in uncovering uncharted genomic territory. Whether extracellular or intracellular, their originality and adaptability underscore their potential to redefine current understandings of ssDNA systems and their roles in nature. Future research will be essential in unraveling their mysteries, potentially opening new avenues in genomics, evolutionary biology, and molecular innovation.

The exceptional nature of these sequences is magnified by their origin: cultures derived from meteorite fragments. This unprecedented source not only highlights the uniqueness of the sequences but also their potential significance in broader scientific contexts. If these findings are validated, they could open doors to transformative discussions in biology and astrobiology. The possibility that such sequences might have an extraterrestrial origin opens the door to intriguing scientific discussions, including the panspermia hypothesis. This theory posits the distribution of life’s building blocks across the universe via celestial bodies. While this study does not provide definitive evidence, the unique characteristics of these sequences invite further investigation into their origins.

To ensure the robustness of these findings, repeating the methodologies applied in this study will be critical. Independent verification of these results, using similar or advanced techniques, could confirm whether these sequences indeed represent extraterrestrial genetic elements. If validated, this study would mark a pivotal moment in scientific history, bridging gaps between molecular biology, evolutionary theory, and our understanding of life’s potential distribution across the cosmos.

### Theoretical Reflections: Connection to the Universal Constructor

The MT_TOTAL and MT_PURE sequences also invite reflection on their potential relationship with the concept of the Universal Constructor, as introduced by John von Neumann. This theoretical framework describes a system capable of self-replication while maintaining the information necessary to reconstruct itself—a hallmark of both life and evolutionary complexity. The exceptional motifs and stable secondary structures identified in these sequences suggest they may operate under similar principles, facilitating replication and interaction in an open evolutionary framework (Binet, Padiolleau-Lefèvre, Octave, Avalle, & Maffucci, Comparative Study of Single-stranded Oligonucleotides Secondary Structure Prediction Tools., 2023). Studies have shown that single-stranded DNA (ssDNA) can form stable secondary structures, such as hairpins and G-quadruplexes, which play crucial roles in replication and other cellular processes (Bonnet, Krichevsky, & Libchaber, 1998). Such features align with the theoretical constraints outlined for living systems, where information and structure converge to enable adaptability and survival (Solé, y otros, 2024).

The unique features of these sequences, including their adaptability and persistence in variable or extreme environments, align with principles of open-ended evolution. These characteristics resonate with the concept of the Universal Constructor, which describes systems capable of self-replication and information retention. While their potential extraterrestrial origin remains speculative, further research could explore their alignment with these theoretical principles, bridging empirical observations with broader evolutionary frameworks. This connection offers profound implications for understanding life’s universality and its capacity to evolve beyond Earth’s confines.

### Abiogenic Considerations and Cultivation Conditions

An intriguing possibility arising from these findings is the potential abiogenic origin of the MT_TOTAL and MT_PURE sequences under the cultivation conditions employed. While bases and short nucleotides have been identified in meteorites such as Murchison, these molecules are typically found in minimal quantities and lack the organized complexity observed in these ssDNA sequences (Martins, y otros, 2008). It is conceivable that the cultivation process provided a microenvironment conducive to abiogenic assembly, mimicking prebiotic conditions where nucleotides polymerize into functional chains. Factors such as temperature fluctuations, ionic concentrations, and aqueous chemistry could have facilitated the emergence of these sequences, offering a glimpse into pathways of molecular evolution in early Earth-like or extraterrestrial contexts (Becker, y otros, Science).

While this hypothesis is speculative, it aligns with experimental studies showing that short nucleotides can spontaneously assemble under certain conditions, forming structures reminiscent of primordial genetic systems (Becker, y otros, Science). However, there are critical limitations to this interpretation. First, there is no direct evidence within this study to confirm that the sequences were synthesized de novo during cultivation. Second, the potential for contamination, although mitigated by the exceptional characteristics of the sequences, cannot be entirely ruled out. The study’s results strongly argue against contamination based on the uniqueness and originality of the sequences, but this remains a cautionary note.

If further studies were to validate the abiogenic assembly of these sequences under cultivation, it could provide valuable insights into the plausibility of nucleotide formation and polymerization in extraterrestrial environments. Such findings would deepen our understanding of prebiotic chemistry and might contribute to discussions around panspermia as a potential mechanism for distributing life’s building blocks across the universe. However, additional experimental evidence is needed to support this hypothesis.

Several prominent scientists, including Francis Crick and Stephen Hawking, have supported the panspermia hypothesis, which suggests that life may have arrived on Earth from outer space, transported via meteorites or comets that impacted the early planet (Crick & Orgel, 1973) (Hawking, 2001). This theory highlights the possibility that the building blocks of life, or even microbial organisms, could have originated elsewhere in the universe and been seeded on Earth. Recent discoveries, such as organic molecules on the comet 67P/Churyumov–Gerasimenko by the Rosetta mission (Altwegg, y otros, 2016) and amino acids in carbonaceous meteorites like Murchison (Oba, y otros, 2022), further support the idea that extraterrestrial bodies can deliver complex organic material. However, to think that this process occurred during a single, finite period of Earth’s history may oversimplify the complexity of the panspermia concept.

I propose a new perspective, termed Live Panspermia, which posits that the exchange of genetic material or precursors of life from extraterrestrial sources is an ongoing and dynamic process. Rather than assuming life arrived in a single primordial form with immense potential, Live Panspermia suggests that life—or its genetic precursors—may continue to arrive periodically. The discovery of unique ssDNA sequences associated with meteorite samples could represent evidence of this ongoing process, offering a new perspective on the dynamic nature of panspermia and its implications for the continual exchange of genetic material across cosmic environments.

## Conclusions

This study represents the first comprehensive investigation into the genomic characteristics of single-stranded DNA (ssDNA) sequences extracted from meteorite-derived samples. Employing advanced metagenomic techniques and rigorous bioinformatics filtering, we identified and characterized novel ssDNA elements with unique structural, functional, and evolutionary features. The findings underscore the originality and potential biological significance of these sequences, opening new avenues for understanding genomic diversity, replication mechanisms, and molecular innovation.

Key findings.

1. Discovery of novel genetic elements:

- The "no-hits zone," comprising sequences unmatched in current genomic databases, accounted for a substantial proportion (∼51%) of the data, emphasizing the vast originality of the identified ssDNA sequences.
- The highly curated MT_PURE dataset revealed conserved and thermodynamically stable motifs, while the broader MT_TOTAL dataset captured a diverse array of structural and functional patterns, highlighting complementary aspects of ssDNA innovation.
2. Structural and functional novelty:

- The AT-rich nature of these ssDNA sequences and their distinctive repetitive motifs suggest adaptations favoring genomic flexibility, stability, and potentially autoreplicative mechanisms.
- Secondary structure predictions, including hairpins and palindromic regions, imply roles in replication and protein binding, drawing parallels with ssDNA viruses while showcasing significant evolutionary divergence.
3. Evolutionary and ecological implications:

- The identified sequences may represent an uncharacterized genomic framework with potential extracellular or intracellular origins. Their originality raises questions about their ecological roles and evolutionary trajectories, particularly in extreme or extraterrestrial environments.
- The lack of homology with terrestrial genomes and the unique compositional features of the sequences support their potential extraterrestrial origin, warranting further exploration.
4. Broader scientific context:

- The study contributes to ongoing discussions in astrobiology regarding the origins of life and the potential for genetic material exchange through processes such as panspermia. The discovery of ssDNA sequences in meteorite-derived samples provides a foundation for investigating the dynamic interaction between Earth’s biosphere and the broader cosmos.

## Future directions

While this research provides groundbreaking insights, it also raises several open questions that demand further investigation:

- The precise origin and ecological roles of these ssDNA sequences remain to be determined.
- Experimental validation of their autoreplicative capacity, protein interactions, and potential regulatory functions is essential.
- Expanding the database of uncharacterized genetic elements will enable more robust comparative analyses, addressing gaps that limit current interpretations.

The findings presented here challenge conventional paradigms in genomic diversity and molecular evolution, bridging the fields of molecular biology, astrobiology, and evolutionary genetics. They underscore the importance of continued exploration of uncharacterized genomic elements to uncover the mechanisms and origins of life’s complexity. By shedding light on these enigmatic ssDNA sequences, this study offers a steppingstone for transformative discoveries, potentially redefining our understanding of life’s diversity and its distribution across the universe.

## Implications and transition to limitations

While these findings pave the way for transformative discoveries, certain methodological and interpretative limitations must be acknowledged to contextualize the results accurately. These limitations provide a framework for guiding future research efforts and refining the experimental approach to further validate and expand upon the conclusions of this study.

## Study limitations

Despite the controls implemented, this study faces certain limitations inherent to the nature of the experiments conducted. First, although stringent protocols were followed to minimize contamination, the possibility that the detected ssDNA fragments are of terrestrial origin cannot be completely ruled out, especially given the sensitivity of sequencing techniques. Second, the lack of matches in genomic databases for sequences in the "no-hits zone" presents challenges in determining their exact origin and function. This phenomenon could reflect gaps in current databases rather than entirely novel structures. Additionally, while the cultivation assays in minimal media suggest extremophilic adaptations, it cannot be conclusively determined that these features are intrinsic to the meteorites rather than laboratory artifacts. Finally, the interpretation of phylogenetic data and repetitive motif analyses would benefit from additional analyses using expanded datasets and functional validation experiments, which will be addressed in future research.

## Supporting information

This document includes the predicted secondary structures of ssDNA sequences analyzed in this study. These structures were generated using RNAfold

## Acknowledgments

the author would like to thank Dr. Alfredo Repáraz Andrade and his team at the Genomics Laboratory for their invaluable assistance with the DNA extraction from the samples.

## Declarations

the author declares no conflict of interest.

## Data availability

The raw sequencing data and genome assemblies are being prepared for submission to a BioProject in GenBank and will be made publicly available upon completion.

## Notes

### Competing Interest Statement

The authors have declared no competing interest.

## References

Abou Assi, H., Garavís, M., González, C., & Damha, M. (2018). i-Motif DNA: structural features and significance to cell biology. Nucleic Acids Res., 46(16), 8038–8056. doi:10.1093/nar/gky735.

Altwegg, K., Balsiger, H., Bar-Nun, A., Berthelier, J., Bieler, A., Bochsler, P., . . . Le. (2016). Prebiotic chemicals-amino acid and phosphorus-in the coma of comet 67P/Churyumov-Gerasimenko. Sci Adv, 2(5), e1600285. doi:10.1126/sciadv.1600285. PMID: 27386550.

Ando, N., Barquera, B., Bartlett, D., Boyd, E., Burnim, A., Byer, A., . . . Watkins, M. (2021). The Molecular Basis for Life in Extreme Environments. Annu Rev Biophys., 50, 343–372. doi:10.1146/annurev-biophys-100120-072804.

Bailey TL, B. M. (2009). MEME SUITE: tools for motif discovery and searching. Nucleic Acids Res., 37(Web Server issue), W202-8. doi:10.1093/nar/gkp335.

Bankevich A, N. S. (2012). SPAdes: a new genome assembly algorithm and its applications to single-cell sequencing. J Comput Biol, 19(5), 455–77. doi:10.1089/cmb.2012.0021.

Becker, S., Feldmann, J., Wiedemann, S., Okamura, H., Schneider, C., Iwan, K., . . . Carell, T. (Science). Unified prebiotically plausible synthesis of pyrimidine and purine RNA ribonucleotides. 366(6461), 76-82. doi:10.1126/science.aax2747.

Binet, T., Padiolleau-Lefèvre, S., Octave, S., & Avalle, B. M. (2023). Comparative Study of Single-stranded Oligonucleotides Secondary Structure Prediction Tools. BMC Bioinformatics., 24(1), 422. doi:10.1186/s12859-023-05532-5.

Binet, T., Padiolleau-Lefèvre, S., Octave, S., Avalle, B., & Maffucci, I. (2023). Comparative Study of Single-stranded Oligonucleotides Secondary Structure Prediction Tools. BMC Bioinformatics, 24(1), 422. doi:10.1186/s12859-023-05532-5.

Bonnet, G., Krichevsky, O., & Libchaber, A. (1998). Kinetics of conformational fluctuations in DNA hairpin-loops. Proc Natl Acad Sci U S A., 95(15), 8602–6. doi:10.1073/pnas.95.15.8602.

Brandes, N., & Linial, M. (2016). Gene overlapping and size constraints in the viral world. Biol Direct., 11, 26. doi:10.1186/s13062-016-0128-3.

Callahan, M., Smith, K., Cleaves, H. 2., Ruzicka, J., Stern, J., Glavin, D., . . . Dworkin, J. (2011). Carbonaceous meteorites contain a wide range of extraterrestrial nucleobases. Proc Natl Acad Sci U S A., 108(34), 13995-8. doi:10.1073/pnas.1106493108.

Camacho, C., Coulouris, G., Avagyan, V., Ma, N., Papadopoulos, J., Bealer, K., & Madden, T. L. (2009). BLAST+: architecture and applications. BMC Bioinformatics, 10, 421. 10.1186/1471-2105-10-421

Cavicchioli, R., Siddiqui, K., Andrews, D., & Sowers, K. (2002). Low-temperature extremophiles and their applications.. Curr Opin Biotechnol., 13(3), 253–61. doi:10.1016/s0958-1669(02)00317-8.

Chadda, R. K., Ahmad, I., Deveryshetty, J., Holehouse, A., Sigurdsson, S., Biswas, G., . . . Antony, E. (2024). Partial wrapping of single-stranded DNA by replication protein A and modulation through phosphorylation. Nucleic Acids Res., 52(19), 11626–11640. doi:10.1093/nar/gkae584.

Cowan, D., Albers, S., Antranikian, G., Atomi, H., Averhoff, B., Basen, M., . . . Vorgias, K. (2024). Extremophiles in a changing world. Extremophiles, 28(2), 26. doi:10.1007/s00792-024-01341-7.

Crick, F., & Orgel, L. (1973). Directed Panspermia. Icarus, 19(3), 341–346. 10.1016/0019-1035(73)90110-3

Dávila-Ramos, S., Castelán-Sánchez, H., Martínez-Ávila, L., Sánchez-Carbente, M., Peralta, R., Hernández-Mendoza, A. D., . . . Batista-García, R. (2019). A Review on Viral Metagenomics in Extreme Environments. Front Microbiol., 10, 2403. doi:10.3389/fmicb.2019.02403.

Delwart, E. (2013). A Roadmap to the Human Virome. PLoS Pathog, 9(2), e1003146. doi:10.1371/journal.ppat.1003146

ENCODE Project Consortium. (s.f.). An integrated encyclopedia of DNA elements in the human genome. Nature, 489(7414), 57-74. doi:10.1038/nature11247.

Ferus M, K. A. (2015). Meteorite-catalyzed synthesis of nucleosides and other prebiotic compounds. Proc Natl Acad Sci U S A, 112(23), 7109–10. doi:10.1073/pnas.1507471112.

Flynn, R., & Zou, L. (2010). Oligonucleotide/oligosaccharide-binding fold proteins: a growing family of genome guardians. Crit Rev Biochem Mol Biol., 45(4), 266–75. doi:10.3109/10409238.2010.488216.

Forterre, P., & Prangishvili, D. (2009). The great billion-year war between ribosome- and capsid-encoding organisms (cells and viruses) as the major source of evolutionary novelties. Ann N Y Acad Sci., 1178, 65–77. doi:10.1111/j.1749-6632.2009.04993.x.

Gama-Castro, S., H, S., Santos-Zavaleta, A., Ledezma-Tejeida, D., Muñiz-Rascado, L., García-Sotelo, J., . . . Pérez-Rueda, E. (2016). RegulonDB version 9.0: high-level integration of gene regulation, coexpression, motif clustering and beyond. Nucleic Acids Res., 44(D1), D133–43. doi:10.1093/nar/gkv1156.

Gemayel, R., Cho, J., Boeynaems, S., & Verstrepen, K. (2012). Beyond junk-variable tandem repeats as facilitators of rapid evolution of regulatory and coding sequences.. Genes (Basel)., 3(3), 461–80. doi:10.3390/genes3030461.

Golden, M., Murrell, B., Martin, D., Pybus, O., & Hein, J. (2020). Evolutionary Analyses of Base-Pairing Interactions in DNA and RNA Secondary Structures. Mol Biol Evol., 37(2), 576–592. doi:10.1093/molbev/msz243.

Guglielmini, J., de la Cruz, F., & Rocha, E. (2013). Evolution of conjugation and type IV secretion systems. Mol Biol Evol., 30(2), 315–31. doi:10.1093/molbev/mss221.

Guo, J., Bolduc, B., Zayed, A. A., Varsani, A., Dominguez-Huerta, G., Delmont, T. O., . . . Roux, S. (2021). VirSorter2: a multi-classifier, expert-guided approach to detect diverse DNA and RNA viruses. Microbiome, 9(1), 37. doi:10.1186/s40168-020-00990-y

Hao, M., Qiao, J., & Qi, H. (2020). Current and Emerging Methods for the Synthesis of Single-Stranded DNA. Genes (Basel), 11(2), 116. doi:10.3390/genes11020116.

Hawking, S. (2001). The Universe in a Nutshell (1 ed.). United Kingdom: Bantam Spectra.

Hu, C., Chen, Z., Wang, G., Yang, H., & Ding, J. (2024). Biochemical and structural characterization of the DNA-binding properties of human TRIP4 ASCH domain reveals insights into its functional role. Structure, 32(8), 1208–1221.e4. doi:10.1016/j.str.2024.05.012.

Irony-Tur Sinai, M., Salamon, A., Stanleigh, N., Goldberg, T., Weiss, A., Wang, Y., & Kerem, B. (2019). AT-dinucleotide rich sequences drive fragile site formation. Nucleic Acids Res., 47(18), 9685–9695. doi:10.1093/nar/gkz689.

Iyer, L., Koonin, E., & Aravind, L. (2002). Classification and evolutionary history of the single-strand annealing proteins, RecT, Redbeta, ERF and RAD52. BMC Genomics, 3, 8. doi:10.1186/1471-2164-3-8.

Jones, P., Binns, D., Chang, H., Fraser, M., Li, W., McAnulla, C., . . . Hunter, S. (2014). InterProScan 5: genome-scale protein function classification. Bioinformatics, 30(9), 1236–40. doi:10.1093/bioinformatics/btu031

Kazazian, H. J. (2004). Mobile elements: drivers of genome evolution. Science, 303(5664), 1626–32. doi:10.1126/science.1089670.

Kazlauskas, D., Dayaram, A., Kraberger, S., Goldstien, S., Varsani, A., & Krupovic, M. (2017). Evolutionary history of ssDNA bacilladnaviruses features horizontal acquisition of the capsid gene from ssRNA nodaviruses. Virology, 504, 114–121. doi:10.1016/j.virol.2017.02.001.

Kazlauskas, D., Varsani, A., & Krupovic, M. (2018). Pervasive Chimerism in the Replication-Associated Proteins of Uncultured Single-Stranded DNA Viruses. Viruses, 10(4), 187. doi:10.3390/v10040187.

Khan, S. (2005). Plasmid rolling-circle replication: highlights of two decades of research. Plasmid, 53(2), 126–36. doi:10.1016/j.plasmid.2004.12.008

Kim, E., & Blair, D. (2015). Function of the Histone-Like Protein H-NS in Motility of Escherichia coli: Multiple Regulatory Roles Rather than Direct Action at the Flagellar Motor. J Bacteriol., 197(19), 3110–20. doi:10.1128/JB.00309-15.

Kim, S., & Lee, T. (2021). Conformational Dynamics of Poly(T) Single-Stranded DNA at the Single-Molecule Level. J Phys Chem Lett., 12(19), 4576–4584. doi:10.1021/acs.jpclett.1c00962.

Koonin, E., Dolja, V., & Krupovic, M. (s.f.). Origins and evolution of viruses of eukaryotes: The ultimate modularity. Virology, 479-480, 2-25. doi:10.1016/j.virol.2015.02.039. Epub 2015 Mar 12.

Längst, G., & Manelyte, L. (2015). Chromatin Remodelers: From Function to Dysfunction. Genes (Basel), 6(2), 299–324. doi:10.3390/genes6020299.

LaTourrette, K., & Garcia-Ruiz, H. (2022). Determinants of Virus Variation, Evolution, and Host Adaptation. Pathogens, 11(9), 1039. doi:10.3390/pathogens11091039. PMID: 36145471

Li, D., Liu, C.-M., Luo, R., Sadakane, K., & Lam, T.-W. (2015). MEGAHIT: an ultra-fast single-node solution for large andcomplex metagenomics assembly via succinct de Bruijn graph. Bioinformatics,, 31(10), 1674–1676. doi:10.1093/bioinformatics/btv033COREMetadata, citation and similar papers at core.ac.ukProvided by HKU Scholars Hub

Lin, M., Malik, F., & Guo, J. (2021). A comparative study of protein-ssDNA interactions. NAR Genom Bioinform., 3(1), lqab006. doi:10.1093/nargab/lqab006.

Lorenz, R., Bernhart, S., Höner, Z. S., Tafer, H., Flamm, C., Stadler, P., & Hofacker, I. (2011). ViennaRNA Package 2.0. Algorithms Mol Biol., 6(26), 1–14. doi:10.1186/1748-7188-6-26.

Löytynoja, A., & Goldman, N. (2008). Phylogeny-aware gap placement prevents errors in sequence alignment and evolutionary analysis. Science, 320(5883), 1632–5. doi:10.1126/science.1158395.

Malathi, V., & Renuka Devi, P. (2019). sDNA viruses: key players in global virome. Virusdisease, 30(1), 3–12. doi:10.1007/s13337-019-00519-4.

Marceau, A. (2012). Functions of single-strand DNA-binding proteins in DNA replication, recombination, and repair. Methods Mol Biol., 1-21. doi:10.1007/978-1-62703-032-8_1.

Martins, Z., Botta, O., Fogel, M., Sephton, M., Glavin, D., Watson, J., . . . Ehrenfreund, P. (2008). Extraterrestrial nucleobases in the Murchison meteorite. Earth and Planetary Science Letters,, 270((1-2)), 130-136. 10.1016/j.epsl.2008.03.026

McKay, C. (2020). What Is Life-and When Do We Search for It on Other Worlds. Astrobiology, 20(2), 163–166. doi:10.1089/ast.2019.2136.

Muhire, B., Golden, M., Murrell, B., Lefeuvre, P., Lett, J., Gray, A., . . . Martin, D. (2014). Evidence of pervasive biologically functional secondary structures within the genomes of eukaryotic single-stranded DNA viruses. J Virol., 88(4), 1972–89. doi:10.1128/JVI.03031-13.

Nüesch, M., Pietrek, L., Holmstrom, E., Nettels, D., von Roten, V., Kronenberg-Tenga, R., . . . Schuler, B. (2024). Nanosecond chain dynamics of single-stranded nucleic acids. Nat Commun., 15(1), 6010. doi:10.1038/s41467-024-50092-8

Oba, Y., Takano, Y., Furukawa, Y., Koga, T., Glavin, D., Dworkin, J., & Naraoka, H. (2022). Identifying the wide diversity of extraterrestrial purine and pyrimidine nucleobases in carbonaceous meteorites. Nat Commun., 13(1), 2008. doi:10.1038/s41467-022-29612-x.

Ondov, B., Bergman, N., & Phillippy, A. (2011). Interactive metagenomic visualization in a Web browser. BMC Bioinformatic, 12, 385. 10.1186/1471-2105-12-385

Padeken, J., Zeller, P., & Gasser, S. (2015). Repeat DNA in genome organization and stability. Curr Opin Genet Dev., 12, 9. doi:10.1016/j.gde.2015.03.009.

Pal, A., & Levy, Y. (2019). Structure, stability and specificity of the binding of ssDNA and ssRNA with proteins. PLoS Comput Biol., 15(4), e1006768. doi:10.1371/journal.pcbi.1006768.

Peng, Y., Leung, H., Yiu, S., & Chin, F. (2012). IDBA-UD: a de novo assembler for single-cell and metagenomic sequencing data with highly uneven depth. 28(11), 1420–8.. doi:10.1093/bioinformatics/bts174.

Pizzarello, S., & Shock, E. (2010). The organic composition of carbonaceous meteorites: the evolutionary story ahead of biochemistry. Cold Spring Harb Perspect Biol., 2(3), a002105. doi:10.1101/cshperspect.a002105.

Podgornaya, O., Vasilyeva, I., & Bespalov, V. (2016). Heterochromatic Tandem Repeats in the Extracellular DNA. Adv Exp Med Biol., 924, 85–89. doi:10.1007/978-3-319-42044-8_16.

Rauluseviciute, I., Riudavets-Puig, R., Blanc-Mathieu, R., Castro-Mondragon, J., Ferenc, K., Kumar, V., . . . Mathelier, A. (2024). JASPAR 2024: 20th anniversary of the open-access database of transcription factor binding profiles. Nucleic Acids Res., 52(D1), D174–D182. doi:10.1093/nar/gkad1059.

Roux, S., Hallam, S., Woyke, T., & Sullivan, M. (Jul de 2015). Viral dark matter and virus-host interactions resolved from publicly available microbial genomes. Elife, 22(4), e08490. doi:10.7554/eLife.08490. PMID: 26200428; PMCID: PMC4533152.

Seemann, T. (2014). Prokka: rapid prokaryotic genome annotation. 30(14), 2068–9. doi:10.1093/bioinformatics/btu153

Sephton, M. (2002). Organic compounds in carbonaceous meteorites. Nat Prod Rep, 19(3), 292–311. doi:10.1039/b103775g.

Simón, D., Cristina, J., & Musto, H. (2021). Nucleotide Composition and Codon Usage Across Viruses and Their Respective Hosts. Front Microbiol, 28(12), 646300. 10.3389/fmicb.2021.646300

Solé, R., Kempes, C., Corominas-Murtra, B., De Domenico, M., Kolchinsky, A., Lachmann, M., . . . Wolpert, D. (2024). Fundamental constraints to the logic of living systems. 14(5), 20240010. doi:10.1098/rsfs.2024.0010.

Stedman, K. (2013). Mechanisms for RNA Capture by ssDNA Viruses: Grand Theft RNA. J Mol Evol., 76(6), 359–64. doi:10.1007/s00239-013-9569-9.

Suttle, C. (2007). Marine viruses--major players in the global ecosystem. Nat Rev Microbiol, 5(10), 801–12. doi:10.1038/nrmicro1750. PMID: 17853907.

Tokuda, M., & Shintani, M. (2024). Microbial evolution through horizontal gene transfer by mobile genetic elements. Microb Biotechnol., 17(1), e14408. doi:10.1111/1751-7915.14408.

Tong, H., Schliekelman, P., & Mrázek, J. (2017). Unsupervised statistical discovery of spaced motifs in prokaryotic genomes. BMC Genomics, 18(1), 27. doi:10.1186/s12864-016-3400-0.

Torralba, B., Blanc, S., & Michalakis, Y. (2024). Reassortments in single-stranded DNA multipartite viruses: Confronting expectations based on molecular constraints with field observations. Virus Evol., 10(1), veae010. doi:10.1093/ve/veae010.

Tóth, G., Gáspári, Z., & Jurka, J. (2000). Microsatellites in different eukaryotic genomes: survey and analysis. Genome Research, 10(7), 967–981. doi:10.1101/gr.10.7.967

Vassetzky, N., & Kramerov, D. (2013). SINEBase: a database and tool for SINE analysis. Nucleic Acids Res., 41(Database issue), D83-9. doi:10.1093/nar/gks1263.

Venkataraman, S., & Selvarajan, R. (2019). Recent advances in understanding the replication initiator protein of the ssDNA plant viruses of the family Nanoviridae. Virusdisease, 30(1), 22–31. doi:10.1007/s13337-019-00514-9.

Wegrzyn, K., Fuentes-Perez, M., Bury, K., Rajewska, M., Moreno-Herrero, F., & Konieczny, I. (2014). Sequence-specific interactions of Rep proteins with ssDNA in the AT-rich region of the plasmid replication origin. Nucleic Acids Res., 42(12), 7807–18. doi:10.1093/nar/gku453.

Wegrzyn, K., Oliwa, M., Nowacka, M., Zabrocka, E., Bury, K., Purzycki, P., . . . Konieczny, I. (2023). Rep protein accommodates together dsDNA and ssDNA which enables a loop-back mechanism to plasmid DNA replication initiation. Nucleic Acids Res., 51(19), 10551–10567. doi:10.1093/nar/gkad740.

Willaert, R., & Kasas, S. (2022). High-Speed Atomic Force Microscopy Visualization of Protein-DNA Interactions Using DNA Origami Frames. Methods Mol Biol., 2516, 157–167. doi:10.1007/978-1-0716-2413-5_10.

Witte, G., Urbanke, C., & Curth, U. (2003). DNA polymerase III chi subunit ties single-stranded DNA binding protein to the bacterial replication machinery. Nucleic Acids Res., 31(15), 4434–40. doi:10.1093/nar/gkg498.

Wood, D. E., Lu, J., & Langmead, B. (2019). Improved metagenomic analysis with Kraken 2. Genome Bio, 20(1), 257. doi:10.1186/s13059-019-1891-0

Yates, L., Aramayo, R., Pokhrel, N., Caldwell, C., Kaplan, J., Perera, R., . . . Zhang, X. (2018). A structural and dynamic model for the assembly of Replication Protein A on single-stranded DNA. Nat Commun., 9(1), 5447. doi:10.1038/s41467-018-07883-7.

Yuan, H., Liu, X., Liu, X., Zhao, L., Mao, S., & Huang, Y. (2024). The evolutionary dynamics of genome sizes and repetitive elements in Ensifera (Insecta: Orthoptera).. BMC Genomics, 25(1), 1041. doi:10.1186/s12864-024-10949-0.

Zawilak-Pawlik, A., Kois, A., Majka, J., Jakimowicz, D., Smulczyk-Krawczyszyn, A., Messer, W., & Zakrzewska-Czerwińska, J. (2005). Architecture of bacterial replication initiation complexes: orisomes from four unrelated bacteria. Biochem J, 389(Pt 2), 471–81. doi:10.1042/BJ20050143.

Zhang, Y., Saini, N., Sheng, Z., & Lobachev, K. (2013). Genome-wide screen reveals replication pathway for quasi-palindrome fragility dependent on homologous recombination. 9(12), e1003979. doi:10.1371/journal.pgen.1003979.

Zheng, Y., Li, X., & Hu, H. (2015). Comprehensive discovery of DNA motifs in 349 human cells and tissues reveals new features of motifs. Nucleic Acids Res., 43(1), 74–83. doi:10.1093/nar/gku1261.

