## Supplementary material for "Discovery of single-stranded DNA in meteorite-derived cultures: evidence of novel genetic elements": This document includes the predicted secondary structures of ssDNA sequences analyzed in this study. These structures were generated using RNAfold

**Supplementary Material for**
"Discovery of Single-Stranded DNA in Meteorite-Derived Cultures: Evidence of Novel Genetic Elements"

This document includes the predicted secondary structures of ssDNA sequences analyzed in this study. These structures were generated using ViennaRNA (RNAfold) and provide insights into potential functional and structural roles.

**Contents:**

1. **MT_TOTAL Sequences:**
   Predicted secondary structures for the 15 ssDNA sequences in the MT_TOTAL dataset.
2. **MT_PURE Sequences:**
   Predicted secondary structures for the 13 ssDNA sequences in the MT_PURE dataset.

Each structure is labeled with its corresponding sequence identifier (e.g., MT_TOTAL_SEQ1, MT_PURE_SEQ1) for easy reference. These results complement the findings in the main text and support the hypothesis of structural stability and autoreplication potential under extreme conditions.

### Section 1:

### Secondary Structures of MT_TOTAL Sequences.

This section includes the predicted secondary structures of the 15 ssDNA sequences in the MT_TOTAL dataset. Each structure is labeled with its corresponding sequence identifier for reference.

### RNAfold WebServer MT_TOTAL_SEQ1

1 Enter Input Parameters 2 View Results

[Home|New job|Help]

##### Results for minimum free energy prediction

The optimal secondary structure in dot-bracket notation with a minimum free energy of **-49.10** kcal/mol is given below.

[color by base-pairing probability | color by positional entropy | no coloring]

```
1       CUUGGUGUUAAAUAUCUAAUAAUUAAUACUCCUGAUCCUCCCACUCCAUUAUUUGUACCUGCAAAAGCUAUACGACCAUUUCCUCCUCCACCACCACCAAAACCAUUUGUGGCAUUUGUACCUUGUAAACCUGCACCUUCGACAGCUCCUUGUCCUGAAU
160     UAUCACAACCUCCAAGUGAUCUAGUACUUGUUGUAUUAGCUUCGUGUGUACCACCUCCUCCCCCACCUGCAUAACAUUUUGUUGUACCGCUUAUACUAAAAUUAGUUCCUAAUCCGCCUCUUCCUCCCUUUGCAAAAUCAUUAGUACCUGACUGAAAUCC
320     UGUUGUAUUAUGUCCCCCUCCUCCAUUAGAACCAUAACCACUUACUCCGUUAUAUGUAGCUCCAUUAUUUCCCUGACCUGCAGUUCCACUUCCAAAGUUUGUAUUUUGUUGUCCAGUACCACCACCA

1       ...((((((((.(((...))).))))))))......................((((..((((((((..((((((................((((.(((....)))))))..........((((....))))).....(((((....)))))......
160     .(((((.........)))))(((((((.....)))))))).....((((((.................))))))........))))))..))))..((((....)))).......................(((((.....(((((.....)))))....
320     ..)))))........................(((((.........))))).(((((.((............)).)))))..........((((((....)))))))))...))))........
```

You can download the minimum free energy (MFE) structure in [Vienna Format| Ct Format]. You can get thermodynamic details on this structure by submitting to our RNAeval web server.

##### Results for thermodynamic ensemble prediction

The free energy of the thermodynamic ensemble is **-60.16** kcal/mol.

The frequency of the MFE structure in the ensemble is **0.00** %.

The ensemble diversity is **112.91.**

You may look at the **dot plot** containing the base pair probabilities [EPS|PDF|IMAGE CONVERTER].

The centroid secondary structure in dot-bracket notation with a minimum free energy of **-25.50** kcal/mol is given below.

[color by base-pairing probability | color by positional entropy | no coloring]

```
1       CUUGGUGUUAAAUAUCUAAUAAUUAAUACUCCUGAUCCUCCCACUCCAUUAUUUGUACCUGCAAAAGCUAUACGACCAUUUCCUCCUCCACCACCACCAAAACCAUUUGUGGCAUUUGUACCUUGUAAACCUGCACCUUCGACAGCUCCUUGUCCUGAAU
160     UAUCACAACCUCCAAGUGAUCUAGUACUUGUUGUAUUAGCUUCGUGUGUACCACCUCCUCCCCCACCUGCAUAACAUUUUGUUGUACCGCUUAUACUAAAAUUAGUUCCUAAUCCGCCUCUUCCUCCCUUUGCAAAAUCAUUAGUACCUGACUGAAAUCC
320     UGUUGUAUUAUGUCCCCCUCCUCCAUUAGAACCAUAACCACUUACUCCGUUAUAUGUAGCUCCAUUAUUUCCCUGACCUGCAGUUCCACUUCCAAAGUUUGUAUUUUGUUGUCCAGUACCACCACCA

1       ...((((............................................................((((.((((((................((((.((......))))))..........((((....))))).....((((......))))......
160     .(((((.........)))))..........))))))..))))....................................................((((....)))).....................................................
320     ...............................(((((.........))))).(((((.((............)).)))))...........(((........)))............))))....
```

You can download the minimum free energy (MFE) structure in [Vienna Format| Ct Format]. You can get thermodynamic details on this structure by submitting to our RNAeval web server.

##### Graphical output

You may look at the interactive drawing of the MFE structure below. If you do not see the interactive drawing and you are using Internet Explorer, please install the Adobe SVG plugin. **A note on base-pairing probabilities:** The structure below is colored by base-pairing probabilities. For unpaired regions the color denotes the probability of being unpaired.

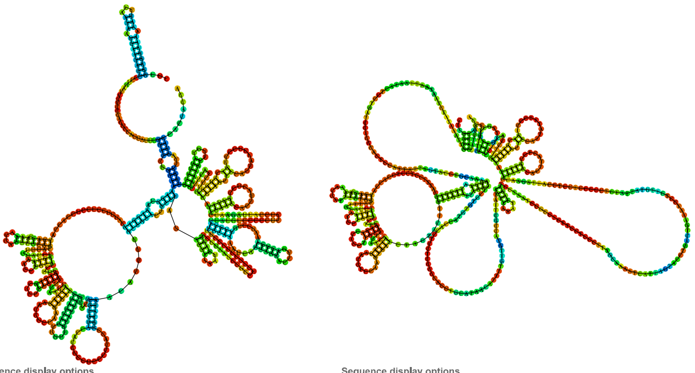

Sequence display options
- (•) Plain Sequence
- ( ) No Sequence

**MFE secondary structure**

Other display options
- (•) Base-pair probabilities
- ( ) Positional entropy
- ( ) None

0 1

Sequence display options
- (•) Plain Sequence
- ( ) No Sequence

**Centroid secondary structure**

Other display options
- (•) Base-pair probabilities
- ( ) Positional entropy
- ( ) None

0 1

| Image description | Download options |
|---|---|
| MFE plain structure drawing | [ EPS \| PDF \| IMAGE CONVERTER \| VIEW IN FORNA ] |
| Centroid plain structure drawing | [ EPS \| PDF \| IMAGE CONVERTER \| VIEW IN FORNA ] |
| MFE structure drawing encoding base-pair probabilities | [ EPS \| PDF \| IMAGE CONVERTER \| VIEW IN FORNA ] |
| Centroid structure drawing encoding base-pair probabilities | [ EPS \| PDF \| IMAGE CONVERTER \| VIEW IN FORNA ] |
| MFE structure drawing encoding positional entropy | [ EPS \| PDF \| IMAGE CONVERTER \| VIEW IN FORNA ] |
| Centroid structure drawing encoding positional entropy | [ EPS \| PDF \| IMAGE CONVERTER \| VIEW IN FORNA ] |

Here you find a mountain plot representation of the MFE structure, the thermodynamic ensemble of RNA structures, and the centroid structure. Additionally we present the positional entropy for each position. Download as [EPS|PDF|IMAGE CONVERTER].

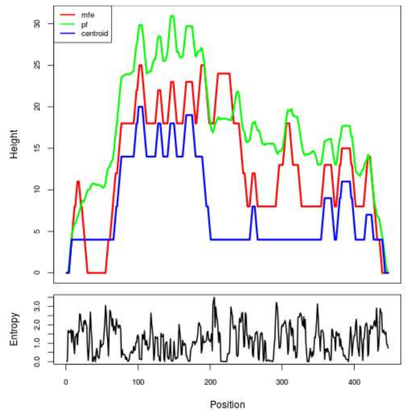


Results have been computed using RNAfold 2.6.3. An equivalent command line call would have been

```
RNAfold -p -d2 --noLP < sequence1.fa > sequence1.out
```

**RNA parameters are described in**

Mathews DH, Disney MD, Childs JL, Schroeder SJ, Zuker M, Turner DH. (2004) Incorporating chemical modification constraints into a dynamic programming algorithm for prediction of RNA secondary structure. *Proc Natl Acad Sci U S A* 101(19):7287-92.

**If you find these results helpful for your work you may want to cite:**


OPEN ACCESS OXFORD JOURNALS Gruber AR, Lorenz R, Bernhart SH, Neuböck R, Hofacker IL.
**The Vienna RNA Websuite.** Nucleic Acids Research, Volume 36, Issue suppl_2, 1 July 2008, Pages W70-W74, DOI: 10.1093/nar/gkn188

Lorenz, R. and Bernhart, S.H. and Höner zu Siederdissen, C. and Tafer, H. and Flamm, C. and Stadler, P.F. and Hofacker, I.L. "ViennaRNA Package 2.0", Algorithms for Molecular Biology, 6:1 page(s): 26, 2011

### MT_TOTAL_SEQ2

[Home|New job|Help]

##### Results for minimum free energy prediction

The optimal secondary structure in dot-bracket notation with a minimum free energy of **-44.20** kcal/mol is given below.
[color by base-pairing probability | color by positional entropy | no coloring]

```
1     UGUAGUCGAUCCAAGUAUCGGGGGAAUAGACGCAACCUAAACAUUUGAAUUGAAUCUUACUCAUUGCUUUGCUCUUCAAGUAUCUUAUCUCUAUUGAUUUUAGCCCAAACACAGACCGUAUAUAAAUCCCACUGAGCAGCCGCGAAACACUCGCUGGAAU
160   GGCUUACCUUAUUUCCUUCGAUCAAGUCCAUCCCGCAUGUCUCGCAAUCAAUAUCAGUAAUAGUAUCGGUAUCACUCUGAUCAGCACAUACUCUGGUUUUCUUCGACACA

1     (((.(((((.......((((((((..(((..(((..(((....)))..)))..)))...)).......((((..((((((.....(((......)))...............((((.((((.......(((((......((((.(((...((.(((.(
160   (((((...................))).)))))).)).))).)))).......)))))....))))))))...))).)))..))))....)))))))).....))))))))
```

You can download the minimum free energy (MFE) structure in [Vienna Format| Ct Format]. You can get thermodynamic details on this structure by submitting to our RNAeval web server.

##### Results for thermodynamic ensemble prediction

The free energy of the thermodynamic ensemble is **-50.12** kcal/mol.
The frequency of the MFE structure in the ensemble is **0.01** %.
The ensemble diversity is **60.31** .
You may look at the **dot plot** containing the base pair probabilities [EPS|PDF|IMAGE CONVERTER].

The centroid secondary structure in dot-bracket notation with a minimum free energy of **-27.27** kcal/mol is given below.
[color by base-pairing probability | color by positional entropy | no coloring]

```
1     UGUAGUCGAUCCAAGUAUCGGGGGAAUAGACGCAACCUAAACAUUUGAAUUGAAUCUUACUCAUUGCUUUGCUCUUCAAGUAUCUUAUCUCUAUUGAUUUUAGCCCAAACACAGACCGUAUAUAAAUCCCACUGAGCAGCCGCGAAACACUCGCUGGAAU
160   GGCUUACCUUAUUUCCUUCGAUCAAGUCCAUCCCGCAUGUCUCGCAAUCAAUAUCAGUAAUAGUAUCGGUAUCACUCUGAUCAGCACAUACUCUGGUUUUCUUCGACACA

1     .((.(((((........................................................................................................((((.((((.......(((((......((((.(((...((.(((.(
160   (((((...................))).)))))).)).))).)))).......)))))....))))))))...............................))))))).
```

You can download the minimum free energy (MFE) structure in [Vienna Format| Ct Format]. You can get thermodynamic details on this structure by submitting to our RNAeval web server.

##### Graphical output

You may look at the interactive drawing of the MFE structure below. If you do not see the interactive drawing and you are using Internet Explorer, please install the Adobe SVG plugin. **A note on base-pairing probabilities:** The structure below is colored by base-pairing probabilities. For unpaired regions the color denotes the probability of being unpaired.

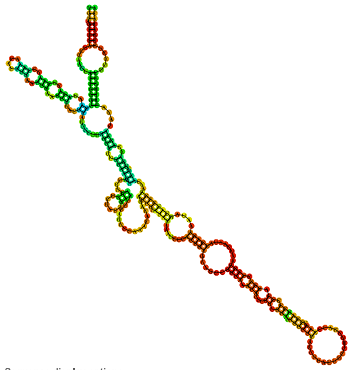

Sequence display options
- ◉ Plain Sequence
- ○ No Sequence

MFE secondary structure

Other display options
- ◉ Base-pair probabilities
- ○ Positional entropy
- ○ None

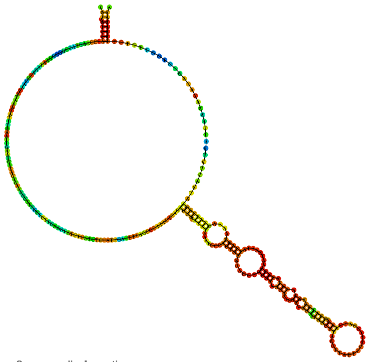

Sequence display options
- ◉ Plain Sequence
- ○ No Sequence

Centroid secondary structure

Other display options
- ◉ Base-pair probabilities
- ○ Positional entropy
- ○ None

| Image description | Download options |
|---|---|
| MFE plain structure drawing | [ EPS \| PDF \| IMAGE CONVERTER \| VIEW IN FORNA ] |
| Centroid plain structure drawing | [ EPS \| PDF \| IMAGE CONVERTER \| VIEW IN FORNA ] |
| MFE structure drawing encoding base-pair probabilities | [ EPS \| PDF \| IMAGE CONVERTER \| VIEW IN FORNA ] |
| Centroid structure drawing encoding base-pair probabilities | [ EPS \| PDF \| IMAGE CONVERTER \| VIEW IN FORNA ] |
| MFE structure drawing encoding positional entropy | [ EPS \| PDF \| IMAGE CONVERTER \| VIEW IN FORNA ] |
| Centroid structure drawing encoding positional entropy | [ EPS \| PDF \| IMAGE CONVERTER \| VIEW IN FORNA ] |

Here you find a mountain plot representation of the MFE structure, the thermodynamic ensemble of RNA structures, and the centroid structure. Additionally we present the positional entropy for each position. Download as [EPS|PDF|IMAGE CONVERTER].

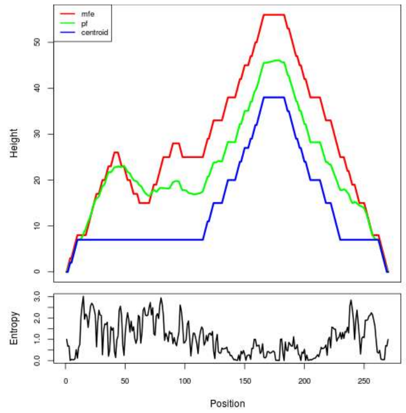


Results have been computed using RNAfold 2.6.3. An equivalent command line call would have been

```
RNAfold -p -d2 --noLP < sequence1.fa > sequence1.out
```

---


**RNA parameters are described in**
Mathews DH, Disney MD, Childs JL, Schroeder SJ, Zuker M, Turner DH. (2004) Incorporating chemical modification constraints into a dynamic programming algorithm for prediction of RNA secondary structure. *Proc Natl Acad Sci U S A* 101(19):7287-92.

**If you find these results helpful for your work you may want to cite:**

OPEN ACCESS OXFORD JOURNALS Gruber AR, Lorenz R, Bernhart SH, Neuböck R, Hofacker IL.
**The Vienna RNA Websuite.** Nucleic Acids Research, Volume 36, Issue suppl_2, 1 July 2008, Pages W70-W74, DOI: 10.1093/nar/gkn188

Lorenz, R. and Bernhart, S.H. and Höner zu Siederdissen, C. and Tafer, H. and Flamm, C. and Stadler, P.F. and Hofacker, I.L. "ViennaRNA Package 2.0", Algorithms for Molecular Biology, 6:1 page(s): 26, 2011

### RNAfold WebServer MT_TOTAL_SEQ3



##### Results for minimum free energy prediction

The optimal secondary structure in dot-bracket notation with a minimum free energy of **-18.75** kcal/mol is given below.
[color by base-pairing probability | color by positional entropy | no coloring]

```
1      UAUAGAUUAAAAAUCACAUAAAAAUAAAAAAUCAAAUAAAAAUAAAAGAACGAAAAAAAAACGAAGAAACAAAGUGACAAAGUAAAAAAGUUCUUAAAAAAUUUGAUUAUAAAAAUGUCGACGACAAACGUGAUAAAAUCUAAAAAAUAAUACAAAAAUUG
160    ACAAAGUCAAAAAAGCCGAAAGUAAAAAGACACAAACACAUAUACACUAUGUAAUACAUUUAACACACACUCAAACACAACAAAUUUCUGCCGUUCCUUACCCC

1      ..((((((....(((((............((((((((........((((((....................................))))))....)))))))).......((((...))))...)))))..))))))...............(((
160    ((...)))))....((.((((((......))......(((((.....)))))............................)))).))............
```

You can download the minimum free energy (MFE) structure in [Vienna Format| Ct Format]. You can get thermodynamic details on this structure by submitting to our RNAeval web server.

##### Results for thermodynamic ensemble prediction

The free energy of the thermodynamic ensemble is **-21.46** kcal/mol.
The frequency of the MFE structure in the ensemble is **1.23** %.
The ensemble diversity is **27.00** .
You may look at the **dot plot** containing the base pair probabilities [EPS|PDF|IMAGE CONVERTER].

The centroid secondary structure in dot-bracket notation with a minimum free energy of **-16.40** kcal/mol is given below.
[color by base-pairing probability | color by positional entropy | no coloring]

```
1      UAUAGAUUAAAAAUCACAUAAAAAUAAAAAAUCAAAUAAAAAUAAAAGAACGAAAAAAAAACGAAGAAACAAAGUGACAAAGUAAAAAAGUUCUUAAAAAAUUUGAUUAUAAAAAUGUCGACGACAAACGUGAUAAAAUCUAAAAAAUAAUACAAAAAUUG
160    ACAAAGUCAAAAAAGCCGAAAGUAAAAAGACACAAACACAUAUACACUAUGUAAUACAUUUAACACACACUCAAACACAACAAAUUUCUGCCGUUCCUUACCCC

1      ..((((((....(((((............((((((((........((((((....................................))))))....)))))))).......((((...))))...)))))..))))))...............(((
160    ((...)))))....((.(((................(((((.....)))))..............................))).))............
```

You can download the minimum free energy (MFE) structure in [Vienna Format| Ct Format]. You can get thermodynamic details on this structure by submitting to our RNAeval web server.

##### Graphical output

You may look at the interactive drawing of the MFE structure below. If you do not see the interactive drawing and you are using Internet Explorer, please install the Adobe SVG plugin. **A note on base-pairing probabilities:** The structure below is colored by base-pairing probabilities. For unpaired regions the color denotes the probability of being unpaired.

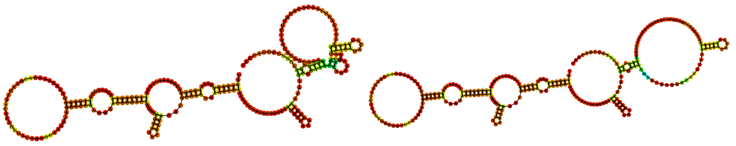

Sequence display options
- (•) Plain Sequence
- ( ) No Sequence

**MFE secondary structure**

Other display options
- (•) Base-pair probabilities
- ( ) Positional entropy
- ( ) None

0 1

Sequence display options
- (•) Plain Sequence
- ( ) No Sequence

**Centroid secondary structure**

Other display options
- (•) Base-pair probabilities
- ( ) Positional entropy
- ( ) None

0 1

| Image description | Download options |
|---|---|
| MFE plain structure drawing | [ EPS \| PDF \| IMAGE CONVERTER \| VIEW IN FORNA ] |
| Centroid plain structure drawing | [ EPS \| PDF \| IMAGE CONVERTER \| VIEW IN FORNA ] |
| MFE structure drawing encoding base-pair probabilities | [ EPS \| PDF \| IMAGE CONVERTER \| VIEW IN FORNA ] |
| Centroid structure drawing encoding base-pair probabilities | [ EPS \| PDF \| IMAGE CONVERTER \| VIEW IN FORNA ] |
| MFE structure drawing encoding positional entropy | [ EPS \| PDF \| IMAGE CONVERTER \| VIEW IN FORNA ] |
| Centroid structure drawing encoding positional entropy | [ EPS \| PDF \| IMAGE CONVERTER \| VIEW IN FORNA ] |

Here you find a mountain plot representation of the MFE structure, the thermodynamic ensemble of RNA structures, and the centroid structure. Additionally we present the positional entropy for each position. Download as [EPS|PDF|IMAGE CONVERTER].

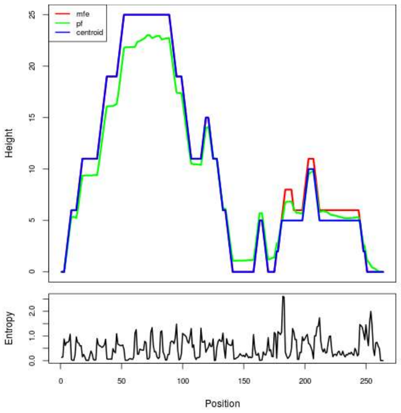


Results have been computed using RNAfold 2.6.3. An equivalent command line call would have been

```
RNAfold -p -d2 --noLP < sequence1.fa > sequence1.out
```

**RNA parameters are described in**
Mathews DH, Disney MD, Childs JL, Schroeder SJ, Zuker M, Turner DH. (2004) Incorporating chemical modification constraints into a dynamic programming algorithm for prediction of RNA secondary structure. *Proc Natl Acad Sci U S A* 101(19):7287-92.

**If you find these results helpful for your work you may want to cite:**

OPEN ACCESS OXFORD JOURNALS Gruber AR, Lorenz R, Bernhart SH, Neuböck R, Hofacker IL.
**The Vienna RNA Websuite.** Nucleic Acids Research, Volume 36, Issue suppl_2, 1 July 2008, Pages W70-W74, DOI: 10.1093/nar/gkn188

Lorenz, R. and Bernhart, S.H. and Höner zu Siederdissen, C. and Tafer, H. and Flamm, C. and Stadler, P.F. and Hofacker, I.L. "ViennaRNA Package 2.0", Algorithms for Molecular Biology, 6:1 page(s): 26, 2011

##### Results for minimum free energy prediction

The optimal secondary structure in dot-bracket notation with a minimum free energy of **-24.50** kcal/mol is given below.

[color by base-pairing probability | color by positional entropy | no coloring]

```
1      GGAUGGGCAUGGGGAUGGGGAUGGGUAUGGGAAUGGGUAUGGGAGUGGGCAUGGGUAUGGGGAUGGGCAUGGGUAUGGGUAUGGGUAUGGGUAUGGGGAUGGGUAUGGGAGUGGGGAUGGGUAUGGGGAUGGGUAUGGAAGAAACGCUAUUAAAAAGGAU
160    CCUAUUAUGAUUGGGAGGCAAAUGUUGAUUUGGGCUGGCCU

1      ....((((....((((.((.((..((.....(((....((((((.....(((...((((...(((..((((.......))))..)))...))))...)))........((((.......((((........)))).......))))...........)
160    )))))....)))).....))..)).)).))))......))))
```

You can download the minimum free energy (MFE) structure in [Vienna Format| Ct Format]. You can get thermodynamic details on this structure by submitting to our RNAeval web server.

##### Results for thermodynamic ensemble prediction

The free energy of the thermodynamic ensemble is **-29.20** kcal/mol.
The frequency of the MFE structure in the ensemble is **0.05** %.
The ensemble diversity is **64.94** .
You may look at the **dot plot** containing the base pair probabilities [EPS|PDF|IMAGE CONVERTER].

The centroid secondary structure in dot-bracket notation with a minimum free energy of **-4.10** kcal/mol is given below.
[color by base-pairing probability | color by positional entropy | no coloring]

```
1      GGAUGGGCAUGGGGAUGGGGAUGGGUAUGGGAAUGGGUAUGGGAGUGGGCAUGGGUAUGGGGAUGGGCAUGGGUAUGGGUAUGGGUAUGGGUAUGGGGAUGGGUAUGGGAGUGGGGAUGGGUAUGGGGAUGGGUAUGGAAGAAACGCUAUUAAAAAGGAU
160    CCUAUUAUGAUUGGGAGGCAAAUGUUGAUUUGGGCUGGCCU

1      ............................................................................................................((((.......((((........)))).......))))............
160    .........................................
```

You can download the minimum free energy (MFE) structure in [Vienna Format| Ct Format]. You can get thermodynamic details on this structure by submitting to our RNAeval web server.

##### Graphical output

You may look at the interactive drawing of the MFE structure below. If you do not see the interactive drawing and you are using Internet Explorer, please install the Adobe SVG plugin. **A note on base-pairing probabilities:** The structure below is colored by base-pairing probabilities. For unpaired regions the color denotes the probability of being unpaired.

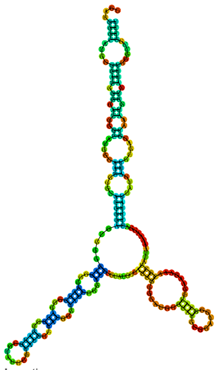

MFE secondary structure

Sequence display options
- (•) Plain Sequence
- ( ) No Sequence

Other display options
- (•) Base-pair probabilities
- ( ) Positional entropy
- ( ) None

0 1

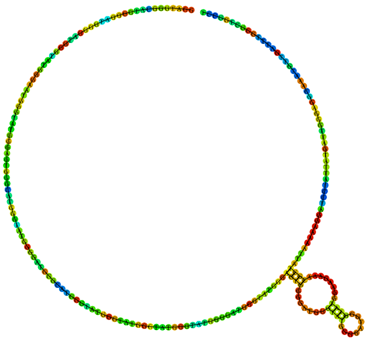

Centroid secondary structure

Sequence display options
- (•) Plain Sequence
- ( ) No Sequence

Other display options
- (•) Base-pair probabilities
- ( ) Positional entropy
- ( ) None

0 1

| Image description | Download options |
|---|---|
| MFE plain structure drawing | [ EPS \| PDF \| IMAGE CONVERTER \| VIEW IN FORNA ] |
| Centroid plain structure drawing | [ EPS \| PDF \| IMAGE CONVERTER \| VIEW IN FORNA ] |
| MFE structure drawing encoding base-pair probabilities | [ EPS \| PDF \| IMAGE CONVERTER \| VIEW IN FORNA ] |
| Centroid structure drawing encoding base-pair probabilities | [ EPS \| PDF \| IMAGE CONVERTER \| VIEW IN FORNA ] |
| MFE structure drawing encoding positional entropy | [ EPS \| PDF \| IMAGE CONVERTER \| VIEW IN FORNA ] |
| Centroid structure drawing encoding positional entropy | [ EPS \| PDF \| IMAGE CONVERTER \| VIEW IN FORNA ] |

Here you find a mountain plot representation of the MFE structure, the thermodynamic ensemble of RNA structures, and the centroid structure. Additionally we present the positional entropy for each position. Download as [EPS|PDF|IMAGE CONVERTER].

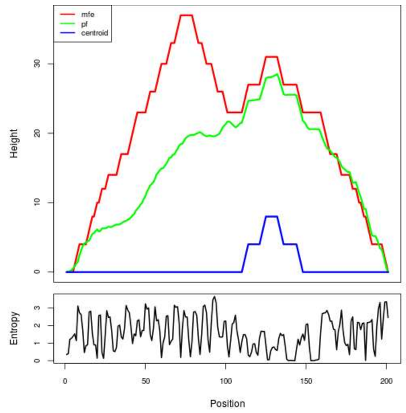


Results have been computed using RNAfold 2.6.3. An equivalent command line call would have been

```
RNAfold -p -d2 --noLP < sequence1.fa > sequence1.out
```

**RNA parameters are described in**

Mathews DH, Disney MD, Childs JL, Schroeder SJ, Zuker M, Turner DH. (2004) Incorporating chemical modification constraints into a dynamic programming algorithm for prediction of RNA secondary structure. *Proc Natl Acad Sci U S A* 101(19):7287-92.

**If you find these results helpful for your work you may want to cite:**

OPEN ACCESS OXFORD JOURNALS Gruber AR, Lorenz R, Bernhart SH, Neuböck R, Hofacker IL.
**The Vienna RNA Websuite.** Nucleic Acids Research, Volume 36, Issue suppl_2, 1 July 2008, Pages W70-W74, DOI: 10.1093/nar/gkn188

Lorenz, R. and Bernhart, S.H. and Höner zu Siederdissen, C. and Tafer, H. and Flamm, C. and Stadler, P.F. and Hofacker, I.L. "ViennaRNA Package 2.0", Algorithms for Molecular Biology, 6:1 page(s): 26, 2011

##### Results for minimum free energy prediction

The optimal secondary structure in dot-bracket notation with a minimum free energy of **-25.40** kcal/mol is given below.

[color by base-pairing probability | color by positional entropy | no coloring]

```
1      GGAUGGGCAUGGGGAUGGGGAUGGGUAUGGGAAUGGGUAUGGGAGUGGGCAUGGGUAUGGGGAUGGGCAUGGGUAUGGGCAUGGGUAUGGGUAUGGGGAUGGGUAUGGGGAUGGGUAUGGGGAUGGGGAUGGGUAUGGAAGAAACGCUAUUAAAAAGGAU
160    CCUAUUAUGAUUGGGAGGCAAAUGUUGAUUUGGGCUGGCCU

1      ....((((....((((.((.((..((.....(((....((((((.....(((...((((...(((..((((.......))))..)))...))))...))).((((.......))))........((((((.........)).)))))........)
160    )))))....)))).....))..)).)).))))......))))
```

You can download the minimum free energy (MFE) structure in [Vienna Format| Ct Format]. You can get thermodynamic details on this structure by submitting to our RNAeval web server.

##### Results for thermodynamic ensemble prediction

The free energy of the thermodynamic ensemble is **-29.67** kcal/mol.
The frequency of the MFE structure in the ensemble is **0.10** %.
The ensemble diversity is **72.22** .
You may look at the **dot plot** containing the base pair probabilities [EPS|PDF|IMAGE CONVERTER].

The centroid secondary structure in dot-bracket notation with a minimum free energy of **-1.40** kcal/mol is given below.
[color by base-pairing probability | color by positional entropy | no coloring]

```
1      GGAUGGGCAUGGGGAUGGGGAUGGGUAUGGGAAUGGGUAUGGGAGUGGGCAUGGGUAUGGGGAUGGGCAUGGGUAUGGGCAUGGGUAUGGGUAUGGGGAUGGGUAUGGGGAUGGGUAUGGGGAUGGGGAUGGGUAUGGAAGAAACGCUAUUAAAAAGGAU
160    CCUAUUAUGAUUGGGAGGCAAAUGUUGAUUUGGGCUGGCCU

1      ..............................................................................................................................(((((..............)))))........
160    .........................................
```

You can download the minimum free energy (MFE) structure in [Vienna Format| Ct Format]. You can get thermodynamic details on this structure by submitting to our RNAeval web server.

##### Graphical output

You may look at the interactive drawing of the MFE structure below. If you do not see the interactive drawing and you are using Internet Explorer, please install the Adobe SVG plugin. **A note on base-pairing probabilities:** The structure below is colored by base-pairing probabilities. For unpaired regions the color denotes the probability of being unpaired.

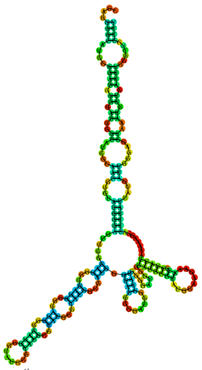

Sequence display options
- Plain Sequence
- No Sequence

**MFE secondary structure**

Other display options
- Base-pair probabilities
- Positional entropy
- None

0 1

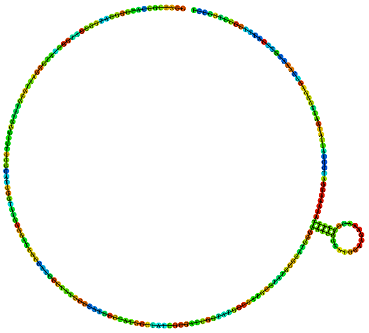

Sequence display options
- Plain Sequence
- No Sequence

**Centroid secondary structure**

Other display options
- Base-pair probabilities
- Positional entropy
- None

0 1

| **Image description** | **Download options** |
|---|---|
| MFE plain structure drawing | [ EPS \| PDF \| IMAGE CONVERTER \| VIEW IN FORNA ] |
| Centroid plain structure drawing | [ EPS \| PDF \| IMAGE CONVERTER \| VIEW IN FORNA ] |
| MFE structure drawing encoding base-pair probabilities | [ EPS \| PDF \| IMAGE CONVERTER \| VIEW IN FORNA ] |
| Centroid structure drawing encoding base-pair probabilities | [ EPS \| PDF \| IMAGE CONVERTER \| VIEW IN FORNA ] |
| MFE structure drawing encoding positional entropy | [ EPS \| PDF \| IMAGE CONVERTER \| VIEW IN FORNA ] |
| Centroid structure drawing encoding positional entropy | [ EPS \| PDF \| IMAGE CONVERTER \| VIEW IN FORNA ] |

Here you find a mountain plot representation of the MFE structure, the thermodynamic ensemble of RNA structures, and the centroid structure. Additionally we present the positional entropy for each position. Download as [EPS|PDF|IMAGE CONVERTER].

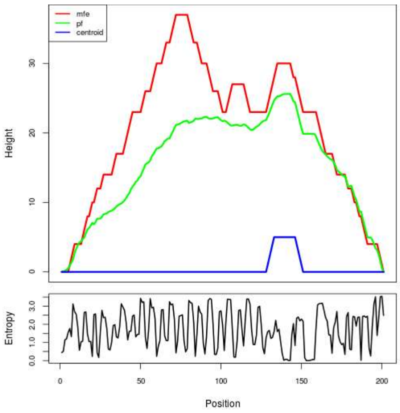


Results have been computed using RNAfold 2.6.3. An equivalent command line call would have been

```
RNAfold -p -d2 --noLP < sequence1.fa > sequence1.out
```

**RNA parameters are described in**

Mathews DH, Disney MD, Childs JL, Schroeder SJ, Zuker M, Turner DH. (2004) Incorporating chemical modification constraints into a dynamic programming algorithm for prediction of RNA secondary structure. *Proc Natl Acad Sci U S A* 101(19):7287-92.

**If you find these results helpful for your work you may want to cite:**

OPEN ACCESS OXFORD JOURNALS Gruber AR, Lorenz R, Bernhart SH, Neuböck R, Hofacker IL.
**The Vienna RNA Websuite.** Nucleic Acids Research, Volume 36, Issue suppl_2, 1 July 2008, Pages W70-W74, DOI: 10.1093/nar/gkn188

Lorenz, R. and Bernhart, S.H. and Höner zu Siederdissen, C. and Tafer, H. and Flamm, C. and Stadler, P.F. and Hofacker, I.L. "ViennaRNA Package 2.0", Algorithms for Molecular Biology, 6:1 page(s): 26, 2011

### RNAfold WebServer MT_TOTAL_SEQ6



##### Results for minimum free energy prediction

The optimal secondary structure in dot-bracket notation with a minimum free energy of **-55.80** kcal/mol is given below.
[color by base-pairing probability | color by positional entropy | no coloring]

```
1     GGCAGGGGGAGGUCUUGGGGCACUGGAAAGGGGCUGAGAGGUAGCUGGAGGGAGCAGAAGGGUCCAUAGGGGCAUGGGUAAGUUGGUUGAGGUCGAGGGAGUGUGGAUACAGGCACUUGGGAGGGGCAGACAGGUCAGGGAGGGUGUGGAGGGUGGAUGA
160   GGGGGCAUUGCAGGGCUUGAGAGGCACCUGGAGGGUGCCCUGGAGCAUUGGGAAGGUC

1     ..(((((.....)))))..(((((......((.(((.......(((......)))......((((....))))...........(((....((.(((..((.((....)).)).))).))...)))...))).)).......)))))............
160   ...(((.((.(((.((((.((.(((((((...)))))))))).)))).))).))..)))
```

You can download the minimum free energy (MFE) structure in [Vienna Format| Ct Format]. You can get thermodynamic details on this structure by submitting to our RNAeval web server.

##### Results for thermodynamic ensemble prediction

The free energy of the thermodynamic ensemble is **-59.81** kcal/mol.
The frequency of the MFE structure in the ensemble is **0.15** %.
The ensemble diversity is **67.26** .
You may look at the **dot plot** containing the base pair probabilities [EPS|PDF|IMAGE CONVERTER].

The centroid secondary structure in dot-bracket notation with a minimum free energy of **-39.10** kcal/mol is given below.
[color by base-pairing probability | color by positional entropy | no coloring]

```
1     GGCAGGGGGAGGUCUUGGGGCACUGGAAAGGGGCUGAGAGGUAGCUGGAGGGAGCAGAAGGGUCCAUAGGGGCAUGGGUAAGUUGGUUGAGGUCGAGGGAGUGUGGAUACAGGCACUUGGGAGGGGCAGACAGGUCAGGGAGGGUGUGGAGGGUGGAUGA
160   GGGGGCAUUGCAGGGCUUGAGAGGCACCUGGAGGGUGCCCUGGAGCAUUGGGAAGGUC

1     ...........................................................((((((....((((......................((((((........))))))........................)))).....))))))..
160   .......((.((..((((.((.((((((((...)))))))))).)))))..)).)).....
```

You can download the minimum free energy (MFE) structure in [Vienna Format| Ct Format]. You can get thermodynamic details on this structure by submitting to our RNAeval web server.

##### Graphical output

You may look at the interactive drawing of the MFE structure below. If you do not see the interactive drawing and you are using Internet Explorer, please install the Adobe SVG plugin. **A note on base-pairing probabilities:** The structure below is colored by base-pairing probabilities. For unpaired regions the color denotes the probability of being unpaired.

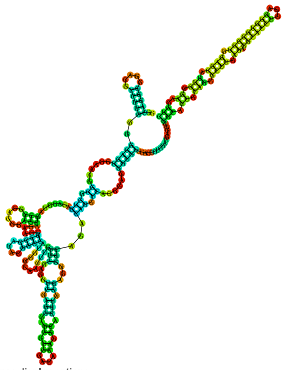

Sequence display options
- Plain Sequence
- No Sequence

**MFE secondary structure**

Other display options
- Base-pair probabilities
- Positional entropy
- None

0 1

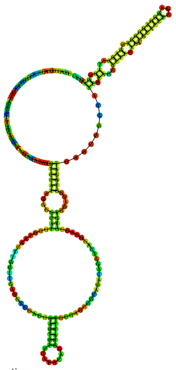

Sequence display options
- Plain Sequence
- No Sequence

**Centroid secondary structure**

Other display options
- Base-pair probabilities
- Positional entropy
- None

0 1

| Image description | Download options |
|---|---|
| MFE plain structure drawing | [ EPS \| PDF \| IMAGE CONVERTER \| VIEW IN FORNA ] |
| Centroid plain structure drawing | [ EPS \| PDF \| IMAGE CONVERTER \| VIEW IN FORNA ] |
| MFE structure drawing encoding base-pair probabilities | [ EPS \| PDF \| IMAGE CONVERTER \| VIEW IN FORNA ] |
| Centroid structure drawing encoding base-pair probabilities | [ EPS \| PDF \| IMAGE CONVERTER \| VIEW IN FORNA ] |
| MFE structure drawing encoding positional entropy | [ EPS \| PDF \| IMAGE CONVERTER \| VIEW IN FORNA ] |
| Centroid structure drawing encoding positional entropy | [ EPS \| PDF \| IMAGE CONVERTER \| VIEW IN FORNA ] |

Here you find a mountain plot representation of the MFE structure, the thermodynamic ensemble of RNA structures, and the centroid structure. Additionally we present the positional entropy for each position. Download as [EPS|PDF|IMAGE CONVERTER].

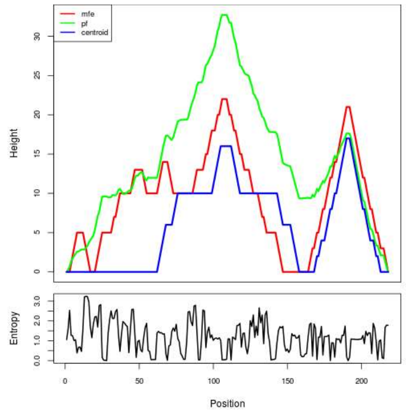


Results have been computed using RNAfold 2.6.3. An equivalent command line call would have been
`RNAfold -p -d2 --noLP < sequence1.fa > sequence1.out`

**RNA parameters are described in**
Mathews DH, Disney MD, Childs JL, Schroeder SJ, Zuker M, Turner DH. (2004) Incorporating chemical modification constraints into a dynamic programming algorithm for prediction of RNA secondary structure. *Proc Natl Acad Sci U S A* 101(19):7287-92.

**If you find these results helpful for your work you may want to cite:**

OPEN ACCESS OXFORD JOURNALS Gruber AR, Lorenz R, Bernhart SH, Neuböck R, Hofacker IL. **The Vienna RNA Websuite.** Nucleic Acids Research, Volume 36, Issue suppl_2, 1 July 2008, Pages W70-W74, DOI: 10.1093/nar/gkn188

Lorenz, R. and Bernhart, S.H. and Höner zu Siederdissen, C. and Tafer, H. and Flamm, C. and Stadler, P.F. and Hofacker, I.L. "ViennaRNA Package 2.0", Algorithms for Molecular Biology, 6:1 page(s): 26, 2011

### RNAfold WebServer MT_TOTAL_SEQ7



##### Results for minimum free energy prediction

The optimal secondary structure in dot-bracket notation with a minimum free energy of **-65.30** kcal/mol is given below.

[color by base-pairing probability | color by positional entropy | no coloring]

```
1     UGUGUCGAAGAAAACCAGAGUAUGUGCUGAUCAGAGUGAUACCGAUACUAUUACUGAUAUUGAUUGCGAGACAUGCGGGAUGGACUUGAUCGAAGGAAAUAAGGUAAGCCAUUCCAGCGAGUGUUUCGCGGCUGCUCAGUGGGAUUUAUAUACGGUCUGU
160   GUUUGGGCUAAAAUCAAUAGAGAUAAGAUACUUGAAGAGCAAAGCAAUGAGUAAGAUUCAAUUCAAAUGUUUAGGUUGCGUCUAUUCCCCGAUACUUGGAUCGACUACA

1     ((((((((...((((((((((((((((((.((((((((((.......)))))))........((((((((((((((((((.(((.(((..........))))))))))))).))..)))))))))))))).)))))).......))))))..)))).
160   ))))..(((....((....(((........)))...))....)))...(((((.(((.(((((.((....)).))))).))))))))..((((...)))).))))).)))
```

You can download the minimum free energy (MFE) structure in [Vienna Format| Ct Format]. You can get thermodynamic details on this structure by submitting to our RNAeval web server.

##### Results for thermodynamic ensemble prediction

The free energy of the thermodynamic ensemble is **-71.06** kcal/mol.
The frequency of the MFE structure in the ensemble is **0.01** %.
The ensemble diversity is **72.88** .
You may look at the **dot plot** containing the base pair probabilities [EPS|PDF|IMAGE CONVERTER].

The centroid secondary structure in dot-bracket notation with a minimum free energy of **-38.90** kcal/mol is given below.
[color by base-pairing probability | color by positional entropy | no coloring]

```
1     UGUGUCGAAGAAAACCAGAGUAUGUGCUGAUCAGAGUGAUACCGAUACUAUUACUGAUAUUGAUUGCGAGACAUGCGGGAUGGACUUGAUCGAAGGAAAUAAGGUAAGCCAUUCCAGCGAGUGUUUCGCGGCUGCUCAGUGGGAUUUAUAUACGGUCUGU
160   GUUUGGGCUAAAAUCAAUAGAGAUAAGAUACUUGAAGAGCAAAGCAAUGAGUAAGAUUCAAUUCAAAUGUUUAGGUUGCGUCUAUUCCCCGAUACUUGGAUCGACUACA

1     .(((((((..........(((....((((.((((((((((.......)))))))........((((((((((((((((((.(((.(((..........)))))))))))))).))..)))))))))).))).)))).............((((.(
160   (...((.......))..)).))))..........................((..(((..............)))..))..........)))....))))).)).
```

You can download the minimum free energy (MFE) structure in [Vienna Format| Ct Format]. You can get thermodynamic details on this structure by submitting to our RNAeval web server.

##### Graphical output

You may look at the interactive drawing of the MFE structure below. If you do not see the interactive drawing and you are using Internet Explorer, please install the Adobe SVG plugin. **A note on base-pairing probabilities:** The structure below is colored by base-pairing probabilities. For unpaired regions the color denotes the probability of being unpaired.

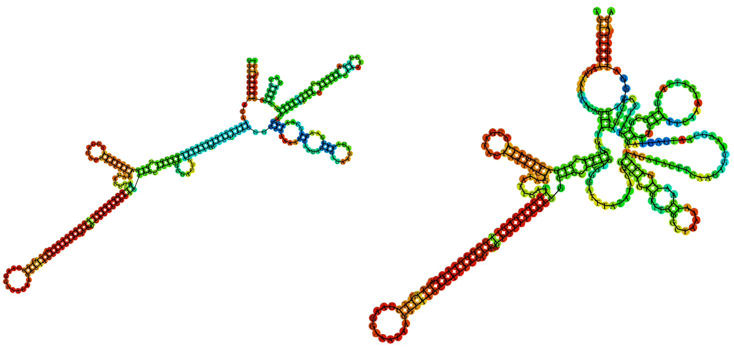

Sequence display options
- (•) Plain Sequence
- ( ) No Sequence

**MFE secondary structure**

Other display options
- (•) Base-pair probabilities
- ( ) Positional entropy
- ( ) None

0 1

Sequence display options
- (•) Plain Sequence
- ( ) No Sequence

**Centroid secondary structure**

Other display options
- (•) Base-pair probabilities
- ( ) Positional entropy
- ( ) None

0 1

| **Image description** | **Download options** |
| --- | --- |
| MFE plain structure drawing | [ EPS \| PDF \| IMAGE CONVERTER \| VIEW IN FORNA ] |
| Centroid plain structure drawing | [ EPS \| PDF \| IMAGE CONVERTER \| VIEW IN FORNA ] |
| MFE structure drawing encoding base-pair probabilities | [ EPS \| PDF \| IMAGE CONVERTER \| VIEW IN FORNA ] |
| Centroid structure drawing encoding base-pair probabilities | [ EPS \| PDF \| IMAGE CONVERTER \| VIEW IN FORNA ] |
| MFE structure drawing encoding positional entropy | [ EPS \| PDF \| IMAGE CONVERTER \| VIEW IN FORNA ] |
| Centroid structure drawing encoding positional entropy | [ EPS \| PDF \| IMAGE CONVERTER \| VIEW IN FORNA ] |

Here you find a mountain plot representation of the MFE structure, the thermodynamic ensemble of RNA structures, and the centroid structure. Additionally we present the positional entropy for each position. Download as [EPS|PDF|IMAGE CONVERTER].

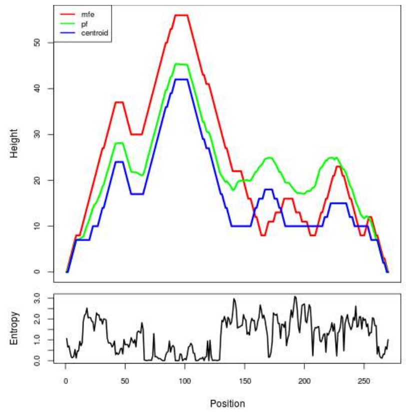


Results have been computed using RNAfold 2.6.3. An equivalent command line call would have been

```
RNAfold -p -d2 --noLP < sequence1.fa > sequence1.out
```

**RNA parameters are described in**

Mathews DH, Disney MD, Childs JL, Schroeder SJ, Zuker M, Turner DH. (2004) Incorporating chemical modification constraints into a dynamic programming algorithm for prediction of RNA secondary structure. *Proc Natl Acad Sci U S A* 101(19):7287-92.

**If you find these results helpful for your work you may want to cite:**

OPEN ACCESS OXFORD JOURNALS Gruber AR, Lorenz R, Bernhart SH, Neuböck R, Hofacker IL.
**The Vienna RNA Websuite.** Nucleic Acids Research, Volume 36, Issue suppl_2, 1 July 2008, Pages W70-W74, DOI: 10.1093/nar/gkn188

Lorenz, R. and Bernhart, S.H. and Höner zu Siederdissen, C. and Tafer, H. and Flamm, C. and Stadler, P.F. and Hofacker, I.L. "ViennaRNA Package 2.0", Algorithms for Molecular Biology, 6:1 page(s): 26, 2011

**RNAfold WebServer** MT_TOTAL_SEQ8

1 Enter Input Parameters 2 View Results



##### Results for minimum free energy prediction

The optimal secondary structure in dot-bracket notation with a minimum free energy of **-56.70** kcal/mol is given below.
[color by base-pairing probability | color by positional entropy | no coloring]

```
1      AUGGGUAUGGGGAUGGGUAUGGGAGUGGGGAUGGGUAUGGGGAUGGGUAUGGAAGAAACGCUAUUAAAAAGGAUCCUAUUAUGAUUGGGAGGCAAAUGUUGAUUUGGGCUGGCCUGUCAUUUGCGUGGGUGGGCUUAGUUGAGGAGGUCACGGCGGAAGG
160    AUACCGGCUCAGUAAUGCCGGUAUGCUGUGUCGUACAGGUGGCAUCUCGUGGA

1      .........((((((....((..((((.......((((........))))........))))..........(((((.......)))))(((.....((((((.(((((.((((...........)))).))))))))))).....)))((((((..((.
160    ((((((((........)))))))).)).))))))).)).....)))))).....
```

You can download the minimum free energy (MFE) structure in [Vienna Format| Ct Format]. You can get thermodynamic details on this structure by submitting to our RNAeval web server.

##### Results for thermodynamic ensemble prediction

The free energy of the thermodynamic ensemble is **-60.34** kcal/mol.
The frequency of the MFE structure in the ensemble is **0.27** %.
The ensemble diversity is **60.92** .
You may look at the **dot plot** containing the base pair probabilities [EPS|PDF|IMAGE CONVERTER].

The centroid secondary structure in dot-bracket notation with a minimum free energy of **-40.57** kcal/mol is given below.
[color by base-pairing probability | color by positional entropy | no coloring]

```
1      AUGGGUAUGGGGAUGGGUAUGGGAGUGGGGAUGGGUAUGGGGAUGGGUAUGGAAGAAACGCUAUUAAAAAGGAUCCUAUUAUGAUUGGGAGGCAAAUGUUGAUUUGGGCUGGCCUGUCAUUUGCGUGGGUGGGCUUAGUUGAGGAGGUCACGGCGGAAGG
160    AUACCGGCUCAGUAAUGCCGGUAUGCUGUGUCGUACAGGUGGCAUCUCGUGGA

1      ..........(((((.........(((..............................)))............((((.......))))................((((.(((.............))).))))..............((((((..((.
160    ((((((((........)))))))).)).)))))).........)))))......
```

You can download the minimum free energy (MFE) structure in [Vienna Format| Ct Format]. You can get thermodynamic details on this structure by submitting to our RNAeval web server.

##### Graphical output

You may look at the interactive drawing of the MFE structure below. If you do not see the interactive drawing and you are using Internet Explorer, please install the Adobe SVG plugin. **A note on base-pairing probabilities:** The structure below is colored by base-pairing probabilities. For unpaired regions the color denotes the probability of being unpaired.

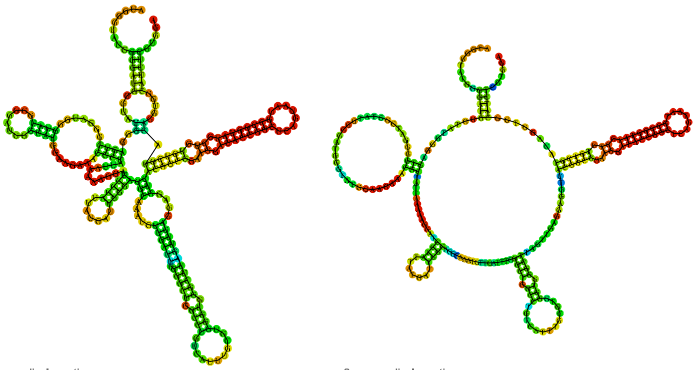

Sequence display options
- Plain Sequence
- No Sequence

**MFE secondary structure**

Other display options
- Base-pair probabilities
- Positional entropy
- None

0 1

Sequence display options
- Plain Sequence
- No Sequence

**Centroid secondary structure**

Other display options
- Base-pair probabilities
- Positional entropy
- None

0 1

| **Image description** | **Download options** |
|---|---|
| MFE plain structure drawing | [ EPS \| PDF \| IMAGE CONVERTER \| VIEW IN FORNA ] |
| Centroid plain structure drawing | [ EPS \| PDF \| IMAGE CONVERTER \| VIEW IN FORNA ] |
| MFE structure drawing encoding base-pair probabilities | [ EPS \| PDF \| IMAGE CONVERTER \| VIEW IN FORNA ] |
| Centroid structure drawing encoding base-pair probabilities | [ EPS \| PDF \| IMAGE CONVERTER \| VIEW IN FORNA ] |
| MFE structure drawing encoding positional entropy | [ EPS \| PDF \| IMAGE CONVERTER \| VIEW IN FORNA ] |
| Centroid structure drawing encoding positional entropy | [ EPS \| PDF \| IMAGE CONVERTER \| VIEW IN FORNA ] |

Here you find a mountain plot representation of the MFE structure, the thermodynamic ensemble of RNA structures, and the centroid structure. Additionally we present the positional entropy for each position. Download as [EPS|PDF|IMAGE CONVERTER].

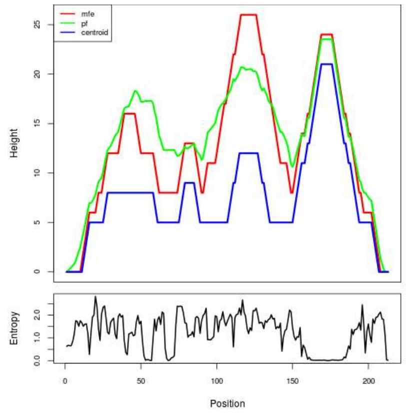


Results have been computed using RNAfold 2.6.3. An equivalent command line call would have been

```
RNAfold -p -d2 --noLP < sequence1.fa > sequence1.out
```

**RNA parameters are described in**
Mathews DH, Disney MD, Childs JL, Schroeder SJ, Zuker M, Turner DH. (2004) Incorporating chemical modification constraints into a dynamic programming algorithm for prediction of RNA secondary structure. *Proc Natl Acad Sci U S A* 101(19):7287-92.

**If you find these results helpful for your work you may want to cite:**

OPEN ACCESS OXFORD JOURNALS Gruber AR, Lorenz R, Bernhart SH, Neuböck R, Hofacker IL.
**The Vienna RNA Websuite.** Nucleic Acids Research, Volume 36, Issue suppl_2, 1 July 2008, Pages W70-W74, DOI: 10.1093/nar/gkn188

Lorenz, R. and Bernhart, S.H. and Höner zu Siederdissen, C. and Tafer, H. and Flamm, C. and Stadler, P.F. and Hofacker, I.L. "ViennaRNA Package 2.0", Algorithms for Molecular Biology, 6:1 page(s): 26, 2011

**RNAfold WebServer**

### MT_TOTAL_SEQ9

1 Enter Input Parameters 2 View Results



##### Results for minimum free energy prediction

The optimal secondary structure in dot-bracket notation with a minimum free energy of **-77.80** kcal/mol is given below.

[color by base-pairing probability | color by positional entropy | no coloring]

```
1      UGGUGGUGGUACUGGACAACAAAAUACAAACUUUGGAAGUGGAACUGCAGGUCAGGGAAAUAAUGGAGCUACAUAUAACGGAGUAAGUGGUUAUGGUUCUAAUGGAGGAGGGGACAUAAUACAACAGGAUUUCAGUCAGGUACUAAUGAUUUUGCAAAG
160    GGAGGAAGAGGCGGAUUAGGAACUAAUUUUAGUAUAAGCGGUACAACAAAAUGUUAUGCAGGUGGGGAGGAGGUGGUACACACGAAGCUAAUACAACAAGUACUAGAUCACUUGGAGGUUGUGAUAAUUCAGGACAAGGAGCUGUCGAAGGUGCAGGUU
320    UACAAGGUACAAAUGCCACAAAUGGUUUUGGUGGUGGUGGAGGAGGAAAUGGUCGUAUAGCUUUUGCAGGUACAAAUAAUGGAGUGGGAGGAUCAGGAGUAUUAAUUAUUAGAUAUUUAACACCAAG

1      (((((.(((((((.(((..((((........))))((..((......))..))...(((((..((.((((((.(((......))).))))))...(((((...........))))).........))..))))).))).))))))).(((((((.((...
160    ..........((((((((...))))))))..((((.(((.(((.((((...)))).))).............(((.....)))...))).))))..(((((........)))))...(((.(((....))).))).((((((((((((..((.....((
320    (((....(((....((((....))))....)))...)))))).......))..))).)))))))))....))..............)).)).))))).(((((((........)))))))..)))))..
```

You can download the minimum free energy (MFE) structure in [Vienna Format| Ct Format]. You can get thermodynamic details on this structure by submitting to our RNAeval web server.

##### Results for thermodynamic ensemble prediction

The free energy of the thermodynamic ensemble is **-88.39** kcal/mol.
The frequency of the MFE structure in the ensemble is **0.00** %.
The ensemble diversity is **129.79**.
You may look at the **dot plot** containing the base pair probabilities [EPS|PDF|IMAGE CONVERTER].

The centroid secondary structure in dot-bracket notation with a minimum free energy of **-35.40** kcal/mol is given below.

[color by base-pairing probability | color by positional entropy | no coloring]

```
1      UGGUGGUGGUACUGGACAACAAAAUACAAACUUUGGAAGUGGAACUGCAGGUCAGGGAAAUAAUGGAGCUACAUAUAACGGAGUAAGUGGUUAUGGUUCUAAUGGAGGAGGGGACAUAAUACAACAGGAUUUCAGUCAGGUACUAAUGAUUUUGCAAAG
160    GGAGGAAGAGGCGGAUUAGGAACUAAUUUUAGUAUAAGCGGUACAACAAAAUGUUAUGCAGGUGGGGAGGAGGUGGUACACACGAAGCUAAUACAACAAGUACUAGAUCACUUGGAGGUUGUGAUAAUUCAGGACAAGGAGCUGUCGAAGGUGCAGGUU
320    UACAAGGUACAAAUGCCACAAAUGGUUUUGGUGGUGGUGGAGGAGGAAAUGGUCGUAUAGCUUUUGCAGGUACAAAUAAUGGAGUGGGAGGAUCAGGAGUAUUAAUUAUUAGAUAUUUAACACCAAG

1      (((((.(((((((.(((.............(((((.........((.((..............)).)).........))))).......(((((....................))))).............))).))))))).......(((...
160    ..............((((((...))))))..............................................(((.....)))...............................(((.(((....))).)))......................
320    ...............((.....))........................................)))............................(((((((........)))))))..)))))..
```

You can download the minimum free energy (MFE) structure in [Vienna Format| Ct Format]. You can get thermodynamic details on this structure by submitting to our RNAeval web server.

##### Graphical output

You may look at the interactive drawing of the MFE structure below. If you do not see the interactive drawing and you are using Internet Explorer, please install the Adobe SVG plugin. **A note on base-pairing probabilities:** The structure below is colored by base-pairing probabilities. For unpaired regions the color denotes the probability of being unpaired.

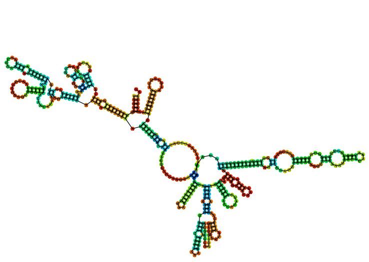

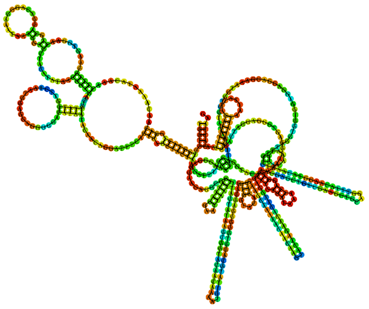

Sequence display options
- Plain Sequence
- No Sequence

**MFE secondary structure**

Other display options
- Base-pair probabilities
- Positional entropy
- None

0 1

Sequence display options
- Plain Sequence
- No Sequence

**Centroid secondary structure**

Other display options
- Base-pair probabilities
- Positional entropy
- None

0 1

| Image description | Download options |
|---|---|
| MFE plain structure drawing | [ EPS \| PDF \| IMAGE CONVERTER \| VIEW IN FORNA ] |
| Centroid plain structure drawing | [ EPS \| PDF \| IMAGE CONVERTER \| VIEW IN FORNA ] |
| MFE structure drawing encoding base-pair probabilities | [ EPS \| PDF \| IMAGE CONVERTER \| VIEW IN FORNA ] |
| Centroid structure drawing encoding base-pair probabilities | [ EPS \| PDF \| IMAGE CONVERTER \| VIEW IN FORNA ] |
| MFE structure drawing encoding positional entropy | [ EPS \| PDF \| IMAGE CONVERTER \| VIEW IN FORNA ] |
| Centroid structure drawing encoding positional entropy | [ EPS \| PDF \| IMAGE CONVERTER \| VIEW IN FORNA ] |

Here you find a mountain plot representation of the MFE structure, the thermodynamic ensemble of RNA structures, and the centroid structure. Additionally we present the positional entropy for each position. Download as [EPS|PDF|IMAGE CONVERTER].

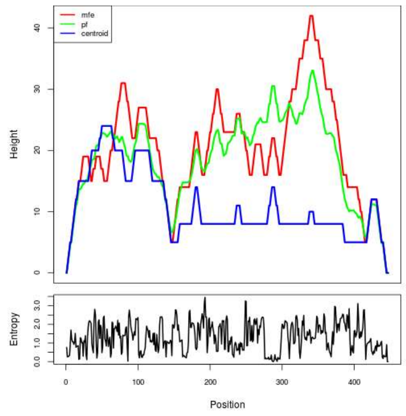


Results have been computed using RNAfold 2.6.3. An equivalent command line call would have been

`RNAfold -p -d2 --noLP < sequence1.fa > sequence1.out`


**RNA parameters are described in**

Mathews DH, Disney MD, Childs JL, Schroeder SJ, Zuker M, Turner DH. (2004) Incorporating chemical modification constraints into a dynamic programming algorithm for prediction of RNA secondary structure. *Proc Natl Acad Sci U S A* 101(19):7287-92.

**If you find these results helpful for your work you may want to cite:**

OPEN ACCESS OXFORD JOURNALS Gruber AR, Lorenz R, Bernhart SH, Neuböck R, Hofacker IL. **The Vienna RNA Websuite.** Nucleic Acids Research, Volume 36, Issue suppl_2, 1 July 2008, Pages W70-W74, DOI: 10.1093/nar/gkn188

Lorenz, R. and Bernhart, S.H. and Höner zu Siederdissen, C. and Tafer, H. and Flamm, C. and Stadler, P.F. and Hofacker, I.L. "ViennaRNA Package 2.0", Algorithms for Molecular Biology, 6:1 page(s): 26, 2011

![RNAfold WebServer] MT_TOTAL_SEQ10

1 Enter Input Parameters 2 View Results

[Home|New job|Help]

##### Results for minimum free energy prediction

The optimal secondary structure in dot-bracket notation with a minimum free energy of **-54.40** kcal/mol is given below.

[color by base-pairing probability | color by positional entropy | no coloring]

```
1     CUUGGUGUUAAAUAUCUAAUAAUUAAUACUCCUGAUCCUCCCACUCCAUUAUUUGUACCUGCAAAAGCUAUACGACCAUUUCCUCCUCCACCACCACCAAAACCAUUUGUGGCAUUUGUACCUUGUAAACCUGCACCUUCGACAGCUCCUUGUCCUGAAU
160   UAUCACAACCUCCAAGUGAUCUAGUACUUGUUGUAUUAGCUUCGUGUGUACCACCUCCUCCCCCACCUGCAUAACAUUUUGUUGUACCGCUUAUACUAAAAUUAGUUCCUAAUCCGCCUCUUCCUCCCUUUGCAAAAUCAUUAGUACCUGACUGAAAUCC
320   UGUUGUAUUAUGUCCCCCUCCUCCAUUAGAACCAUAACCACUUACUCCGUUAUAUGUAGCUCCAUUAUUUCCCUGACCUGCAGUUCCACUUCCAAAGUUUGUAUUUUGUUGUCCAGUACCACCACCACUACCUCCAUUUGAUGCUACACUUGUUAAAGCU
480   CCUGUACUUCCGUUACCU

1     ...((((((((.(((...))).)))))))).............................((((..((((....((........))..................((..((((((((.......((((....)))).....(((((...))))).......
160   .(((((.........)))))(((((((.....)))))))....(((.((((.................((((((((...))))))...))....((((....))))........................((((.....(((((.....))))).....
320   ..))))).........................(((((..........))))).(((((.((............)).)))))..........((((((....))))))........)))))))................)))))))))..))....))))
480   ..)))).........
```

You can download the minimum free energy (MFE) structure in [Vienna Format| Ct Format]. You can get thermodynamic details on this structure by submitting to our RNAeval web server.

##### Results for thermodynamic ensemble prediction

The free energy of the thermodynamic ensemble is **-64.47** kcal/mol.
The frequency of the MFE structure in the ensemble is **0.00** %.
The ensemble diversity is **127.20**.
You may look at the **dot plot** containing the base pair probabilities [EPS|PDF|IMAGE CONVERTER].

The centroid secondary structure in dot-bracket notation with a minimum free energy of **-18.80** kcal/mol is given below.

[color by base-pairing probability | color by positional entropy | no coloring]

```
1     CUUGGUGUUAAAUAUCUAAUAAUUAAUACUCCUGAUCCUCCCACUCCAUUAUUUGUACCUGCAAAAGCUAUACGACCAUUUCCUCCUCCACCACCACCAAAACCAUUUGUGGCAUUUGUACCUUGUAAACCUGCACCUUCGACAGCUCCUUGUCCUGAAU
160   UAUCACAACCUCCAAGUGAUCUAGUACUUGUUGUAUUAGCUUCGUGUGUACCACCUCCUCCCCCACCUGCAUAACAUUUUGUUGUACCGCUUAUACUAAAAUUAGUUCCUAAUCCGCCUCUUCCUCCCUUUGCAAAAUCAUUAGUACCUGACUGAAAUCC
320   UGUUGUAUUAUGUCCCCCUCCUCCAUUAGAACCAUAACCACUUACUCCGUUAUAUGUAGCUCCAUUAUUUCCCUGACCUGCAGUUCCACUUCCAAAGUUUGUAUUUUGUUGUCCAGUACCACCACCACUACCUCCAUUUGAUGCUACACUUGUUAAAGCU
480   CCUGUACUUCCGUUACCU

1     ...((((((((.((.....)).))))))))..............................................................(((.............)))...........((((....))))......((((......))))......
160   .(((((.........)))))(((((((.....)))))))......................................................((((....))))..................................................
320   ................................(((((..........)))))..................................((((((....))))))..................................................
480   .................
```

You can download the minimum free energy (MFE) structure in [Vienna Format| Ct Format]. You can get thermodynamic details on this structure by submitting to our RNAeval web server.

##### Graphical output

You may look at the interactive drawing of the MFE structure below. If you do not see the interactive drawing and you are using Internet Explorer, please install the Adobe SVG plugin. **A note on base-pairing probabilities:** The structure below is colored by base-pairing probabilities. For unpaired regions the color denotes the probability of being unpaired.

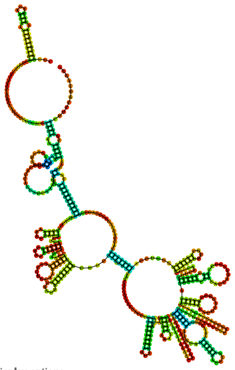

Sequence display options
- (•) Plain Sequence
- ( ) No Sequence

MFE secondary structure

Other display options
- (•) Base-pair probabilities
- ( ) Positional entropy
- ( ) None

0 — 1

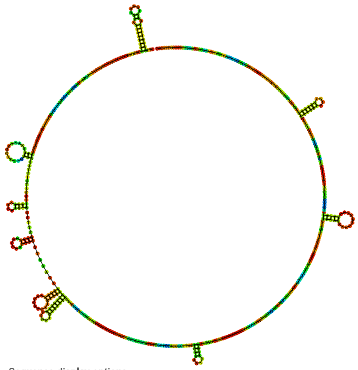

Sequence display options
- (•) Plain Sequence
- ( ) No Sequence

Centroid secondary structure

Other display options
- (•) Base-pair probabilities
- ( ) Positional entropy
- ( ) None

0 — 1

| Image description | Download options |
|---|---|
| MFE plain structure drawing | [ EPS \| PDF \| IMAGE CONVERTER \| VIEW IN FORNA ] |
| Centroid plain structure drawing | [ EPS \| PDF \| IMAGE CONVERTER \| VIEW IN FORNA ] |
| MFE structure drawing encoding base-pair probabilities | [ EPS \| PDF \| IMAGE CONVERTER \| VIEW IN FORNA ] |
| Centroid structure drawing encoding base-pair probabilities | [ EPS \| PDF \| IMAGE CONVERTER \| VIEW IN FORNA ] |
| MFE structure drawing encoding positional entropy | [ EPS \| PDF \| IMAGE CONVERTER \| VIEW IN FORNA ] |
| Centroid structure drawing encoding positional entropy | [ EPS \| PDF \| IMAGE CONVERTER \| VIEW IN FORNA ] |

Here you find a mountain plot representation of the MFE structure, the thermodynamic ensemble of RNA structures, and the centroid structure. Additionally we present the positional entropy for each position. Download as [EPS|PDF|IMAGE CONVERTER].

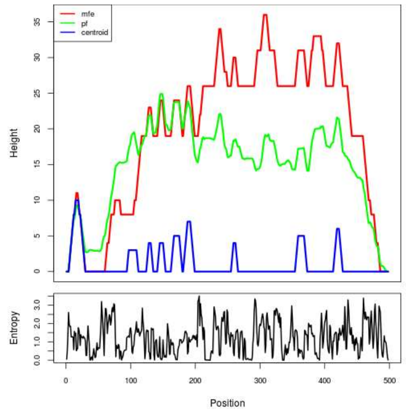


Results have been computed using RNAfold 2.6.3. An equivalent command line call would have been

```
RNAfold -p -d2 --noLP < sequence1.fa > sequence1.out
```

**RNA parameters are described in**


Mathews DH, Disney MD, Childs JL, Schroeder SJ, Zuker M, Turner DH. (2004) Incorporating chemical modification constraints into a dynamic programming algorithm for prediction of RNA secondary structure. *Proc Natl Acad Sci U S A* 101(19):7287-92.


**If you find these results helpful for your work you may want to cite:**


OPEN ACCESS OXFORD JOURNALS Gruber AR, Lorenz R, Bernhart SH, Neuböck R, Hofacker IL.
**The Vienna RNA Websuite.** Nucleic Acids Research, Volume 36, Issue suppl_2, 1 July 2008, Pages W70-W74, DOI: 10.1093/nar/gkn188

Lorenz, R. and Bernhart, S.H. and Höner zu Siederdissen, C. and Tafer, H. and Flamm, C. and Stadler, P.F. and Hofacker, I.L. "ViennaRNA Package 2.0", Algorithms for Molecular Biology, 6:1 page(s): 26, 2011

### RNAfold WebServer MT_TOTAL_SEQ11



##### Results for minimum free energy prediction

The optimal secondary structure in dot-bracket notation with a minimum free energy of **-46.30** kcal/mol is given below.

[color by base-pairing probability | color by positional entropy | no coloring]

```
1     AUACCUGUGCGGCCUUCGCCUGCUGCCAGGUGCAUCCACGAGAUGCCACCUGUACGACACAGCAUACCGGCAUUACUGAGCCGGUAUCCUUCCGCCGUGACCUCCUCAACUAAGCCCACCCACGCAAAUGACAGGCCAGCCCAAAUCAACAUUUGCCUCC
160   CAAUCAUAAUAGGAUCCUUUUUAAUAGCGUUUCUUCCAUACCCAUCCCCAUACCCAUCCCCACUCCCAUACCCAUCCCCAUACCCAUACCCAUCCCCAUACCCAUGCCCACCCCCAUACCCAUUCCCAUACCCAUCCCCAUCC

1     ...((((((((((........)))))((((((((((.....)))).))))))).....(((.((((((((((........))))))))......)).)))......................(((((((...................)))))))....
160   ........)))))...............................................................................................................................
```

You can download the minimum free energy (MFE) structure in [Vienna Format| Ct Format]. You can get thermodynamic details on this structure by submitting to our RNAeval web server.

##### Results for thermodynamic ensemble prediction

The free energy of the thermodynamic ensemble is **-50.57** kcal/mol.
The frequency of the MFE structure in the ensemble is **0.10** %.
The ensemble diversity is **53.47** .
You may look at the **dot plot** containing the base pair probabilities [EPS|PDF|IMAGE CONVERTER].

The centroid secondary structure in dot-bracket notation with a minimum free energy of **-17.90** kcal/mol is given below.

[color by base-pairing probability | color by positional entropy | no coloring]

```
1     AUACCUGUGCGGCCUUCGCCUGCUGCCAGGUGCAUCCACGAGAUGCCACCUGUACGACACAGCAUACCGGCAUUACUGAGCCGGUAUCCUUCCGCCGUGACCUCCUCAACUAAGCCCACCCACGCAAAUGACAGGCCAGCCCAAAUCAACAUUUGCCUCC
160   CAAUCAUAAUAGGAUCCUUUUUAAUAGCGUUUCUUCCAUACCCAUCCCCAUACCCAUCCCCACUCCCAUACCCAUCCCCAUACCCAUACCCAUCCCCAUACCCAUGCCCACCCCCAUACCCAUUCCCAUACCCAUCCCCAUCC

1     .........................(((((.((((.....))))..)))))..........((((((((........))))))))...................................................((........))......
160   ...........................................................................................................................................
```

You can download the minimum free energy (MFE) structure in [Vienna Format| Ct Format]. You can get thermodynamic details on this structure by submitting to our RNAeval web server.

##### Graphical output

You may look at the interactive drawing of the MFE structure below. If you do not see the interactive drawing and you are using Internet Explorer, please install the Adobe SVG plugin. **A note on base-pairing probabilities:** The structure below is colored by base-pairing probabilities. For unpaired regions the color denotes the probability of being unpaired.

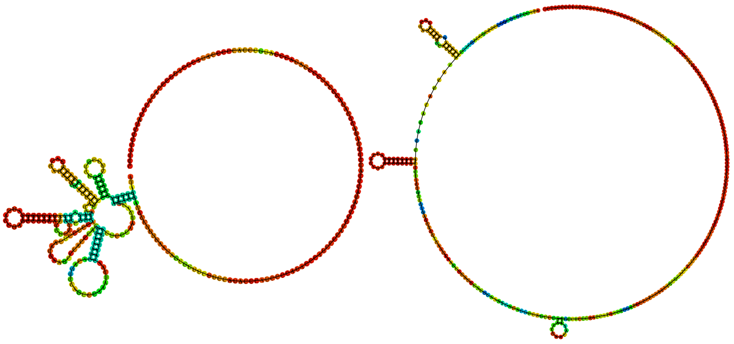

Sequence display options
- Plain Sequence
- No Sequence

**MFE secondary structure**

Other display options
- Base-pair probabilities
- Positional entropy
- None

0 1

Sequence display options
- Plain Sequence
- No Sequence

**Centroid secondary structure**

Other display options
- Base-pair probabilities
- Positional entropy
- None

0 1

| **Image description** | **Download options** |
|---|---|
| MFE plain structure drawing | [ EPS \| PDF \| IMAGE CONVERTER \| VIEW IN FORNA ] |
| Centroid plain structure drawing | [ EPS \| PDF \| IMAGE CONVERTER \| VIEW IN FORNA ] |
| MFE structure drawing encoding base-pair probabilities | [ EPS \| PDF \| IMAGE CONVERTER \| VIEW IN FORNA ] |
| Centroid structure drawing encoding base-pair probabilities | [ EPS \| PDF \| IMAGE CONVERTER \| VIEW IN FORNA ] |
| MFE structure drawing encoding positional entropy | [ EPS \| PDF \| IMAGE CONVERTER \| VIEW IN FORNA ] |
| Centroid structure drawing encoding positional entropy | [ EPS \| PDF \| IMAGE CONVERTER \| VIEW IN FORNA ] |

Here you find a mountain plot representation of the MFE structure, the thermodynamic ensemble of RNA structures, and the centroid structure. Additionally we present the positional entropy for each position. Download as [EPS|PDF|IMAGE CONVERTER].

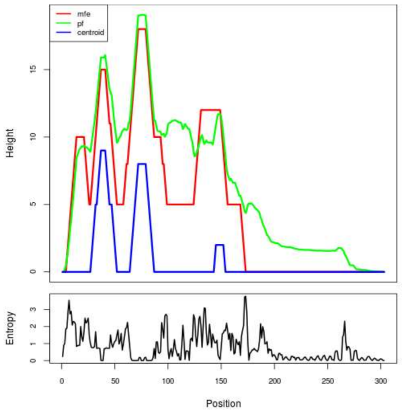


Results have been computed using RNAfold 2.6.3. An equivalent command line call would have been

```
RNAfold -p -d2 --noLP < sequence1.fa > sequence1.out
```

**RNA parameters are described in**

Mathews DH, Disney MD, Childs JL, Schroeder SJ, Zuker M, Turner DH. (2004) Incorporating chemical modification constraints into a dynamic programming algorithm for prediction of RNA secondary structure. *Proc Natl Acad Sci U S A* 101(19):7287-92.

**If you find these results helpful for your work you may want to cite:**

OPEN ACCESS OXFORD JOURNALS Gruber AR, Lorenz R, Bernhart SH, Neuböck R, Hofacker IL. **The Vienna RNA Websuite.** Nucleic Acids Research, Volume 36, Issue suppl_2, 1 July 2008, Pages W70-W74, DOI: 10.1093/nar/gkn188

Lorenz, R. and Bernhart, S.H. and Höner zu Siederdissen, C. and Tafer, H. and Flamm, C. and Stadler, P.F. and Hofacker, I.L. "ViennaRNA Package 2.0", Algorithms for Molecular Biology, 6:1 page(s): 26, 2011

### RNAfold WebServer MT_TOTAL_SEQ12

1 Enter Input Parameters 2 View Results

[Home|New job|Help]

##### Results for minimum free energy prediction

The optimal secondary structure in dot-bracket notation with a minimum free energy of **-26.10** kcal/mol is given below.

[color by base-pairing probability | color by positional entropy | no coloring]

```
1       CGCCACUCACCGACCACCGCGGAUAAUACCCCUCCACCCGAUCCACCGACCCACCCCCCGUACCCUCCACAUAAUGCCCCCUCGCCGGCGCCAACAAAUACUGCGCCCCAAAAAACUCCUCAUUAUCCAUGCACUUCGUAUCCACCGUGUCCGCAUACAU
160     CCCAGCCCCACGCUGAUCAUCCGACCCCCGAAACCCCGACCCAUACCGAUACUGCCCCAUCACCACCUGAUCCCCCUGCG

1       (((...............((((((..............((......))...........................((......)).(((((...........)))))....................((((.....))))......)))))).......
160     ..((((.....))))......((.....))......((........))..........((((.....))))......)))
```

You can download the minimum free energy (MFE) structure in [Vienna Format| Ct Format]. You can get thermodynamic details on this structure by submitting to our RNAeval web server.

##### Results for thermodynamic ensemble prediction

The free energy of the thermodynamic ensemble is **-30.07** kcal/mol.
The frequency of the MFE structure in the ensemble is **0.16** %.
The ensemble diversity is **37.09** .
You may look at the **dot plot** containing the base pair probabilities [EPS|PDF|IMAGE CONVERTER].

The centroid secondary structure in dot-bracket notation with a minimum free energy of **-12.40** kcal/mol is given below.

[color by base-pairing probability | color by positional entropy | no coloring]

```
1       CGCCACUCACCGACCACCGCGGAUAAUACCCCUCCACCCGAUCCACCGACCCACCCCCCGUACCCUCCACAUAAUGCCCCCUCGCCGGCGCCAACAAAUACUGCGCCCCAAAAAACUCCUCAUUAUCCAUGCACUUCGUAUCCACCGUGUCCGCAUACAU
160     CCCAGCCCCACGCUGAUCAUCCGACCCCCGAAACCCCGACCCAUACCGAUACUGCCCCAUCACCACCUGAUCCCCCUGCG

1       ......................................((......))...........................((......)).(((((...........)))))....................................................
160     ..((((.....))))...........................................((((.....)))).........
```

You can download the minimum free energy (MFE) structure in [Vienna Format| Ct Format]. You can get thermodynamic details on this structure by submitting to our RNAeval web server.

##### Graphical output

You may look at the interactive drawing of the MFE structure below. If you do not see the interactive drawing and you are using Internet Explorer, please install the Adobe SVG plugin. **A note on base-pairing probabilities:** The structure below is colored by base-pairing probabilities. For unpaired regions the color denotes the probability of being unpaired.

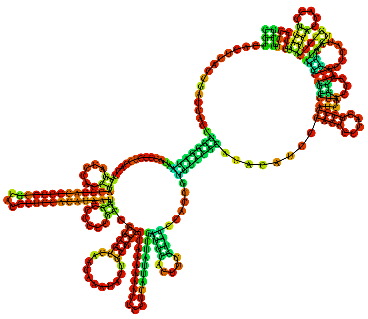

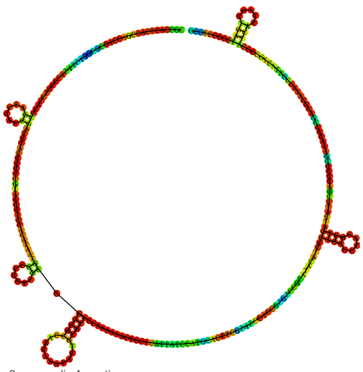

**MFE secondary structure**

Sequence display options
- (•) Plain Sequence
- ( ) No Sequence

Other display options
- (•) Base-pair probabilities
- ( ) Positional entropy
- ( ) None

0 — 1

**Centroid secondary structure**

Sequence display options
- (•) Plain Sequence
- ( ) No Sequence

Other display options
- (•) Base-pair probabilities
- ( ) Positional entropy
- ( ) None

0 — 1

| Image description | Download options |
|---|---|
| MFE plain structure drawing | [ EPS \| PDF \| IMAGE CONVERTER \| VIEW IN FORNA ] |
| Centroid plain structure drawing | [ EPS \| PDF \| IMAGE CONVERTER \| VIEW IN FORNA ] |
| MFE structure drawing encoding base-pair probabilities | [ EPS \| PDF \| IMAGE CONVERTER \| VIEW IN FORNA ] |
| Centroid structure drawing encoding base-pair probabilities | [ EPS \| PDF \| IMAGE CONVERTER \| VIEW IN FORNA ] |
| MFE structure drawing encoding positional entropy | [ EPS \| PDF \| IMAGE CONVERTER \| VIEW IN FORNA ] |
| Centroid structure drawing encoding positional entropy | [ EPS \| PDF \| IMAGE CONVERTER \| VIEW IN FORNA ] |

Here you find a mountain plot representation of the MFE structure, the thermodynamic ensemble of RNA structures, and the centroid structure. Additionally we present the positional entropy for each position. Download as [EPS|PDF|IMAGE CONVERTER].

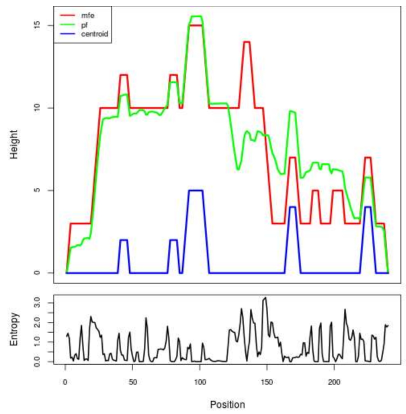


Results have been computed using RNAfold 2.6.3. An equivalent command line call would have been

```
RNAfold -p -d2 --noLP < sequence1.fa > sequence1.out
```


**RNA parameters are described in**

Mathews DH, Disney MD, Childs JL, Schroeder SJ, Zuker M, Turner DH. (2004) Incorporating chemical modification constraints into a dynamic programming algorithm for prediction of RNA secondary structure. *Proc Natl Acad Sci U S A* 101(19):7287-92.

**If you find these results helpful for your work you may want to cite:**

OPEN ACCESS OXFORD JOURNALS Gruber AR, Lorenz R, Bernhart SH, Neuböck R, Hofacker IL. **The Vienna RNA Websuite.** Nucleic Acids Research, Volume 36, Issue suppl_2, 1 July 2008, Pages W70-W74, DOI: 10.1093/nar/gkn188

Lorenz, R. and Bernhart, S.H. and Höner zu Siederdissen, C. and Tafer, H. and Flamm, C. and Stadler, P.F. and Hofacker, I.L. "ViennaRNA Package 2.0", Algorithms for Molecular Biology, 6:1 page(s): 26, 2011

RNAfold WebServer

### MT_TOTAL_SEQ13

1 Enter Input Parameters 2 View Results

[Home|New job|Help]

#### Results for minimum free energy prediction

The optimal secondary structure in dot-bracket notation with a minimum free energy of **-2.20** kcal/mol is given below.

[color by base-pairing probability | color by positional entropy | no coloring]

```
1      UUCUUCUUCUUCUUCUUCUUCUUCUUCUUCUUCUUCUUCUCCUUCUUCUCCUUCUUCUGCUUCUUCUUCUUCUUCUCCUUCUUCUGCUUCUGCUUCUCCUUCUCCUCCUCCUCCUCCUUCUUCUUCUUCUUCUGCUUCUGCUUCUGCUUCUUCUUCUUCUUCU
160    UCUUCUUCUCCUCCUGCUUCUCAUCCUUCUUCUCCUCCUUCUUCUCCUCCUCCUCCUCCUCCUUCUCCUUCUU

1      ..................................................................................((....))......................................((....))......................
160    .........................................................................
```

You can download the minimum free energy (MFE) structure in [Vienna Format| Ct Format]. You can get thermodynamic details on this structure by submitting to our RNAeval web server.

#### Results for thermodynamic ensemble prediction

The free energy of the thermodynamic ensemble is **-3.15** kcal/mol.
The frequency of the MFE structure in the ensemble is **21.35** %.
The ensemble diversity is **4.65** .
You may look at the **dot plot** containing the base pair probabilities [EPS|PDF|IMAGE CONVERTER].

The centroid secondary structure in dot-bracket notation with a minimum free energy of **-1.10** kcal/mol is given below.

[color by base-pairing probability | color by positional entropy | no coloring]

```
1      UUCUUCUUCUUCUUCUUCUUCUUCUUCUUCUUCUUCUUCUCCUUCUUCUCCUUCUUCUGCUUCUUCUUCUUCUUCUCCUUCUUCUGCUUCUGCUUCUCCUUCUCCUCCUCCUCCUCCUUCUUCUUCUUCUUCUGCUUCUGCUUCUGCUUCUUCUUCUUCUUCU
160    UCUUCUUCUCCUCCUGCUUCUCAUCCUUCUUCUCCUCCUUCUUCUCCUCCUCCUCCUCCUCCUUCUCCUUCUU

1      ..................................................................................((....)).....................................................................
160    .........................................................................
```

You can download the minimum free energy (MFE) structure in [Vienna Format| Ct Format]. You can get thermodynamic details on this structure by submitting to our RNAeval web server.

#### Graphical output

You may look at the interactive drawing of the MFE structure below. If you do not see the interactive drawing and you are using Internet Explorer, please install the Adobe SVG plugin. **A note on base-pairing probabilities:** The structure below is colored by base-pairing probabilities. For unpaired regions the color denotes the probability of being unpaired.

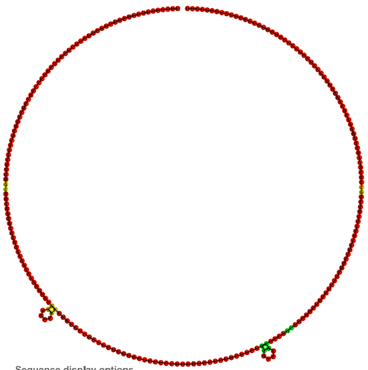

MFE secondary structure

Sequence display options
- Plain Sequence
- No Sequence

Other display options
- Base-pair probabilities
- Positional entropy
- None

0 1

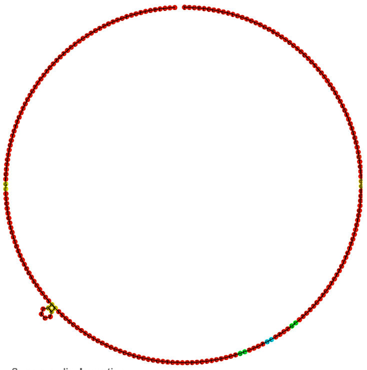

Centroid secondary structure

Sequence display options
- Plain Sequence
- No Sequence

Other display options
- Base-pair probabilities
- Positional entropy
- None

0 1

| Image description | Download options |
|---|---|
| MFE plain structure drawing | [ EPS \| PDF \| IMAGE CONVERTER \| VIEW IN FORNA ] |
| Centroid plain structure drawing | [ EPS \| PDF \| IMAGE CONVERTER \| VIEW IN FORNA ] |
| MFE structure drawing encoding base-pair probabilities | [ EPS \| PDF \| IMAGE CONVERTER \| VIEW IN FORNA ] |
| Centroid structure drawing encoding base-pair probabilities | [ EPS \| PDF \| IMAGE CONVERTER \| VIEW IN FORNA ] |
| MFE structure drawing encoding positional entropy | [ EPS \| PDF \| IMAGE CONVERTER \| VIEW IN FORNA ] |
| Centroid structure drawing encoding positional entropy | [ EPS \| PDF \| IMAGE CONVERTER \| VIEW IN FORNA ] |

Here you find a mountain plot representation of the MFE structure, the thermodynamic ensemble of RNA structures, and the centroid structure. Additionally we present the positional entropy for each position. Download as [EPS|PDF|IMAGE CONVERTER].

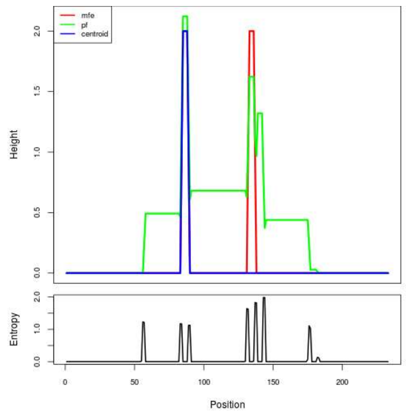


Results have been computed using RNAfold 2.6.3. An equivalent command line call would have been

```
RNAfold -p -d2 --noLP < sequence1.fa > sequence1.out
```


**RNA parameters are described in**

Mathews DH, Disney MD, Childs JL, Schroeder SJ, Zuker M, Turner DH. (2004) Incorporating chemical modification constraints into a dynamic programming algorithm for prediction of RNA secondary structure. *Proc Natl Acad Sci U S A* 101(19):7287-92.

**If you find these results helpful for your work you may want to cite:**

OPEN ACCESS OXFORD JOURNALS Gruber AR, Lorenz R, Bernhart SH, Neuböck R, Hofacker IL.
**The Vienna RNA Websuite.** Nucleic Acids Research, Volume 36, Issue suppl_2, 1 July 2008, Pages W70-W74, DOI: 10.1093/nar/gkn188

Lorenz, R. and Bernhart, S.H. and Höner zu Siederdissen, C. and Tafer, H. and Flamm, C. and Stadler, P.F. and Hofacker, I.L. "ViennaRNA Package 2.0", Algorithms for Molecular Biology, 6:1 page(s): 26, 2011

RNAfold WebServer

### MT_TOTAL_SEQ14

1 Enter Input Parameters 2 View Results

[Home|New job|Help]

##### Results for minimum free energy prediction

The optimal secondary structure in dot-bracket notation with a minimum free energy of **-32.50** kcal/mol is given below.
[color by base-pairing probability | color by positional entropy | no coloring]

```
1      AGAGGAGGGAGGAGGAGCUCACCGGAGAGGAGGGAGGAGGAGCUCACCGGAGAGGAGGGAGGAGAAACCGUGAGGGGAUGGGAGGGGAGGAGAGGAGGGAGGAGGAGGUCGCCGGAGAGGAGGAGGAGGUCGCCGGAGAGGAGGAGGAGGAGGUCGCCGG
160    AGAGGAGGAGGGUAGUAUGGUGGAGGAGAGAAGGGAAGAUGGAGUGGAGGAGAGGUGGA

1      ..........((..((((((.((............))..)))))).))...................((((......))))..........................((((((...............((.((((.((..............)).))))
160    .))..............))))))....................................
```

You can download the minimum free energy (MFE) structure in [Vienna Format| Ct Format]. You can get thermodynamic details on this structure by submitting to our RNAeval web server.

##### Results for thermodynamic ensemble prediction

The free energy of the thermodynamic ensemble is **-33.97** kcal/mol.
The frequency of the MFE structure in the ensemble is **9.28** %.
The ensemble diversity is **30.06** .
You may look at the **dot plot** containing the base pair probabilities [EPS|PDF|IMAGE CONVERTER].

The centroid secondary structure in dot-bracket notation with a minimum free energy of **-30.00** kcal/mol is given below.
[color by base-pairing probability | color by positional entropy | no coloring]

```
1      AGAGGAGGGAGGAGGAGCUCACCGGAGAGGAGGGAGGAGGAGCUCACCGGAGAGGAGGGAGGAGAAACCGUGAGGGGAUGGGAGGGGAGGAGAGGAGGGAGGAGGAGGUCGCCGGAGAGGAGGAGGAGGUCGCCGGAGAGGAGGAGGAGGAGGUCGCCGG
160    AGAGGAGGAGGGUAGUAUGGUGGAGGAGAGAAGGGAAGAUGGAGUGGAGGAGAGGUGGA

1      ..............((((((.((............))..))))))......................((((......))))...........................(((((...............((.((((.((..............)).))))
160    .))..............))))).....................................
```

You can download the minimum free energy (MFE) structure in [Vienna Format| Ct Format]. You can get thermodynamic details on this structure by submitting to our RNAeval web server.

##### Graphical output

You may look at the interactive drawing of the MFE structure below. If you do not see the interactive drawing and you are using Internet Explorer, please install the Adobe SVG plugin. **A note on base-pairing probabilities:** The structure below is colored by base-pairing probabilities. For unpaired regions the color denotes the probability of being unpaired.

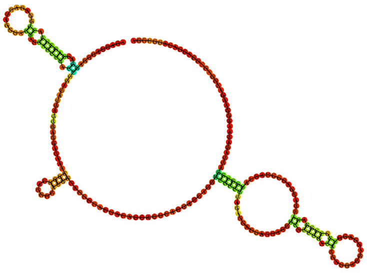

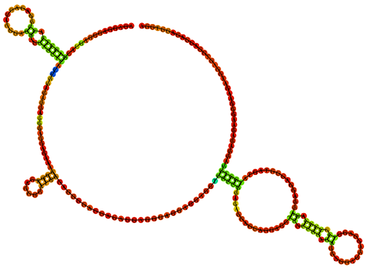

Sequence display options
- (•) Plain Sequence
- ( ) No Sequence

MFE secondary structure

Other display options
- (•) Base-pair probabilities
- ( ) Positional entropy
- ( ) None

0 1

Sequence display options
- (•) Plain Sequence
- ( ) No Sequence

Centroid secondary structure

Other display options
- (•) Base-pair probabilities
- ( ) Positional entropy
- ( ) None

0 1

| Image description | Download options |
|---|---|
| MFE plain structure drawing | [ EPS \| PDF \| IMAGE CONVERTER \| VIEW IN FORNA ] |
| Centroid plain structure drawing | [ EPS \| PDF \| IMAGE CONVERTER \| VIEW IN FORNA ] |
| MFE structure drawing encoding base-pair probabilities | [ EPS \| PDF \| IMAGE CONVERTER \| VIEW IN FORNA ] |
| Centroid structure drawing encoding base-pair probabilities | [ EPS \| PDF \| IMAGE CONVERTER \| VIEW IN FORNA ] |
| MFE structure drawing encoding positional entropy | [ EPS \| PDF \| IMAGE CONVERTER \| VIEW IN FORNA ] |
| Centroid structure drawing encoding positional entropy | [ EPS \| PDF \| IMAGE CONVERTER \| VIEW IN FORNA ] |

Here you find a mountain plot representation of the MFE structure, the thermodynamic ensemble of RNA structures, and the centroid structure. Additionally we present the positional entropy for each position. Download as [EPS|PDF|IMAGE CONVERTER].

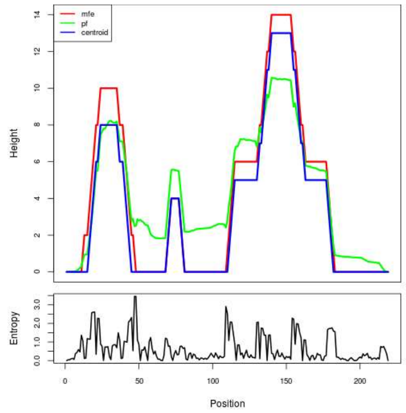


Results have been computed using RNAfold 2.6.3. An equivalent command line call would have been
`RNAfold -p -d2 --noLP < sequence1.fa > sequence1.out`

**RNA parameters are described in**
Mathews DH, Disney MD, Childs JL, Schroeder SJ, Zuker M, Turner DH. (2004) Incorporating chemical modification constraints into a dynamic programming algorithm for prediction of RNA secondary structure. *Proc Natl Acad Sci U S A* 101(19):7287-92.

**If you find these results helpful for your work you may want to cite:**

OPEN ACCESS OXFORD JOURNALS Gruber AR, Lorenz R, Bernhart SH, Neuböck R, Hofacker IL.
**The Vienna RNA Websuite.** Nucleic Acids Research, Volume 36, Issue suppl_2, 1 July 2008, Pages W70-W74, DOI: 10.1093/nar/gkn188

Lorenz, R. and Bernhart, S.H. and Höner zu Siederdissen, C. and Tafer, H. and Flamm, C. and Stadler, P.F. and Hofacker, I.L. "ViennaRNA Package 2.0", Algorithms for Molecular Biology, 6:1 page(s): 26, 2011

### MT_TOTAL_SEQ15



##### Results for minimum free energy prediction

The optimal secondary structure in dot-bracket notation with a minimum free energy of **-23.90** kcal/mol is given below.

[color by base-pairing probability | color by positional entropy | no coloring]

```
1      GACCUUCCCAAUGCUCCAGGGCACCCUCCAGGUGCCUCUCAAGCCCUGCAAUGCCCCCUCAUCCACCCUCCACACCCUCCCUGACCUGUCUGCCCCUCCCAAGUGCCUGUAUCCACACUCCCUCGACCUCAACCAACUUACCCAUGCCCCUAUGGACCCU
160    UCUGCUCCCUCCAGCUACCUCUCAGCCCCUUUCCAGUGCCCCAAGACCUCCCCCUGCC

1      ........((..(((..((((((((.....)))))).))..)))..)).....................................((((..........((((..........)))).......................(((((....))))).....
160    ...(((......)))............................))))...........
```

You can download the minimum free energy (MFE) structure in [Vienna Format| Ct Format]. You can get thermodynamic details on this structure by submitting to our RNAeval web server.

##### Results for thermodynamic ensemble prediction

The free energy of the thermodynamic ensemble is **-27.97** kcal/mol.
The frequency of the MFE structure in the ensemble is **0.14** %.
The ensemble diversity is **34.25** .
You may look at the **dot plot** containing the base pair probabilities [EPS|PDF|IMAGE CONVERTER].

The centroid secondary structure in dot-bracket notation with a minimum free energy of **-20.10** kcal/mol is given below.

[color by base-pairing probability | color by positional entropy | no coloring]

```
1      GACCUUCCCAAUGCUCCAGGGCACCCUCCAGGUGCCUCUCAAGCCCUGCAAUGCCCCCUCAUCCACCCUCCACACCCUCCCUGACCUGUCUGCCCCUCCCAAGUGCCUGUAUCCACACUCCCUCGACCUCAACCAACUUACCCAUGCCCCUAUGGACCCU
160    UCUGCUCCCUCCAGCUACCUCUCAGCCCCUUUCCAGUGCCCCAAGACCUCCCCCUGCC

1      ...........(((..((((((((.....)))))).))..))).........................................................(((..........))).......................(((((....))))).....
160    ...(((......)))...........................................
```

You can download the minimum free energy (MFE) structure in [Vienna Format| Ct Format]. You can get thermodynamic details on this structure by submitting to our RNAeval web server.

##### Graphical output

You may look at the interactive drawing of the MFE structure below. If you do not see the interactive drawing and you are using Internet Explorer, please install the Adobe SVG plugin. **A note on base-pairing probabilities:** The structure below is colored by base-pairing probabilities. For unpaired regions the color denotes the probability of being unpaired.

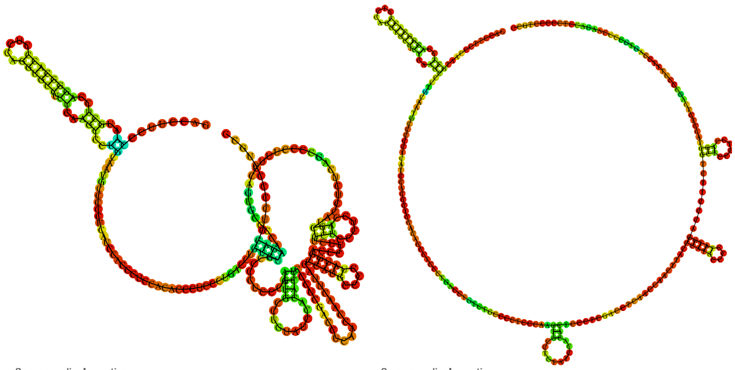

Sequence display options
- (•) Plain Sequence
- ( ) No Sequence

**MFE secondary structure**

Other display options
- (•) Base-pair probabilities
- ( ) Positional entropy
- ( ) None

0 1

Sequence display options
- (•) Plain Sequence
- ( ) No Sequence

**Centroid secondary structure**

Other display options
- (•) Base-pair probabilities
- ( ) Positional entropy
- ( ) None

0 1

| **Image description** | **Download options** |
|---|---|
| MFE plain structure drawing | [ EPS \| PDF \| IMAGE CONVERTER \| VIEW IN FORNA ] |
| Centroid plain structure drawing | [ EPS \| PDF \| IMAGE CONVERTER \| VIEW IN FORNA ] |
| MFE structure drawing encoding base-pair probabilities | [ EPS \| PDF \| IMAGE CONVERTER \| VIEW IN FORNA ] |
| Centroid structure drawing encoding base-pair probabilities | [ EPS \| PDF \| IMAGE CONVERTER \| VIEW IN FORNA ] |
| MFE structure drawing encoding positional entropy | [ EPS \| PDF \| IMAGE CONVERTER \| VIEW IN FORNA ] |
| Centroid structure drawing encoding positional entropy | [ EPS \| PDF \| IMAGE CONVERTER \| VIEW IN FORNA ] |

Here you find a mountain plot representation of the MFE structure, the thermodynamic ensemble of RNA structures, and the centroid structure. Additionally we present the positional entropy for each position. Download as [EPS|PDF|IMAGE CONVERTER].

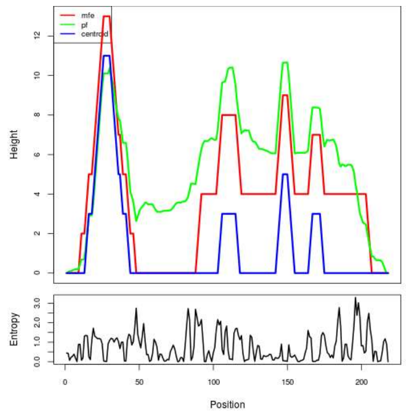


Results have been computed using RNAfold 2.6.3. An equivalent command line call would have been

```
RNAfold -p -d2 --noLP < sequence1.fa > sequence1.out
```

**RNA parameters are described in**

Mathews DH, Disney MD, Childs JL, Schroeder SJ, Zuker M, Turner DH. (2004) Incorporating chemical modification constraints into a dynamic programming algorithm for prediction of RNA secondary structure. *Proc Natl Acad Sci U S A* 101(19):7287-92.

**If you find these results helpful for your work you may want to cite:**

OPEN ACCESS OXFORD JOURNALS Gruber AR, Lorenz R, Bernhart SH, Neuböck R, Hofacker IL.
**The Vienna RNA Websuite.** Nucleic Acids Research, Volume 36, Issue suppl_2, 1 July 2008, Pages W70-W74, DOI: 10.1093/nar/gkn188

Lorenz, R. and Bernhart, S.H. and Höner zu Siederdissen, C. and Tafer, H. and Flamm, C. and Stadler, P.F. and Hofacker, I.L. "ViennaRNA Package 2.0", Algorithms for Molecular Biology, 6:1 page(s): 26, 2011

#### Section 2:

#### Secondary Structures of MT_PURE Sequences.

This section includes the predicted secondary structures of the 13 ssDNA sequences in the MT_PURE dataset. Each structure is labeled with its corresponding sequence identifier for reference.

RNAfold WebServer

### MT_PURE_SEQ1

1 Enter Input Parameters 2 View Results

[Home|New job|Help]

#### Results for minimum free energy prediction

The optimal secondary structure in dot-bracket notation with a minimum free energy of **-77.80** kcal/mol is given below.

[color by base-pairing probability | color by positional entropy | no coloring]

```
1      UGGUGGUGGUACUGGACAACAAAAUACAAACUUUGGAAGUGGAACUGCAGGUCAGGGAAAUAAUGGAGCUACAUAUAACGGAGUAAGUGGUUAUGGUUCUAAUGGAGGAGGGGGACAUAAUACAACAGGAUUUCAGUCAGGUACUAAUGAUUUUGCAAAG
160    GGAGGAAGAGGCGGAUUAGGAACUAAUUUUAGUAUAAGCGGUACAACAAAAUGUUAUGCAGGUGGGGGAGGAGGUGGUACACACGAAGCUAAUACAACAAGUACUAGAUCACUUGGAGGUUGUGAUAAUUCAGGACAAGGAGCUGUCGAAGGUGCAGGUU
320    UACAAGGUACAAAUGCCACAAAUGGUUUUGGUGGUGGUGGAGGAGGAAAUGGUCGUAUAGCUUUUGCAGGUACAAAUAAUGGAGUGGGAGGAUCAGGAGUAUUAAUUAUUAGAUAUUUAACACCAAG

1      (((((.(((((((.(((..((((........))))((..((......))..))...(((((..((.((((((.(((......))).))))))...(((((...........))))).........))..))))).))).))))))).(((((((.((...
160    ..........((((((((...))))))))..((((.(((.(((.((((...)))).)))..............(((.....)))...))).))))..(((((........)))))...(((.(((....))).))).((((((((((((..((.....((
320    (((....(((....((((....))))....)))...)))))......))..))).)))))))))....))............)).)).))))).(((((((........))))))).)))))..
```

You can download the minimum free energy (MFE) structure in [Vienna Format| Ct Format]. You can get thermodynamic details on this structure by submitting to our RNAeval web server.

#### Results for thermodynamic ensemble prediction

The free energy of the thermodynamic ensemble is **-88.39** kcal/mol.

The frequency of the MFE structure in the ensemble is **0.00** %.

The ensemble diversity is **129.79.**

You may look at the **dot plot** containing the base pair probabilities [EPS|PDF|IMAGE CONVERTER].

The centroid secondary structure in dot-bracket notation with a minimum free energy of **-35.40** kcal/mol is given below.

[color by base-pairing probability | color by positional entropy | no coloring]

```
1      UGGUGGUGGUACUGGACAACAAAAUACAAACUUUGGAAGUGGAACUGCAGGUCAGGGAAAUAAUGGAGCUACAUAUAACGGAGUAAGUGGUUAUGGUUCUAAUGGAGGAGGGGGACAUAAUACAACAGGAUUUCAGUCAGGUACUAAUGAUUUUGCAAAG
160    GGAGGAAGAGGCGGAUUAGGAACUAAUUUUAGUAUAAGCGGUACAACAAAAUGUUAUGCAGGUGGGGGAGGAGGUGGUACACACGAAGCUAAUACAACAAGUACUAGAUCACUUGGAGGUUGUGAUAAUUCAGGACAAGGAGCUGUCGAAGGUGCAGGUU
320    UACAAGGUACAAAUGCCACAAAUGGUUUUGGUGGUGGUGGAGGAGGAAAUGGUCGUAUAGCUUUUGCAGGUACAAAUAAUGGAGUGGGAGGAUCAGGAGUAUUAAUUAUUAGAUAUUUAACACCAAG

1      (((((.(((((((.(((............(((((........((.((.............)).))..........))))).......(((((....................)))))...............))).))))))).......(((...
160    ............((((((...))))))...............................................(((.....)))...................................(((.(((....))).)))......................
320    ...............((......))......................................)))............................(((((((........)))))))..)))))..
```

You can download the minimum free energy (MFE) structure in [Vienna Format| Ct Format]. You can get thermodynamic details on this structure by submitting to our RNAeval web server.

#### Graphical output

You may look at the interactive drawing of the MFE structure below. If you do not see the interactive drawing and you are using Internet Explorer, please install the Adobe SVG plugin. **A note on base-pairing probabilities:** The structure below is colored by base-pairing probabilities. For unpaired regions the color denotes the probability of being unpaired.

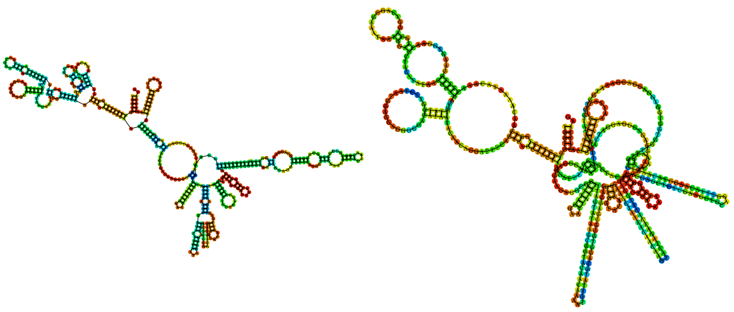

| Sequence display options | MFE secondary structure | Sequence display options | Centroid secondary structure |
|---|---|---|---|
| ◉ Plain Sequence | | ◉ Plain Sequence | |
| ○ No Sequence | | ○ No Sequence | |
| Other display options | | Other display options | |
| ◉ Base-pair probabilities | 0 — 1 | ◉ Base-pair probabilities | 0 — 1 |
| ○ Positional entropy | | ○ Positional entropy | |
| ○ None | | ○ None | |

| Image description | Download options |
|---|---|
| MFE plain structure drawing | [ EPS \| PDF \| IMAGE CONVERTER \| VIEW IN FORNA ] |
| Centroid plain structure drawing | [ EPS \| PDF \| IMAGE CONVERTER \| VIEW IN FORNA ] |
| MFE structure drawing encoding base-pair probabilities | [ EPS \| PDF \| IMAGE CONVERTER \| VIEW IN FORNA ] |
| Centroid structure drawing encoding base-pair probabilities | [ EPS \| PDF \| IMAGE CONVERTER \| VIEW IN FORNA ] |
| MFE structure drawing encoding positional entropy | [ EPS \| PDF \| IMAGE CONVERTER \| VIEW IN FORNA ] |
| Centroid structure drawing encoding positional entropy | [ EPS \| PDF \| IMAGE CONVERTER \| VIEW IN FORNA ] |

Here you find a mountain plot representation of the MFE structure, the thermodynamic ensemble of RNA structures, and the centroid structure. Additionally we present the positional entropy for each position. Download as [EPS|PDF|IMAGE CONVERTER].

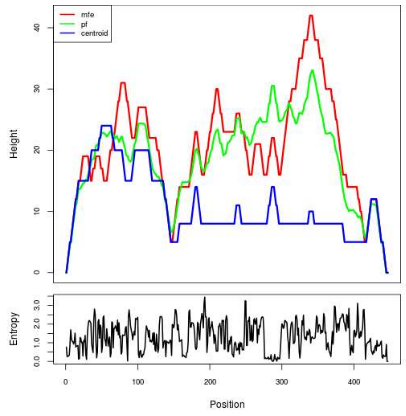


Results have been computed using RNAfold 2.6.3. An equivalent command line call would have been

`RNAfold -p -d2 --noLP < sequence1.fa > sequence1.out`


**RNA parameters are described in**

Mathews DH, Disney MD, Childs JL, Schroeder SJ, Zuker M, Turner DH. (2004) Incorporating chemical modification constraints into a dynamic programming algorithm for prediction of RNA secondary structure. *Proc Natl Acad Sci U S A* 101(19):7287-92.

**If you find these results helpful for your work you may want to cite:**

OPEN ACCESS OXFORD JOURNALS Gruber AR, Lorenz R, Bernhart SH, Neuböck R, Hofacker IL.
**The Vienna RNA Websuite.** Nucleic Acids Research, Volume 36, Issue suppl_2, 1 July 2008, Pages W70-W74, DOI: 10.1093/nar/gkn188

Lorenz, R. and Bernhart, S.H. and Höner zu Siederdissen, C. and Tafer, H. and Flamm, C. and Stadler, P.F. and Hofacker, I.L. "ViennaRNA Package 2.0", Algorithms for Molecular Biology, 6:1 page(s): 26, 2011

### RNAfold WebServer MT_PURE_SEQ2

1 Enter Input Parameters 2 View Results

[Home|New job|Help]

##### Results for minimum free energy prediction

The optimal secondary structure in dot-bracket notation with a minimum free energy of **-44.20** kcal/mol is given below.

[color by base-pairing probability | color by positional entropy | no coloring]

```
1       UGUAGUCGAUCCAAGUAUCGGGGGAAUAGACGCAACCUAAACAUUUGAAUUGAAUCUUACUCAUUGCUUUGCUCUUCAAGUAUCUUAUCUCUAUUGAUUUUAGCCCAAACACAGACCGUAUAUAAAUCCCACUGAGCAGCCGCGAAACACUCGCUGGAAU
160     GGCUUACCUUAUUUCCUUCGAUCAAGUCCAUCCCGCAUGUCUCGCAAUCAAUAUCAGUAAUAGUAUCGGUAUCACUCUGAUCAGCACAUACUCUGGUUUUCUUCGACACA

1       (((.(((((.......((((((((..(((..(((..(((....)))..)))..)))...))........((((..((((((.....(((......)))...............((((.((((.......(((((......((((.(((...((.(((.(
160     (((((..................))).))))))).)).))).))))........)))))....))))))))...))).)))..))))....)))))))).....))))))))
```

You can download the minimum free energy (MFE) structure in [Vienna Format| Ct Format]. You can get thermodynamic details on this structure by submitting to our RNAeval web server.

##### Results for thermodynamic ensemble prediction

The free energy of the thermodynamic ensemble is **-50.12** kcal/mol.
The frequency of the MFE structure in the ensemble is **0.01** %.
The ensemble diversity is **60.31** .
You may look at the **dot plot** containing the base pair probabilities [EPS|PDF|IMAGE CONVERTER].

The centroid secondary structure in dot-bracket notation with a minimum free energy of **-27.27** kcal/mol is given below.

[color by base-pairing probability | color by positional entropy | no coloring]

```
1       UGUAGUCGAUCCAAGUAUCGGGGGAAUAGACGCAACCUAAACAUUUGAAUUGAAUCUUACUCAUUGCUUUGCUCUUCAAGUAUCUUAUCUCUAUUGAUUUUAGCCCAAACACAGACCGUAUAUAAAUCCCACUGAGCAGCCGCGAAACACUCGCUGGAAU
160     GGCUUACCUUAUUUCCUUCGAUCAAGUCCAUCCCGCAUGUCUCGCAAUCAAUAUCAGUAAUAGUAUCGGUAUCACUCUGAUCAGCACAUACUCUGGUUUUCUUCGACACA

1       .((.(((((.................................................................................................((((.((((.......(((((......((((.(((...((.(((.(
160     (((((..................))).))))))).)).))).))))........)))))....))))))))..............................))))))).
```

You can download the minimum free energy (MFE) structure in [Vienna Format| Ct Format]. You can get thermodynamic details on this structure by submitting to our RNAeval web server.

##### Graphical output

You may look at the interactive drawing of the MFE structure below. If you do not see the interactive drawing and you are using Internet Explorer, please install the Adobe SVG plugin. **A note on base-pairing probabilities:** The structure below is colored by base-pairing probabilities. For unpaired regions the color denotes the probability of being unpaired.

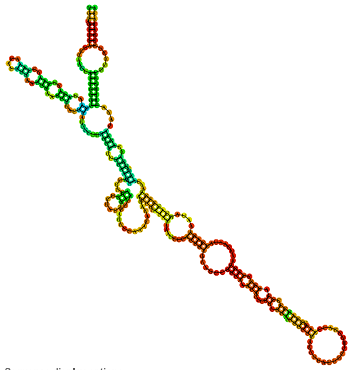

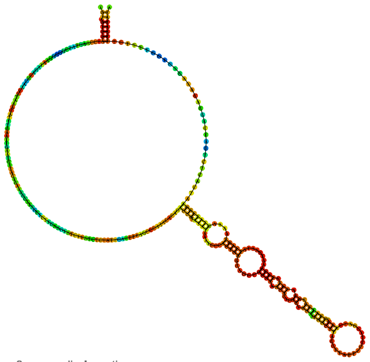

Sequence display options
◉ Plain Sequence
○ No Sequence

**MFE secondary structure**

Other display options
◉ Base-pair probabilities
○ Positional entropy
○ None

0 1

Sequence display options
◉ Plain Sequence
○ No Sequence

**Centroid secondary structure**

Other display options
◉ Base-pair probabilities
○ Positional entropy
○ None

0 1

| **Image description** | **Download options** |
|---|---|
| MFE plain structure drawing | [ EPS \| PDF \| IMAGE CONVERTER \| VIEW IN FORNA ] |
| Centroid plain structure drawing | [ EPS \| PDF \| IMAGE CONVERTER \| VIEW IN FORNA ] |
| MFE structure drawing encoding base-pair probabilities | [ EPS \| PDF \| IMAGE CONVERTER \| VIEW IN FORNA ] |
| Centroid structure drawing encoding base-pair probabilities | [ EPS \| PDF \| IMAGE CONVERTER \| VIEW IN FORNA ] |
| MFE structure drawing encoding positional entropy | [ EPS \| PDF \| IMAGE CONVERTER \| VIEW IN FORNA ] |
| Centroid structure drawing encoding positional entropy | [ EPS \| PDF \| IMAGE CONVERTER \| VIEW IN FORNA ] |

Here you find a mountain plot representation of the MFE structure, the thermodynamic ensemble of RNA structures, and the centroid structure. Additionally we present the positional entropy for each position. Download as [EPS|PDF|IMAGE CONVERTER].

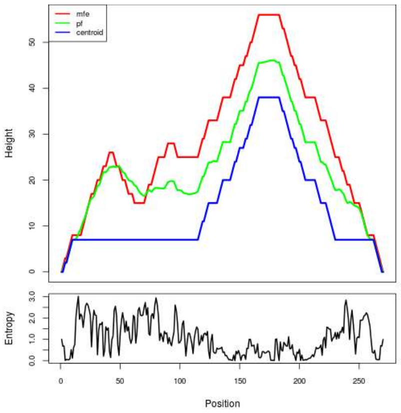


Results have been computed using RNAfold 2.6.3. An equivalent command line call would have been

```
RNAfold -p -d2 --noLP < sequence1.fa > sequence1.out
```

**RNA parameters are described in**

Mathews DH, Disney MD, Childs JL, Schroeder SJ, Zuker M, Turner DH. (2004) Incorporating chemical modification constraints into a dynamic programming algorithm for prediction of RNA secondary structure. *Proc Natl Acad Sci U S A* 101(19):7287-92.

**If you find these results helpful for your work you may want to cite:**

OPEN ACCESS OXFORD JOURNALS Gruber AR, Lorenz R, Bernhart SH, Neuböck R, Hofacker IL.
**The Vienna RNA Websuite.** Nucleic Acids Research, Volume 36, Issue suppl_2, 1 July 2008, Pages W70-W74, DOI: 10.1093/nar/gkn188

Lorenz, R. and Bernhart, S.H. and Höner zu Siederdissen, C. and Tafer, H. and Flamm, C. and Stadler, P.F. and Hofacker, I.L. "ViennaRNA Package 2.0", Algorithms for Molecular Biology, 6:1 page(s): 26, 2011

### MT_PURE_SEQ3



##### Results for minimum free energy prediction

The optimal secondary structure in dot-bracket notation with a minimum free energy of **-14.80** kcal/mol is given below.
[color by base-pairing probability | color by positional entropy | no coloring]

```
1     AAAAAAUCAAAUAAAAAUAAAAGAACGAAAAAAAAACGAAGAAACAAAGUGACAAAGUAAAAAAGUUCUUAAAAAAUUUGAUUAUAAAAAUGUCGACGACAAACGUGAUAAAAUCUAAAAAAUAAUACAAAAAUUGACAAAGUCAAAAAAGCCGAAAGUAA
160   AAAGACACAAACACAUAUACACUAUGUAAUACAUUUAACACACACUCAAACACAACAAAUUUCUGCCGUUCCUUACCCC

1     ..................................((((.(((((....(((.....((((..(((((......)))))..)))).....((((.(((.....)))))))......................(((((...)))))...............
160   ............(((((.....)))))).......................)))......)))))..))))........
```

You can download the minimum free energy (MFE) structure in [Vienna Format| Ct Format]. You can get thermodynamic details on this structure by submitting to our RNAeval web server.

##### Results for thermodynamic ensemble prediction

The free energy of the thermodynamic ensemble is **-19.39** kcal/mol.
The frequency of the MFE structure in the ensemble is **0.06** %.
The ensemble diversity is **41.89** .
You may look at the **dot plot** containing the base pair probabilities [EPS|PDF|IMAGE CONVERTER].

The centroid secondary structure in dot-bracket notation with a minimum free energy of **-11.20** kcal/mol is given below.
[color by base-pairing probability | color by positional entropy | no coloring]

```
1     AAAAAAUCAAAUAAAAAUAAAAGAACGAAAAAAAAACGAAGAAACAAAGUGACAAAGUAAAAAAGUUCUUAAAAAAUUUGAUUAUAAAAAUGUCGACGACAAACGUGAUAAAAUCUAAAAAAUAAUACAAAAAUUGACAAAGUCAAAAAAGCCGAAAGUAA
160   AAAGACACAAACACAUAUACACUAUGUAAUACAUUUAACACACACUCAAACACAACAAAUUUCUGCCGUUCCUUACCCC

1     ......................(((((...............................................................(((.(((.....))))))).......................(((((...)))))...............
160   ............(((((.....)))))......................................)))))........
```

You can download the minimum free energy (MFE) structure in [Vienna Format| Ct Format]. You can get thermodynamic details on this structure by submitting to our RNAeval web server.

##### Graphical output

You may look at the interactive drawing of the MFE structure below. If you do not see the interactive drawing and you are using Internet Explorer, please install the Adobe SVG plugin. **A note on base-pairing probabilities:** The structure below is colored by base-pairing probabilities. For unpaired regions the color denotes the probability of being unpaired.

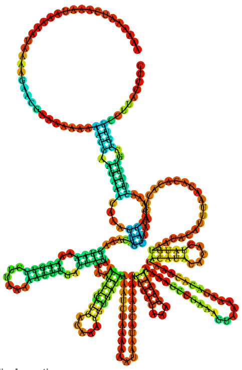

Sequence display options
- Plain Sequence
- No Sequence

**MFE secondary structure**

Other display options
- Base-pair probabilities
- Positional entropy
- None

0 1

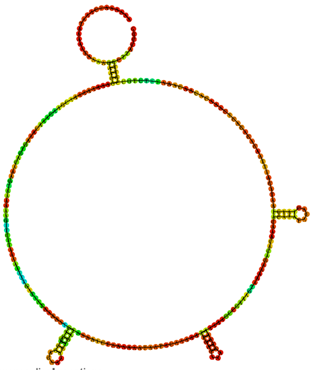

Sequence display options
- Plain Sequence
- No Sequence

**Centroid secondary structure**

Other display options
- Base-pair probabilities
- Positional entropy
- None

0 1

| Image description | Download options |
|---|---|
| MFE plain structure drawing | [ EPS \| PDF \| IMAGE CONVERTER \| VIEW IN FORNA ] |
| Centroid plain structure drawing | [ EPS \| PDF \| IMAGE CONVERTER \| VIEW IN FORNA ] |
| MFE structure drawing encoding base-pair probabilities | [ EPS \| PDF \| IMAGE CONVERTER \| VIEW IN FORNA ] |
| Centroid structure drawing encoding base-pair probabilities | [ EPS \| PDF \| IMAGE CONVERTER \| VIEW IN FORNA ] |
| MFE structure drawing encoding positional entropy | [ EPS \| PDF \| IMAGE CONVERTER \| VIEW IN FORNA ] |
| Centroid structure drawing encoding positional entropy | [ EPS \| PDF \| IMAGE CONVERTER \| VIEW IN FORNA ] |

Here you find a mountain plot representation of the MFE structure, the thermodynamic ensemble of RNA structures, and the centroid structure. Additionally we present the positional entropy for each position. Download as [EPS|PDF|IMAGE CONVERTER].

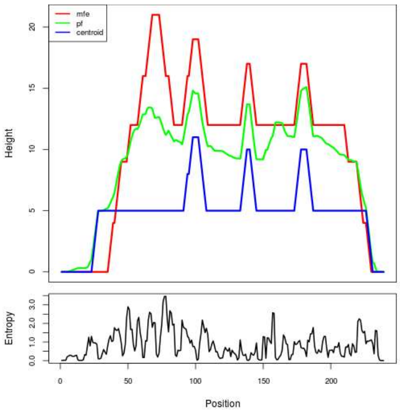


Results have been computed using RNAfold 2.6.3. An equivalent command line call would have been
`RNAfold -p -d2 --noLP < sequence1.fa > sequence1.out`

**RNA parameters are described in**
Mathews DH, Disney MD, Childs JL, Schroeder SJ, Zuker M, Turner DH. (2004) Incorporating chemical modification constraints into a dynamic programming algorithm for prediction of RNA secondary structure. *Proc Natl Acad Sci U S A* 101(19):7287-92.

**If you find these results helpful for your work you may want to cite:**

OPEN ACCESS OXFORD JOURNALS Gruber AR, Lorenz R, Bernhart SH, Neuböck R, Hofacker IL.
**The Vienna RNA Websuite.** Nucleic Acids Research, Volume 36, Issue suppl_2, 1 July 2008, Pages W70-W74, DOI: 10.1093/nar/gkn188

Lorenz, R. and Bernhart, S.H. and Höner zu Siederdissen, C. and Tafer, H. and Flamm, C. and Stadler, P.F. and Hofacker, I.L. "ViennaRNA Package 2.0", Algorithms for Molecular Biology, 6:1 page(s): 26, 2011

### RNAfold WebServer MT_PURE_SEQ4

1 Enter Input Parameters 2 View Results

[Home|New job|Help]

##### Results for minimum free energy prediction

The optimal secondary structure in dot-bracket notation with a minimum free energy of **-14.80** kcal/mol is given below.

[color by base-pairing probability | color by positional entropy | no coloring]

```
1      AAAAAAAACGAAGAAACAAAGUGACAAAGUAAAAAAGUUCUUAAAAAUUUGAUUAUAAAAAUGUCGACGACAAACGUGAUAAAAUCUAAAAAAUAAUACAAAAAUUGACAAAGUCAAAAAAGCCGAAAGUAAAAAGACACAAACACAUAUACACUAUGUA
160    AUACAUUUAACACACACUCAAACACAACAAAUUUCUGCCGUUCCUUACCCC

1      ......((((.(((((....(((.....((((..(((((......)))))..)))).....((((.(((.....))))))).....................(((((...))))).........................(((((.....))))).
160    ......................)))......)))))..)))).........
```

You can download the minimum free energy (MFE) structure in [Vienna Format| Ct Format]. You can get thermodynamic details on this structure by submitting to our RNAeval web server.

##### Results for thermodynamic ensemble prediction

The free energy of the thermodynamic ensemble is **-18.34** kcal/mol.
The frequency of the MFE structure in the ensemble is **0.32** %.
The ensemble diversity is **30.96** .
You may look at the **dot plot** containing the base pair probabilities [EPS|PDF|IMAGE CONVERTER].

The centroid secondary structure in dot-bracket notation with a minimum free energy of **-10.10** kcal/mol is given below.

[color by base-pairing probability | color by positional entropy | no coloring]

```
1      AAAAAAAACGAAGAAACAAAGUGACAAAGUAAAAAAGUUCUUAAAAAUUUGAUUAUAAAAAUGUCGACGACAAACGUGAUAAAAUCUAAAAAAUAAUACAAAAAUUGACAAAGUCAAAAAAGCCGAAAGUAAAAAGACACAAACACAUAUACACUAUGUA
160    AUACAUUUAACACACACUCAAACACAACAAAUUUCUGCCGUUCCUUACCCC

1      ......((((.(((((..................((((........))))............(((.(((.....))))))......................(((((...))))).........................(((((.....))))).
160    ..............................)))))..)))).........
```

You can download the minimum free energy (MFE) structure in [Vienna Format| Ct Format]. You can get thermodynamic details on this structure by submitting to our RNAeval web server.

##### Graphical output

You may look at the interactive drawing of the MFE structure below. If you do not see the interactive drawing and you are using Internet Explorer, please install the Adobe SVG plugin. **A note on base-pairing probabilities:** The structure below is colored by base-pairing probabilities. For unpaired regions the color denotes the probability of being unpaired.

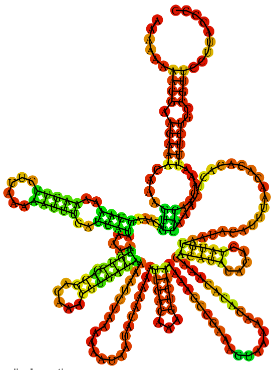

MFE secondary structure

Sequence display options
- Plain Sequence
- No Sequence

Other display options
- Base-pair probabilities
- Positional entropy
- None

0 1

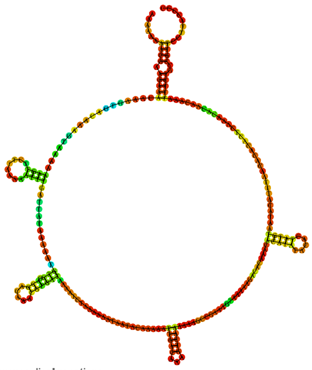

Centroid secondary structure

Sequence display options
- Plain Sequence
- No Sequence

Other display options
- Base-pair probabilities
- Positional entropy
- None

0 1

| Image description | Download options |
|---|---|
| MFE plain structure drawing | [ EPS \| PDF \| IMAGE CONVERTER \| VIEW IN FORNA ] |
| Centroid plain structure drawing | [ EPS \| PDF \| IMAGE CONVERTER \| VIEW IN FORNA ] |
| MFE structure drawing encoding base-pair probabilities | [ EPS \| PDF \| IMAGE CONVERTER \| VIEW IN FORNA ] |
| Centroid structure drawing encoding base-pair probabilities | [ EPS \| PDF \| IMAGE CONVERTER \| VIEW IN FORNA ] |
| MFE structure drawing encoding positional entropy | [ EPS \| PDF \| IMAGE CONVERTER \| VIEW IN FORNA ] |
| Centroid structure drawing encoding positional entropy | [ EPS \| PDF \| IMAGE CONVERTER \| VIEW IN FORNA ] |

Here you find a mountain plot representation of the MFE structure, the thermodynamic ensemble of RNA structures, and the centroid structure. Additionally we present the positional entropy for each position. Download as [EPS|PDF|IMAGE CONVERTER].

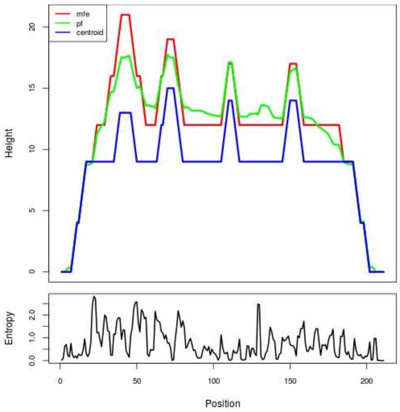


Results have been computed using RNAfold 2.6.3. An equivalent command line call would have been

```
RNAfold -p -d2 --noLP < sequence1.fa > sequence1.out
```


**RNA parameters are described in**

Mathews DH, Disney MD, Childs JL, Schroeder SJ, Zuker M, Turner DH. (2004) Incorporating chemical modification constraints into a dynamic programming algorithm for prediction of RNA secondary structure. *Proc Natl Acad Sci U S A* 101(19):7287-92.

**If you find these results helpful for your work you may want to cite:**

OPEN ACCESS OXFORD JOURNALS Gruber AR, Lorenz R, Bernhart SH, Neuböck R, Hofacker IL. **The Vienna RNA Websuite.** Nucleic Acids Research, Volume 36, Issue suppl_2, 1 July 2008, Pages W70-W74, DOI: 10.1093/nar/gkn188

Lorenz, R. and Bernhart, S.H. and Höner zu Siederdissen, C. and Tafer, H. and Flamm, C. and Stadler, P.F. and Hofacker, I.L. "ViennaRNA Package 2.0", Algorithms for Molecular Biology, 6:1 page(s): 26, 2011

### RNAfold WebServer MT_PURE_SEQ5

1 Enter Input Parameters 2 View Results

[Home|New job|Help]

##### Results for minimum free energy prediction

The optimal secondary structure in dot-bracket notation with a minimum free energy of **-55.80** kcal/mol is given below.

[color by base-pairing probability | color by positional entropy | no coloring]

```
1       GGCAGGGGGAGGUCUUGGGGCACUGGAAAGGGGCUGAGAGGUAGCUGGAGGGAGCAGAAGGGUCCAUAGGGGCAUGGGUAAGUUGGUUGAGGUCGAGGGAGUGUGGAUACAGGCACUUGGGAGGGGCAGACAGGUCAGGGAGGGUGUGGAGGGUGGAUGA
160     GGGGGCAUUGCAGGGCUUGAGAGGCACCUGGAGGGUGCCCUGGAGCAUUGGGAAGGUC

1       ..(((((.....)))))..(((((......((.(((.......(((......)))......((((....)))).........(((....((.(((..((.((....)).)).))).))...)))...))).))......))))).............
160     ...(((.((.(((.((((.((.(((((((...))))))))).)))).))).))..)))
```

You can download the minimum free energy (MFE) structure in [Vienna Format| Ct Format]. You can get thermodynamic details on this structure by submitting to our RNAeval web server.

##### Results for thermodynamic ensemble prediction

The free energy of the thermodynamic ensemble is **-59.81** kcal/mol.

The frequency of the MFE structure in the ensemble is **0.15** %.

The ensemble diversity is **67.26** .

You may look at the **dot plot** containing the base pair probabilities [EPS|PDF|IMAGE CONVERTER].

The centroid secondary structure in dot-bracket notation with a minimum free energy of **-39.10** kcal/mol is given below.

[color by base-pairing probability | color by positional entropy | no coloring]

```
1       GGCAGGGGGAGGUCUUGGGGCACUGGAAAGGGGCUGAGAGGUAGCUGGAGGGAGCAGAAGGGUCCAUAGGGGCAUGGGUAAGUUGGUUGAGGUCGAGGGAGUGUGGAUACAGGCACUUGGGAGGGGCAGACAGGUCAGGGAGGGUGUGGAGGGUGGAUGA
160     GGGGGCAUUGCAGGGCUUGAGAGGCACCUGGAGGGUGCCCUGGAGCAUUGGGAAGGUC

1       .............................................................((((((....((((......................((((((........))))))..........................)))).....))))))..
160     .......((.((..((((.((.(((((((...))))))))).))))..)).)).....
```

You can download the minimum free energy (MFE) structure in [Vienna Format| Ct Format]. You can get thermodynamic details on this structure by submitting to our RNAeval web server.

##### Graphical output

You may look at the interactive drawing of the MFE structure below. If you do not see the interactive drawing and you are using Internet Explorer, please install the Adobe SVG plugin. **A note on base-pairing probabilities:** The structure below is colored by base-pairing probabilities. For unpaired regions the color denotes the probability of being unpaired.

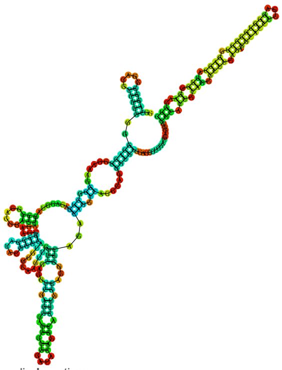

Sequence display options
- (•) Plain Sequence
- ( ) No Sequence

**MFE secondary structure**

Other display options
- (•) Base-pair probabilities
- ( ) Positional entropy
- ( ) None

0 — 1

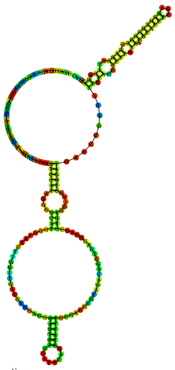

Sequence display options
- (•) Plain Sequence
- ( ) No Sequence

**Centroid secondary structure**

Other display options
- (•) Base-pair probabilities
- ( ) Positional entropy
- ( ) None

0 — 1

| Image description | Download options |
|---|---|
| MFE plain structure drawing | [ EPS \| PDF \| IMAGE CONVERTER \| VIEW IN FORNA ] |
| Centroid plain structure drawing | [ EPS \| PDF \| IMAGE CONVERTER \| VIEW IN FORNA ] |
| MFE structure drawing encoding base-pair probabilities | [ EPS \| PDF \| IMAGE CONVERTER \| VIEW IN FORNA ] |
| Centroid structure drawing encoding base-pair probabilities | [ EPS \| PDF \| IMAGE CONVERTER \| VIEW IN FORNA ] |
| MFE structure drawing encoding positional entropy | [ EPS \| PDF \| IMAGE CONVERTER \| VIEW IN FORNA ] |
| Centroid structure drawing encoding positional entropy | [ EPS \| PDF \| IMAGE CONVERTER \| VIEW IN FORNA ] |

Here you find a mountain plot representation of the MFE structure, the thermodynamic ensemble of RNA structures, and the centroid structure. Additionally we present the positional entropy for each position. Download as [EPS|PDF|IMAGE CONVERTER].

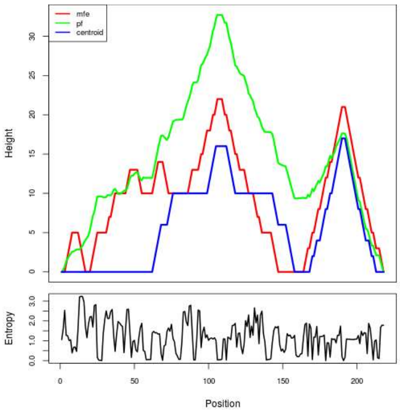


Results have been computed using RNAfold 2.6.3. An equivalent command line call would have been

`RNAfold -p -d2 --noLP < sequence1.fa > sequence1.out`


**RNA parameters are described in**

Mathews DH, Disney MD, Childs JL, Schroeder SJ, Zuker M, Turner DH. (2004) Incorporating chemical modification constraints into a dynamic programming algorithm for prediction of RNA secondary structure. *Proc Natl Acad Sci U S A* 101(19):7287-92.

**If you find these results helpful for your work you may want to cite:**

OPEN ACCESS OXFORD JOURNALS Gruber AR, Lorenz R, Bernhart SH, Neuböck R, Hofacker IL. **The Vienna RNA Websuite.** Nucleic Acids Research, Volume 36, Issue suppl_2, 1 July 2008, Pages W70-W74, DOI: 10.1093/nar/gkn188

Lorenz, R. and Bernhart, S.H. and Höner zu Siederdissen, C. and Tafer, H. and Flamm, C. and Stadler, P.F. and Hofacker, I.L. "ViennaRNA Package 2.0", Algorithms for Molecular Biology, 6:1 page(s): 26, 2011

### MT_PURE_SEQ6



##### Results for minimum free energy prediction

The optimal secondary structure in dot-bracket notation with a minimum free energy of **-65.30** kcal/mol is given below.

[color by base-pairing probability | color by positional entropy | no coloring]

```
1     UGUGUCGAAGAAAACCAGAGUAUGUGCUGAUCAGAGUGAUACCGAUACUAUUACUGAUAUUGAUUGCGAGACAUGCGGGAUGGACUUGAUCGAAGGAAAUAAGGUAAGCCAUUCCAGCGAGUGUUUCGCGGCUGCUCAGUGGGAUUUAUAUACGGUCUGU
160   GUUUGGGCUAAAAUCAAUAGAGAUAAGAUACUUGAAGAGCAAAGCAAUGAGUAAGAUUCAAUUCAAAUGUUUAGGUUGCGUCUAUUCCCCCGAUACUUGGAUCGACUACA

1     (((((((...((((((((((((((((((.(((((((((.......)))))))........(((((((((((((((((((.(((.(((...........))))))))))))).))..))))))))))))))).)))))).......))))))..)))).
160   ))))..(((....((....(((........)))...))....)))...(((((.(((.(((((.((....)).))))).))))))))..((((...)))).))))).)))
```

You can download the minimum free energy (MFE) structure in [Vienna Format| Ct Format]. You can get thermodynamic details on this structure by submitting to our RNAeval web server.

##### Results for thermodynamic ensemble prediction

The free energy of the thermodynamic ensemble is **-71.06** kcal/mol.
The frequency of the MFE structure in the ensemble is **0.01** %.
The ensemble diversity is **72.88** .
You may look at the **dot plot** containing the base pair probabilities [EPS|PDF|IMAGE CONVERTER].

The centroid secondary structure in dot-bracket notation with a minimum free energy of **-38.90** kcal/mol is given below.
[color by base-pairing probability | color by positional entropy | no coloring]

```
1     UGUGUCGAAGAAAACCAGAGUAUGUGCUGAUCAGAGUGAUACCGAUACUAUUACUGAUAUUGAUUGCGAGACAUGCGGGAUGGACUUGAUCGAAGGAAAUAAGGUAAGCCAUUCCAGCGAGUGUUUCGCGGCUGCUCAGUGGGAUUUAUAUACGGUCUGU
160   GUUUGGGCUAAAAUCAAUAGAGAUAAGAUACUUGAAGAGCAAAGCAAUGAGUAAGAUUCAAUUCAAAUGUUUAGGUUGCGUCUAUUCCCCCGAUACUUGGAUCGACUACA

1     .(((((((...........(((....((((.(((((((((.......))))))))........(((((((((((((((((((.(((.(((...........))))))))))))).))..))))))))))).))).))))...............((((.(
160   (...((.......))..)).))))............................((..(((..............)))..))...........)))....)))))).)).
```

You can download the minimum free energy (MFE) structure in [Vienna Format| Ct Format]. You can get thermodynamic details on this structure by submitting to our RNAeval web server.

##### Graphical output

You may look at the interactive drawing of the MFE structure below. If you do not see the interactive drawing and you are using Internet Explorer, please install the Adobe SVG plugin. **A note on base-pairing probabilities:** The structure below is colored by base-pairing probabilities. For unpaired regions the color denotes the probability of being unpaired.

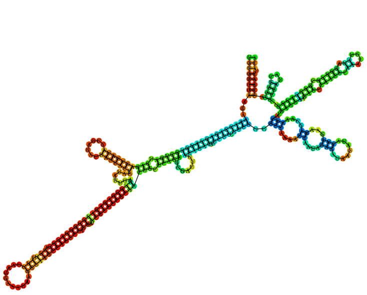

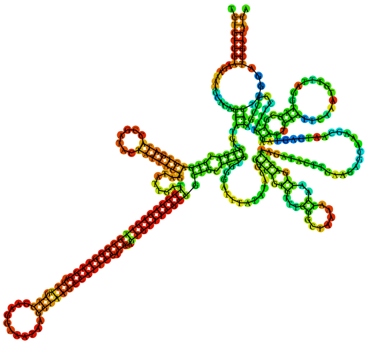

Sequence display options
- (•) Plain Sequence
- ( ) No Sequence

**MFE secondary structure**

Other display options
- (•) Base-pair probabilities
- ( ) Positional entropy
- ( ) None

0 1

Sequence display options
- (•) Plain Sequence
- ( ) No Sequence

**Centroid secondary structure**

Other display options
- (•) Base-pair probabilities
- ( ) Positional entropy
- ( ) None

0 1

| **Image description** | **Download options** |
|---|---|
| MFE plain structure drawing | [ EPS \| PDF \| IMAGE CONVERTER \| VIEW IN FORNA ] |
| Centroid plain structure drawing | [ EPS \| PDF \| IMAGE CONVERTER \| VIEW IN FORNA ] |
| MFE structure drawing encoding base-pair probabilities | [ EPS \| PDF \| IMAGE CONVERTER \| VIEW IN FORNA ] |
| Centroid structure drawing encoding base-pair probabilities | [ EPS \| PDF \| IMAGE CONVERTER \| VIEW IN FORNA ] |
| MFE structure drawing encoding positional entropy | [ EPS \| PDF \| IMAGE CONVERTER \| VIEW IN FORNA ] |
| Centroid structure drawing encoding positional entropy | [ EPS \| PDF \| IMAGE CONVERTER \| VIEW IN FORNA ] |

Here you find a mountain plot representation of the MFE structure, the thermodynamic ensemble of RNA structures, and the centroid structure. Additionally we present the positional entropy for each position. Download as [EPS|PDF|IMAGE CONVERTER].

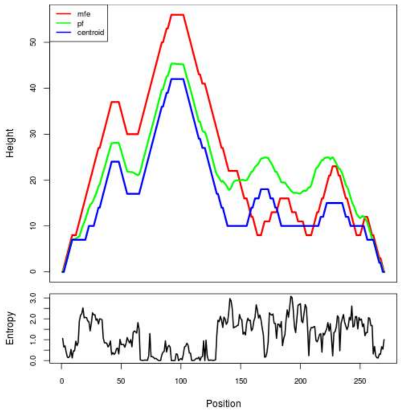


Results have been computed using RNAfold 2.6.3. An equivalent command line call would have been

```
RNAfold -p -d2 --noLP < sequence1.fa > sequence1.out
```

**RNA parameters are described in**

Mathews DH, Disney MD, Childs JL, Schroeder SJ, Zuker M, Turner DH. (2004) Incorporating chemical modification constraints into a dynamic programming algorithm for prediction of RNA secondary structure. *Proc Natl Acad Sci U S A* 101(19):7287-92.

**If you find these results helpful for your work you may want to cite:**

OPEN ACCESS OXFORD JOURNALS Gruber AR, Lorenz R, Bernhart SH, Neuböck R, Hofacker IL. **The Vienna RNA Websuite.** Nucleic Acids Research, Volume 36, Issue suppl_2, 1 July 2008, Pages W70–W74, DOI: 10.1093/nar/gkn188

Lorenz, R. and Bernhart, S.H. and Höner zu Siederdissen, C. and Tafer, H. and Flamm, C. and Stadler, P.F. and Hofacker, I.L. "ViennaRNA Package 2.0", Algorithms for Molecular Biology, 6:1 page(s): 26, 2011

##### Results for minimum free energy prediction

The optimal secondary structure in dot-bracket notation with a minimum free energy of **-24.70** kcal/mol is given below.
[color by base-pairing probability | color by positional entropy | no coloring]

```
1      UGCCACCUGUACGACACAGCAUACCGGCAUUACUGAGCCGGUAUCCUUCCGCCGUGACCUCCUCAACUAAGCCCACCCACGCAAAUGACAGGCCAGCCCAAAUCAACAUUUGCCUCCCAAUCAUAAUAGGAUCCUUUUUAAUAGCGUUUCUUCCAUACCC
160    AUCCCCAUACCCAUCCCCACUCCCAUACCCACCCCCAUCCCCA

1      .....((((....(((.(((((((((........)))))))).......)).)))......................(((((((..................)))))))...........)))))..............................
160    .........................................
```

You can download the minimum free energy (MFE) structure in [Vienna Format| Ct Format]. You can get thermodynamic details on this structure by submitting to our RNAeval web server.

##### Results for thermodynamic ensemble prediction

The free energy of the thermodynamic ensemble is **-27.72** kcal/mol.
The frequency of the MFE structure in the ensemble is **0.75** %.
The ensemble diversity is **35.16** .
You may look at the **dot plot** containing the base pair probabilities [EPS|PDF|IMAGE CONVERTER].

The centroid secondary structure in dot-bracket notation with a minimum free energy of **-13.10** kcal/mol is given below.
[color by base-pairing probability | color by positional entropy | no coloring]

```
1      UGCCACCUGUACGACACAGCAUACCGGCAUUACUGAGCCGGUAUCCUUCCGCCGUGACCUCCUCAACUAAGCCCACCCACGCAAAUGACAGGCCAGCCCAAAUCAACAUUUGCCUCCCAAUCAUAAUAGGAUCCUUUUUAAUAGCGUUUCUUCCAUACCC
160    AUCCCCAUACCCAUCCCCACUCCCAUACCCACCCCCAUCCCCA

1      ....................((((((((........))))))))...............................................................................................................
160    .........................................
```

You can download the minimum free energy (MFE) structure in [Vienna Format| Ct Format]. You can get thermodynamic details on this structure by submitting to our RNAeval web server.

##### Graphical output

You may look at the interactive drawing of the MFE structure below. If you do not see the interactive drawing and you are using Internet Explorer, please install the Adobe SVG plugin. **A note on base-pairing probabilities:** The structure below is colored by base-pairing probabilities. For unpaired regions the color denotes the probability of being unpaired.

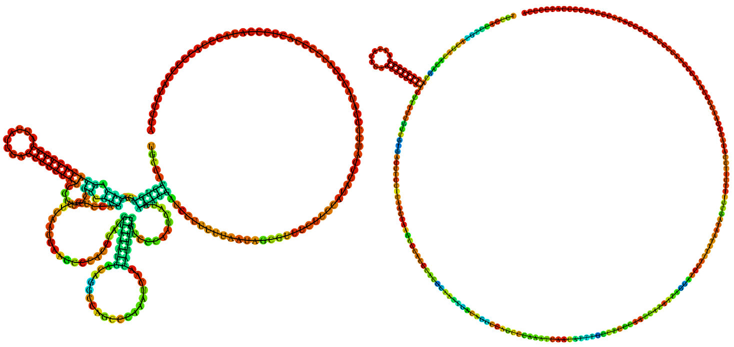

| Sequence display options | MFE secondary structure | Sequence display options | Centroid secondary structure |
|---|---|---|---|
| ◉ Plain Sequence | | ◉ Plain Sequence | |
| ○ No Sequence | | ○ No Sequence | |
| Other display options | | Other display options | |
| ◉ Base-pair probabilities | 0 — 1 | ◉ Base-pair probabilities | 0 — 1 |
| ○ Positional entropy | | ○ Positional entropy | |
| ○ None | | ○ None | |

| Image description | Download options |
|---|---|
| MFE plain structure drawing | [ EPS \| PDF \| IMAGE CONVERTER \| VIEW IN FORNA ] |
| Centroid plain structure drawing | [ EPS \| PDF \| IMAGE CONVERTER \| VIEW IN FORNA ] |
| MFE structure drawing encoding base-pair probabilities | [ EPS \| PDF \| IMAGE CONVERTER \| VIEW IN FORNA ] |
| Centroid structure drawing encoding base-pair probabilities | [ EPS \| PDF \| IMAGE CONVERTER \| VIEW IN FORNA ] |
| MFE structure drawing encoding positional entropy | [ EPS \| PDF \| IMAGE CONVERTER \| VIEW IN FORNA ] |
| Centroid structure drawing encoding positional entropy | [ EPS \| PDF \| IMAGE CONVERTER \| VIEW IN FORNA ] |

Here you find a mountain plot representation of the MFE structure, the thermodynamic ensemble of RNA structures, and the centroid structure. Additionally we present the positional entropy for each position. Download as [EPS|PDF|IMAGE CONVERTER].

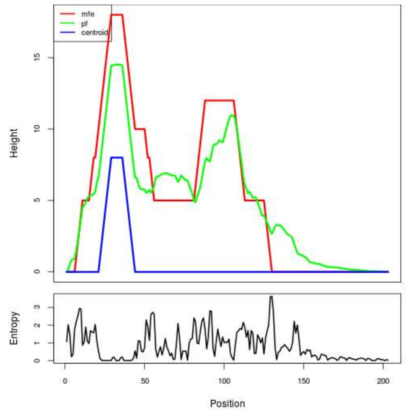


Results have been computed using RNAfold 2.6.3. An equivalent command line call would have been

```
RNAfold -p -d2 --noLP < sequence1.fa > sequence1.out
```

**RNA parameters are described in**
Mathews DH, Disney MD, Childs JL, Schroeder SJ, Zuker M, Turner DH. (2004) Incorporating chemical modification constraints into a dynamic programming algorithm for prediction of RNA secondary structure. *Proc Natl Acad Sci U S A* 101(19):7287-92.

**If you find these results helpful for your work you may want to cite:**

OPEN ACCESS OXFORD JOURNALS Gruber AR, Lorenz R, Bernhart SH, Neuböck R, Hofacker IL.
**The Vienna RNA Websuite.** Nucleic Acids Research, Volume 36, Issue suppl_2, 1 July 2008, Pages W70-W74, DOI: 10.1093/nar/gkn188

Lorenz, R. and Bernhart, S.H. and Höner zu Siederdissen, C. and Tafer, H. and Flamm, C. and Stadler, P.F. and Hofacker, I.L. "ViennaRNA Package 2.0", Algorithms for Molecular Biology, 6:1 page(s): 26, 2011

### MT_PURE_SEQ8



#### Results for minimum free energy prediction

The optimal secondary structure in dot-bracket notation with a minimum free energy of **-54.40** kcal/mol is given below.

[color by base-pairing probability | color by positional entropy | no coloring]

```
1      CUUGGUGUUAAAUAUCUAAUAAUUAAUACUCCUGAUCCUCCCACUCCAUUAUUUGUACCUGCAAAAGCUAUACGACCAUUUCCUCCUCCACCACCACCAAAACCAUUUGUGGCAUUUGUACCUUGUAAACCUGCACCUUCGACAGCUCCUUGUCCUGAAU
160    UAUCACAACCUCCAAGUGAUCUAGUACUUGUUGUAUUAGCUUCGUGUGUACCACCUCCUCCCCCACCUGCAUAACAUUUUGUUGUACCGCUUAUACUAAAAUUAGUUCCUAAUCCGCCUCUUCCUCCCUUUGCAAAAUCAUUAGUACCUGACUGAAAUCC
320    UGUUGUAUUAUGUCCCCCUCCUCCAUUAGAACCAUAACCACUUACUCCGUUAUAUGUAGCUCCAUUAUUUCCCUGACCUGCAGUUCCACUUCCAAAGUUUGUAUUUUGUUGUCCAGUACCACCACCACUACCUCCAUUUGAUGCUACACUUGUUAAAGCU
480    CCUGUACUUCCGUUACCU

1      ...((((((((.(((...))).))))))))............................((((..((((....((........))..................((..((((((((.......((((....))))......(((((....)))))......
160    .(((((.........)))))(((((((.....)))))))....(((.((((.................((((((((...))))))...))....((((....)))).......................(((((.....(((((.....)))))....
320    ..)))))..........................(((((..........))))).(((((.((............)).)))))..........((((((....))))))......))))))).................)))))))))..))....))))
480    ..))))............
```

You can download the minimum free energy (MFE) structure in [Vienna Format| Ct Format]. You can get thermodynamic details on this structure by submitting to our RNAeval web server.

#### Results for thermodynamic ensemble prediction

The free energy of the thermodynamic ensemble is **-64.47** kcal/mol.

The frequency of the MFE structure in the ensemble is **0.00** %.

The ensemble diversity is **127.20**.

You may look at the **dot plot** containing the base pair probabilities [EPS|PDF|IMAGE CONVERTER].

The centroid secondary structure in dot-bracket notation with a minimum free energy of **-18.80** kcal/mol is given below.

[color by base-pairing probability | color by positional entropy | no coloring]

```
1      CUUGGUGUUAAAUAUCUAAUAAUUAAUACUCCUGAUCCUCCCACUCCAUUAUUUGUACCUGCAAAAGCUAUACGACCAUUUCCUCCUCCACCACCACCAAAACCAUUUGUGGCAUUUGUACCUUGUAAACCUGCACCUUCGACAGCUCCUUGUCCUGAAU
160    UAUCACAACCUCCAAGUGAUCUAGUACUUGUUGUAUUAGCUUCGUGUGUACCACCUCCUCCCCCACCUGCAUAACAUUUUGUUGUACCGCUUAUACUAAAAUUAGUUCCUAAUCCGCCUCUUCCUCCCUUUGCAAAAUCAUUAGUACCUGACUGAAAUCC
320    UGUUGUAUUAUGUCCCCCUCCUCCAUUAGAACCAUAACCACUUACUCCGUUAUAUGUAGCUCCAUUAUUUCCCUGACCUGCAGUUCCACUUCCAAAGUUUGUAUUUUGUUGUCCAGUACCACCACCACUACCUCCAUUUGAUGCUACACUUGUUAAAGCU
480    CCUGUACUUCCGUUACCU

1      ...((((((((.((.....)).))))))))...............................................................(((.............)))..........((((....))))......((((......))))......
160    .(((((.........)))))(((((((.....))))))).....................................................((((....)))).......................................................
320    .................................(((((..........)))))................................((((((....))))))..........................................................
480    ..................
```

You can download the minimum free energy (MFE) structure in [Vienna Format| Ct Format]. You can get thermodynamic details on this structure by submitting to our RNAeval web server.

#### Graphical output

You may look at the interactive drawing of the MFE structure below. If you do not see the interactive drawing and you are using Internet Explorer, please install the Adobe SVG plugin. **A note on base-pairing probabilities:** The structure below is colored by base-pairing probabilities. For unpaired regions the color denotes the probability of being unpaired.

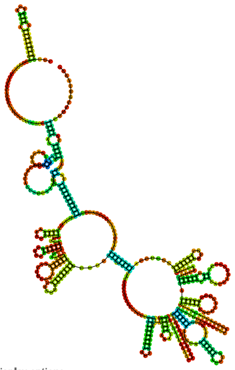

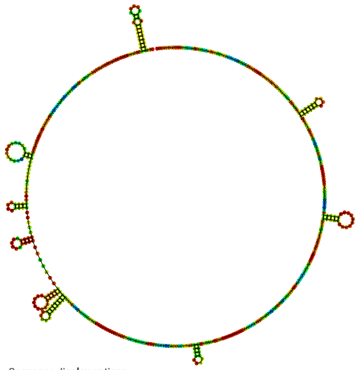

Sequence display options
- Plain Sequence
- No Sequence

MFE secondary structure

Other display options
- Base-pair probabilities
- Positional entropy
- None

0 1

Sequence display options
- Plain Sequence
- No Sequence

Centroid secondary structure

Other display options
- Base-pair probabilities
- Positional entropy
- None

0 1

| Image description | Download options |
| --- | --- |
| MFE plain structure drawing | [ EPS \| PDF \| IMAGE CONVERTER \| VIEW IN FORNA ] |
| Centroid plain structure drawing | [ EPS \| PDF \| IMAGE CONVERTER \| VIEW IN FORNA ] |
| MFE structure drawing encoding base-pair probabilities | [ EPS \| PDF \| IMAGE CONVERTER \| VIEW IN FORNA ] |
| Centroid structure drawing encoding base-pair probabilities | [ EPS \| PDF \| IMAGE CONVERTER \| VIEW IN FORNA ] |
| MFE structure drawing encoding positional entropy | [ EPS \| PDF \| IMAGE CONVERTER \| VIEW IN FORNA ] |
| Centroid structure drawing encoding positional entropy | [ EPS \| PDF \| IMAGE CONVERTER \| VIEW IN FORNA ] |

Here you find a mountain plot representation of the MFE structure, the thermodynamic ensemble of RNA structures, and the centroid structure. Additionally we present the positional entropy for each position. Download as [EPS|PDF|IMAGE CONVERTER].

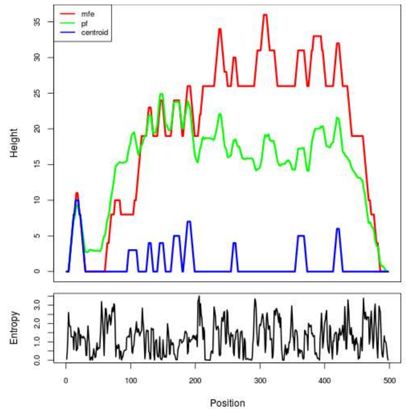


Results have been computed using RNAfold 2.6.3. An equivalent command line call would have been

```
RNAfold -p -d2 --noLP < sequence1.fa > sequence1.out
```

**RNA parameters are described in**

Mathews DH, Disney MD, Childs JL, Schroeder SJ, Zuker M, Turner DH. (2004) Incorporating chemical modification constraints into a dynamic programming algorithm for prediction of RNA secondary structure. *Proc Natl Acad Sci U S A* 101(19):7287-92.

**If you find these results helpful for your work you may want to cite:**

OPEN ACCESS OXFORD JOURNALS Gruber AR, Lorenz R, Bernhart SH, Neuböck R, Hofacker IL. **The Vienna RNA Websuite.** Nucleic Acids Research, Volume 36, Issue suppl_2, 1 July 2008, Pages W70-W74, DOI: 10.1093/nar/gkn188

Lorenz, R. and Bernhart, S.H. and Höner zu Siederdissen, C. and Tafer, H. and Flamm, C. and Stadler, P.F. and Hofacker, I.L. "ViennaRNA Package 2.0", Algorithms for Molecular Biology, 6:1 page(s): 26, 2011

RNAfold WebServer

### MT_PURE_SEQ9

1 Enter Input Parameters 2 View Results

[Home|New job|Help]

#### Results for minimum free energy prediction

The optimal secondary structure in dot-bracket notation with a minimum free energy of **-26.10** kcal/mol is given below.

[color by base-pairing probability | color by positional entropy | no coloring]

```
1     CGCCACUCACCGACCACCGCGGAUAAUACCCCUCCACCCGAUCCACCGACCCACCCCCCGUACCCUCCACAUAAUGCCCCCUCGCCGGCGCCAACAAAUACUGCGCCCCAAAAAACUCCUCAUUAUCCAUGCACUUCGUAUCCACCGUGUCCGCAUACAU
160   CCCAGCCCCACGCUGAUCAUCCGACCCCCGAAACCCCGACCCAUACCGAUACUGCCCCAUCACCACCUGAUCCCCCUGCG

1     (((...............((((((.............((......)).........................((......)).(((((...........))))).....................((((.....))))......)))))))......
160   ..((((.....))))......((.....))......((........))..........((((.....))))......)))
```

You can download the minimum free energy (MFE) structure in [Vienna Format| Ct Format]. You can get thermodynamic details on this structure by submitting to our RNAeval web server.

#### Results for thermodynamic ensemble prediction

The free energy of the thermodynamic ensemble is **-30.07** kcal/mol.
The frequency of the MFE structure in the ensemble is **0.16** %.
The ensemble diversity is **37.09** .
You may look at the **dot plot** containing the base pair probabilities [EPS|PDF|IMAGE CONVERTER].

The centroid secondary structure in dot-bracket notation with a minimum free energy of **-12.40** kcal/mol is given below.

[color by base-pairing probability | color by positional entropy | no coloring]

```
1     CGCCACUCACCGACCACCGCGGAUAAUACCCCUCCACCCGAUCCACCGACCCACCCCCCGUACCCUCCACAUAAUGCCCCCUCGCCGGCGCCAACAAAUACUGCGCCCCAAAAAACUCCUCAUUAUCCAUGCACUUCGUAUCCACCGUGUCCGCAUACAU
160   CCCAGCCCCACGCUGAUCAUCCGACCCCCGAAACCCCGACCCAUACCGAUACUGCCCCAUCACCACCUGAUCCCCCUGCG

1     ......................................((......))..........................((......)).(((((...........))))).......................................................
160   ..((((.....))))..........................................((((.....))))........
```

You can download the minimum free energy (MFE) structure in [Vienna Format| Ct Format]. You can get thermodynamic details on this structure by submitting to our RNAeval web server.

#### Graphical output

You may look at the interactive drawing of the MFE structure below. If you do not see the interactive drawing and you are using Internet Explorer, please install the Adobe SVG plugin. **A note on base-pairing probabilities:** The structure below is colored by base-pairing probabilities. For unpaired regions the color denotes the probability of being unpaired.

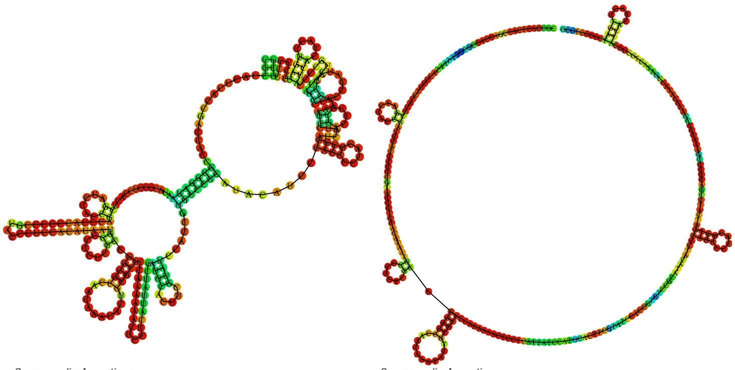

| Sequence display options | MFE secondary structure | Sequence display options | Centroid secondary structure |
|---|---|---|---|
| ◉ Plain Sequence | | ◉ Plain Sequence | |
| ○ No Sequence | | ○ No Sequence | |
| Other display options | | Other display options | |
| ◉ Base-pair probabilities | 0 — 1 | ◉ Base-pair probabilities | 0 — 1 |
| ○ Positional entropy | | ○ Positional entropy | |
| ○ None | | ○ None | |

| Image description | Download options |
|---|---|
| MFE plain structure drawing | [ EPS \| PDF \| IMAGE CONVERTER \| VIEW IN FORNA ] |
| Centroid plain structure drawing | [ EPS \| PDF \| IMAGE CONVERTER \| VIEW IN FORNA ] |
| MFE structure drawing encoding base-pair probabilities | [ EPS \| PDF \| IMAGE CONVERTER \| VIEW IN FORNA ] |
| Centroid structure drawing encoding base-pair probabilities | [ EPS \| PDF \| IMAGE CONVERTER \| VIEW IN FORNA ] |
| MFE structure drawing encoding positional entropy | [ EPS \| PDF \| IMAGE CONVERTER \| VIEW IN FORNA ] |
| Centroid structure drawing encoding positional entropy | [ EPS \| PDF \| IMAGE CONVERTER \| VIEW IN FORNA ] |

Here you find a mountain plot representation of the MFE structure, the thermodynamic ensemble of RNA structures, and the centroid structure. Additionally we present the positional entropy for each position. Download as [EPS|PDF|IMAGE CONVERTER].

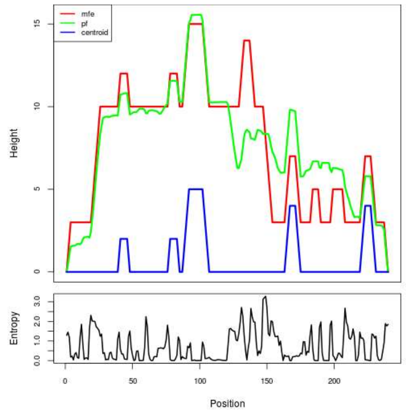


Results have been computed using RNAfold 2.6.3. An equivalent command line call would have been

```
RNAfold -p -d2 --noLP < sequence1.fa > sequence1.out
```

**RNA parameters are described in**


Mathews DH, Disney MD, Childs JL, Schroeder SJ, Zuker M, Turner DH. (2004) Incorporating chemical modification constraints into a dynamic programming algorithm for prediction of RNA secondary structure. *Proc Natl Acad Sci U S A* 101(19):7287-92.


**If you find these results helpful for your work you may want to cite:**


OPEN ACCESS OXFORD JOURNALS Gruber AR, Lorenz R, Bernhart SH, Neuböck R, Hofacker IL.
**The Vienna RNA Websuite.** Nucleic Acids Research, Volume 36, Issue suppl_2, 1 July 2008, Pages W70-W74, DOI: 10.1093/nar/gkn188

Lorenz, R. and Bernhart, S.H. and Höner zu Siederdissen, C. and Tafer, H. and Flamm, C. and Stadler, P.F. and Hofacker, I.L. "ViennaRNA Package 2.0", Algorithms for Molecular Biology, 6:1 page(s): 26, 2011

### RNAfold WebServer MT_PURE_SEQ10

1 Enter Input Parameters 2 View Results

[Home|New job|Help]

##### Results for minimum free energy prediction

The optimal secondary structure in dot-bracket notation with a minimum free energy of **-31.80** kcal/mol is given below.

[color by base-pairing probability | color by positional entropy | no coloring]

```
1      GGGGUAAGGAACGGCAGAAAUUUGUUGUGUUUGAGUGUGUGUUAAAUGUAUUACAUAGUGUAUAUGUGUUUGUGUCUUUUUACUUUCGGCUUUUUUGACUUUGUCAAUUUUUGUAUUAUUUUUUAGAUUUUAUCACGUUUGUCGUCGACAUUUUUAUAAU
160    CAAAUUUUUAAGAACUUUUUUACUUUGUCACUUUGUUUCUUCGUUUUUUUUUCGUUCUUUUAUUUUUAUUUGAUUUUUU

1      .(((.((((((((.((((((.....(((...((((((((((..........)))))......(((((.....(((...........((....(((((...)))))).....))..........)))).....))))).((((...))))((((((.((.
160    .....)).))))))......)))))...)))...))))))))))))))).)))..........................
```

You can download the minimum free energy (MFE) structure in [Vienna Format| Ct Format]. You can get thermodynamic details on this structure by submitting to our RNAeval web server.

##### Results for thermodynamic ensemble prediction

The free energy of the thermodynamic ensemble is **-35.55** kcal/mol.
The frequency of the MFE structure in the ensemble is **0.23** %.
The ensemble diversity is **67.59** .
You may look at the **dot plot** containing the base pair probabilities [EPS|PDF|IMAGE CONVERTER].

The centroid secondary structure in dot-bracket notation with a minimum free energy of **-19.50** kcal/mol is given below.

[color by base-pairing probability | color by positional entropy | no coloring]

```
1      GGGGUAAGGAACGGCAGAAAUUUGUUGUGUUUGAGUGUGUGUUAAAUGUAUUACAUAGUGUAUAUGUGUUUGUGUCUUUUUACUUUCGGCUUUUUUGACUUUGUCAAUUUUUGUAUUAUUUUUUAGAUUUUAUCACGUUUGUCGUCGACAUUUUUAUAAU
160    CAAAUUUUUAAGAACUUUUUUACUUUGUCACUUUGUUUCUUCGUUUUUUUUUCGUUCUUUUAUUUUUAUUUGAUUUUUU

1      .....((((((((.((((((.....(((...((((.................................((((...........((....(((((...))))).....))..........))))...........(((...)))..........
160    ....................))))...)))...)))))))))))))))..........................
```

You can download the minimum free energy (MFE) structure in [Vienna Format| Ct Format]. You can get thermodynamic details on this structure by submitting to our RNAeval web server.

##### Graphical output

You may look at the interactive drawing of the MFE structure below. If you do not see the interactive drawing and you are using Internet Explorer, please install the Adobe SVG plugin. **A note on base-pairing probabilities:** The structure below is colored by base-pairing probabilities. For unpaired regions the color denotes the probability of being unpaired.

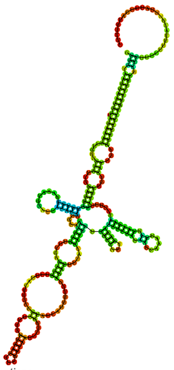

Sequence display options
- Plain Sequence
- No Sequence

MFE secondary structure

Other display options
- Base-pair probabilities
- Positional entropy
- None

0 1

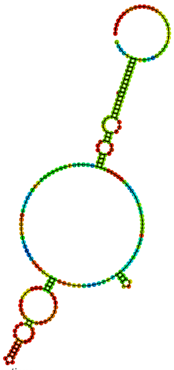

Sequence display options
- Plain Sequence
- No Sequence

Centroid secondary structure

Other display options
- Base-pair probabilities
- Positional entropy
- None

0 1

| Image description | Download options |
|---|---|
| MFE plain structure drawing | [ EPS \| PDF \| IMAGE CONVERTER \| VIEW IN FORNA ] |
| Centroid plain structure drawing | [ EPS \| PDF \| IMAGE CONVERTER \| VIEW IN FORNA ] |
| MFE structure drawing encoding base-pair probabilities | [ EPS \| PDF \| IMAGE CONVERTER \| VIEW IN FORNA ] |
| Centroid structure drawing encoding base-pair probabilities | [ EPS \| PDF \| IMAGE CONVERTER \| VIEW IN FORNA ] |
| MFE structure drawing encoding positional entropy | [ EPS \| PDF \| IMAGE CONVERTER \| VIEW IN FORNA ] |
| Centroid structure drawing encoding positional entropy | [ EPS \| PDF \| IMAGE CONVERTER \| VIEW IN FORNA ] |

Here you find a mountain plot representation of the MFE structure, the thermodynamic ensemble of RNA structures, and the centroid structure. Additionally we present the positional entropy for each position. Download as [EPS|PDF|IMAGE CONVERTER].

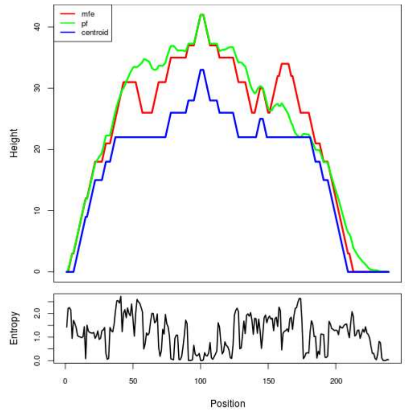


Results have been computed using RNAfold 2.6.3. An equivalent command line call would have been

```
RNAfold -p -d2 --noLP < sequence1.fa > sequence1.out
```

**RNA parameters are described in**

Mathews DH, Disney MD, Childs JL, Schroeder SJ, Zuker M, Turner DH. (2004) Incorporating chemical modification constraints into a dynamic programming algorithm for prediction of RNA secondary structure. *Proc Natl Acad Sci U S A* 101(19):7287-92.

**If you find these results helpful for your work you may want to cite:**


OPEN ACCESS OXFORD JOURNALS Gruber AR, Lorenz R, Bernhart SH, Neuböck R, Hofacker IL.
**The Vienna RNA Websuite.** Nucleic Acids Research, Volume 36, Issue suppl_2, 1 July 2008, Pages W70-W74, DOI: 10.1093/nar/gkn188

Lorenz, R. and Bernhart, S.H. and Höner zu Siederdissen, C. and Tafer, H. and Flamm, C. and Stadler, P.F. and Hofacker, I.L. "ViennaRNA Package 2.0", Algorithms for Molecular Biology, 6:1 page(s): 26, 2011

### RNAfold WebServer MT_PURE_SEQ11

1 Enter Input Parameters 2 View Results

[Home|New job|Help]

##### Results for minimum free energy prediction

The optimal secondary structure in dot-bracket notation with a minimum free energy of **-24.70** kcal/mol is given below.

[color by base-pairing probability | color by positional entropy | no coloring]

```
1      UGCCACCUGUACGACACAGCAUACCGGCAUUACUGAGCCGGUAUCCUUCCGCCGUGACCUCCUCAACUAAGCCCACCCACGCAAAUGACAGGCCAGCCCAAAUCAACAUUUGCCUCCCAAUCAUAAUAGGAUCCUUUUUAAUAGCGUUUCUUCCAUACCC
160    AUCCCCAUACCCAUCCCCACUCCCAUACCCAUCCCCAUACCCACCCCCAUCCCCAUACCCAUG

1      .....((((....(((.(((((((((........))))))))......)).)))......................(((((((..................)))))))...........))))).............................
160    ...............................................................
```

You can download the minimum free energy (MFE) structure in [Vienna Format| Ct Format]. You can get thermodynamic details on this structure by submitting to our RNAeval web server.

##### Results for thermodynamic ensemble prediction

The free energy of the thermodynamic ensemble is **-27.82** kcal/mol.
The frequency of the MFE structure in the ensemble is **0.63** %.
The ensemble diversity is **35.83** .
You may look at the **dot plot** containing the base pair probabilities [EPS|PDF|IMAGE CONVERTER].

The centroid secondary structure in dot-bracket notation with a minimum free energy of **-13.10** kcal/mol is given below.

[color by base-pairing probability | color by positional entropy | no coloring]

```
1      UGCCACCUGUACGACACAGCAUACCGGCAUUACUGAGCCGGUAUCCUUCCGCCGUGACCUCCUCAACUAAGCCCACCCACGCAAAUGACAGGCCAGCCCAAAUCAACAUUUGCCUCCCAAUCAUAAUAGGAUCCUUUUUAAUAGCGUUUCUUCCAUACCC
160    AUCCCCAUACCCAUCCCCACUCCCAUACCCAUCCCCAUACCCACCCCCAUCCCCAUACCCAUG

1      ...................((((((((........))))))))....................................................................................................................
160    ...............................................................
```

You can download the minimum free energy (MFE) structure in [Vienna Format| Ct Format]. You can get thermodynamic details on this structure by submitting to our RNAeval web server.

##### Graphical output

You may look at the interactive drawing of the MFE structure below. If you do not see the interactive drawing and you are using Internet Explorer, please install the Adobe SVG plugin. **A note on base-pairing probabilities:** The structure below is colored by base-pairing probabilities. For unpaired regions the color denotes the probability of being unpaired.

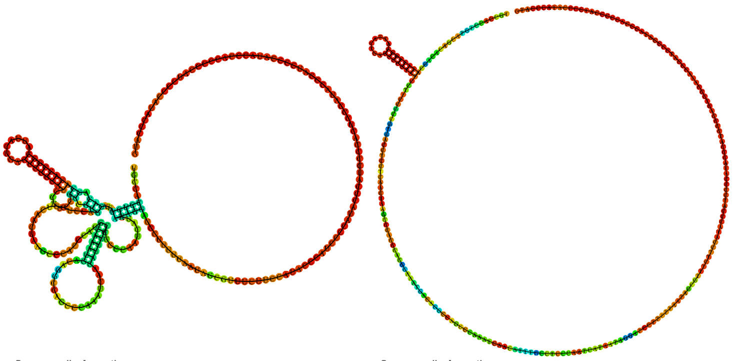

| Sequence display options | MFE secondary structure | Sequence display options | Centroid secondary structure |
|---|---|---|---|
| ◉ Plain Sequence | | ◉ Plain Sequence | |
| ○ No Sequence | | ○ No Sequence | |
| Other display options | | Other display options | |
| ◉ Base-pair probabilities | 0 – 1 | ◉ Base-pair probabilities | 0 – 1 |
| ○ Positional entropy | | ○ Positional entropy | |
| ○ None | | ○ None | |

| Image description | Download options |
|---|---|
| MFE plain structure drawing | [ EPS \| PDF \| IMAGE CONVERTER \| VIEW IN FORNA ] |
| Centroid plain structure drawing | [ EPS \| PDF \| IMAGE CONVERTER \| VIEW IN FORNA ] |
| MFE structure drawing encoding base-pair probabilities | [ EPS \| PDF \| IMAGE CONVERTER \| VIEW IN FORNA ] |
| Centroid structure drawing encoding base-pair probabilities | [ EPS \| PDF \| IMAGE CONVERTER \| VIEW IN FORNA ] |
| MFE structure drawing encoding positional entropy | [ EPS \| PDF \| IMAGE CONVERTER \| VIEW IN FORNA ] |
| Centroid structure drawing encoding positional entropy | [ EPS \| PDF \| IMAGE CONVERTER \| VIEW IN FORNA ] |

Here you find a mountain plot representation of the MFE structure, the thermodynamic ensemble of RNA structures, and the centroid structure. Additionally we present the positional entropy for each position. Download as [EPS|PDF|IMAGE CONVERTER].

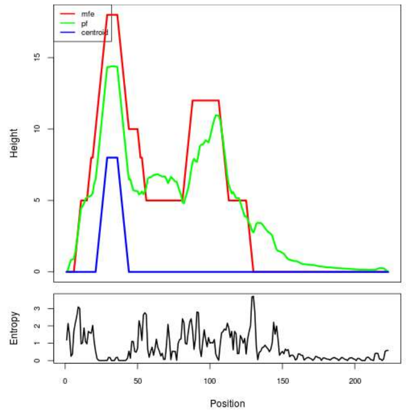


Results have been computed using RNAfold 2.6.3. An equivalent command line call would have been

```
RNAfold -p -d2 --noLP < sequence1.fa > sequence1.out
```

**RNA parameters are described in**

Mathews DH, Disney MD, Childs JL, Schroeder SJ, Zuker M, Turner DH. (2004) Incorporating chemical modification constraints into a dynamic programming algorithm for prediction of RNA secondary structure. *Proc Natl Acad Sci U S A* 101(19):7287-92.

**If you find these results helpful for your work you may want to cite:**


OPEN ACCESS OXFORD JOURNALS Gruber AR, Lorenz R, Bernhart SH, Neuböck R, Hofacker IL.
**The Vienna RNA Websuite.** Nucleic Acids Research, Volume 36, Issue suppl_2, 1 July 2008, Pages W70-W74, DOI: 10.1093/nar/gkn188

Lorenz, R. and Bernhart, S.H. and Höner zu Siederdissen, C. and Tafer, H. and Flamm, C. and Stadler, P.F. and Hofacker, I.L. "ViennaRNA Package 2.0", Algorithms for Molecular Biology, 6:1 page(s): 26, 2011

##### Results for minimum free energy prediction

The optimal secondary structure in dot-bracket notation with a minimum free energy of **-57.90** kcal/mol is given below.

[color by base-pairing probability | color by positional entropy | no coloring]

```
1     GGCAGGGGGAGUUCUUGGGGCACUGGAAAGGGGCUGAGAGGUAGCUGGAGGGAGCAGAAGGGUCCAUAGGGGCAUGGGUAAGUUGGUUGAGGUCGAGGGAGUGUGGAUACAGGCACUUGGGAGGGGCAGACAGGUCAGGGAGGGUGUGGAGGGUGGAUGA
160   GGGGGCAUUGCAGGGCUUGAGAGGCACCUGGAGGGUGCCCUGGAGCAUUGGGAAGGUC

1     ..........(((((.((((((((....(((.(((...((((..(((.((...((......((((((....((((.......(((.(((..(((....((((((........))))))......)))...)))..)))......))))......))))))..
160   ....)).)).))).))))....))).)))....)))))))))))))...........
```

You can download the minimum free energy (MFE) structure in [Vienna Format| Ct Format]. You can get thermodynamic details on this structure by submitting to our RNAeval web server.

##### Results for thermodynamic ensemble prediction

The free energy of the thermodynamic ensemble is **-61.48** kcal/mol.
The frequency of the MFE structure in the ensemble is **0.30** %.
The ensemble diversity is **39.42** .
You may look at the **dot plot** containing the base pair probabilities [EPS|PDF|IMAGE CONVERTER].

The centroid secondary structure in dot-bracket notation with a minimum free energy of **-52.50** kcal/mol is given below.

[color by base-pairing probability | color by positional entropy | no coloring]

```
1     GGCAGGGGGAGUUCUUGGGGCACUGGAAAGGGGCUGAGAGGUAGCUGGAGGGAGCAGAAGGGUCCAUAGGGGCAUGGGUAAGUUGGUUGAGGUCGAGGGAGUGUGGAUACAGGCACUUGGGAGGGGCAGACAGGUCAGGGAGGGUGUGGAGGGUGGAUGA
160   GGGGGCAUUGCAGGGCUUGAGAGGCACCUGGAGGGUGCCCUGGAGCAUUGGGAAGGUC

1     (((.......(((((.((((((((....(((.(((...((((..(((.((...((......((((((....((((.......................((((((........))))))........................))))......))))))..
160   ....)).)).))).))))....))).)))....)))))))))))))........)))
```

You can download the minimum free energy (MFE) structure in [Vienna Format| Ct Format]. You can get thermodynamic details on this structure by submitting to our RNAeval web server.

##### Graphical output

You may look at the interactive drawing of the MFE structure below. If you do not see the interactive drawing and you are using Internet Explorer, please install the Adobe SVG plugin. **A note on base-pairing probabilities:** The structure below is colored by base-pairing probabilities. For unpaired regions the color denotes the probability of being unpaired.

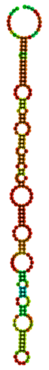

Sequence display options
- Plain Sequence
- No Sequence

**MFE secondary structure**

Other display options
- Base-pair probabilities
- Positional entropy
- None

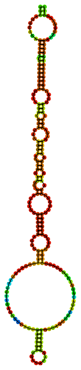

Sequence display options
- Plain Sequence
- No Sequence

**Centroid secondary structure**

Other display options
- Base-pair probabilities
- Positional entropy
- None

| **Image description** | **Download options** |
| --- | --- |
| MFE plain structure drawing | [ EPS \| PDF \| IMAGE CONVERTER \| VIEW IN FORNA ] |
| Centroid plain structure drawing | [ EPS \| PDF \| IMAGE CONVERTER \| VIEW IN FORNA ] |
| MFE structure drawing encoding base-pair probabilities | [ EPS \| PDF \| IMAGE CONVERTER \| VIEW IN FORNA ] |
| Centroid structure drawing encoding base-pair probabilities | [ EPS \| PDF \| IMAGE CONVERTER \| VIEW IN FORNA ] |
| MFE structure drawing encoding positional entropy | [ EPS \| PDF \| IMAGE CONVERTER \| VIEW IN FORNA ] |
| Centroid structure drawing encoding positional entropy | [ EPS \| PDF \| IMAGE CONVERTER \| VIEW IN FORNA ] |

Here you find a mountain plot representation of the MFE structure, the thermodynamic ensemble of RNA structures, and the centroid structure. Additionally we present the positional entropy for each position. Download as [EPS|PDF|IMAGE CONVERTER].

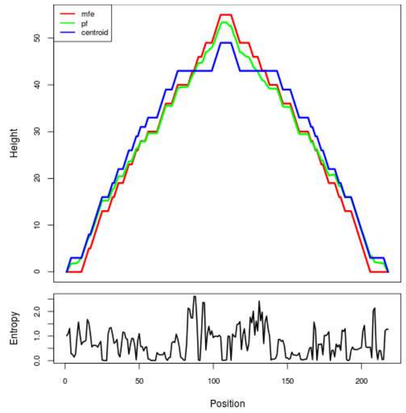


Results have been computed using RNAfold 2.6.3. An equivalent command line call would have been

```
RNAfold -p -d2 --noLP < sequence1.fa > sequence1.out
```

**RNA parameters are described in**

Mathews DH, Disney MD, Childs JL, Schroeder SJ, Zuker M, Turner DH. (2004) Incorporating chemical modification constraints into a dynamic programming algorithm for prediction of RNA secondary structure. *Proc Natl Acad Sci U S A* 101(19):7287-92.

**If you find these results helpful for your work you may want to cite:**

OPEN ACCESS OXFORD JOURNALS Gruber AR, Lorenz R, Bernhart SH, Neuböck R, Hofacker IL. **The Vienna RNA Websuite.** Nucleic Acids Research, Volume 36, Issue suppl_2, 1 July 2008, Pages W70-W74, DOI: 10.1093/nar/gkn188

Lorenz, R. and Bernhart, S.H. and Höner zu Siederdissen, C. and Tafer, H. and Flamm, C. and Stadler, P.F. and Hofacker, I.L. "ViennaRNA Package 2.0", Algorithms for Molecular Biology, 6:1 page(s): 26, 2011

### RNAfold WebServer MT_PURE_SEQ13

1 Enter Input Parameters 2 View Results

[Home|New job|Help]

#### Results for minimum free energy prediction

The optimal secondary structure in dot-bracket notation with a minimum free energy of **-8.14** kcal/mol is given below.

[color by base-pairing probability | color by positional entropy | no coloring]

```
1      CCACCUCCCUCUCCCUCGUUUCCUCCCCGCUCGCUCCUCCCUCCCUCGCUCCCUCGCUCCCUCCCCGCUCGCUCCUCCCUCCCUCGCCCGCUCGUCCCCGCUCCCUCCUCCCUCUCCCUCAGUCAUUGCCUCGCUCCCUCACCCACUAGUCUCCUCCUUC
160    UCCUCUCCCUCCCUCUCCCUCCUCCUUCUCCCUAGUUUCCUCCUUGCUUGGUGCU

1      ..............................((.............((......)).........((..((.............))..))..................................(((..((..........(((((..........
160    ..............................))))).........))..))))).
```

You can download the minimum free energy (MFE) structure in [Vienna Format| Ct Format]. You can get thermodynamic details on this structure by submitting to our RNAeval web server.

#### Results for thermodynamic ensemble prediction

The free energy of the thermodynamic ensemble is **-10.61** kcal/mol.
The frequency of the MFE structure in the ensemble is **1.81** %.
The ensemble diversity is **17.02** .
You may look at the **dot plot** containing the base pair probabilities [EPS|PDF|IMAGE CONVERTER].

The centroid secondary structure in dot-bracket notation with a minimum free energy of **-5.64** kcal/mol is given below.

[color by base-pairing probability | color by positional entropy | no coloring]

```
1      CCACCUCCCUCUCCCUCGUUUCCUCCCCGCUCGCUCCUCCCUCCCUCGCUCCCUCGCUCCCUCCCCGCUCGCUCCUCCCUCCCUCGCCCGCUCGUCCCCGCUCCCUCCUCCCUCUCCCUCAGUCAUUGCCUCGCUCCCUCACCCACUAGUCUCCUCCUUC
160    UCCUCUCCCUCCCUCUCCCUCCUCCUUCUCCCUAGUUUCCUCCUUGCUUGGUGCU

1      ..............................................((......))............((.............))....................................(((..((..........(((((..........
160    ..............................))))).........))..)))...
```

You can download the minimum free energy (MFE) structure in [Vienna Format| Ct Format]. You can get thermodynamic details on this structure by submitting to our RNAeval web server.

#### Graphical output

You may look at the interactive drawing of the MFE structure below. If you do not see the interactive drawing and you are using Internet Explorer, please install the Adobe SVG plugin. **A note on base-pairing probabilities:** The structure below is colored by base-pairing probabilities. For unpaired regions the color denotes the probability of being unpaired.

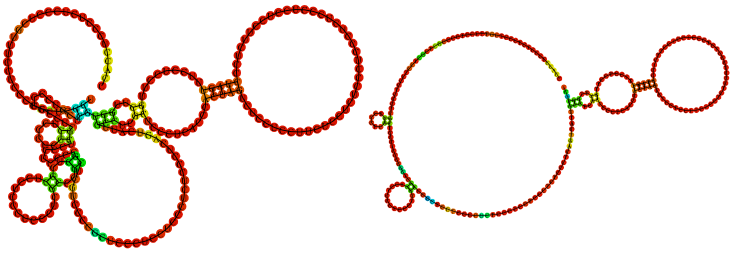

Sequence display options
- Plain Sequence
- No Sequence

MFE secondary structure

Other display options
- Base-pair probabilities
- Positional entropy
- None

0 1

Sequence display options
- Plain Sequence
- No Sequence

Centroid secondary structure

Other display options
- Base-pair probabilities
- Positional entropy
- None

0 1

| Image description | Download options |
|---|---|
| MFE plain structure drawing | [ EPS \| PDF \| IMAGE CONVERTER \| VIEW IN FORNA ] |
| Centroid plain structure drawing | [ EPS \| PDF \| IMAGE CONVERTER \| VIEW IN FORNA ] |
| MFE structure drawing encoding base-pair probabilities | [ EPS \| PDF \| IMAGE CONVERTER \| VIEW IN FORNA ] |
| Centroid structure drawing encoding base-pair probabilities | [ EPS \| PDF \| IMAGE CONVERTER \| VIEW IN FORNA ] |
| MFE structure drawing encoding positional entropy | [ EPS \| PDF \| IMAGE CONVERTER \| VIEW IN FORNA ] |
| Centroid structure drawing encoding positional entropy | [ EPS \| PDF \| IMAGE CONVERTER \| VIEW IN FORNA ] |

Here you find a mountain plot representation of the MFE structure, the thermodynamic ensemble of RNA structures, and the centroid structure. Additionally we present the positional entropy for each position. Download as [EPS|PDF|IMAGE CONVERTER].

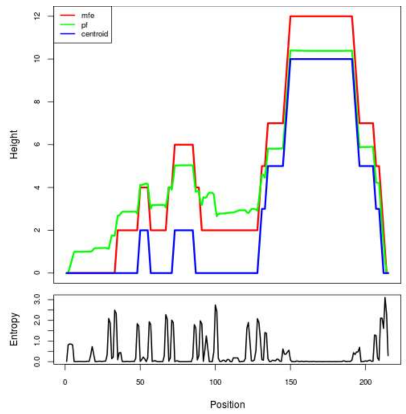


Results have been computed using RNAfold 2.6.3. An equivalent command line call would have been

```
RNAfold -p -d2 --noLP < sequence1.fa > sequence1.out
```

**RNA parameters are described in**

Mathews DH, Disney MD, Childs JL, Schroeder SJ, Zuker M, Turner DH. (2004) Incorporating chemical modification constraints into a dynamic programming algorithm for prediction of RNA secondary structure. *Proc Natl Acad Sci U S A* 101(19):7287-92.

**If you find these results helpful for your work you may want to cite:**


OPEN ACCESS OXFORD JOURNALS Gruber AR, Lorenz R, Bernhart SH, Neuböck R, Hofacker IL. **The Vienna RNA Websuite.** Nucleic Acids Research, Volume 36, Issue suppl_2, 1 July 2008, Pages W70-W74, DOI: 10.1093/nar/gkn188

Lorenz, R. and Bernhart, S.H. and Höner zu Siederdissen, C. and Tafer, H. and Flamm, C. and Stadler, P.F. and Hofacker, I.L. "ViennaRNA Package 2.0", Algorithms for Molecular Biology, 6:1 page(s): 26, 2011
